# Metabolic regulation of RNA methylation by the m^6^A-reader IGF2BP3

**DOI:** 10.1101/2024.10.31.621399

**Authors:** Gunjan Sharma, Martin Gutierrez, Anthony E Jones, Amit Kumar Jaiswal, Zachary T Neeb, Amy Rios, Michelle L Thaxton, Tasha L Lin, Tiffany M Tran, Lyna E S Kabbani, Alexander J Ritter, Linsey Stiles, Johanna ten Hoeve, Ajit S Divakaruni, Jeremy R Sanford, Dinesh S Rao

**Affiliations:** Department of Pathology and Laboratory Medicine, University of California, Los Angeles, Los Angeles, CA; Department of Molecular, Cell and Developmental Biology and Center for Molecular Biology of RNA, University of California Santa Cruz, Santa Cruz, CA; Department of Molecular and Medical Pharmacology, University of California, Los Angeles, Los Angeles, CA, United States; Division of Hematology and Oncology, Department of Medicine, University of California, Los Angeles, Los Angeles, CA; Department of Medicine, University of California, Los Angeles, Los Angeles, CA, United States; UCLA Metabolomics Center, University of California, Los Angeles, CA, 90095, USA; Center for Biomolecular Science & Engineering, University of California Santa Cruz, Santa Cruz, CA; Jonsson Comprehensive Cancer Center, University of California, Los Angeles, Los Angeles, CA; Broad Stem Cell Research Center, University of California, Los Angeles, Los Angeles, CA

## Abstract

The interplay of RNA modifications – deposited by “writers”, removed by “erasers” and identified by RNA binding proteins known as “readers” – forms the basis of the epitranscriptomic gene regulation hypothesis. Recent studies have identified the oncofetal RNA-binding protein IGF2BP3 as a “reader” of the N6-methyladenosine (m^6^A) modification and crucial for regulating gene expression. Yet, how its function as a reader overlaps with its critical oncogenic function in leukemia remains an open question. Here, we report the novel finding that the reader IGF2BP3 reprograms cellular metabolism, resulting in an altered ability of the “writers” to modify the epitranscriptome. In leukemia cells, IGF2BP3 supports increased glycolytic flux and one-carbon metabolism, leading to increased production of S-adenosyl methionine (SAM), a key substrate for methylation reactions within the cell. IGF2BP3 directly regulates the translation of MAT2B, the regulatory subunit of the methionine-adenosyltransferase complex, which is the final enzyme in a pathway leading to SAM production. This, in turn, results in increased m^6^A modifications on RNA, resulting in positive feedback regulation. This novel mechanism illustrates how metabolism mutually acts with epitranscriptomic modifications, underscoring the pervasive impact of IGF2BP3 in gene regulatory mechanisms governing a broad range of cancer-specific processes.

## INTRODUCTION

Methylation, the addition of a methyl group (CH₃) to DNA, RNA, or proteins, has broad and important effects on gene expression, protein function, and cellular signaling. Although the existence of the N6-methyladenosine (m^6^A) RNA modification has been known since 1974, more recent work has revealed key regulatory roles for this RNA modification^1–3^. The discovery of RNA methylases, followed by the identification of RNA demethylases and methylation-sensitive RNA binding proteins, referred to as writers, erasers and readers, forms the basis of the epitranscriptome hypothesis, which posits that RNA modification contributes to gene expression regulation^4^. One of the most striking features of m^6^A methylation is its predominant localization within 3’ UTRs near the stop codon^5,6^. This localization of m^6^A modifications, overlapping with RNA-binding proteins (RBPs) and microRNA binding sites may underlie its reported function in a host of RNA homeostatic processes^7^.

Insulin-like Growth Factor 2 mRNA Binding Protein 3 (IGF2BP3) is an oncofetal RBP which regulates mRNA localization, stability, and translation^8^. The recent discovery that IGF2BP3 is an m^6^A reader is consistent with its binding preference for 3’UTRs. IGF2BP3 regulates mRNA targets enriched for genes important in various aspects of oncogenesis and differentiation^9–12^. Prior work from our group and others established IGF2BP3 as a critical regulator of leukemogenesis in *MLL*-translocated B-acute lymphoblastic leukemia^10,11^. Recently, IGF2BP3 has also been shown to regulate lipid and other metabolic pathways in epithelial cancer cells^13,14^. In this rapidly progressing field, IGF2BP3 is thought to regulate the stability of a number of coding and non-coding RNAs, which then directly or indirectly impact enzymes regulating various metabolic pathways.

The generation of RNA methylation is dependent on the presence of S-adenosyl methionine (SAM), which is the key methyl donor for most cellular methylation reactions. The one-carbon metabolism (OCM) pathway plays a crucial role in generating methyl donors required for DNA synthesis and DNA/RNA methylation reactions^15^, in addition to other critical cancer-specific processes. Dysregulation of OCM has been implicated in various cancers, including leukemia, and has emerged as an important regulator of leukemic stem cell (LSC) function^16–18^. Glycine and serine, two key OCM metabolites, are known to play a key role in oncogenesis^19,20^. In T-cell acute lymphoblastic leukemia, serine hydroxy methyltransferases (SHMT1/2) were discovered to have vulnerabilities and a valuable drug target^21^. Together, these findings point towards an intriguing, yet not fully understood, role between OCM, cancer, and the availability of substrates for cellular methylation reactions.

In our efforts to understand the effects of IGF2BP3 on leukemia cell metabolism, we uncovered an impact on glycolysis and OCM. Metabolic profiling analyses revealed a deficit in the glycolytic metabolites pyruvate and lactate as well as those linked to one-carbon metabolism including serine, glycine, S-adenosylmethionine (SAM), and cystathionine. By adopting a combined high through-put analysis approach using RNA binding data from enhanced cross-linking immunoprecipitation (eCLIP) with IGF2BP3 and the metabolomics data, we identified several direct targets of IGF2BP3 in the Glycine-Serine cycle and the Methyl/Folate cycle. Targeted western blot analysis showed that IGF2BP3-deficient cells had reduced levels of several metabolic regulators, including MAT2A and MAT2B, the rate-limiting enzyme in the production of SAM. Critically, we found that IGF2BP3 depletion consistently led to decreased overall m^6^A modifications in a range of cell lines and systems, including genetic knockout and small molecule inhibition, while exogenous expression of IGF2BP3 rescued the metabolic-epitranscriptomic phenotype. Our work uncovers the unexpected phenomenon of how an m^6^A reader, IGF2BP3, modulates its affinity to its target mRNAs by driving changes in RNA methylation, thereby generating a pervasive shift in gene expression, and maintaining a cancerous phenotype.

## RESULTS

### IGF2BP3 promotes glycolysis in leukemic cells

Given prior reports of IGF2BP family proteins impacting metabolism^22^ and our observations of an important role of IGF2BP3 in acute leukemia, we undertook metabolic profiling experiments to understand the role of this protein in regulating cancer cell metabolism. Seahorse XF analysis showed that the depletion of IGF2BP3 decreased cellular lactate efflux, as calculated from standard Seahorse XF parameters^23^, using independent CRISPR/Cas9 strategies in SEM cells^12^ (Figs. 1A-B, 1C *left*). Reductions in the lactate efflux rate were also seen in both NALM6 cells as well as in murine bone marrow cells transduced with MLL-Af4 depleted of IGF2BP3 (Fig. 1C, *center* and *right*). For an orthogonal measurement of glycolysis to confirm our findings, we conducted gas chromatography/mass spectrometric (GC/MS) analysis. Consistent with previous findings, we observed a reduction in steady-state levels of pyruvate and lactate (Fig. 1D and 1E), as well as reduced enrichment from uniformly labelled ^13^C_6_ glucose into these metabolites (Fig. 1E). To discriminate between a specific, targeted reduction in glycolytic flux or a global decrease in cellular energy demand and metabolic rate, we conducted respirometry and stable isotope tracing to identify potential changes in oxidative phosphorylation and mitochondrial function. Importantly, we did not observe reproducible changes in any oxygen consumption rate (OCR) parameters or enrichment from uniformly labelled ^13^C_6_-glucose or ^13^C_5_-glutamine into the TCA cycle, and steady-state abundances of TCA cycle intermediates were mostly unchanged (Suppl. Fig. 1). Altogether, our initial characterization of metabolism in Fig. 1 indicates that IGF2BP3 uniquely supports glycolytic flux without an appreciable effect on oxidative phosphorylation.

**Figure 1.**
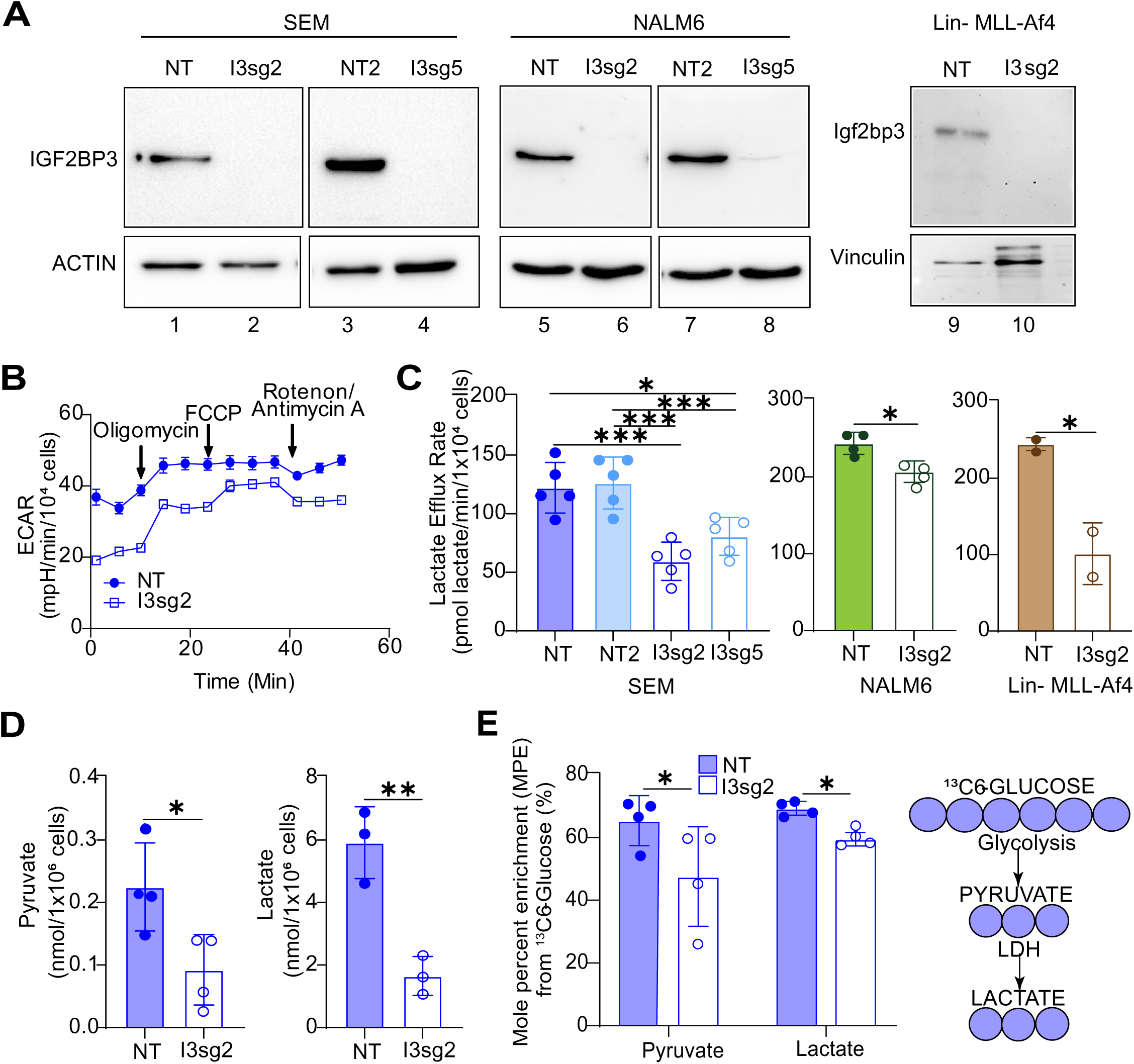
IGF2BP3 impacts glycolytic metabolism in B-acute lymphoblastic leukemia cells. **A.** Western blots for IGF2BP3-deleted (I3sg2, I3sg5) in SEM, NALM6 and Lin-/MLL-Af4 murine cells. **B.** Seahorse XF Extracellular acidification rate (ECAR) kinetic trace in control and IGF2BP3-deleted SEM cells (I3sg2). **C.** Aggregate lactate efflux rates from Seahorse XF Analysis in control (NT, NT2) versus IGF2BP3-deleted (I3sg2, I3sg5) in SEM, NALM6 and Lin-/MLL-Af4 murine cells. **D.** Pyruvate and Lactate amounts measured by GC-MS in control versus IGF2BP3-deleted (I3sg2) SEM cells **E.** Incorporation of carbon from ^13^C-labeled glucose into pyruvate and lactate, measured as mole percent enrichment (MPE) from GC-MS experiments. All data are n>3 biological replicates. *, p<0.05; **, p<0.01; ***, p<0.001.

### IGF2BP3 supports one-carbon metabolism and the generation of S-adenosyl-methionine (SAM)

Given our initial findings of alterations in glycolytic metabolism in IGF2BP3-depleted cells, we undertook a liquid chromatography/mass spectrometric (LC/MS) analysis of metabolites in SEM cells to more thoroughly characterize changes in glycolysis-linked pathways beyond the information available from Seahorse XF and GC/MS analysis. These LC/MS studies revealed twenty-nine metabolites of central carbon metabolism that showed statistically significant changes in at least one of the two CRISPR knockout lines that were queried (Fig. 2A) (Supp. Table 1). Looking at the metabolites noted to be altered by both GC/MS and LC/MS, we found reductions in glycolytic metabolites (lactate and fructose-1,6-bisphosphate) (Suppl. Fig. 2A-B) as well as metabolites in the one-carbon and sulfur-containing amino acid pathways such as serine (Fig. 2C), glycine (Fig. 2E), glutathione (Fig. 2I) and cystathionine (Suppl. Fig. 2C). Furthermore, isotopologue distribution patterns from uniformly labelled glucose revealed these reduced levels were attributable to reduced synthesis. (Fig. 2D, 2F, 2J and Suppl. Fig. 2D, respectively). Importantly, there was a significant change in steady-state levels of S-adenosyl methionine (SAM), a product of the methionine cycle that derives some carbons from flux through the Serine-Glycine pathway (Fig. 2G). The incorporation of ^13^C_6_-glucose into SAM was also reduced, again indicating that decreased flux through the Serine-Glycine pathway may be at least partially responsible for the reduced SAM levels (Fig. 2H). Together, our findings indicate that IGF2BP3 supports the generation of SAM, which is the key methyl donor for a number of methylation reactions in cells.

**Figure 2.**
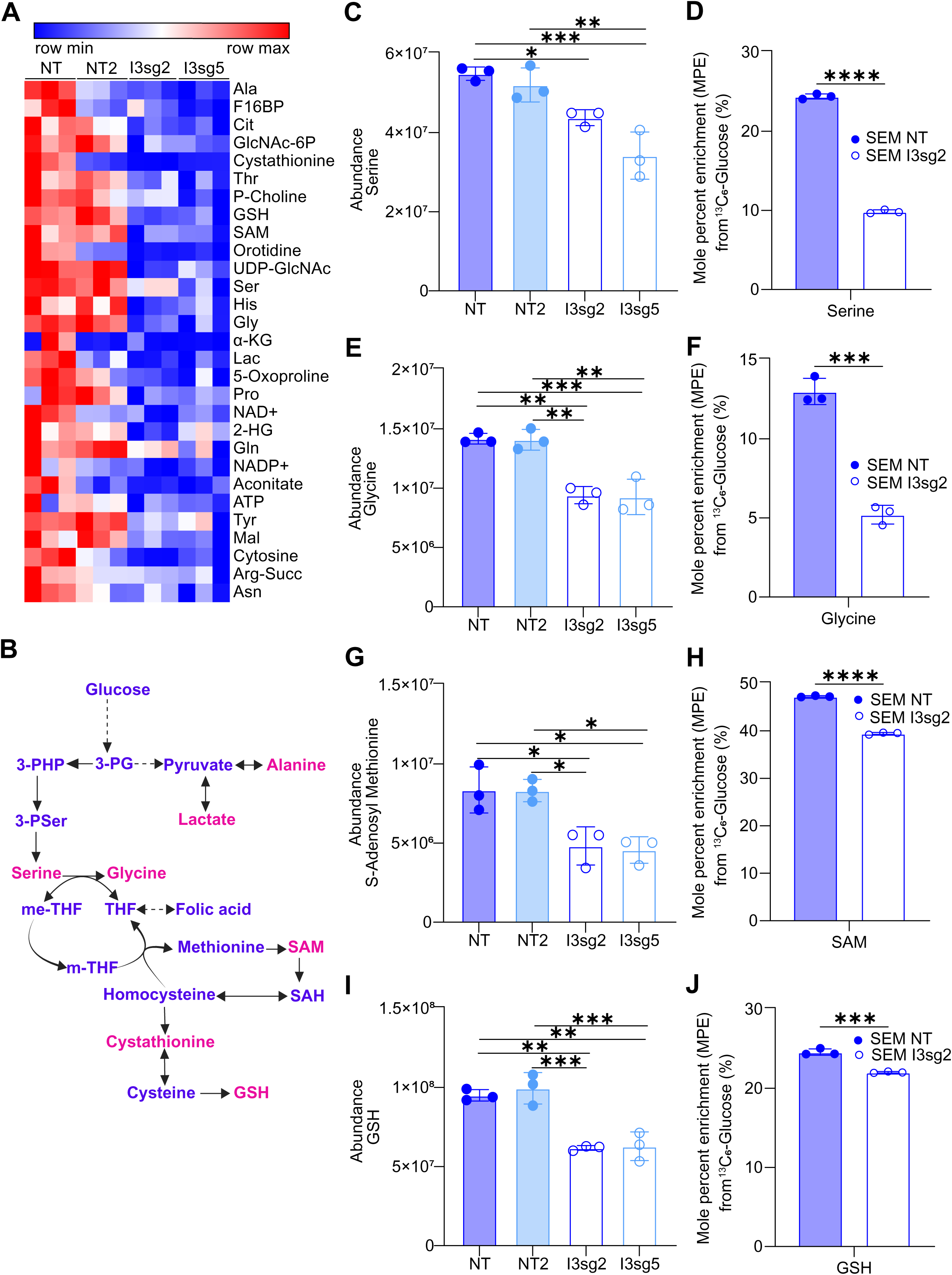
IGF2BP3 supports one-carbon metabolism pathways that serve as methyl donors. **A.** Heatmap depicting significantly altered metabolites from control versus IGF2BP3-deleted SEM cells, as indicated, using targeted analysis of polar central carbon metabolites by LC-MS. Shown are metabolites with a consistent change in both IGF2BP3-deleted lines. **B.** Schematic of metabolites that are produced in one-carbon metabolism. **C-J.** Intracellular abundance and steady-state incorporation of carbon from ^13^C-labeled glucose, measured as mole percent enrichment (MPE), into one-carbon pathway metabolites serine, glycine, S-adenosyl-methionine (SAM) and glutathione (GSH) in control versus IGF2BP3-deleted SEM cells. All data are n>3 biological replicates. *, p<0.05; **, p<0.01; ***, p<0.001.

### IGF2BP3 promotes m^6^A modifications on RNA

Because SAM serves as a methyl donor for a variety metabolic and gene regulatory processes, we hypothesized that IGF2BP3 depletion may impact protein and nucleic acid methylation. Given prior reports that SAM levels impact histone methylation^24^, we examined H3K4me1 and H3K4me3 marks in our model systems. Indeed, both methylation marks were reduced in SEM cells (two distinct sgRNAs) and in Lin-MLL-Af4 cells that had been depleted for IGF2BP3 (Fig. 3A). With this finding in hand, we next examined m^6^A marks on RNA using an ELISA-based assay. Strikingly, IGF2BP3 deletion reduced the relative m^6^A levels in both SEM and Lin-MLL-AF4 cells (Fig. 3B). Similar findings were also observed in NALM6 cells that had been depleted of IGF2BP3 (Fig. 3C), where we had also noted the reduction in glycolytic flux. To confirm these observations, we performed dot blots on total RNA purified from control or IGF2BP3-depleted cells. Staining with the m^6^A antibody (see Materials & Methods) was significantly reduced in IGF2BP3-depleted cells relative to control, despite equivalent levels of RNA in each sample as visualized by methylene blue staining (Fig. 3D). Importantly, this change in m^6^A levels was not accompanied by a change in either RNA methylase or demethylase activity within the cells (Fig. 3E-F). Similarly, the protein levels of METTL3, METTL14, METTL16, and FTO, key m^6^A writers and erasers were unchanged as a function of IGF2BP3 (Fig. 3H).

**Figure 3.**
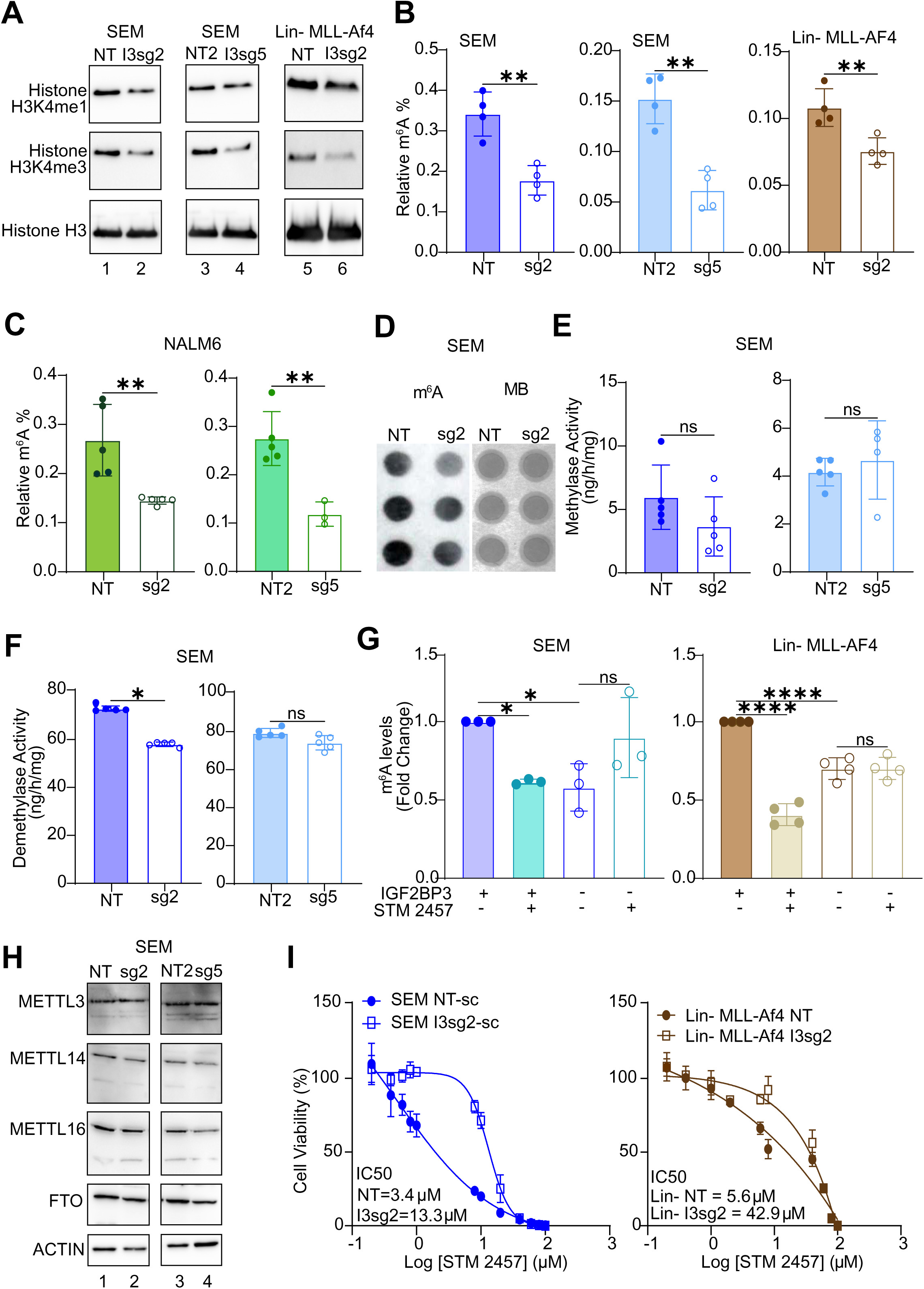
IGF2BP3 regulates N6-methyladenosine marks in RNA. **A.** Western blot analysis of histone methylation (H3K4me1 and H3K4me4) in SEM and Lin-MLL-Af4 cells, control or deficient for IGF2BP3. **B.** ELISA measurement of m^6^A modification on RNA isolated from SEM and Lin-MLL-Af4 cells as above. **C.** ELISA measurement of m^6^A modification on RNA isolated from NALM6 cells, control or deficient for IGF2BP3. **D.** Dot blot analysis of m^6^A modification (left) and methylene blue staining in SEM cells, control or deficient for IGF2BP3. **E.** RNA m^6^A methylase activity (colorimetric assay, expressed as enzymatic activity) in SEM cells, control or deficient for IGF2BP3. **F.** RNA m^6^A demethylase activity (colorimetric assay, expressed as enzymatic activity) in SEM cells, control or deficient for IGF2BP3. **G.** ELISA measurement of m^6^A modification on RNA isolated from control or IGF2BP3-deficient SEM and Lin-MLL-Af4 cells, following treatment with METTL3 inhibitor STM2457 at 5 µM concentration. **H.** Western blot analysis of RNA m^6^A-methylase and demethylase enzymes in SEM cells, control or deficient for IGF2BP3. **I.** Cell viability assays (Cell Titer Glo) on control versus IGF2BP3 deleted SEM (left) and Lin-MLL-Af4 cells (right) cells treated with STM2457. Cells were grown for 3 days in the presence of inhibitor prior to measurement of cell viability.

Given the known role of IGF2BP3 as an m^6^A reader, we then queried whether upstream inhibition of METTL3, the catalytic unit of the RNA m^6^A writer enzyme, by STM2457 would show an additional effect on m^6^A levels. Remarkably, there was no further reduction in m^6^A levels by the addition of STM2457, and IC50 curves demonstrated that IGF2BP3-depleted cells were more resistant to STM2457-based cell growth inhibition in both cell types that were tested (Fig. 3G and 3I, respectively). The effect on m6A levels following treatment with PF-9366, an allosteric inhibitor of MAT2A^25^, was similarly attenuated in the IGF2BP3-depleted cells (data not shown). These data suggest that IGF2BP3 directly impacts the production of SAM through the MAT2A/B complex, a phenotype that was not additive to the loss of METTL3 or MAT2A.

### IGF2BP3 regulation of metabolic genes involves specific translational control

Our prior work^11^ demonstrated that IGF2BP3 supports leukemogenesis. A re-analysis of differentially regulated gene sets from those studies revealed that there was also an enrichment of gene sets related to metabolism (Supp Fig. 3A-D). We, therefore, used Metaboanalyst ^26^ to analyze deregulated metabolites identified in the LC/MS analysis (Fig. 2). This revealed highly statistically significant enrichment for metabolites related to glycine, serine and threonine metabolism, aminoacyl tRNA biosynthesis and cysteine and methionine metabolism (Fig. 4A). Lesser enrichments were seen for glutathione metabolism as well as other pathways. Next, we utilized recently generated eCLIP data for IGF2BP3 in SEM cells to identify target mRNAs (manuscript in preparation; Fig. 4B). Indeed, we found genes related to these same pathways as IGF2BP3 targets (Fig. 4B). Next, we queried whether there were concordant alterations in gene expression based on differential expression analysis by RNA-sequencing. A majority of the genes did not show changes at the RNA level, implying either non-functional protein-RNA interactions or a mechanism that does not rely on changes in RNA levels (Fig. 4C). Because of the relatively small number of altered metabolites, we pursued an alternative strategy by analyzing the expression of key regulators of glycolysis and one-carbon metabolism by western blot analysis. We found that there were small but consistent changes in the expression of proteins related to glycolysis, serine/glycine biosynthesis, one-carbon cycle, and methyl cycle (Fig. 4D). Of these, the change in MAT2A was most concordant with a functional role in the observed metabolic changes, particularly the decrease in SAM. Broadly, the same changes in protein expression were also observed in the Lin-murine system (Supp. Fig. 3E).

**Figure 4.**
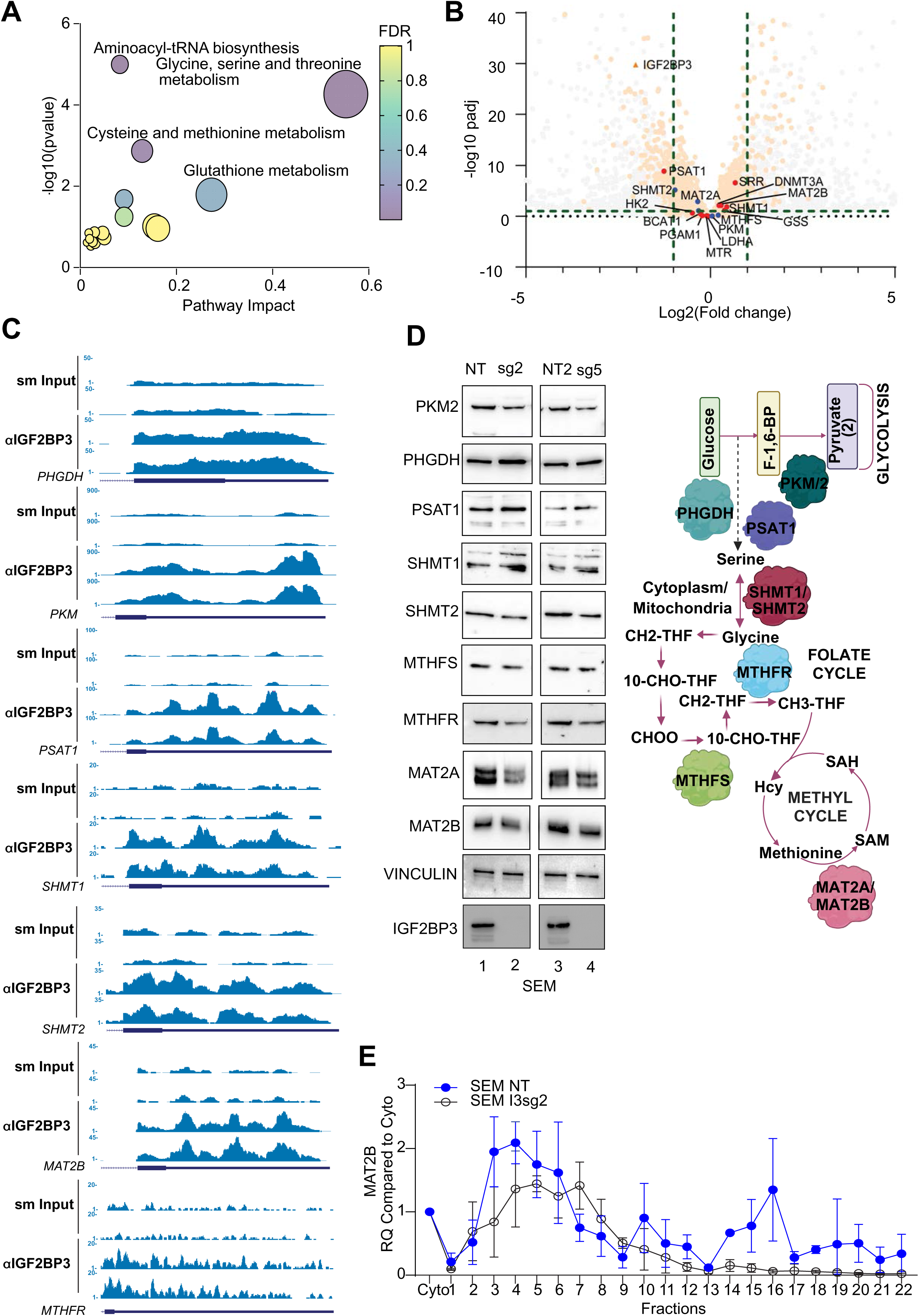
IGF2BP3 regulates translation of metabolic genes. **A.** MetaboAnalyst-based pathway enrichment analysis of consistently differentially regulated metabolites in SEM cells with knockout of IGF2BP3. **B.** Volcano plot showing differentially expressed genes and IGF2BP3 targets defined by eCLIP analysis (dots exceeding the thresholds depicted by dashed lines), Putative IGF2BP3 targets which were differentially expressed are highlighted as transparent orange, IGF2BP3 metabolic targets which were identified using Skipper (see ref. ^65^) are highlighted in red, while metabolic targets that were not IGF2BP3 are in blue. Grey dots are not IGF2BP3 targets. Green dashed lines mark the significant cutoffs for diff. expression (−1/1) and sig pvalue (1). **C.** Genome browser snapshots of eCLIP read coverage across some putative IGF2BP3 target genes. Depicted are the genes with key roles in glycolysis and one-carbon metabolism and map to the enriched terms in (A). **D.** Western Blot analysis of key genes in metabolic pathways (left) and simplified schematic depiction of genes that control metabolic pathways altered in IGF2BP3-depleted cells. **E.** 10-45% Sucrose gradient fractionation of cytosolic extracts from control or IGF2BP3-depleted SEM cells. MAT2B mRNA distribution was measured by RT-qPCR.

Given the surprising observation that IGF2BP3 deletion only subtly alters steady-state levels of metabolic transcripts, we explored the possibility of IGF2BP3-dependent translational control. Importantly, global translation was not altered per SUnSET assay^27^, where puromycin incorporation into elongating protein strands is assessed (Supp. Fig 4A). To query IGF2BP3-related mechanisms of translational regulation, we profiled polyribosome association of mRNA transcripts from control or IGF2BP3-depleted SEM cytosolic extracts using sucrose gradient centrifugation and RT-qPCR. Out of the 7 candidate transcripts identified from the eCLIP analyses (Fig. 4C), we found that polyribosome association on MAT2B mRNA, which encodes the regulatory subunit of the MAT2 complex^28^, was strongly reduced in IGF2BP3-depleted cells (Fig. 4E). Western blot analysis also revealed a reduction of steady-state protein levels in the IGF2BP3-depleted cells (Fig. 4D). A similar result was obtained for the PKM gene, whereas PSAT1, SHMT1 and MTHFR transcript distribution was not significantly altered across gradients from control or IGF2BP3-depleted cells (Supp. Fig. 3F-I). We confirmed that ribosome integrity was required for the observed changes, as lysates treated with EDTA, which causes ribosomal subunit dissociation, attenuates transcript sedimentation (data not shown). Interestingly, MAT2A, the catalytic subunit of the MAT2 complex, did not show any changes in polyribosome association despite showing an alteration in protein levels (Supp. Fig. 4B). Given this lack of change in translation, we next queried protein stability, using a Cycloheximide Chase Assay^29^, finding that protein levels of MAT2A showed declining levels in IGF2BP3-deficient cells, but not so in IGF2BP3-expressing cells (Supp. Fig 4B). These findings are consistent with reduced stability of the MAT2A enzyme in the absence of the MAT2B subunit, as previously reported^28^. Taken together with our previous data indicating unchanged steady-state protein levels of canonical writers and erasers (Fig. 3H, Suppl. Fig. 3E), we suggest that altered translation of MAT2B drives changes in MAT2A protein stability, resulting in a concomitant decrease in SAM and m^6^A in IGF2BP3-depleted cells.

### IGF2BP3 promotes the metabolic-epitranscriptomic axis *in vivo*

To further query the significance of our observations relating IGF2BP3 to metabolism and the epitranscriptome, we set out to understand if the changes we observed were also noted after the loss of function *in vivo*. First, we examined mouse bone marrow from mice that had been transplanted with Lin-MLL-Af4 cells as previously described^12^, finding that m^6^A levels were reduced with depletion of IGF2BP3 (Fig. 5A). Next, we turned to our recently developed small molecule inhibitor of IGF2BP3, I3IN-002, that inhibits cell growth with an IC50 of 2-3.5 uM in SEM cells (Fig, 5B-C; manuscript under consideration). Similar to genetic depletion, I3IN-002 reduced ECAR and lactate efflux levels in SEM cells that were treated with 5 uM I3IN-002 (Fig. 5D-E). I3IN-002 resulted in a reduction of m^6^A levels, similar to treatment with STM2457, the previously mentioned METTL3 inhibitor (Fig. 5F). *In vivo*, treatment of murine Lin-MLL-Af4 tumors as well as a PDX model of B-ALL, PDX#22694^30^ resulted in statistically significant reductions of m^6^A levels in both cases (Fig. 5G-H). Together, these data support the idea that IGF2BP3 promotes altered metabolism and RNA modifications *in vivo*.

**Figure 5.**
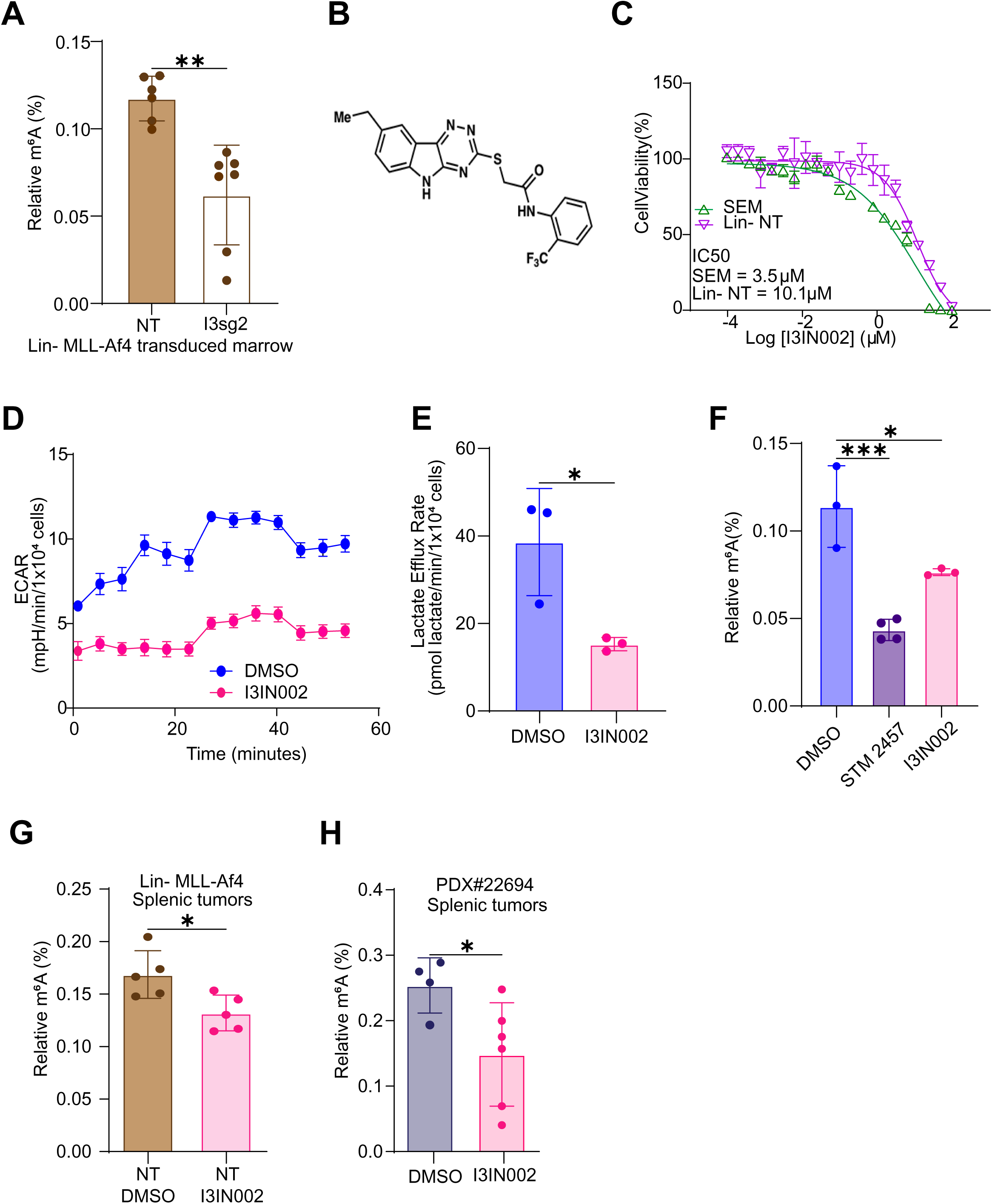
IGF2BP3 loss of function impacts glycolytic metabolism and m^6^A RNA modifications in vivo. **A.** ELISA Measurement of m^6^A modification from murine bone marrow isolated following transplantation with Lin-MLL-Af4 bone marrow (see ref.^12^) **B.** Chemical structure of I3IN-002 **C.** IC50 based on cell viability, measured by CellTiterGlo, in SEM and Lin-cells, following treatment with I3IN-002. **D.** Seahorse XF Extracellular acidification rate (ECAR) kinetic trace in SEM cells treated with vehicle or I3IN-002, a small molecule inhibitor of IGF2BP3. **E.** Aggregate lactate efflux rates from Seahorse XF Analysis in SEM cells treated with vehicle of I3IN-002. **F.** ELISA measurement of m^6^A RNA modifications in SEM cells treated with vehicle, STM2457, or I3IN-002. **G.** ELISA measurement of m^6^A RNA modifications in splenic tumors isolated from mice transplanted with Lin-MLL-Af4 cells, subsequently treated in vivo with I3IN-002. **H.** ELISA measurement of m^6^A RNA modifications in splenic tumors isolated from mice transplanted with human PDX B-ALL cells, subsequently treated in vivo with I3IN-002.

### Enforced expression of IGF2BP3 rescues the metabolic-epitranscriptomic phenotype

If IGF2BP3 depletion causes a reduction in SAM through regulation of the MAT2 complex, then we expect exogenous IGF2BP3 to rescue these phenotypes *in vitro*. To test this hypothesis, we re-introduced IGF2BP3 in SEM and Lin-MLL-Af4 cells in which IGF2BP3 had been deleted using CRISPR. Here, we utilized a codon-altered IGF2BP3 that retained the same amino acid sequence but had an altered nucleotide sequence to escape CRISPR/Cas9-mediated degradation. Western blotting confirmed that the codon-altered protein was efficiently expressed in both model systems (Fig. 6A). Following the re-expression of IGF2BP3 led to the rescue of PKM2 and MAT2A/MAT2B protein expression, cell growth and m^6^A modification of RNA in both SEM and Lin-MLL-Af4 cells (Fig. 6B-G, respectively). In an orthogonal set of assays, we turned to our previously described mouse model with germline deletion of *Igf2bp3* (*Igf2bp3^del/del^*)^11^. Lin-cells were collected from the bone marrow of these mice and transduced with MLL-Af4 as previously described, which in wild-type mice leads to overexpression of MLL-Af4 protein. Next, we used retroviral transduction to constitutively express wild-type IGF2BP3 in these cells and compared it with an empty-vector control (Fig. 6H). Constitutive exogenous expression of IGF2BP3 led to increased expression levels of MAT2A and MAT2B (Fig. 6H), consistent with the model of gene expression regulation presented previously. Re-expression of IGF2BP3 also led to increased cell growth, as measured by cell viability in Cell Titer Glo measurements (Fig. 6I). In terms of metabolic changes, we observed an increased extracellular acidification rate (ECAR; Fig. 6J) and increased lactate efflux rate (Fig. 6K). Colony formation in methylcellulose was increased with re-expression of IGF2BP3 and, importantly, m^6^A modifications in RNA were also increased (Fig. 6L-M).

**Figure 6.**
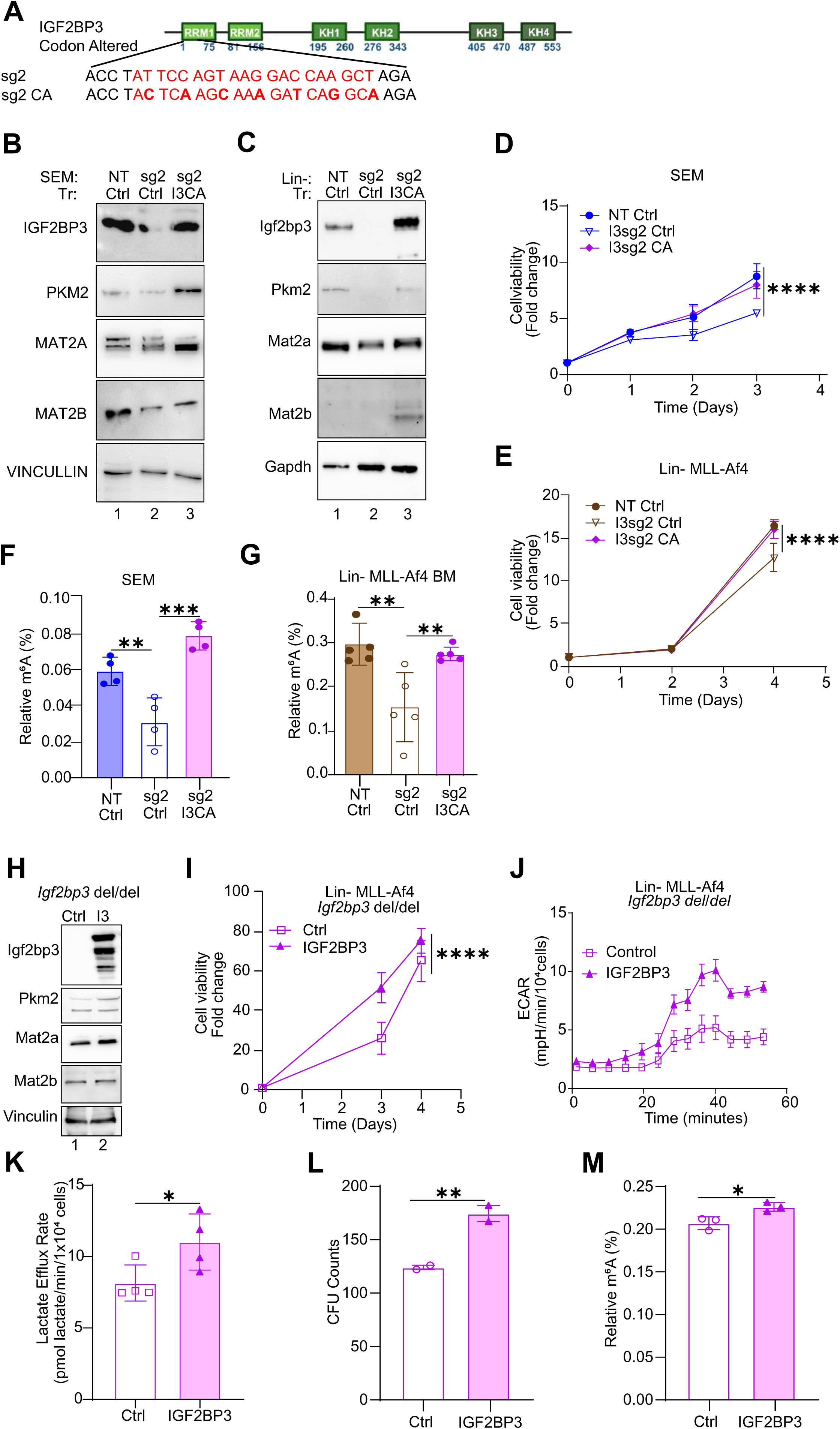
Re-expression of IGF2BP3 recovers metabolism, cell growth and RNA m^6^A modifications. **A.** MSCV-based construct showing bases altered to render it insensitive to sg2-mediated CRISPR/Cas9 activity (“codon-altered”, I3CA) **B.** Western blot analysis of enforced expression of IGF2BP3 in SEM cells that were previously deleted for IGF2BP3. NT/Ctrl, SEM cells sufficient for IGF2BP3, transduced with control vector; sg2/MIG, SEM cells deleted for IGF2BP3, transduced with control vector; sg2/I3CA, SEM cells deleted for IGF2BP3 then transduced with codon-altered IGF2BP3. Additionally, Western blot analysis for PKM2, MAT2A, MAT2B in SEM cells is shown. **C.** Western blot analysis of enforced expression of IGF2BP3 in Lin-/MLL-Af4 cells that were previously deleted for IGF2BP3. NT/Ctrl, cells sufficient for IGF2BP3, transduced with control vector; sg2/MIG, cells deleted for IGF2BP3, transduced with control vector; sg2/I3CA, cells deleted for IGF2BP3 then transduced with codon-altered IGF2BP3. Additionally, Western blot analysis for PKM2, MAT2A, MAT2B in SEM cells is shown. **D.** Cell growth curves measured by Cell Titer Glo, over three days in SEM cells, notated as in (B). **E.** Cell growth curves measured by Cell Titer Glo over three days in Lin-/MLL-Af4 cells, notated as in (C). **F.** ELISA measurement of m^6^A modification in RNA isolated from SEM cells notated as in (B). **G.** ELISA Measurement of m^6^A modification in RNA isolated from Lin-MLL-Af4 cells notated as in (E). **H.** Western blot analysis of Lin-cells from *Igf2bp3^del/del^* mice. Briefly, cells were isolated from mice with a germline deletion of Igf2bp3, transformed with MLL-Af4, and then subjected to transduction with MSCV-based constructs carrying the wild-type murine Igf2bp3. Proteins that were analyzed are: Igf2bp3, Myc, Mat2a, Mat2b and Actin. **I.** Cell growth, measured by Cell titer Glo, over 4 days, in *Igf2bp3^del/del^*Lin-MLL-Af4 cells with enforced IGF2BP3 expression as above. **J.** Seahorse XF Extracellular acidification rate (ECAR) kinetic trace in cells described above. **K.** Aggregate lactate efflux rates from Seahorse XF Analysis in cells described above. **L.** Colony formation assays from Lin-MLL-Af4 cells as described above. **M.** ELISA measurement of m^6^A modification on RNA isolated from *Igf2bp3^del/del^*Lin-MLL-Af4 cells with enforced IGF2BP3 expression as above.

To validate the findings *in vivo*, we utilized bone marrow transplant assays to evaluate the phenotype of MLL-Af4 *Igf2bp3* (*Igf2bp3^del/del^*) Lin-cells with and without enforced IGF2BP3 expression. Following transplantation of transduced cells, IGF2BP3 re-expressing mice showed a significant increase in engrafted cells, bone marrow counts, spleen weights and spleen counts at 6 weeks post-transplant compared to control mice (Fig. 7A-D). Consistent with our previous findings, IGF2BP3 re-expressing mice displayed significantly higher counts for CD11b+, Lineage-negative cells, LSK (Lin-ckit+Sca1-) cells, including potential leukemia-initiating cell (LIC) population^11,12^ in both bone marrow and spleen (Fig. 7E-I and Suppl. Fig. 5, respectively). Next, we queried the metabolic state of IGF2BP3 re-expressing and control mice using respirometry. Consistent with our other findings, IGF2BP3 re-expression increased ECAR and Lactate efflux rate (Figure 7J-K). Importantly, we also observed an increase in the m^6^A modifications on RNA in the IGF2BP3 re-expression group compared to the control group (Figure 7L). Together, these findings confirm that IGF2BP3 regulates cell growth, metabolic flux through the glycolytic pathway, expression of key regulators of SAM synthesis, and the m^6^A modification in mRNA.

**Figure 7.**
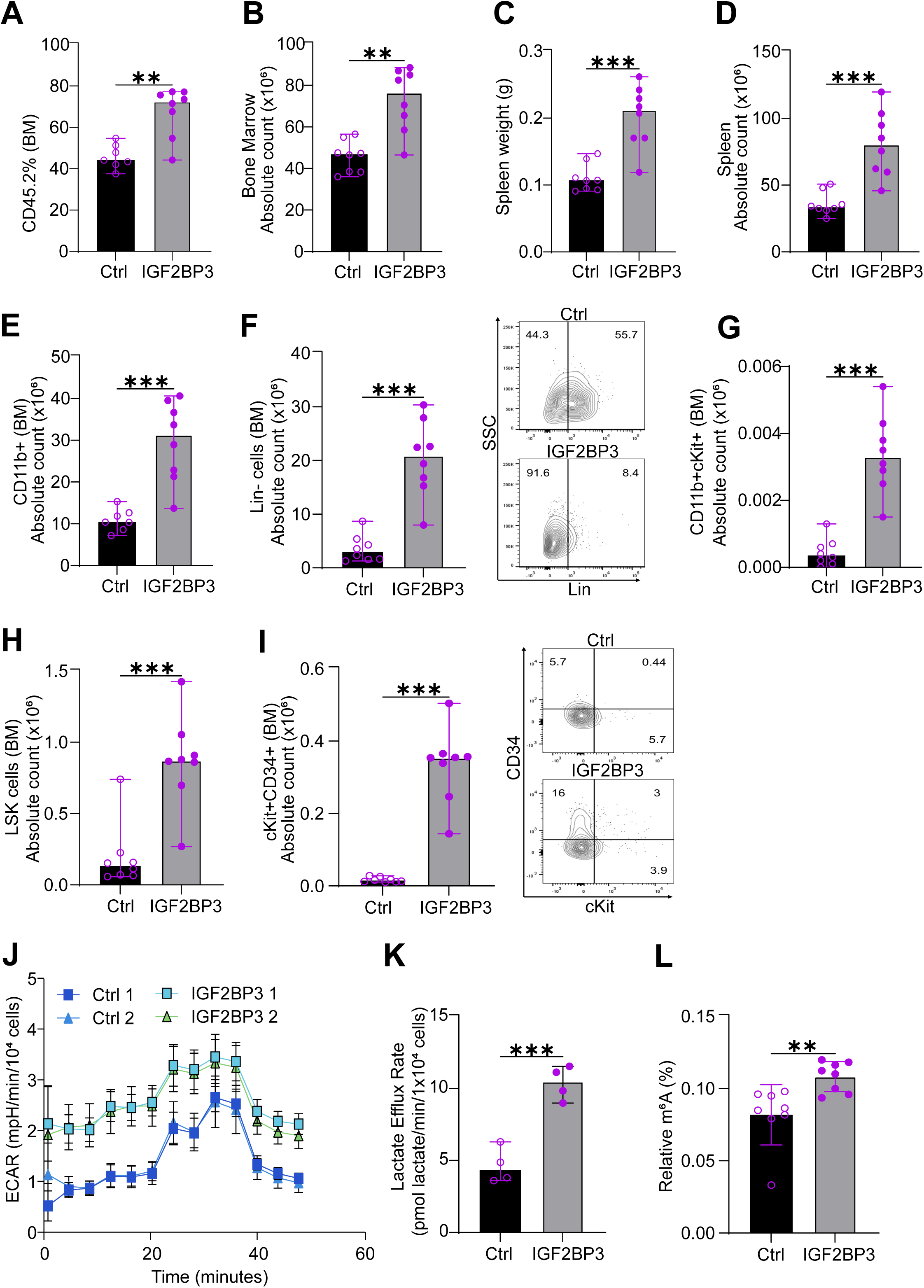
IGF2BP3 promotes glycolytic metabolism and m^6^A RNA modifications in vivo. **A.** Percentage engraftment of CD45.2 Lin-cells in bone marrow from *Igf2bp3^del/del^* mice transduced with MLL-Af4 re-expressing empty vector (Ctrl) or IGF2BP3 in the two groups at 6 weeks. **B.** Quantitation of bone marrow count in mice transplanted with MLL-Af4 re-expressing empty vector (Ctrl) or IGF2BP3 in the two groups at 6 weeks. **C.** Spleen weights of mice transplanted with MLL-Af4 re-expressing empty vector (Ctrl) or IGF2BP3 in the two groups at 6 weeks. **D.** Quantitation of spleen cell count in mice transplanted with MLL-Af4 re-expressing empty vector (Ctrl) or IGF2BP3 in the two groups at 6 weeks. **E.** Quantitation of bone marrow CD11b+ cell count in mice transplanted with MLL-Af4 re-expressing empty vector (Ctrl) or IGF2BP3 in the two groups at 6 weeks. **F.** Quantitation of bone marrow Lin-cell count along with representative FACS plots in mice transplanted with MLL-Af4 re-expressing empty vector (Ctrl) or IGF2BP3 in the two groups at 6 weeks. **G.** Quantitation of bone marrow CD11b+cKit+ cell count in mice transplanted with MLL-Af4 re-expressing empty vector (Ctrl) or IGF2BP3 in the two groups at 6 weeks. **H.** Quantitation of bone marrow LSK (Lin-cKit+Sca1-) cell count in mice transplanted with MLL-Af4 re-expressing empty vector (Ctrl) or IGF2BP3 in the two groups at 6 weeks. **I.** Quantitation of bone marrow CD11b+Sca1-(potential LIC; Tran et al.) cell count in mice transplanted with MLL-Af4 re-expressing empty vector (Ctrl) or IGF2BP3 in the two groups at 6 weeks. **J.** Seahorse XF Extracellular acidification rate (ECAR) kinetic trace for bone marrow cells isolated from the empty vector (Ctrl) or IGF2BP3 re-expression group at 6 weeks (n = 4, each group; for representation n = 2). **K.** Aggregate lactate efflux rates from Seahorse XF Analysis in cells described above. **L.** ELISA measurement of m^6^A RNA modifications in splenic tumors isolated from mice transplanted with MLL-Af4 re-expressing empty vector (Ctrl) or IGF2BP3 in the two groups at 6 weeks. All data are n = 2 biological replicates. *, p<0.05; **, p<0.01; ***, p<0.001.

## DISCUSSION

Despite having a 0.1%-0.4%^31^ occupancy on RNA, aberrant m^6^A RNA modification has been implicated in leukemogenesis^32^. To date, the role of m^6^A in oncogenesis has been best elucidated in the case of m^6^A writers (METTL3/14 complex) and erasers (FTO/ALKBH5)^33–37^. These studies have determined that both m6A writers and erasers can promote oncogenesis. This suggests that interpretation of the m^6^A modification within the cell is key in determining the effect on cellular homeostasis; in line with this idea, the YTH family of m^6^A readers were found to play a role in oncogenesis^38,39^. With the identification of the IGF2BP family of proteins and others as m^6^A readers, a general trend for several of these reader proteins playing critical roles in cancer pathogenesis is emerging^40–42^. In this study, we report the unexpected finding that the m^6^A reader IGF2BP3 drives changes in RNA methylation via the regulation of cancer cell metabolism, potentiating post-transcriptional amplification of oncogenic gene expression.

Using a combined approach involving Seahorse XF Analysis and mass spectrometry, we observed significant and consistent reductions in glycolytic flux in IGF2BP3-depleted cells. We also observed a reduction in the steady-state levels and *de novo* synthesis of serine and glycine, which use the glycolytic intermediate 3-phosphoglycerate as a biosynthetic precursor. Increased glycolysis and associated flux into serine biosynthesis have been shown to support tumor growth and survival by fueling the production of S-adenosyl methionine (SAM), the primary methyl donor for global methylation reactions^43,44^, which was also decreased in the IGF2BP3-depleted cells. Interestingly, there was no consistent decrease in oxidative phosphorylation or relative incorporation from glucose or glutamine into TCA cycle intermediates upon IGF2BP3 loss, suggesting a specific and targeted effect of IGF2BP3 on glycolysis and one-carbon metabolism rather than global depression of overall metabolic rates in these leukemia models. It may be that subtle changes in transcription and translation of mitochondrial proteins may not manifest in functional changes in these cells but could in other model systems rely more on oxidative phosphorylation to fuel energetics. Nonetheless, we did observe a specific decrease in α-ketoglutarate (α-kg) levels without commensurate changes in other TCA cycle intermediates. Given that the intracellular α-KG/succinate and α-KG/fumarate ratios regulate the α-kg-dependent dioxygenase family of demethylases and prolyl hydroxylases^45^, it may be that IGF2BP3 regulates broader epigenetic, epitranscriptomic, and transcriptional control via modulation of these metabolites.

Given the observed impact on one-carbon metabolism and SAM, we queried the impact on methylation reactions within cells depleted for IGF2BP3. We observed reduced m^6^A levels following IGF2BP3 depletion. This reduction was not related to a change in RNA m^6^A-methylase or demethylase activity within the cells, and the key enzymes involved in m^6^A showed no change in protein levels. Further arguing for a direct effect on methylation, STM2457 (METTL3 inhibitor) and PF-9366 (MAT2A inhibitor) showed robust growth inhibition in wild-type but not IGF2BP3 depleted cells. This suggests that IGF2BP3 supports SAM biosynthesis, and the availability of SAM is suggested to be rate-limiting for METTL3 activity. While this study was in progress, another group reported modulation of m^6^A RNA methylation by IGF2BP3 via the m^6^A eraser FTO via overexpression of IGF2BP3^46^. While some portions of the study are in agreement with our observations, we did not observe changes in either FTO levels or in RNA demethylase activity. Nonetheless, this work does perhaps illuminate another facet of the intriguing role of IGF2BP3 beyond simply being a passive reader of m^6^A modifications.

We next queried the targets of IGF2BP3 in search of mechanistic understanding. In the current study, enhanced Cross-linking immunoprecipitation (eCLIP-seq) of IGF2BP3 identified several direct targets of IGF2BP3 in the central pathways of the Glycine-Serine cycle, Folate Cycle and the Methyl cycle, which together constitute one-carbon metabolism (OCM)^47^. Because the current RNA-seq dataset did not show significant changes in the RNA levels of many of these genes, we undertook a targeted western blotting approach to query if key regulators were indeed de-regulated. Importantly, we found that MAT2A, a component of the MAT2 rate-limiting enzyme for SAM production, was significantly and consistently reduced. Interestingly, while MAT2A itself was not a direct target of IGF2BP3, the MAT2B mRNA, encoding the other component of the MAT2 enzyme and an allosteric regulator of MAT2A^28^, demonstrated IGF2BP3 CLIP sites within its 3’UTR. Concordantly, MAT2B protein levels were reduced on western blot. Nonetheless, there could be other factors also contributing to the observed downregulation of SAM. Decreased glycolytic flux may play a role, given our observations of reduced ECAR and lactate efflux rate. We also noted downregulation of PKM2, an isoform of *PKM* specifically overexpressed in cancers, in the absence of IGF2BP3. Interestingly, PKM2 activity is reduced in response to serine deprivation, and when in excess, serine binds to and activates PKM2 to increase glycolysis and decrease flux to serine production^48^. Hence, multiple mechanisms are likely to contribute to the observed changes in SAM.

Interestingly, mRNA level measurements from wild-type and knockout cells did not reveal altered abundance of mRNA transcripts encoding the metabolic regulators. To query this aspect of protein translation, we turned to polysome profiling. The polysome profiling revealed no change in the polysome gradient fractionation of MAT2A mRNA, implying no change in translation. On the other hand, both PKM and MAT2B demonstrated an alteration consistent with the model that IGF2BP3’s presence promotes their translation. Therefore, we suspected that the decrease in MAT2A protein levels could be due to the loss of MAT2B, which stabilizes MAT2A, which was confirmed in cycloheximide chase experiment^28^. Hence, our experiments point to a novel mechanism of gene expression regulation by IGF2BP3, where IGF2BP3 impacts translation. We also observed a downregulation in the protein expression of different genes involved in the folate cycle, which has also been linked to RNA methylation^49,50^. However, only MTR is a direct target of IGF2BP3, and did not show notable changes in polysome profiles. Notably, some of the enzymes involved in serine and glycine biosynthesis (PHGDH, PSAT1, SHMT1) showed upregulation in IGF2BP3-deficient cells. This may be the result of a feedback mechanism in response to decreased substrate availability. Hence, we do not fully understand the basis of IGF2BP3-based regulation of metabolic genes. This is further highlighted by the fact that IGF2BP3-based binding and regulation are likely combinatorial-based not only on m^6^A but also sequence, spacing and other RNA modifications, such as m^7^G^51,52^.

We note that MYC is a direct target of IGF2BP3 based both on our own studies and that of others^10,40^. Indeed, we found downregulation of MYC in the current study with depletion of IGF2BP3 (data not shown). Further experiments downregulating MYC demonstrated altered glycolysis but failed to recapitulate other aspects of the IGF2BP3 metabolic-epitranscriptomic phenotype. Particularly, m^6^A was not altered following MYC downregulation (data not shown). Still, recent studies have purported to link m^6^A and MYC in mature B-cell neoplasms and as a regulator of glutamine metabolism in AML^53^ ^54^. Hence, the full extent of involvement of MYC in this pathway is not yet known, but does not detract, from the novelty of our central finding of an m^6^A-reader influencing deposition of the very same modification.

Significantly, rescue experiments with overexpression of IGF2BP3 successfully restored the metabolic-epitranscriptomic changes that we altered by depletion of the protein in multiple model systems. This suggests a model where aberrant expression of IGF2BP3, in our case driven by MLL-AF4, begins a sequence of events resulting in altered metabolism. This leads us to propose a model where, once activated, IGF2BP3 expression reinforces a cellular metabolic and epitranscriptomic phenotype that maintains oncogenesis. Consistently, IGF2BP3 re/overexpression not only rescued cell growth and leukemogenesis *in vitro* and *in vivo*, but also increased m^6^A modifications in the Lin-*Igf2bp3 del/del*, SEM and Lin-MLL-Af4 systems. We also confirmed decreased m^6^A levels, as a consequence of *Igf2bp3* deletion, from *in vivo* leukemia samples^12^. Additionally, we utilized a small molecule inhibitor of IGF2BP3, I3IN-002 (manuscript under review), finding a decrease in glycolysis and m^6^A levels both in vitro and *in vivo,* using a PDX model. Altogether, we provide multiple orthogonal lines of evidence to show that the expression of IGF2BP3 results in the metabolic-epitranscriptomic phenotype described here.

While our study provides evidence for the role of IGF2BP3 in the metabolic control of the cancer cell epitranscriptome, many questions remain unanswered. What is the transcript-level impact of IGF2BP3 depletion on methylation? How do multiple RNA modifications interact to regulate IGF2BP3 function and, more broadly, other epitranscriptomic readers? How does the abundance of methyl donors impact DNA and histone methylation, and how does that impact gene expression in the context of IGF2BP3-driven oncogenesis? We acknowledge that there are many important and interesting questions to arise from our work, and these form several focal points for further research.

In conclusion, IGF2BP3, a non-canonical m^6^A reader, regulates the abundance of the m^6^A modification *in vitro* and *in vivo*, via an effect on metabolism. Our data suggest that the translational control of MAT2B mRNA, and potentially others, is an important regulatory interaction controlled by IGF2BP3. There are specific mRNA targets regulated via translational control that contribute to this function of IGF2BP3. Our findings provide a novel insight into how m^6^A modifications may be propagated and retained by a change in the cellular metabolic milieu. The implications for positive feedback regulation may underlie the potency of IGF2BP3 as a key regulator of leukemogenesis. In the future, a detailed understanding of the many aspects of IGF2BP3 function may aid in designing rational combinatorial therapies that will pre-empt resistance and relapse.

## METHODS

### Cell lines and cell culture

All cell lines were maintained in standard conditions in an incubator at 37 °C and 5% CO_2_. Human cell line HEK 293T (*ATCC*® *CRL3216*™) and B-ALL cell lines, RS4;11 (ATCC CRL-1873), NALM6 (ATCC CRL-3273), and SEM (DMZ-ACC 546) were cultured as previously described^55^. Immortalized MLL-Af4 transformed hematopoietic stem and progenitor cells derived from mouse bone marrow (MLL-Af4 Lin-cells) were cultured in IMDM with 15% FBS, supplemented with recombinant mouse stem cell factor (SCF, 100 ng/mL, Thermo Fisher), recombinant mouse Interleukin-6 (IL-6, 4 ng/mL, Thermo Fisher), recombinant human FMS like tyrosine kinase 3 ligand (FLT3-L, 50 ng/mL, Thermo Fisher) and mouse thrombopoietin (TPO,50 ng/mL, Thermo Fisher).

### CRISPR/Cas9-mediated deletion/overexpression of IGF2BP3 in cell lines

Human B-ALL cell lines SEM, RS4;11, and NALM6 were depleted for IGF2BP3 using lentiviral delivery of CRISPR/Cas9 components in a two-vector system and sgRNA sequence as previously described^12,55^. For the rescue experiments, the codon-optimized IGF2BP3 (MIG-CO IGF2BP3) sequence was synthesized and cloned in MSCV-IRES-GFP (MIG) by GeneScript. The constructs were delivered using pGAG/POL helper plasmids and pseudotyped with pVsVG.

The MSCV-MLL-FLAG-Af4 plasmid was generously provided by Michael Thirman (University of Chicago) through a material transfer agreement^56^. Immortalized MLL-Af4 Lin-cells were initially isolated from bone marrow of Cas9-GFP mice and then transformed using retroviral transduction with MLL-Af4 retroviral supernatant, with four rounds of transduction with MLL-Af4 retroviral supernatant, followed by selection in G418 supplemented media at 400 µg/mL for 7 days, as previously described^11^. Cells were then stably transduced with lentiviral supernatant containing sgRNA against *Igf2bp3* (I3sg2) or non-targeting (NT), and sorted on GFP and mCherry positivity ^12,55^ For rescue experiments, germline Igf2bp3 MLL-Af4 Lin cells underwent second transduction with retroviral supernatant containing MSCV-IRES-GFP (MIG) or MSCV-IRES-GFP-IGF2BP3 (MIG-IGF2BP3) and were sorted according to GFP positivity.

### Protein extraction and Western blot

Cell lysates were made in non-denaturing cell lysis buffer and RIPA lysis buffer. Lysates were electrophoresed using SDS-PAGE using standard conditions^10^. The complete list of antibodies used is listed in the Supp. Table 3.

### Methylcellulose-based colony-forming unit assays

The assay was performed by seeding MLL-Af4 Igf2bp3 germline knockout Lin– cells and IGF2BP3 re-expressed line into MethoCult colony-forming media (STEMCELL Technologies, M3434) at seeding densities of 250 to 2500. Cells were cultured in MethoCult media for 10 to 12 days and counted for total colony number.

### Cell Proliferation, drug cytotoxicity and viability assays

*IGF2BP3* KO (Knockout) and NT (Non-Targeting Control) cells were seeded at 1500 cells/well in 96-well plates and cultured for 72 hours at 5% CO_2_ and 37°C. Cell titre Glo (CTG) reagents were added according to the manufacturer’s instructions (Promega CellTiter kit), and luminescence was measured (Varioskan LUX multimode microplate reader, ThermoFisher). Five technical replicates were prepared for each sample. For inhibitor treatment, cells were treated with the drug or a 0.1% dimethyl sulfoxide (DMSO) control, using concentrations and periods specified in the figure legends. The following inhibitors were used: I3IN002 (Lab synthesized), STM2457 (Catalog No. S9870, Selleckchem), and PF9366 (Catalog No. S0435, Selleckchem).

### Cycloheximide CHASE assay

Briefly, 50 mg/mL cycloheximide (CHX) (01810, Sigma-Aldrich) was added to SEM cells with or without IGF2BP3 knockout at different time points, ranging from 30 minutes to 8 hours before collection. Cells were harvested and collected for translation analysis of different proteins.

### SUnSET Assay

The assay was performed as described earlier^27^. Briefly, 50 mg/mL Puromycin (P4512, Sigma-Aldrich) was added to SEM cells with or without IGF2BP3 knockout at different time points, ranging from 45 minutes to 3 hours before collection. Cells were harvested and collected for Western blot analysis using a monoclonal antibody against puromycin (clone 12D10 MABE343, Sigma-Aldrich) to monitor translation directly.

### Colorimetric measurement of m^6^A levels

Total RNA was extracted from cells with or without IGF2BP3 knockout or other conditions and the corresponding reagents were added according to the manufacturer’s instructions for the m^6^A detection kit (EpiGentek, USA). Finally, changes in the OD value in each well were detected by an enzymatic labeling system at a wavelength of 450 nm within 2–15 min of reagent addition. The following formula was used:

m^6^A% = [(SampleOD-NCOD) + S]/[(PCOD-NCOD) + P] * 100%

S: The total amount of sample RNA added (ng)

P: Total amount of positive control RNA added (ng)

### Colorimetric measurement of Methylase and Demethylase Activity

RNA methylase and demethylase activity were measured using the commercially available Epigenase m^6^A Methylase Activity/Inhibition Assay Kit (Epigentek; P-9019) and Demethylase Activity/Inhibition Assay Kit (Epigentek; P-9013), respectively. The assays were performed per the manufacturer’s protocol. Briefly, 10 μg of total protein lysate was used for control and knockouts. The samples were incubated on the plate for 90 min, washed with wash buffer, and incubated with capture antibody, detection antibody, and enhancer antibody for 60, 30, and 30 min, respectively. After washing five times with a wash buffer, the developer solution was added, and the colour change was monitored for 5 minutes. The reaction was stopped using a stop solution and read at 450 nm using a Varioskan Lux multimode microplate reader (Thermo Fisher). Methylase and Demethylase activity were reported in units of OD/h/mg and normalized against the standard curve.

### Respirometry

All oxygen consumption and extracellular acidification measurements were conducted using an Agilent Seahorse XF Pro or XF^e^96 Analyzer. Experiments were performed at 37°C and pH 7.4. All respiratory parameters were corrected for non-mitochondrial respiration and background signal from the instrument with the addition of 200 nM rotenone and 1 μM antimycin A. Where appropriate, oligomycin was used at 2 μM unless otherwise specified, and FCCP concentrations were titrated to determine an optimal concentration for a given experiment. Unbuffered DMEM assay medium was composed of DMEM (Sigma #5030; pH 7.4) supplemented with 31.6 mM NaCl, 3 mg/l phenol red, and 5 mM HEPES unless otherwise indicated.

### GC/MS Analysis and stable isotope tracing

Experiments were performed as described previously^57^. Metabolite extraction was conducted with a Folch-like method using a 5:2:5 ratio of methanol: water: chloroform. For the isotope tracing experiment, 5 million SEM cells per technical replicate were plated in a medium containing either 10 mM ^13^C_6_ glucose (Cambridge Isotope Laboratories) or 6 mM ^13^C_5_ glutamine (Cambridge Isotope Laboratories) for 24 hours. After incubation, the cells were washed with ice-cold 0.9% (w/v) NaCl and then resuspended in a mix of ice-cold methanol, water containing 5 µg/mL norvaline (Sigma #N7502; an internal standard) and chloroform. Samples were then vortexed for 1 min and centrifuged at 10,000g for 5 min at 4°C. The polar fraction (top layer) was removed, and the samples were dried overnight using a refrigerated CentriVap vacuum concentrator (LabConco). Metabolites (50 nmol to 23 pmol) were extracted alongside the cell samples to ensure the signal fell within the linear detection range of the instrument. The dried polar metabolites were reconstituted in 20 µL of 2% (w/v) methoxyamine in pyridine prior to a 45-min incubation at 37°C. Subsequently, 20 µL of MTBSTFA with 1% tertbutyldimethylchlorosilane was added to samples, followed by an additional 45-min incubation at 37°C. Samples were analyzed using Agilent MassHunter software and FluxFix software (http://fluxfix.science) to correct for the abundance of natural heavy isotopes against an in-house reference set of unlabeled metabolite standards^58^.

### Metabolite extraction and mass-spectrometry-based metabolomics analysis

SEM cells with or without IGF2BP3 knockout were cultured in their regular culture medium without glucose but supplemented with ^13^C_6_ Glucose (10 mM, Cambridge Isotope Laboratories, Inc.). 24 h later, 5×10^6^ cells were collected and rinsed with PBS, and 1 mL cold 80% methanol (Optima* LC/MS, Fisher Scientific) was added to cells. As an internal standard, 1 µM norvaline (Sigma-Aldrich) was added to each sample. Samples were then vortexed every 5 min, three times and spun down at top speed for 5 min at 4 °C. The supernatants were transferred to a newtubes, and the pellet was resuspended in 0.5 M NaOH for protein estimation. The extracts were dried overnight using a refrigerated CentriVap vacuum concentrator (LabConco) and stored at −80°C. The mass spectrometry-based analysis of extracted metabolites was conducted at UCLA Metabolomics Center. Dried metabolites were resuspended in 100 µl 50% ACN:water and 5 µl was loaded onto a Luna 3µm NH2 100A (150 × 2.0 mm) column (Phenomenex). The chromatographic separation was performed on a Vanquish Flex (Thermo Scientific) with mobile phases A (5 mM NH4AcO pH 9.9) and B (ACN) and a flow rate of 200 μl/min. A linear gradient from 15% A to 95% A over 18 min was followed by 7 min isocratic flow at 95% A and re-equilibration to 15% A. Metabolites were detected with a Thermo Scientific Q Exactive mass spectrometer run with polarity switching in full scan mode with an m/z range of 70-975 and 70.000 resolution. Maven (v 8.1.27.11) was utilized to quantify the targeted metabolites by AreaTop using accurate mass measurements (< 5 ppm) and expected retention time previously verified with standards. Values were normalized to cell number or protein content of extracted material, where applicable. ^13^C natural abundance corrections were made using AccuCor. Total amounts were calculated by summing up the intensities of all detected isotopologues of a given metabolite. Data analysis was performed using in-house R scripts. MetaboAnalyst (v6.0) was then used to analyze the enriched metabolic pathways of significantly changed metabolites with default parameters^26^.

### Sucrose gradient preparation for polysome profiling

Linear sucrose gradients (15-45%) were made by dissolving sucrose in a polysome gradient buffer (20 mM Tris-HCl, pH 7.5, 200 mM KCl, 25 mM MgCl₂). Approximately 6.5 mL of the 15% sucrose solution was layered into SW41 centrifuge tubes, followed by the 45% solution underneath. The gradients were mixed using the Biocomp Gradient Station, then stabilized at 4°C overnight.

### Polyribosome Profiling and Fractionation

The experiment was performed as previously described^59^. Briefly, SEM cells were treated with 100 µg/mL CHX for 10 minutes at 37°C, then pelleted (200 rpm for 10 minutes). The pellet was resuspended in 9 mL PBS (without MgCl₂ and KCl) and pelleted again. After removing the supernatant, cells were lysed in an ice-cold lysis buffer (20 mM Tris-HCl pH 7.5, 200 mM KCl, 25 mM MgCl₂, 0.5% NP-40, 100 µg/mL CHX, supplemented with protease inhibitors). The lysate was passed through a 23-gauge needle 3-5 times and incubated on ice for 10 minutes. Following incubation, the lysate was centrifuged (10,000 rcf for 10 minutes at 4°C) to separate nuclear and membrane fractions. The cytosolic supernatant was then layered onto the pre-formed sucrose gradient and ultracentrifuged at 40,000 rpm for 2 hours at 4°C using a Beckman SW41 rotor. The gradients were manually fractionated (non-continuous) from top to bottom into 22 samples of 0.4 mL each using the Biocomp Gradient Station (Piston Gradient Fractionator (Model 153)). Fractions were pooled into 96-well plates with a 1:1 ratio of 2x RNA Shield containing spike-in RNA (SARS-CoV N1).

### RNA extraction and cDNA synthesis for Polysome profiling

Total RNA was extracted from cell culture pellets using an Agilent Bravo automated liquid handling platform and Quick-DNA/RNA Viral Magbead kit following the manufacturer’s protocol (Zymo Research). Samples were treated with RQ1 RNase-Free DNase following the manufacturer’s protocol (Promega). cDNA synthesis was then performed using a High-Capacity cDNA Reverse Transcription Kit following the manufacturer’s protocol (Applied Biosystems). cDNA was diluted 1:10 and qPCR was performed.

### Polysome profiling qPCR

qPCR experiments were performed on a QuantStudio 6 Flex Real-Time PCR System (Applied Biosystems). Experiments were performed using Luna Universal qPCR Master Mix, following the manufacturer’s protocol (New England Biolabs). SARS-CoV2 N1 RNA spike-in was used to normalize the relative expression levels of target mRNAs to the cytosol fraction with the ΔΔCt method. Primer sequences are provided in Supp. Table 2.

### Animal Experiments

For *in vivo* studies, C57BL/6J, B6.SJL-*Ptprc^a^ Pepc^b,^*/BoyJ (B6 CD45.1), and B6J.129(Cg)-Gt(ROSA)26Sor^tm1.1(CAG-cas9,-EGFP)Fezh/J^ (Cas9-GFP, BL/6J) were procured from The Jackson Laboratory. The *Igf2bp3* (*Igf2bp3^del/del^*) mice used in this study were generated and maintained as previously described^11^. The bone marrow transplant experiments were performed as previously described^12^. For *in vivo* drug treatment, one week after transplantation, mice were injected with vehicle or I3IN-002 three times a week at a dose of 25 mg/kg, for three weeks, with an endpoint at four weeks post-transplantation. Flow cytometry was performed to check the peripheral blood engraftment of the leukemic cells at week 2 and week 4. Once the peripheral blood engraftment reached >20%, the experiment was terminated and tissues were harvested to be analyzed by Flow cytometry, histology and m^6^A ELISA. For rescue experiments, MSCV-IRES-GFP (Control) or MSCV-IRES-GFP-IGF2BP3 (MIG-IGF2BP3) overexpressed MLL-Af4 Lin-cells were transplanted as described previously^11^. All the animal experiments received Institutional Animal Research Committee approval at UCLA.

### PDX experiments

NOD.Cg-PrkdcSCIDIl2rgtm1Wjl/SzJ (NOD/SCID/IL-2Rγ−/−, NSG) mice were maintained in the animal facilities at the University of California, Los Angeles (UCLA). Six-to ten-week-old females were utilized to study the in vivo efficacy of the small molecule inhibitor against IGF2BP3. 5,00,000 PDX cells (PDX#22694) were engrafted in the NSG mice and one week following transplantation, mice were injected with vehicle or I3IN-002, three times a week at a dose of 25 mg/kg, for three weeks, with an end point at four weeks post-transplantation. The engraftment of the leukemic cells was checked by Flow cytometry at week 2 and week 4 in the peripheral blood. Once the peripheral blood engraftment reached >20% at 4 weeks, the experiment was terminated, and tissues were harvested to be analyzed by Flow cytometry and m^6^A ELISA.

### eCLIP-Seq

eCLIP studies were performed by Eclipse Bioinnovations Inc https://eclipsebio.com/ according to the published single-end eCLIP protocol^60^. Briefly, 20×10^6^ SEM cells were grown and UV cross-linked at 400 mJoules/cm^2^ with 254 nm radiation, flash frozen, and stored until use at −80°C. Crosslinked cell pellets were further processed by Eclipse Bioinnovations for eCLIP using a rabbit anti-IGF2BP3 antibody (MBL, RN009P). A parallel Size-Matched Input (SMInput) library was also generated as a control where the samples were treated identically, except they were not immunoprecipitated with anti-IGF2BP3 antibody. Protein-RNA complexes were separated on an SDS-PAGE gel, transferred to a PVDF membrane and isolated using standard iCLIP protocol^61^. Libraries were then amplified as previously described^62^. Three replicates using 20 million SEM cells per replicate (3 IP libraries and 3 size-matched input libraries) were processed, yielding six libraries. Sequencing was performed as SE72 on the NextSeq platform.

### eCLIP-seq Read Processing

Data were processed similarly to the standard eCLIP pipeline^62^. Briefly, reads were trimmed (cutadapt) and FASTQs were aligned with STAR v2.7.8a^63^ to the human genome (GRCh38, GENCODE v38 annotation). To remove all the repetitive elements, a reverse intersection of all peak files with the repeatmasker bed file (downloaded from the University of California Santa Cruz (UCSC) table viewer) was performed^64^. PCR duplicate reads were removed, and the aligned files were further processed and analyzed for peaks enriched over the background using Skipper v1.0.0^65^. IGF2BP3 eCLIP fine-mapped peak sets were filtered for peaks with log2(fold change) ≥ and ≥ 3.0, respectively, in terms of mean read counts in IP vs. size-matched input^62^.

### Joint Analysis of Metabolomics and eCLIP datasets

For finding the enriched metabolic pathways, using the IGF2BP3 eCLIP datasets, MetaboAnalyst^26^ (v6.0) “joint pathway analysis” module was used. For the enrichment analysis, a list of significantly changed metabolites along with IGF2BP3’s mRNA target list was used with default parameters^26^. Additionally, for the enrichment of metabolic pathways using HumanCyc^66^ and Metabolomics Workbench Metabolites^67^ from the IGF2BP3 knockout datasets, enrichR^68^ web module was used.

### RNA extraction

Total RNA was extracted with TRI Reagent (Zymo Research) using the manufacturer’s protocol for the RNA isolation with the following modification: one additional RNA ethanol wash step was included. After the total RNA was solubilized in ddH_2_0, one overnight ethanol precipitation step was included for further purification of the total RNA.

### Illumina sequencing of mRNA libraries

Total RNA was isolated for cell culture pellets as described above. 2 μg of total RNA for each sample was used for mRNA library preparation using the NEXT-FLEX Rapid Directional RNA-Seq Kit 2.0 following the manufacturer’s protocol (Perkin Elmer Applied Genomics). Before library preparation, total RNA samples were subjected to Poly(A) selection and purification using the NEXTFLEX Poly(A) Beads Kit 2.0 following the manufacturer’s protocol (Perkin Elmer Applied Genomics). Pooled mRNA sequencing libraries were sequenced on an Illumina NovaSeq S4 at the UC Davis Sequencing Core Facility, generating 150 bp paired-end reads.

### m^6^A Dot Blots

Total RNA was isolated from cell culture pellets as described above. RNA was first denatured for 5 minutes at 95° C and then placed on ice for 5 minutes. Hybond-N^+^ nylon membrane (Amersham Biosciences) was pre-soaked for 5 minutes in 2x SSC before RNA blotting was performed using a commercial apparatus with a vacuum manifold (Schleicer and Schuell, Inc.). The membrane was cross-linked twice using the auto-cross link function on a UV Stratalinker 2400. The membrane was then blocked for 1 hour at room temperature in a 10% blocking solution/0.1% PBST before incubation overnight at 4°C with primary m^6^A antibody (Millipore MABE-1006) (1:1,000) in 5% blocking solution/0.1% PBST. The membrane was washed 3 times for 5 minutes in 0.1% PBST and incubated for 1 hour at room temperature in HRP mouse secondary antibody (1:10,000) in 5% blocking solution/0.1% TBST. The membrane was washed 3 times for 5 minutes in 0.1% PBST and then incubated for 5 minutes in SuperSignal West Pico PLUS substrate (Thermo Scientific). The membrane then visualized using a BioRad Chemidoc using the optimal auto-exposure setting. As a loading control, samples were run in parallel on a separate membrane and stained with 0.1% methylene blue for two hours before de-staining in ddH_2_O until the dots were visible.

### Statistical analysis

The data shown represents mean ± SD for continuous numerical data. Two-tailed student’s t-tests or one-way ANOVA followed by Bonferroni’s multiple comparisons test were performed using GraphPad Prism software and conducted as described in the figure legends. Survival analyses were performed using the Kaplan-Meier method, with comparisons made using log-rank tests, followed by Bonferroni’s correction for multiple comparisons. A *P* value less than 0.05 was considered significant.

## Supporting information

Supplementary Images

## ACKNOWLEDGMENTS

We are grateful to Prof. Neil Garg, Georgia Scherer and Dr. Jacob Sorrentino for the co-development and synthesis of the small molecule compound I3IN-002. We thank prior members of the Rao lab for meaningful conversations about this work. These studies were supported by R01CA264986 from the National Institutes of Health (DSR, JRS), the Jonsson Comprehensive Cancer Center (JCCC) (DSR and ASD), CIRM DISC-2-13456 from the California Institute of Regenerative Medicine (DSR), and R03CA251854 (DSR). This project has partly been supported by the UCLA Jonsson Comprehensive Cancer Center’s Office of Cancer Training and Education (JCCC OCTE) (GS). Flow cytometry was performed in the UCLA JCCC and Center for AIDS Research Flow Cytometry Core Facility that is supported by National Institutes of Health awards AI-28697, and award number P30CA016042, the JCCC, the UCLA AIDS Institute, and the David Geffen School of Medicine at UCLA.

## CONFLICT OF INTEREST DISCLOSURES

DSR has served as a consultant to AbbVie, a pharmaceutical company that develops and markets drugs for hematologic disorders. DSR, MLT and AKJ are inventors on a patent application that includes the compound I3IN-002.

## AUTHOR CONTRIBUTIONS

G.S..: Designed research, Performed experiments, Acquired data, Analyzed data, Generated Figures, Wrote manuscript

M.G.: Performed experiments, Acquired data, Analyzed data, Assisted in generating figures

A.E.J.: Performed experiments, Acquired data, Analyzed data

A.K.J..: Performed experiments, Acquired data, Analyzed data

Z.T.N.: Performed experiments, Acquired data, Analyzed data

A.R.: Performed experiments

M.L.T.: Performed experiments

T.L.L.: Performed experiments, Acquired data

T.M.T.: Performed experiments

L.E.S.K.: Analyzed data, Assisted in generating figures

A.J.R.: Analyzed data

L.S.: Performed experiments

J. t. H.: Performed experiments

A.S.D.: Designed research, Analyzed data, Edited manuscript, Secured Funding

J.R.S..: Designed research, Analyzed data, Edited manuscript, Secured Funding

D.S.R.: Designed research, Analyzed data, Wrote manuscript, Secured Funding, Project leader

## SUPPLEMENTARY FIGURE LEGENDS

**Supplementary Figure 1. IGF2BP3 does not grossly regulate oxidative phosphorylation in B-ALL cells.**

**A.** Seahorse XF kinetic trace for Oxygen consumption rate (OCR) for control versus IGF2BP3 deleted SEM cells.

**B.** Maximal respiration rate measurements for control versus IGF2BP3 deleted SEM cells, as measured in Seahorse experiments.

**C.** Rate of ATP generation from oxidative phosphorylation for control versus IGF2BP3 deleted SEM cells, as measured in Seahorse experiments.

**D.** ATP-linked respiration measurement for control versus IGF2BP3 deleted NALM6 cells, as measured in Seahorse experiments.

**E.** Maximal respiration measurements for control versus IGF2BP3 deleted NALM6 cells, as measured in Seahorse experiments.

**F.** Rate of ATP generation from oxidative phosphorylation for control versus IGF2BP3 deleted NALM6 cells, as measured in Seahorse experiments.

**G.** Steady-state levels of TCA-cycle intermediates measured by GC/MS in control versus IGF2BP3-deleted (I3sg2) SEM cells.

**H.** Steady-state levels of amino acids measured by GC/MS in control versus IGF2BP3-deleted (I3sg2) SEM cells.

**I.** Incorporation of carbon from ^13^C-labeled glucose, into citric acid cycle intermediates (citrate, alpha-ketoglutarate, fumarate and malate) measured as mole percent enrichment (MPE) from GC-MS experiments.

**J.** Incorporation of carbon from ^13^C-labeled glutamine into citric acid cycle intermediates (citrate, alpha-ketoglutarate, fumarate and malate), measured as mole percent enrichment (MPE) from GC-MS experiments.

All data are n>3 biological replicates. *, p<0.05; **, p<0.01; ***, p<0.001.

**Supplementary Figure 2. Additional metabolites show consistent changes in both knockout lines of IGF2BP3.**

**A-B.** Abundance of Lactate and Fructose-1,6-bisphosphate measured by LC-MS in control versus IGF2BP3-deleted (I3sg2) SEM cells

**C-D.** Abundance and incorporation of carbon from ^13^C-labeled glucose into Cystathionine measured as mole percent enrichment (MPE) from LC-MS experiments.

**Supplementary Figure 3. Related to Figure 4.**

**A.** Metabolism-specific analyses of prior datasets produced by our groups (Human Cyc 2016, dataset from ref.^10^).

**B.** Metabolism-specific analyses of prior datasets produced by our groups (Metabolomics workbench metabolites, dataset from ref ^10^).

**C.** Metabolism-specific analyses of prior datasets produced by our groups (Human Cyc 2016, dataset from ref.^11^).

**D.** Metabolism-specific analyses of prior datasets produced by our groups (Metabolomics workbench metabolites, dataset from ref.^11^).

**E.** Western blot analysis of key metabolic enzymes in Lin-MLL-Af4 cells.

**F.** 10-45% Sucrose gradient fractionation of cytosolic extracts from control or IGF2BP3-depleted SEM cells. PKM mRNA distribution was measured by RT-qPCR.

**G-I**. As in F, sucrose gradient fractionation of PSAT1, SHMT1, MTHFR mRNAs, IGF2BP3 targets that showed increases or mild decreases in protein expression levels.

**Supplementary Figure 4. Related to Figure 4.**

**A.** Western blot analysis of puromycin incorporation for studying changes in the global translation at different time points (SuNSET Assay) in IGF2BP3 expressing and depleted cells (left: sg2; right sg5). β-Actin (ACTIN) was used as a loading control.

**B.** 10-45% Sucrose gradient fractionation of cytosolic extracts from control or IGF2BP3-depleted SEM cells. MAT2A mRNA distribution was measured by RT-qPCR.

**C.** Western blot analysis of the Cycloheximide Chase Assay in IGF2BP3 sufficient and depleted cells (left: sg2; right sg5) for MAT2A. β-Actin (ACTIN) was used as a loading control and to normalize the change in MAT2A expression over time in the IGF2BP3 sufficient and depleted cells. ImageJ software was used to quantify the change in protein amounts of MAT2A over time and plotted as a graph in terms of fold change (IGF2BP3-depleted/NT).

**Supplementary Figure 5. Related to Figure 6.**

**A.** Quantitation of bone marrow B220+ cell count in mice transplanted with MLL-Af4 re-expressing empty vector (Ctrl) or IGF2BP3 in the two groups at 6 weeks.

**B.** Quantitation of bone marrow CD11b+Sca1-cell count in mice transplanted with MLL-Af4 re-expressing empty vector (Ctrl) or IGF2BP3 in the two groups at 6 weeks.

**C.** Percentage engraftment of CD45.2 Lin-cells in spleen from *Igf2bp3^del/del^* mice transduced with MLL-Af4 re-expressing empty vector (Ctrl) or IGF2BP3 in the two groups at 6 weeks.

**D.** Quantitation of splenic CD11b+ cell count in mice transplanted with MLL-Af4 re-expressing empty vector (Ctrl) or IGF2BP3 in the two groups at 6 weeks.

**E.** Quantitation of splenic lineage-negative cell count in mice transplanted with MLL-Af4 re-expressing empty vector (Ctrl) or IGF2BP3 in the two groups at 6 weeks.

**F.** Quantitation of splenic LSK (Lin-cKit+Sca1-) cell count in mice transplanted with MLL-Af4 re-expressing empty vector (Ctrl) or IGF2BP3 in the two groups at 6 weeks.

**G.** Quantitation of splenic CD11b+Sca1-(potential LIC; Tran et al.) cell count in mice transplanted with MLL-Af4 re-expressing empty vector (Ctrl) or IGF2BP3 in the two groups at 6 weeks.

**H.** Quantitation of splenic cKit+CD34+ (potential LIC; Lin et al.) cell count in mice transplanted with MLL-Af4 re-expressing empty vector (Ctrl) or IGF2BP3 in the two groups at 6 weeks.

**I.** Quantitation of splenic CD11b+cKit+ (potential LIC; Lin et al.) cell count in mice transplanted with MLL-Af4 re-expressing empty vector (Ctrl) or IGF2BP3 in the two groups at 6 weeks.

All data are n = 2 biological replicates. *, p<0.05; **, p<0.01; ***, p<0.001.

## References

1. Desrosiers, R., Friderici, K., and Rottman, F. (1974). Identification of methylated nucleosides in messenger RNA from Novikoff hepatoma cells. Proc Natl Acad Sci U S A 71, 3971–3975. 10.1073/pnas.71.10.3971.

2. Wang, X., Zhao, B.S., Roundtree, I.A., Lu, Z., Han, D., Ma, H., Weng, X., Chen, K., Shi, H., and He, C. (2015). N(6)-methyladenosine Modulates Messenger RNA Translation Efficiency. Cell 161, 1388–1399. 10.1016/j.cell.2015.05.014.

3. Wang, Y., and Zhao, J.C. (2016). Update: Mechanisms Underlying N(6)-Methyladenosine Modification of Eukaryotic mRNA. Trends Genet 32, 763–773. 10.1016/j.tig.2016.09.006.

4. Roundtree, I.A., Evans, M.E., Pan, T., and He, C. (2017). Dynamic RNA Modifications in Gene Expression Regulation. Cell 169, 1187–1200. 10.1016/j.cell.2017.05.045.

5. Meyer, K.D., Saletore, Y., Zumbo, P., Elemento, O., Mason, C.E., and Jaffrey, S.R. (2012). Comprehensive analysis of mRNA methylation reveals enrichment in 3’ UTRs and near stop codons. Cell 149, 1635–1646. 10.1016/j.cell.2012.05.003 S0092-8674(12)00536-3 [pii].

6. Ke, S., Alemu, E.A., Mertens, C., Gantman, E.C., Fak, J.J., Mele, A., Haripal, B., Zucker-Scharff, I., Moore, M.J., Park, C.Y., et al. (2015). A majority of m6A residues are in the last exons, allowing the potential for 3’ UTR regulation. Genes Dev 29, 2037–2053. 10.1101/gad.269415.115.

7. Roignant, J.Y., and Soller, M. (2017). m(6)A in mRNA: An Ancient Mechanism for Fine-Tuning Gene Expression. Trends Genet 33, 380–390. 10.1016/j.tig.2017.04.003.

8. Bell, J.L., Wächter, K., Mühleck, B., Pazaitis, N., Köhn, M., Lederer, M., and Hüttelmaier, S. (2013). Insulin-like growth factor 2 mRNA-binding proteins (IGF2BPs): post-transcriptional drivers of cancer progression? Cellular and Molecular Life Sciences 70, 2657–2675. 10.1007/s00018-012-1186-z.

9. Ennajdaoui, H., Howard, Jonathan M., Sterne-Weiler, T., Jahanbani, F., Coyne, Doyle J., Uren, Philip J., Dargyte, M., Katzman, S., Draper, Jolene M., Wallace, A., et al. (2016). IGF2BP3 Modulates the Interaction of Invasion-Associated Transcripts with RISC. Cell Reports 15, 1876–1883. 10.1016/j.celrep.2016.04.083.

10. Palanichamy, J.K., Tran, T.M., Howard, J.M., Contreras, J.R., Fernando, T.R., Sterne-Weiler, T., Katzman, S., Toloue, M., Yan, W., Basso, G., et al. (2016). RNA-binding protein IGF2BP3 targeting of oncogenic transcripts promotes hematopoietic progenitor proliferation. J Clin Invest 126, 1495–1511. 10.1172/JCI80046.

11. Tran, T.M., Philipp, J., Bassi, J.S., Nibber, N., Draper, J.M., Lin, T.L., Palanichamy, J.K., Jaiswal, A.K., Silva, O., Paing, M., et al. (2022). The RNA-binding protein IGF2BP3 is critical for MLL-AF4-mediated leukemogenesis. Leukemia 36, 68–79. 10.1038/s41375-021-01346-7.

12. Lin, T.L., Jaiswal, A.K., Ritter, A.J., Reppas, J., Tran, T.M., Neeb, Z.T., Katzman, S., Thaxton, M.L., Cohen, A., Sanford, J.R., and Rao, D.S. (2024). Targeting IGF2BP3 enhances antileukemic effects of menin-MLL inhibition in MLL-AF4 leukemia. Blood Adv 8, 261–275. 10.1182/bloodadvances.2023011132.

13. Zhou, T., Xiao, Z., Lu, J., Zhang, L., Bo, L., and Wang, J. (2023 Nov 15). IGF2BP3-mediated regulation of GLS and GLUD1 gene expression promotes treg-induced immune escape in human cervical cancer. Am J Cancer Res 13(*11*), 5289–5305.

14. Lin, Z.A.-O., Li, J.A.-O., Zhang, J.A.-O., Feng, W.A.-O., Lu, J.A.-O., Ma, X.A.-O., Ding, W.A.-O., Ouyang, S.A.-O., Lu, J.A.-O., Yue, P.A.-O., et al. (2023 Jul 5). Metabolic Reprogramming Driven by IGF2BP3 Promotes Acquired Resistance to EGFR Inhibitors in Non-Small Cell Lung Cancer. Cancer Res 83(*13*), 2187–2207. 10.1158/0008-5472.CAN-22-3059.

15. Mentch, S.J., and Locasale, J.W. (2016 Jan). One-carbon metabolism and epigenetics: understanding the specificity. Ann N Y Acad Sci 1363(*1*), 91–98. 10.1111/nyas.12956.

16. Ducker, G.S., and Rabinowitz, J.D. (2017 Jan 10). One-Carbon Metabolism in Health and Disease. Cell Metab 25(*1*), 27–42. 10.1016/j.cmet.2016.08.009.

17. Patel, S.B., Nemkov, T., D’Alessandro, A., and Welner, R.S. (2022). Deciphering Metabolic Adaptability of Leukemic Stem Cells. Front Oncol 12, 846149. 10.3389/fonc.2022.846149.

18. Zarou, M.M., Rattigan, K.M., Sarnello, D., Shokry, E., Dawson, A., Ianniciello, A., Dunn, K., Copland, M., Sumpton, D., Vazquez, A., and Helgason, G.V. (2024). Inhibition of mitochondrial folate metabolism drives differentiation through mTORC1 mediated purine sensing. Nat Commun 15, 1931. 10.1038/s41467-024-46114-0.

19. Jain, M., Nilsson, R., Sharma, S., Madhusudhan, N., Kitami, T., Souza, A.L., Kafri, R., Kirschner, M.W., Clish, C.B., and Mootha, V.K. (2012). Metabolite profiling identifies a key role for glycine in rapid cancer cell proliferation. Science 336, 1040–1044. 10.1126/science.1218595.

20. Maddocks, O.D., Berkers, C.R., Mason, S.M., Zheng, L., Blyth, K., Gottlieb, E., and Vousden, K.H. (2013). Serine starvation induces stress and p53-dependent metabolic remodelling in cancer cells. Nature 493, 542–546. 10.1038/nature11743.

21. Pikman, Y., Ocasio-Martinez, N., Alexe, G., Dimitrov, B., Kitara, S., Diehl, F.F., Robichaud, A.L., Conway, A.S., Ross, L., Su, A., et al. (2022). Targeting serine hydroxymethyltransferases 1 and 2 for T-cell acute lymphoblastic leukemia therapy. Leukemia 36, 348–360. 10.1038/s41375-021-01361-8.

22. Duan, M., Liu, H., Xu, S., Yang, Z., Zhang, F., Wang, G., Wang, Y., Zhao, S., and Jiang, X. (2023 Jul 20). IGF2BPs as novel m(6)A readers: Diverse roles in regulating cancer cell biological functions, hypoxia adaptation, metabolism, and immunosuppressive tumor microenvironment. Genes Dis 11(*2*), 890–920. 10.1016/j.gendis.2023.06.017.

23. Desousa, B.R., Kim, K.K.O., Jones, A.E., Ball, A.B., Hsieh, W.Y., Swain, P., Morrow, D.H., Brownstein, A.J., Ferrick, D.A., Shirihai, O.S., et al. (2023). Calculation of ATP production rates using the Seahorse XF Analyzer. EMBO reports 24, e56380. 10.15252/embr.202256380.

24. Mentch, S.J., Mehrmohamadi, M., Huang, L., Liu, X., Gupta, D., Mattocks, D., Gómez Padilla, P., Ables, G., Bamman, M.M., Thalacker-Mercer, A.E., et al. (2015 Nov 3). Histone Methylation Dynamics and Gene Regulation Occur through the Sensing of One-Carbon Metabolism. Cell Metab 22(*5*), 861–873. 10.1016/j.cmet.2015.08.024.

25. Quinlan, C.A.-O., Kaiser, S.E., Bolaños, B., Nowlin, D., Grantner, R., Karlicek-Bryant, S., Feng, J.L., Jenkinson, S., Freeman-Cook, K., Dann, S.G., et al. (2017 Jul). Targeting S-adenosylmethionine biosynthesis with a novel allosteric inhibitor of Mat2A. Nat Chem Biol 13(*7*), 785–792. 10.1038/nchembio.2384.

26. Pang, Z., Lu, Y., Zhou, G., Hui, F., Xu, L., Viau, C., Spigelman, Aliya F., MacDonald, Patrick E., Wishart, David S., Li, S., and Xia, J. (2024). MetaboAnalyst 6.0: towards a unified platform for metabolomics data processing, analysis and interpretation. Nucleic Acids Research 52, W398–W406. 10.1093/nar/gkae253.

27. Schmidt, E.K., Clavarino G Fau - Ceppi, M., Ceppi M Fau - Pierre, P., and Pierre, P. (2009 Apr). SUnSET, a nonradioactive method to monitor protein synthesis. Nat Methods 6(*4*), 275–277. 10.1038/nmeth.1314.

28. Bailey, J., Douglas, H., Masino, L., de Carvalho, L.P.S., and Argyrou, A. (2021). Human Mat2A Uses an Ordered Kinetic Mechanism and Is Stabilized but Not Regulated by Mat2B. Biochemistry 60, 3621–3632. 10.1021/acs.biochem.1c00672.

29. Miao, Y., Du, Q., Zhang, H.G., Yuan, Y., Zuo, Y., and Zheng, H. (2023 Jun 5). Cycloheximide (CHX) Chase Assay to Examine Protein Half-life. Bio Protoc 13(*11*), e4690. 10.21769/BioProtoc.4690.

30. Townsend, E.C., Murakami, M.A., Christodoulou, A., Christie, A.L., Köster, J., DeSouza, T.A., Morgan, E.A., Kallgren, S.P., Liu, H., Wu, S.C., et al. (2016 Apr 11). The Public Repository of Xenografts Enables Discovery and Randomized Phase II-like Trials in Mice. Cancer Cell 29(*4*), 574–586. 10.1016/j.ccell.2016.03.008.

31. Wei CM, G.A., Moss B (1975 Apr). Methylated nucleotides block 5’ terminus of HeLa cell messenger RNA. Cell 4(*4*), 379–386. 10.1016/0092-8674(75)90158-0.

32. Zhang, N., Shen, Y., Li, H., Chen, Y., Zhang, P., Lou, S., and Deng, J. (2022). The m6A reader IGF2BP3 promotes acute myeloid leukemia progression by enhancing RCC2 stability. Experimental & Molecular Medicine 54, 194–205. 10.1038/s12276-022-00735-x.

33. Bartosovic, M., Molares, H.C., Gregorova, P., Hrossova, D., Kudla, G., and Vanacova, S. (2017). N6-methyladenosine demethylase FTO targets pre-mRNAs and regulates alternative splicing and 3’-end processing. Nucleic Acids Res 45, 11356–11370. 10.1093/nar/gkx778.

34. Cui, Q., Shi, H., Ye, P., Li, L., Qu, Q., Sun, G., Sun, G., Lu, Z., Huang, Y., Yang, C.G., et al. (2017). m(6)A RNA Methylation Regulates the Self-Renewal and Tumorigenesis of Glioblastoma Stem Cells. Cell Rep 18, 2622–2634. 10.1016/j.celrep.2017.02.059.

35. Li, Z., Weng, H., Su, R., Weng, X., Zuo, Z., Li, C., Huang, H., Nachtergaele, S., Dong, L., Hu, C., et al. (2017). FTO Plays an Oncogenic Role in Acute Myeloid Leukemia as a N(6)-Methyladenosine RNA Demethylase. Cancer cell 31, 127–141. 10.1016/j.ccell.2016.11.017.

36. Wang, L., and Tang, Y. (2023). N6-methyladenosine (m6A) in cancer stem cell: From molecular mechanisms to therapeutic implications. Biomed Pharmacother 163, 114846. 10.1016/j.biopha.2023.114846.

37. Chen, T., Hao, Y.J., Zhang, Y., Li, M.M., Wang, M., Han, W., Wu, Y., Lv, Y., Hao, J., Wang, L., et al. (2015). m(6)A RNA methylation is regulated by microRNAs and promotes reprogramming to pluripotency. Cell Stem Cell 16, 289–301. 10.1016/j.stem.2015.01.016.

38. Li, L., Tang, C., Ye, J., Xu, D., Chu, C., Wang, L., Zhou, Q., Gan, S., and Liu, B. (2023). Bioinformatic analysis of m6A “reader” YTH family in pan-cancer as a clinical prognosis biomarker. Scientific Reports 13, 17350. 10.1038/s41598-023-44143-1.

39. Shi, R., Ying, S., Li, Y., Zhu, L., Wang, X., and Jin, H. (2021). Linking the YTH domain to cancer: the importance of YTH family proteins in epigenetics. Cell Death & Disease 12, 346. 10.1038/s41419-021-03625-8.

40. Huang, H., Weng, H., Sun, W., Qin, X., Shi, H., Wu, H., Zhao, B.S., Mesquita, A., Liu, C., Yuan, C.L., et al. (2018). Recognition of RNA N6-methyladenosine by IGF2BP proteins enhances mRNA stability and translation. Nature Cell Biology 20, 285–295. 10.1038/s41556-018-0045-z.

41. Liu, N., Zhou, K.I., Parisien, M., Dai, Q., Diatchenko, L., and Pan, T. (2017). N6-methyladenosine alters RNA structure to regulate binding of a low-complexity protein. Nucleic Acids Res 45, 6051–6063. 10.1093/nar/gkx141.

42. Covelo-Molares, H., Bartosovic, M., and Vanacova, S. (2018). RNA methylation in nuclear pre-mRNA processing. Wiley Interdiscip Rev RNA 9, e1489. 10.1002/wrna.1489.

43. Geeraerts, S.L., Heylen, E., De Keersmaecker, K., and Kampen, K.R. (2021). The ins and outs of serine and glycine metabolism in cancer. Nature Metabolism 3, 131–141. 10.1038/s42255-020-00329-9.

44. Maddocks, Oliver D.K., Labuschagne, Christiaan F., Adams, Peter D., and Vousden, Karen H. (2016). Serine Metabolism Supports the Methionine Cycle and DNA/RNA Methylation through De Novo ATP Synthesis in Cancer Cells. Molecular Cell 61, 210–221. 10.1016/j.molcel.2015.12.014.

45. Baksh, S.C., and Finley, L.W.S. (2021 Jan). Metabolic Coordination of Cell Fate by α-Ketoglutarate-Dependent Dioxygenases. Trends Cell Biol 31(*1*), 24–36. 10.1016/j.tcb.2020.09.010.

46. Dai, W., Tian, R., Yu, L., Bian, S., Chen, Y., Yin, B., Luan, Y., Chen, S., Fan, Z., Yan, R., et al. (2024). Overcoming therapeutic resistance in oncolytic herpes virotherapy by targeting IGF2BP3-induced NETosis in malignant glioma. Nature Communications 15, 131. 10.1038/s41467-023-44576-2.

47. Newman, A.C., and Maddocks, O.D.K. (2017). One-carbon metabolism in cancer. British Journal of Cancer 116, 1499–1504. 10.1038/bjc.2017.118.

48. Chaneton, B., Hillmann, P., Zheng, L., Martin, A.C.L., Maddocks, O.D.K., Chokkathukalam, A., Coyle, J.E., Jankevics, A., Holding, F.P., Vousden, K.H., et al. (2012). Serine is a natural ligand and allosteric activator of pyruvate kinase M2. Nature 491, 458–462. 10.1038/nature11540.

49. Green, N.H., Galvan, D.L., Badal, S.S., Chang, B.H., LeBleu, V.S., Long, J., Jonasch, E., and Danesh, F.R. (2019). MTHFD2 links RNA methylation to metabolic reprogramming in renal cell carcinoma. Oncogene 38, 6211–6225. 10.1038/s41388-019-0869-4.

50. Zhang, W., Bai, Y., Hao, L., Zhao, Y., Zhang, L., Ding, W., Qi, Y., and Xu, Q.A.-O. (2024 Sep 2). One-carbon metabolism supports S-adenosylmethionine and m6A methylation to control the osteogenesis of bone marrow stem cells and bone formation. J Bone Miner Res 39(*9*), 1356–1370. 10.1093/jbmr/zjae121.

51. Schneider, T., Hung, L.-H., Aziz, M., Wilmen, A., Thaum, S., Wagner, J., Janowski, R., Müller, S., Schreiner, S., Friedhoff, P., et al. (2019). Combinatorial recognition of clustered RNA elements by the multidomain RNA-binding protein IMP3. Nat. Commun. 10, 2266. 10.1038/s41467-019-09769-8.

52. Liu, C., Dou, X., Zhao, Y., Zhang, L., Zhang, L., Dai, Q., Liu, J., Wu, T., Xiao, Y., and He, C. (2024). IGF2BP3 promotes mRNA degradation through internal m7G modification. Nature Communications 15, 7421. 10.1038/s41467-024-51634-w.

53. Wu, G.A.-O., Suo, C.A.-O., Yang, Y., Shen, S., Sun, L., Li, S.T., Zhou, Y., Yang, D., Wang, Y.A.-O., Cai, Y., et al. (2021 Mar 3). MYC promotes cancer progression by modulating m(6) A modifications to suppress target gene translation. EMBO Rep 22(3), e51519. 10.15252/embr.202051519.

54. Weng, H., Huang, F., Yu, Z., Chen, Z., Prince, E., Kang, Y., Zhou, K., Li, W., Hu, J., Fu, C., et al. (2022). The m(6)A reader IGF2BP2 regulates glutamine metabolism and represents a therapeutic target in acute myeloid leukemia. Cancer Cell 40, 1566–1582.e1510. 10.1016/j.ccell.2022.10.004.

55. Jaiswal, A.K., Truong, H., Tran, T.M., Lin, T.L., Casero, D., Alberti, M.O., and Rao, D.S. (2021). Focused CRISPR-Cas9 genetic screening reveals USO1 as a vulnerability in B-cell acute lymphoblastic leukemia. Scientific Reports 11, 13158. 10.1038/s41598-021-92448-w.

56. Lin, S., Luo, R.T., Ptasinska, A., Kerry, J., Assi, S.A., Wunderlich, M., Imamura, T., Kaberlein, J.J., Rayes, A., Althoff, M.J., et al. (2016 Nov 14). Instructive Role of MLL-Fusion Proteins Revealed by a Model of t(4;11) Pro-B Acute Lymphoblastic Leukemia. Cancer Cell 30(*5*), 737–749. 10.1016/j.ccell.2016.10.008.

57. Cordes, T., and Metallo, C.M. (2019). Quantifying Intermediary Metabolism and Lipogenesis in Cultured Mammalian Cells Using Stable Isotope Tracing and Mass Spectrometry. In High-Throughput Metabolomics: Methods and Protocols, A. D’Alessandro, ed. (Springer New York), pp. 219–241. 10.1007/978-1-4939-9236-2_14.

58. Trefely, S.A.-O., Ashwell, P., and Snyder, N.W. (2016 Nov 25). FluxFix: automatic isotopologue normalization for metabolic tracer analysis. BMC Bioinformatics 17(*1*), 485. 10.1186/s12859-016-1360-7.

59. Martinez-Nunez, R.T., and Sanford, J.R. (2016). Studying Isoform-Specific mRNA Recruitment to Polyribosomes with Frac-seq. Methods Mol Biol 1358, 99–108. 10.1007/978-1-4939-3067-8_6.

60. Conway, A.E., Van Nostrand, E.L., Pratt, G.A., Aigner, S., Wilbert, M.L., Sundararaman, B., Freese, P., Lambert, N.J., Sathe, S., Liang, T.Y., et al. (2016). Enhanced CLIP Uncovers IMP Protein-RNA Targets in Human Pluripotent Stem Cells Important for Cell Adhesion and Survival. Cell Rep 15, 666–679. 10.1016/j.celrep.2016.03.052.

61. Huppertz, I., Attig, J., D’Ambrogio, A., Easton, L.E., Sibley, C.R., Sugimoto, Y., Tajnik, M., König, J., and Ule, J. (2014 Feb). iCLIP: protein-RNA interactions at nucleotide resolution. Methods 65(*3*), 274–287. 10.1016/j.ymeth.2013.10.011.

62. Van Nostrand, E.L., Pratt, G.A., Shishkin, A.A., Gelboin-Burkhart, C., Fang, M.Y., Sundararaman, B., Blue, S.M., Nguyen, T.B., Surka, C., Elkins, K., et al. (2016). Robust transcriptome-wide discovery of RNA-binding protein binding sites with enhanced CLIP (eCLIP). Nat Methods 13, 508–514. 10.1038/nmeth.3810.

63. Dobin, A., Davis, C.A., Schlesinger, F., Drenkow, J., Zaleski, C., Jha, S., Batut, P., Chaisson, M., and Gingeras, T.R. (2013). STAR: ultrafast universal RNA-seq aligner. Bioinformatics 29, 15–21. 10.1093/bioinformatics/bts635.

64. Karolchik, D., Baertsch R Fau - Diekhans, M., Diekhans M Fau - Furey, T.S., Furey Ts Fau - Hinrichs, A., Hinrichs A Fau - Lu, Y.T., Lu Yt Fau - Roskin, K.M., Roskin Km Fau - Schwartz, M., Schwartz M Fau - Sugnet, C.W., Sugnet Cw Fau - Thomas, D.J., Thomas Dj Fau - Weber, R.J., et al. (2003 Jan 1). The UCSC Genome Browser Database. Nucleic Acids Res 31(*1*), 51-54. 10.1093/nar/gkg129.

65. Boyle, E.A., Her, H.L., Mueller, J.R., Naritomi, J.T., Nguyen, G.G., and Yeo, G.W. (2023 May 4). Skipper analysis of eCLIP datasets enables sensitive detection of constrained translation factor binding sites. Cell Genom 3(*6*), 100317. 10.1016/j.xgen.2023.100317.

66. Karp, P.D., Billington, R., Caspi, R., Fulcher, C.A., Latendresse, M., Kothari, A., Keseler, I.M., Krummenacker, M., Midford, P.E., Ong, Q., et al. (2019 Jul 19). The BioCyc collection of microbial genomes and metabolic pathways. Brief Bioinform 20(*4*), 1085–1093. 10.1093/bib/bbx085.

67. Sud, M., Fahy, E., Cotter, D., Azam, K., Vadivelu, I., Burant, C., Edison, A., Fiehn, O., Higashi, R., Nair, K.S., et al. (2016 Jan 4). Metabolomics Workbench: An international repository for metabolomics data and metadata, metabolite standards, protocols, tutorials and training, and analysis tools. Nucleic Acids Res 44(*D1*), D463–470. 10.1093/nar/gkv1042.

68. Kuleshov, M.V., Jones, M.R., Rouillard, A.D., Fernandez, N.F., Duan, Q., Wang, Z., Koplev, S., Jenkins, S.L., Jagodnik, K.M., Lachmann, A., et al. (2016). Enrichr: a comprehensive gene set enrichment analysis web server 2016 update. Nucleic acids research 44, W90–W97. 10.1093/nar/gkw377.

