## Supplementary Table 1 for "Metabolic regulation of RNA methylation by the m^6^A-reader IGF2BP3"

| | | | | | | | Abundance | | | | | | | |
|---|---|---|---|---|---|---|---|---|---|---|---|---|---|---|
| **Name** | **KEGG.ID** | **Condition** | **Av** | **Std** | **CV** | **Exp001** | **Exp002** | **Exp003** | **ANOVA** | **Sig** | **Av2** | **denom** | **Norm_Av** | **Norm_Std** |
| 2-HG | C02630 | NT | 2859093.575 | 504035.2637 | 0.176291979 | 3397034.376 | 2397726.922 | 2782519.428 | 0.010276591 | * | 2859093.575 | 1472383.027 | 1.941813728 | 0.342326184 |
| 2-HG | C02630 | I3sg2 | 1472383.027 | 149725.2701 | 0.101689076 | 1328818.62 | 1627589.461 | 1460741 | 0.010276591 | * | 1472383.027 | 1472383.027 | 1 | 0.101689076 |
| 5-Oxoproline | C01879 | NT | 18863038.67 | 779789.2118 | 0.041339533 | 18164844.8 | 19704543.16 | 18719728.05 | 0.000262781 | *** | 18863038.67 | 12510231.2 | 1.507808957 | 0.062332118 |
| 5-Oxoproline | C01879 | I3sg2 | 12510231.2 | 459601.7737 | 0.036738072 | 12862811.41 | 12677451.21 | 11990431 | 0.000262781 | *** | 12510231.2 | 12510231.2 | 1 | 0.036738072 |
| Aconitate | C00417 | NT | 721821.5876 | 92815.00153 | 0.128584408 | 723441.2934 | 813816.1361 | 628207.3333 | 0.002131325 | ** | 721821.5876 | 342258.8069 | 2.108993467 | 0.271183676 |
| Aconitate | C00417 | I3sg2 | 342258.8069 | 8631.109144 | 0.025218078 | 338252.5729 | 336358.8478 | 352165 | 0.002131325 | ** | 342258.8069 | 342258.8069 | 1 | 0.025218078 |
| ADP/ATP | NA | NT | 0.323362593 | 0.075073275 | 0.232164377 | 0.240170398 | 0.343856782 | 0.3860606 | 0.029523822 | * | 0.323362593 | 0.323362593 | 1 | 0.232164377 |
| ADP/ATP | NA | I3sg2 | 0.596193324 | 0.12119501 | 0.203281394 | 0.735189085 | 0.512611257 | 0.540779632 | 0.029523822 | * | 0.596193324 | 0.323362593 | 1.843730032 | 0.37479601 |
| Ala | C00041 | NT | 38355759.67 | 2461590.506 | 0.064177858 | 37997701.18 | 40976770.36 | 36092807.46 | 0.000181364 | *** | 38355759.67 | 14346365.02 | 2.673552472 | 0.171582872 |
| Ala | C00041 | I3sg2 | 14346365.02 | 1904125.407 | 0.132725287 | 14905739.1 | 15908149.96 | 12225206 | 0.000181364 | *** | 14346365.02 | 14346365.02 | 1 | 0.132725287 |
| AMP/ATP | NA | NT | 0.13773589 | 0.069453305 | 0.504249874 | 0.103621127 | 0.091937096 | 0.217649448 | 0.005794947 | ** | 0.13773589 | 0.13773589 | 1 | 0.504249874 |
| AMP/ATP | NA | I3sg2 | 0.366865806 | 0.02513092 | 0.068501669 | 0.395884483 | 0.352335404 | 0.35237753 | 0.005794947 | ** | 0.366865806 | 0.13773589 | 2.663545462 | 0.182457311 |
| Arg-Succ | C03406 | NT | 180405.7208 | 44737.43405 | 0.247982347 | 232000.0307 | 156835.3106 | 152381.8211 | 0.034126665 | * | 180405.7208 | 97023.80907 | 1.859396395 | 0.461097482 |
| Arg-Succ | C03406 | I3sg2 | 97023.80907 | 9220.625882 | 0.095034672 | 107517.3796 | 93337.04758 | 90217 | 0.034126665 | * | 97023.80907 | 97023.80907 | 1 | 0.095034672 |
| Asn | C00152 | NT | 3638780.038 | 273384.0218 | 0.075130681 | 3945569.897 | 3549796.368 | 3420973.85 | 0.007091273 | ** | 3638780.038 | 2469811.942 | 1.473302471 | 0.110690218 |
| Asn | C00152 | I3sg2 | 2469811.942 | 290256.4239 | 0.11752167 | 2189942.035 | 2769444.791 | 2450049 | 0.007091273 | ** | 2469811.942 | 2469811.942 | 1 | 0.11752167 |
| C | D07769 | NT | 26961.09829 | 17866.44755 | 0.662675065 | 14622.93965 | 18811.05302 | 47449.30221 | 0.008742237 | ** | 26961.09829 | 26961.09829 | 1 | 0.662675065 |
| C | D07769 | I3sg2 | 114667.6104 | 26241.50935 | 0.228848489 | 94223.49258 | 144258.3385 | 105521 | 0.008742237 | ** | 114667.6104 | 26961.09829 | 4.253076382 | 0.973310103 |
| CDP-choline | C00307 | NT | 90801.01786 | 15304.95083 | 0.168554838 | 91780.08954 | 75030.03629 | 105592.9278 | 0.001094597 | ** | 90801.01786 | 90801.01786 | 1 | 0.168554838 |
| CDP-choline | C00307 | I3sg2 | 258699.8323 | 31011.04404 | 0.119872687 | 239923.5896 | 294493.9072 | 241682 | 0.001094597 | ** | 258699.8323 | 90801.01786 | 2.849085157 | 0.341527494 |
| CDP-EtA | C00570 | NT | 1020971.933 | 121002.4052 | 0.118516877 | 989035.2091 | 919141.2446 | 1154739.346 | 0.049192741 | * | 1020971.933 | 1020971.933 | 1 | 0.118516877 |
| CDP-EtA | C00570 | I3sg2 | 1225912.478 | 38961.52869 | 0.031781656 | 1199734.576 | 1207314.856 | 1270688 | 0.049192741 | * | 1225912.478 | 1020971.933 | 1.200730831 | 0.038161214 |
| Cit | C00158 | NT | 126509850.5 | 16754130.57 | 0.132433407 | 142747162.3 | 109282782.6 | 127499606.6 | 0.011703467 | * | 126509850.5 | 81266230.7 | 1.556733337 | 0.2061635 |
| Cit | C00158 | I3sg2 | 81266230.7 | 6057371.556 | 0.074537376 | 75988180.48 | 87879974.62 | 79930537 | 0.011703467 | * | 81266230.7 | 81266230.7 | 1 | 0.074537376 |
| CMP | C00055 | NT | 99197.01866 | 65194.82721 | 0.657225672 | 86453.35639 | 41314.94097 | 169822.7586 | 0.02355065 | * | 99197.01866 | 99197.01866 | 1 | 0.657225672 |
| CMP | C00055 | I3sg2 | 272383.2625 | 53317.80832 | 0.195745538 | 221123.8017 | 268480.9859 | 327545 | 0.02355065 | * | 272383.2625 | 99197.01866 | 2.745881542 | 0.53749406 |
| Cystathionine | C02291 | NT | 4287607.921 | 710307.4741 | 0.165665212 | 5044794.14 | 4182032.175 | 3635997.449 | 0.000709469 | *** | 4287607.921 | 425386.633 | 10.07931982 | 1.669792652 |
| Cystathionine | C02291 | I3sg2 | 425386.633 | 18135.04051 | 0.0426319 | 438599.9223 | 432848.9767 | 404711 | 0.000709469 | *** | 425386.633 | 425386.633 | 1 | 0.0426319 |
| Cytosine | C00380 | NT | 410561.4359 | 51520.30167 | 0.125487435 | 359120.0278 | 410404.0094 | 462160.2704 | 0.000205362 | *** | 410561.4359 | 16195.68674 | 25.35004797 | 3.181112508 |
| Cytosine | C00380 | I3sg2 | 16195.68674 | 11379.82805 | 0.702645601 | 29329.40986 | 9988.650363 | 9269 | 0.000205362 | *** | 16195.68674 | 16195.68674 | 1 | 0.702645601 |
| F16BP | C00354 | NT | 2671150.925 | 589170.5546 | 0.220568051 | 1999656.987 | 2912325.907 | 3101469.882 | 0.044236093 | * | 2671150.925 | 1363877.605 | 1.958497533 | 0.431981985 |
| F16BP | C00354 | I3sg2 | 1363877.605 | 513438.4898 | 0.37645496 | 1943042.506 | 1184045.311 | 964545 | 0.044236093 | * | 1363877.605 | 1363877.605 | 1 | 0.37645496 |
| Folate | C00504 | NT | 172610.8509 | 46141.23435 | 0.267313637 | 140941.5396 | 151340.1895 | 225550.8235 | 0.023071132 | * | 172610.8509 | 172610.8509 | 1 | 0.267313637 |
| Folate | C00504 | I3sg2 | 370707.5128 | 83861.04954 | 0.22621891 | 273945.1132 | 415855.4252 | 422322 | 0.023071132 | * | 370707.5128 | 172610.8509 | 2.147648951 | 0.485838805 |
| GABA | C00334 | NT | 1717606.077 | 273922.8397 | 0.159479431 | 1980267.269 | 1433663.078 | 1738887.885 | 0.012556498 | * | 1717606.077 | 1717606.077 | 1 | 0.159479431 |
| GABA | C00334 | I3sg2 | 4455348.432 | 1065843.72 | 0.239227916 | 3227139.389 | 5001263.907 | 5137642 | 0.012556498 | * | 4455348.432 | 1717606.077 | 2.593929127 | 0.620540259 |
| Glc | C00031 | NT | 8974296.029 | 2170922.97 | 0.241904542 | 6655628.935 | 10958701.53 | 9308557.627 | 0.004848842 | ** | 8974296.029 | 8974296.029 | 1 | 0.241904542 |
| Glc | C00031 | I3sg2 | 19475786.93 | 2380910.005 | 0.122249746 | 18233737.61 | 17972731.19 | 22220892 | 0.004848842 | ** | 19475786.93 | 8974296.029 | 2.170174337 | 0.265303261 |
| GlcA | C00257 | NT | 703620.2939 | 151018.6014 | 0.214630821 | 831366.6361 | 742544.3602 | 536949.8855 | 0.010939104 | * | 703620.2939 | 270793.1466 | 2.598368175 | 0.557689895 |
| GlcA | C00257 | I3sg2 | 270793.1466 | 71557.7823 | 0.26425256 | 234335.8749 | 353237.5648 | 224806 | 0.010939104 | * | 270793.1466 | 270793.1466 | 1 | 0.26425256 |
| GlcNAc-6P | C00357 | NT | 476299.7703 | 54697.72013 | 0.114838855 | 534641.0003 | 426175.2476 | 468083.0629 | 0.022694534 | * | 476299.7703 | 276884.7344 | 1.720209571 | 0.197546897 |
| GlcNAc-6P | C00357 | I3sg2 | 276884.7344 | 78722.76686 | 0.284316024 | 189879.5986 | 343187.6046 | 297587 | 0.022694534 | * | 276884.7344 | 276884.7344 | 1 | 0.284316024 |
| GlucA | C00191 | NT | 185972.0512 | 26507.36616 | 0.142534139 | 208670.8493 | 156840.2484 | 192405.056 | 0.001188349 | ** | 185972.0512 | 52972.48466 | 3.510729247 | 0.50039877 |
| GlucA | C00191 | I3sg2 | 52972.48466 | 8989.068793 | 0.169693169 | 61208.97614 | 54324.47784 | 43384 | 0.001188349 | ** | 52972.48466 | 52972.48466 | 1 | 0.169693169 |
| Gly | C00037 | NT | 14176813.3 | 450961.996 | 0.031809828 | 13918783.72 | 14697532.4 | 13914123.76 | 0.000633787 | *** | 14176813.3 | 9412258.545 | 1.506207381 | 0.047912198 |
| Gly | C00037 | I3sg2 | 9412258.545 | 722037.384 | 0.076712447 | 8606369.704 | 9630147.931 | 10000258 | 0.000633787 | *** | 9412258.545 | 9412258.545 | 1 | 0.076712447 |
| GSH | C00051 | NT | 95036256.62 | 3630394.158 | 0.038200096 | 99076639.06 | 93983662.48 | 92048468.33 | 0.000126431 | *** | 95036256.62 | 61795077.87 | 1.537925995 | 0.058748921 |
| GSH | C00051 | I3sg2 | 61795077.87 | 1507732.581 | 0.024398911 | 62842120.68 | 60066967.92 | 62476145 | 0.000126431 | *** | 61795077.87 | 61795077.87 | 1 | 0.024398911 |
| His | C00135 | NT | 36016885.1 | 5019268.83 | 0.139358771 | 41744607.8 | 33919805.4 | 32386242.1 | 0.016346722 | * | 36016885.1 | 23768619.18 | 1.515312473 | 0.211172083 |
| His | C00135 | I3sg2 | 23768619.18 | 1778295.24 | 0.074816935 | 21798644.78 | 24251874.75 | 25255338 | 0.016346722 | * | 23768619.18 | 23768619.18 | 1 | 0.074816935 |
| HS2O3 | C00320 | NT | 498005.4162 | 63378.68714 | 0.127265056 | 515024.8451 | 427854.7067 | 551136.6967 | 0.04195572 | * | 498005.4162 | 498005.4162 | 1 | 0.127265056 |
| HS2O3 | C00320 | I3sg2 | 1052878.372 | 319524.6751 | 0.30347729 | 698154.4942 | 1142352.622 | 1318128 | 0.04195572 | * | 1052878.372 | 498005.4162 | 2.114190605 | 0.641608835 |
| IMP | C00130 | NT | 165383.195 | 56795.29316 | 0.34341635 | 139149.8203 | 126446.4208 | 230553.3439 | 0.001606668 | ** | 165383.195 | 165383.195 | 1 | 0.34341635 |
| IMP | C00130 | I3sg2 | 430850.8506 | 20800.42562 | 0.048277555 | 410038.1705 | 430875.3814 | 451639 | 0.001606668 | ** | 430850.8506 | 165383.195 | 2.605167052 | 0.125771096 |
| Lac | C00186 | NT | 16460403.35 | 1457930.558 | 0.088571983 | 15055158.69 | 16360201.24 | 17965850.13 | 0.000431732 | *** | 16460403.35 | 6573700.565 | 2.503978268 | 0.221782319 |
| Lac | C00186 | I3sg2 | 6573700.565 | 658876.5604 | 0.100229171 | 6128783.081 | 6261692.614 | 7330626 | 0.000431732 | *** | 6573700.565 | 6573700.565 | 1 | 0.100229171 |
| NAD+ | C00003 | NT | 7753075.47 | 960044.3437 | 0.12382755 | 8651006.803 | 7867114.807 | 6741104.801 | 0.004195068 | ** | 7753075.47 | 1419297.671 | 5.4626141 | 0.676422123 |
| NAD+ | C00003 | I3sg2 | 1419297.671 | 1601841.318 | 1.128615477 | 3267650.321 | 555037.692 | 435205 | 0.004195068 | ** | 1419297.671 | 1419297.671 | 1 | 1.128615477 |
| NADP+ | C00006 | NT | 745186.4261 | 331962.2055 | 0.445475379 | 1098326.678 | 439512.1651 | 697720.4351 | 0.044005874 | * | 745186.4261 | 141118.0511 | 5.280588985 | 2.352372379 |
| NADP+ | C00006 | I3sg2 | 141118.0511 | 140497.6456 | 0.995603642 | 303097.8399 | 67970.31332 | 52286 | 0.044005874 | * | 141118.0511 | 141118.0511 | 1 | 0.995603642 |
| Orotidine | X00006 | NT | 8260900.083 | 1412560.986 | 0.170993593 | 9890462.581 | 7507135.925 | 7385101.742 | 0.00199713 | ** | 8260900.083 | 2372257.076 | 3.482295475 | 0.595450215 |
| Orotidine | X00006 | I3sg2 | 2372257.076 | 157733.7762 | 0.066491013 | 2204853.922 | 2518104.306 | 2393813 | 0.00199713 | ** | 2372257.076 | 2372257.076 | 1 | 0.066491013 |
| P-Choline | C00588 | NT | 92779212.72 | 6890712.471 | 0.074270004 | 100485643.9 | 87211322.94 | 90640671.37 | 0.0034778 | ** | 92779212.72 | 65853590.06 | 1.408870992 | 0.104636854 |
| P-Choline | C00588 | I3sg2 | 65853590.06 | 3069148.409 | 0.046605635 | 62310970.97 | 67540879.21 | 67708920 | 0.0034778 | ** | 65853590.06 | 65853590.06 | 1 | 0.046605635 |
| P-EtA | C00346 | NT | 829498.1741 | 49590.65872 | 0.059783928 | 885399.2166 | 812296.5373 | 790798.7683 | 0.000244299 | *** | 829498.1741 | 829498.1741 | 1 | 0.059783928 |
| P-EtA | C00346 | I3sg2 | 2174198.317 | 181400.6195 | 0.083433336 | 1968041.872 | 2245170.079 | 2309383 | 0.000244299 | *** | 2174198.317 | 829498.1741 | 2.621100787 | 0.218687184 |
| P-Ser | C01005 | NT | 279396.4551 | 33141.94593 | 0.11861978 | 317585.1501 | 258155.8874 | 262448.3278 | 0.001275843 | ** | 279396.4551 | 118324.9158 | 2.361264771 | 0.280092707 |
| P-Ser | C01005 | I3sg2 | 118324.9158 | 9712.563603 | 0.082083841 | 110271.2004 | 115592.5469 | 129111 | 0.001275843 | ** | 118324.9158 | 118324.9158 | 1 | 0.082083841 |
| Pantothenate | C00864 | NT | 19312766.56 | 3247291.566 | 0.168142226 | 22811047.88 | 16394684.78 | 18732567.01 | 0.010834687 | * | 19312766.56 | 10555780.29 | 1.829591563 | 0.307631599 |
| Pantothenate | C00864 | I3sg2 | 10555780.29 | 907305.1319 | 0.085953393 | 9698084.703 | 11505657.17 | 10463599 | 0.010834687 | * | 10555780.29 | 10555780.29 | 1 | 0.085953393 |
| PO4 | C00009 | NT | 89153786.38 | 5782076.699 | 0.064855088 | 84042857.55 | 95429629.25 | 87988872.35 | 0.000548914 | *** | 89153786.38 | 89153786.38 | 1 | 0.064855088 |
| PO4 | C00009 | I3sg2 | 142603318.1 | 7157955.285 | 0.050194872 | 135135730.3 | 143269104 | 149405120 | 0.000548914 | *** | 142603318.1 | 89153786.38 | 1.599520602 | 0.080287732 |
| Pro-OH | C01157 | NT | 3180612.581 | 492377.0322 | 0.154805724 | 2616356.278 | 3523125.002 | 3402356.463 | 0.026894371 | * | 3180612.581 | 3180612.581 | 1 | 0.154805724 |
| Pro-OH | C01157 | I3sg2 | 4161440.561 | 70592.67516 | 0.016963519 | 4129993.111 | 4112036.57 | 4242292 | 0.026894371 | * | 4161440.561 | 3180612.581 | 1.308377067 | 0.022194679 |
| S7P | C05382 | NT | 328220.9533 | 45014.66599 | 0.137147448 | 307939.7391 | 296914.915 | 379808.2058 | 0.022438136 | * | 328220.9533 | 220418.0378 | 1.489083909 | 0.204224057 |
| S7P | C05382 | I3sg2 | 220418.0378 | 25301.90201 | 0.114790524 | 230038.0345 | 191717.079 | 239499 | 0.022438136 | * | 220418.0378 | 220418.0378 | 1 | 0.114790524 |
| SAM | C00019 | NT | 8361055.992 | 1455552.301 | 0.174087137 | 9959019.888 | 7110946.522 | 8013201.567 | 0.031534885 | * | 8361055.992 | 4824351.543 | 1.733094265 | 0.301709419 |
| SAM | C00019 | I3sg2 | 4824351.543 | 1202389.672 | 0.249233428 | 3435970.219 | 5512305.409 | 5524779 | 0.031534885 | * | 4824351.543 | 4824351.543 | 1 | 0.249233428 |
| Ser | C00065 | NT | 54943554.99 | 1704431.257 | 0.031021496 | 55601550.26 | 56220909.62 | 53008205.1 | 0.001887042 | ** | 54943554.99 | 43796874.77 | 1.254508576 | 0.038916732 |
| Ser | C00065 | I3sg2 | 43796874.77 | 2029439.418 | 0.04633754 | 41453488.83 | 44974144.47 | 44962991 | 0.001887042 | ** | 43796874.77 | 43796874.77 | 1 | 0.04633754 |
| Thr | C00188 | NT | 113115732.6 | 13230038.39 | 0.116960197 | 128118722 | 103120917.5 | 108107558.3 | 0.042640425 | * | 113115732.6 | 86103647.52 | 1.313715921 | 0.153652473 |
| Thr | C00188 | I3sg2 | 86103647.52 | 8901523.835 | 0.103381495 | 82524078.19 | 79549144.37 | 96237720 | 0.042640425 | * | 86103647.52 | 86103647.52 | 1 | 0.103381495 |
| Tyr | C00082 | NT | 91517436.81 | 1987713.391 | 0.021719505 | 93630453.41 | 89684800.19 | 91237056.82 | 0.000602219 | *** | 91517436.81 | 69088720.66 | 1.324636438 | 0.028770447 |
| Tyr | C00082 | I3sg2 | 69088720.66 | 3418958.319 | 0.049486491 | 66265979.83 | 72890358.14 | 68109824 | 0.000602219 | *** | 69088720.66 | 69088720.66 | 1 | 0.049486491 |
| UDP-GlcNAc | C00043 | NT | 22392831.38 | 1306118.14 | 0.058327512 | 23180047.95 | 23113297.68 | 20885148.53 | 0.000400772 | *** | 22392831.38 | 12724474.01 | 1.759823735 | 0.10264614 |
| UDP-GlcNAc | C00043 | I3sg2 | 12724474.01 | 806242.361 | 0.063361547 | 11794407.28 | 13224984.76 | 13154030 | 0.000400772 | *** | 12724474.01 | 12724474.01 | 1 | 0.063361547 |
| Xanthine | C00385 | NT | 34789.29897 | 12519.07779 | 0.35985427 | 20407.3217 | 43243.75747 | 40716.81774 | 6.08E-05 | *** | 34789.29897 | 34789.29897 | 1 | 0.35985427 |
| Xanthine | C00385 | I3sg2 | 189626.2972 | 8645.220673 | 0.045590832 | 188418.2633 | 181648.6284 | 198812 | 6.08E-05 | *** | 189626.2972 | 34789.29897 | 5.450707627 | 0.248502296 |

| Name | KEGG.ID | Condition | Iso | Nr.C | Exp001 | Exp002 | Exp003 | Norm_Av | Norm_Std | CV | MID001 | MID002 | MID003 | Av | ANOVA | Sig |
|---|---|---|---|---|---|---|---|---|---|---|---|---|---|---|---|---|
| | | | | | Isotopologue Distribution | | | | | | | | | | | |
| 2-HG | C02630 | NT | C12 PARENT | 5 | 2271349.59 | 1625827.395 | 1951086.888 | 65.76536 | 1.5724 | 0.0239 | 64.4451 | 65.3461 | 67.5049 | 65.7654 | 3.66E-05 | *** |
| 2-HG | C02630 | I3sg2 | C12 PARENT | 5 | 1269493.211 | 1595707.463 | 1449432.882 | 92.88287 | 1.7379 | 0.0187 | 91.0192 | 93.1701 | 94.4593 | 92.8829 | 3.66E-05 | *** |
| 2-HG | C02630 | NT | C13-02 | 5 | 627376.0082 | 376907.4751 | 455884.7993 | 16.2408 | 1.3864 | 0.0854 | 17.8006 | 15.1489 | 15.773 | 16.2408 | 0.00201 | ** |
| 2-HG | C02630 | I3sg2 | C13-02 | 5 | 120346.0607 | 104329.679 | 75764.15413 | 6.55254 | 1.8882 | 0.2882 | 8.62849 | 6.0916 | 4.93754 | 6.55254 | 0.00201 | ** |
| 2-HG | C02630 | NT | C13-03 | 5 | 196940.6132 | 154646.8542 | 137384.6061 | 5.518922 | 0.7336 | 0.1329 | 5.5878 | 6.21564 | 4.75332 | 5.51892 | 0.00034 | *** |
| 2-HG | C02630 | I3sg2 | C13-03 | 5 | 4913.771116 | 0 | 9255.226371 | 0.318489 | 0.303 | 0.9514 | 0.3523 | 0 | 0.60316 | 0.31849 | 0.00034 | *** |
| 2-HG | C02630 | NT | C13-04 | 5 | 271939.4282 | 188381.7349 | 219132.9223 | 7.622993 | 0.0805 | 0.0106 | 7.71575 | 7.57153 | 7.5817 | 7.62299 | 7.91E-06 | *** |
| 2-HG | C02630 | I3sg2 | C13-04 | 5 | 0 | 12644.96109 | 0 | 0.246104 | 0.4263 | 1.7321 | 0 | 0.73831 | 0 | 0.2461 | 7.91E-06 | *** |
| 2-HG | C02630 | NT | C13-05 | 5 | 156867.4937 | 142263.1647 | 126798.1703 | 4.851921 | 0.7506 | 0.1547 | 4.45081 | 5.71791 | 4.38704 | 4.85192 | 0.00036 | *** |
| 2-HG | C02630 | I3sg2 | C13-05 | 5 | 0 | 0 | 0 | 0 | 0 | NA | 0 | 0 | 0 | 0 | 0.00036 | *** |
| 3PG | C00197 | NT | C12 PARENT | 3 | 120371.2962 | 57828.54109 | 49565.87046 | 8.950552 | 0.104 | 0.0116 | 9.06221 | 8.85633 | 8.93312 | 8.95055 | 0.93228 | |
| 3PG | C00197 | I3sg2 | C12 PARENT | 3 | 62693.69187 | 73090.9014 | 120119.8278 | 9.126449 | 3.3666 | 0.3689 | 12.3699 | 5.64899 | 9.36047 | 9.12645 | 0.93228 | |
| 3PG | C00197 | NT | C13-03 | 3 | 1207906.012 | 595134.503 | 505289.4191 | 91.04945 | 0.104 | 0.0011 | 90.9378 | 91.1437 | 91.0669 | 91.0494 | 0.93228 | |
| 3PG | C00197 | I3sg2 | C13-03 | 3 | 444131.5663 | 1220785.05 | 1163146.548 | 90.87355 | 3.3666 | 0.037 | 87.6301 | 94.351 | 90.6395 | 90.8736 | 0.93228 | |
| 5-Oxoproline | C01879 | NT | C12 PARENT | 5 | 16752451.52 | 18849129.95 | 16775304.91 | 91.70346 | 3.1103 | 0.0339 | 91.1211 | 95.0638 | 88.9255 | 91.7035 | 0.10625 | |
| 5-Oxoproline | C01879 | I3sg2 | C12 PARENT | 5 | 12449711.95 | 12090181.77 | 11645690.55 | 95.52006 | 0.6672 | 0.007 | 96.1152 | 94.7988 | 95.6462 | 95.5201 | 0.10625 | |
| 5-Oxoproline | C01879 | NT | C13-01 | 5 | 0 | 19349.52191 | 162469.1814 | 0.319611 | 0.4716 | 1.4755 | 0 | 0.09759 | 0.86125 | 0.31961 | 0.78879 | |
| 5-Oxoproline | C01879 | I3sg2 | C13-01 | 5 | 0 | 182460.1644 | 0 | 0.476889 | 0.826 | 1.7321 | 0 | 1.43067 | 0 | 0.47689 | 0.78879 | |
| 5-Oxoproline | C01879 | NT | C13-02 | 5 | 1054234.972 | 589752.5589 | 1261958.177 | 5.132746 | 1.9293 | 0.3759 | 5.73427 | 2.97436 | 6.68961 | 5.13275 | 0.15388 | |
| 5-Oxoproline | C01879 | I3sg2 | C13-02 | 5 | 373759.9054 | 369459.4906 | 439784.7565 | 3.131469 | 0.4162 | 0.1329 | 2.88553 | 2.89692 | 3.61196 | 3.13147 | 0.15388 | |
| 5-Oxoproline | C01879 | NT | C13-04 | 5 | 380576.051 | 276736.4205 | 383603.6864 | 1.833075 | 0.3792 | 0.2069 | 2.07005 | 1.39569 | 2.03347 | 1.83307 | 0.00834 | ** |
| 5-Oxoproline | C01879 | I3sg2 | C13-04 | 5 | 101847.0388 | 86748.57716 | 45455.67323 | 0.613269 | 0.2145 | 0.3497 | 0.78629 | 0.68019 | 0.37333 | 0.61327 | 0.00834 | ** |
| 5-Oxoproline | C01879 | NT | C13-05 | 5 | 197565.9747 | 92901.32043 | 281110.0035 | 1.011104 | 0.5138 | 0.5081 | 1.07461 | 0.46854 | 1.49016 | 1.0111 | 0.06714 | |
| 5-Oxoproline | C01879 | I3sg2 | C13-05 | 5 | 27589.50421 | 24664.49623 | 44874.18899 | 0.258315 | 0.096 | 0.3715 | 0.213 | 0.19339 | 0.36855 | 0.25831 | 0.06714 | |
| A | C00212 | NT | C12 PARENT | 10 | 93664.75249 | 87654.01645 | 13415.65534 | 56.07845 | 39.9354 | 0.7121 | 21.9518 | 46.2836 | 100 | 56.0784 | 0.72387 | |
| A | C00212 | I3sg2 | C12 PARENT | 10 | 46888.86549 | 249598.3752 | 190530.1793 | 47.20594 | 6.9488 | 0.1472 | 53.4233 | 39.7047 | 48.4898 | 47.2059 | 0.72387 | |
| A | C00212 | NT | C13-05 | 10 | 282342.5644 | 78804.5574 | 0 | 35.92738 | 33.4498 | 0.931 | 66.1713 | 41.6108 | 0 | 35.9274 | 0.45224 | |
| A | C00212 | I3sg2 | C13-05 | 10 | 40879.68339 | 368916.903 | 202397.9199 | 52.25735 | 6.0887 | 0.1165 | 46.5767 | 58.6852 | 51.5102 | 52.2573 | 0.45224 | |
| A | C00212 | NT | C13-06 | 10 | 50677.01512 | 22926.14921 | 0 | 7.994177 | 6.9241 | 0.8661 | 11.8769 | 12.1056 | 0 | 7.99418 | 0.13817 | |
| A | C00212 | I3sg2 | C13-06 | 10 | 0 | 10121.86951 | 0 | 0.53671 | 0.9296 | 1.7321 | 0 | 1.61013 | 0 | 0.53671 | 0.13817 | |
| a-KG | C00026 | NT | C12 PARENT | 5 | 221950.4663 | 2040182.852 | 1505859.682 | 73.51454 | 23.0306 | 0.3133 | 100 | 58.1979 | 62.3457 | 73.5145 | 0.20091 | |
| a-KG | C00026 | I3sg2 | C12 PARENT | 5 | 897901.686 | 157555.7015 | 123928.8261 | 95.15702 | 8.3883 | 0.0882 | 85.4711 | 100 | 100 | 95.157 | 0.20091 | |
| a-KG | C00026 | NT | C13-02 | 5 | 0 | 666386.777 | 441036.4304 | 12.42302 | 10.7652 | 0.8666 | 0 | 19.0092 | 18.2598 | 12.423 | 0.33725 | |
| a-KG | C00026 | I3sg2 | C13-02 | 5 | 133378.4825 | 0 | 0 | 4.232089 | 7.3302 | 1.7321 | 12.6963 | 0 | 0 | 4.23209 | 0.33725 | |
| a-KG | C00026 | NT | C13-03 | 5 | 0 | 204069.2913 | 139393.0111 | 3.864135 | 3.3465 | 0.866 | 0 | 5.82125 | 5.77116 | 3.86414 | 0.18367 | |
| a-KG | C00026 | I3sg2 | C13-03 | 5 | 19252.92448 | 0 | 0 | 0.610894 | 1.0581 | 1.7321 | 1.83268 | 0 | 0 | 0.61089 | 0.18367 | |
| a-KG | C00026 | NT | C13-04 | 5 | 0 | 372905.5368 | 174362.3419 | 5.952135 | 5.4307 | 0.9124 | 0 | 10.6374 | 7.21896 | 5.95214 | 0.13049 | |
| a-KG | C00026 | I3sg2 | C13-04 | 5 | 0 | 0 | 0 | 0 | 0 | NA | 0 | 0 | 0 | 0 | 0.13049 | |
| a-KG | C00026 | NT | C13-05 | 5 | 0 | 222048.3237 | 154687.6025 | 4.246167 | 3.6775 | 0.8661 | 0 | 6.33412 | 6.40438 | 4.24617 | 0.11613 | |
| a-KG | C00026 | I3sg2 | C13-05 | 5 | 0 | 0 | 0 | 0 | 0 | NA | 0 | 0 | 0 | 0 | 0.11613 | |
| Ac-carnitine | C02571 | NT | C12 PARENT | 9 | 8150464.274 | 10533812.82 | 9595272.951 | 64.96349 | 0.2891 | 0.0045 | 64.643 | 65.0427 | 65.2047 | 64.9635 | 7.52E-06 | *** |
| Ac-carnitine | C02571 | I3sg2 | C12 PARENT | 9 | 12128843.46 | 18575729.3 | 11616093.67 | 74.46248 | 0.4696 | 0.0063 | 74.8515 | 73.9408 | 74.5951 | 74.4625 | 7.52E-06 | *** |
| Ac-carnitine | C02571 | NT | C13-01 | 9 | 0 | 0 | 0 | 0 | 0 | NA | 0 | 0 | 0 | 0 | NA | |
| Ac-carnitine | C02571 | I3sg2 | C13-01 | 9 | 0 | 0 | 0 | 0 | 0 | NA | 0 | 0 | 0 | 0 | NA | |
| Ac-carnitine | C02571 | NT | C13-02 | 9 | 4457959.467 | 5661402.284 | 5120337.462 | 35.03651 | 0.2891 | 0.0083 | 35.357 | 34.9573 | 34.7953 | 35.0365 | 7.52E-06 | *** |
| Ac-carnitine | C02571 | I3sg2 | C13-02 | 9 | 4075028.508 | 6546694.83 | 3956100.255 | 25.53752 | 0.4696 | 0.0184 | 25.1485 | 26.0592 | 25.4049 | 25.5375 | 7.52E-06 | *** |
| Ac-carnitine | C02571 | NT | C13-03 | 9 | 0 | 0 | 0 | 0 | 0 | NA | 0 | 0 | 0 | 0 | NA | |
| Ac-carnitine | C02571 | I3sg2 | C13-03 | 9 | 0 | 0 | 0 | 0 | 0 | NA | 0 | 0 | 0 | 0 | NA | |
| Acetyl-CoA | C00024 | NT | C12 PARENT | 23 | 9517.416111 | 3373.770277 | 4204.517733 | 6.638982 | 0.3184 | 0.048 | 6.94109 | 6.66941 | 6.30645 | 6.63898 | 0.00015 | *** |
| Acetyl-CoA | C00024 | I3sg2 | C12 PARENT | 23 | 12564.41385 | 17037.49396 | 19468.52453 | 19.54804 | 1.5727 | 0.0805 | 19.6596 | 21.062 | 17.9226 | 19.548 | 0.00015 | *** |
| Acetyl-CoA | C00024 | NT | C13-01 | 23 | 0 | 0 | 0 | 0 | 0 | NA | 0 | 0 | 0 | 0 | NA | |
| Acetyl-CoA | C00024 | I3sg2 | C13-01 | 23 | 0 | 0 | 0 | 0 | 0 | NA | 0 | 0 | 0 | 0 | NA | |
| Acetyl-CoA | C00024 | NT | C13-02 | 23 | 11945.82337 | 4352.101653 | 5776.803397 | 8.660101 | 0.0545 | 0.0063 | 8.71213 | 8.60341 | 8.66476 | 8.6601 | 0.00534 | ** |
| Acetyl-CoA | C00024 | I3sg2 | C13-02 | 23 | 8631.768088 | 12051.61594 | 19169.82603 | 15.3507 | 2.1074 | 0.1373 | 13.5062 | 14.8984 | 17.6476 | 15.3507 | 0.00534 | ** |
| Acetyl-CoA | C00024 | NT | C13-03 | 23 | 688.5289125 | 0 | 423.8138992 | 0.379278 | 0.3352 | 0.8837 | 0.50215 | 0 | 0.63569 | 0.37928 | 0.13518 | |
| Acetyl-CoA | C00024 | I3sg2 | C13-03 | 23 | 983.0885913 | 1919.678362 | 522.1161192 | 1.46401 | 0.9484 | 0.6478 | 1.53825 | 2.37313 | 0.48066 | 1.46401 | 0.13518 | |
| Acetyl-CoA | C00024 | NT | C13-04 | 23 | 0 | 46.55930087 | 473.1523745 | 0.267244 | 0.3859 | 1.4441 | 0 | 0.09204 | 0.70969 | 0.26724 | 0.45704 | |
| Acetyl-CoA | C00024 | I3sg2 | C13-04 | 23 | 0 | 470.3394876 | 1407.872616 | 0.625838 | 0.6492 | 1.0373 | 0 | 0.58144 | 1.29608 | 0.62584 | 0.45704 | |
| Acetyl-CoA | C00024 | NT | C13-05 | 23 | 42622.53069 | 16339.56552 | 19101.95396 | 30.67898 | 1.8582 | 0.0606 | 31.0848 | 32.3007 | 28.6515 | 30.679 | 0.14764 | |
| Acetyl-CoA | C00024 | I3sg2 | C13-05 | 23 | 22283.46494 | 25832.60228 | 35510.43622 | 33.16409 | 1.5225 | 0.0459 | 34.8671 | 31.9346 | 32.6906 | 33.1641 | 0.14764 | |
| Acetyl-CoA | C00024 | NT | C13-06 | 23 | 11851.68769 | 2725.319166 | 4611.00851 | 6.982386 | 1.629 | 0.2333 | 8.64348 | 5.38752 | 6.91616 | 6.98239 | 0.01693 | * |
| Acetyl-CoA | C00024 | I3sg2 | C13-06 | 23 | 1868.981717 | 2438.297673 | 736.7150909 | 2.205625 | 1.3235 | 0.6001 | 2.92441 | 3.01425 | 0.67821 | 2.20562 | 0.01693 | * |
| Acetyl-CoA | C00024 | NT | C13-07 | 23 | 50169.42649 | 20302.75257 | 26790.01556 | 38.969 | 2.0615 | 0.0529 | 36.5887 | 40.1353 | 40.183 | 38.969 | 0.00086 | *** |
| Acetyl-CoA | C00024 | I3sg2 | C13-07 | 23 | 17089.60655 | 20961.60536 | 30491.8857 | 26.90792 | 1.0885 | 0.0405 | 26.7402 | 25.913 | 28.0706 | 26.9079 | 0.00086 | *** |
| Acetyl-CoA | C00024 | NT | C13-08 | 23 | 9377.814665 | 2641.986542 | 5288.818574 | 6.664961 | 1.3634 | 0.2046 | 6.83927 | 5.22279 | 7.93282 | 6.66496 | 0.0021 | ** |
| Acetyl-CoA | C00024 | I3sg2 | C13-08 | 23 | 488.4243267 | 180.6484393 | 1318.444882 | 0.73377 | 0.4959 | 0.6758 | 0.76424 | 0.22332 | 1.21375 | 0.73377 | 0.0021 | ** |
| Acetyl-CoA | C00024 | NT | C13-09 | 23 | 943.877252 | 803.7136464 | 0 | 0.759062 | 0.7968 | 1.0497 | 0.68837 | 1.58881 | 0 | 0.75906 | 0.17427 | |
| Acetyl-CoA | C00024 | I3sg2 | C13-09 | 23 | 0 | 0 | 0 | 0 | 0 | NA | 0 | 0 | 0 | 0 | 0.17427 | |
| Aconitate | C00417 | NT | C12 PARENT | 6 | 306105.6172 | 321157.8528 | 229645.8561 | 39.14358 | 2.7452 | 0.0701 | 41.9163 | 39.0877 | 36.4267 | 39.1436 | 0.06398 | |
| Aconitate | C00417 | I3sg2 | C12 PARENT | 6 | 158702.4047 | 239241.4387 | 184542.3688 | 55.53043 | 10.8318 | 0.1951 | 46.8551 | 67.6707 | 52.0654 | 55.5304 | 0.06398 | |
| Aconitate | C00417 | NT | C13-01 | 6 | 0 | 0 | 7900.366321 | 0.417722 | 0.7235 | 1.7321 | 0 | 0 | 1.25317 | 0.41772 | 0.74 | |
| Aconitate | C00417 | I3sg2 | C13-01 | 6 | 5916.819614 | 0 | 852.4398255 | 0.662459 | 0.9468 | 1.4292 | 1.74688 | 0 | 0.2405 | 0.66246 | 0.74 | |
| Aconitate | C00417 | NT | C13-02 | 6 | 184957.4537 | 283962.3207 | 206849.5504 | 30.89947 | 4.9046 | 0.1587 | 25.327 | 34.5607 | 32.8107 | 30.8995 | 0.25098 | |
| Aconitate | C00417 | I3sg2 | C13-02 | 6 | 134590.2475 | 104158.2573 | 162796.6008 | 38.37608 | 8.3181 | 0.2168 | 39.7363 | 29.4617 | 45.9302 | 38.3761 | 0.25098 | |
| Aconitate | C00417 | NT | C13-03 | 6 | 68446.26244 | 51022.50367 | 50520.54211 | 7.865381 | 1.5866 | 0.2017 | 9.37263 | 6.20989 | 8.01363 | 7.86538 | 0.02459 | * |
| Aconitate | C00417 | I3sg2 | C13-03 | 6 | 16990.69606 | 10137.78683 | 6251.943922 | 3.215907 | 1.654 | 0.5143 | 5.01632 | 2.86753 | 1.76388 | 3.21591 | 0.02459 | * |
| Aconitate | C00417 | NT | C13-04 | 6 | 86683.9243 | 95200.65809 | 90034.98055 | 12.5794 | 1.4808 | 0.1177 | 11.87 | 11.5868 | 14.2815 | 12.5794 | 0.00084 | *** |
| Aconitate | C00417 | I3sg2 | C13-04 | 6 | 9786.918692 | 0 | 0 | 0.96316 | 1.6682 | 1.7321 | 2.88948 | 0 | 0 | 0.96316 | 0.00084 | *** |
| Aconitate | C00417 | NT | C13-05 | 6 | 84085.2004 | 70290.02897 | 45481.39193 | 9.094452 | 2.2001 | 0.2419 | 11.5141 | 8.55491 | 7.21431 | 9.09445 | 0.01172 | * |
| Aconitate | C00417 | I3sg2 | C13-05 | 6 | 12721.61523 | 0 | 0 | 1.251972 | 2.1685 | 1.7321 | 3.75592 | 0 | 0 | 1.25197 | 0.01172 | * |
| Adenine | C00147 | NT | C12 PARENT | 5 | 151731.6979 | 1647103.756 | 338150.3243 | 86.15939 | 3.8904 | 0.0452 | 90.6254 | 83.506 | 84.3468 | 86.1594 | 0.0045 | ** |
| Adenine | C00147 | I3sg2 | C12 PARENT | 5 | 906301.1967 | 376169.3074 | 232806.6615 | 99.46573 | 0.9254 | 0.0093 | 98.3972 | 100 | 100 | 99.4657 | 0.0045 | ** |
| Adenine | C00147 | NT | C13-01 | 5 | 15695.69512 | 325334.5252 | 62754.25539 | 13.84061 | 3.8904 | 0.2811 | 9.37463 | 16.494 | 15.6532 | 13.8406 | 0.0045 | ** |
| Adenine | C00147 | I3sg2 | C13-01 | 5 | 14762.95296 | 0 | 0 | 0.534272 | 0.9254 | 1.7321 | 1.60281 | 0 | 0 | 0.53427 | 0.0045 | ** |
| ADP | C00008 | NT | C12 PARENT | 10 | 2729789.403 | 1532386.982 | 3064516.517 | 12.41678 | 0.6224 | 0.0501 | 11.994 | 13.1314 | 12.1249 | 12.4168 | 2.27E-05 | *** |
| ADP | C00008 | I3sg2 | C12 PARENT | 10 | 4385422.931 | 7303359.395 | 6671131.807 | 29.70394 | 1.169 | 0.0394 | 28.4193 | 30.7053 | 29.9873 | 29.7039 | 2.27E-05 | *** |
| ADP | C00008 | NT | C13-01 | 10 | 0 | 0 | 0 | 0 | 0 | NA | 0 | 0 | 0 | 0 | NA | |
| ADP | C00008 | I3sg2 | C13-01 | 10 | 0 | 0 | 0 | 0 | 0 | NA | 0 | 0 | 0 | 0 | NA | |
| ADP | C00008 | NT | C13-02 | 10 | 135128.9071 | 52414.00636 | 169663.1001 | 0.571384 | 0.1127 | 0.1973 | 0.59372 | 0.44915 | 0.67128 | 0.57138 | 0.0314 | * |
| ADP | C00008 | I3sg2 | C13-02 | 10 | 162305.1653 | 194288.5019 | 183872.2195 | 0.898388 | 0.1329 | 0.148 | 1.0518 | 0.81684 | 0.82652 | 0.89839 | 0.0314 | * |
| ADP | C00008 | NT | C13-03 | 10 | 217661.6721 | 98573.66926 | 230621.9033 | 0.904507 | 0.0562 | 0.0622 | 0.95635 | 0.8447 | 0.91247 | 0.90451 | 0.01419 | * |

| | | | | | | | | | | | | | | | | |
|---|---|---|---|---|---|---|---|---|---|---|---|---|---|---|---|---|
| ADP | C00008 | I3sg2 | C13-03 | 10 | 190129.9806 | 402722.7825 | 335001.0518 | 1.477043 | 0.2319 | 0.157 | 1.23212 | 1.69315 | 1.50586 | 1.47704 | 0.01419 | * |
| ADP | C00008 | NT | C13-04 | 10 | 385177.2014 | 208834.2685 | 497746.8222 | 1.817095 | 0.1405 | 0.0773 | 1.69236 | 1.78956 | 1.96936 | 1.8171 | 0.72143 | |
| ADP | C00008 | I3sg2 | C13-04 | 10 | 356265.8731 | 238373.4014 | 377560.6147 | 1.669365 | 0.6537 | 0.3916 | 2.30874 | 1.00218 | 1.69717 | 1.66936 | 0.72143 | |
| ADP | C00008 | NT | C13-05 | 10 | 14776533.31 | 7396534.357 | 16516605.71 | 64.55197 | 1.0345 | 0.016 | 64.9241 | 63.3829 | 65.3489 | 64.552 | 0.87876 | |
| ADP | C00008 | I3sg2 | C13-05 | 10 | 10009853.6 | 15308179.83 | 14264016.37 | 64.44846 | 0.3827 | 0.0059 | 64.8678 | 64.3596 | 64.118 | 64.4485 | 0.87876 | |
| ADP | C00008 | NT | C13-06 | 10 | 3186576.699 | 1743914.838 | 3355333.68 | 14.07354 | 0.8366 | 0.0594 | 14.001 | 14.9441 | 13.2756 | 14.0735 | 1.53E-05 | *** |
| ADP | C00008 | I3sg2 | C13-06 | 10 | 215236.4313 | 197210.3387 | 299950.5481 | 1.190748 | 0.314 | 0.2637 | 1.39482 | 0.82912 | 1.3483 | 1.19075 | 1.53E-05 | *** |
| ADP | C00008 | NT | C13-07 | 10 | 1104882.937 | 547235.7931 | 1248426.348 | 4.827813 | 0.1272 | 0.0263 | 4.85456 | 4.68941 | 4.93947 | 4.82781 | 6.22E-07 | *** |
| ADP | C00008 | I3sg2 | C13-07 | 10 | 85793.60144 | 141236.8263 | 114985.4586 | 0.555548 | 0.0385 | 0.0692 | 0.55598 | 0.5938 | 0.51687 | 0.55555 | 6.22E-07 | *** |
| ADP | C00008 | NT | C13-08 | 10 | 223953.6598 | 89705.19147 | 191594.6887 | 0.836918 | 0.1275 | 0.1523 | 0.98399 | 0.76871 | 0.75806 | 0.83692 | 0.00109 | ** |
| ADP | C00008 | I3sg2 | C13-08 | 10 | 26158.95232 | 0 | 0 | 0.056507 | 0.0979 | 1.7321 | 0.16952 | 0 | 0 | 0.05651 | 0.00109 | ** |
| ADP | C00008 | NT | C13-09 | 10 | 0 | 0 | 0 | 0 | 0 | NA | 0 | 0 | 0 | 0 | NA | |
| ADP | C00008 | I3sg2 | C13-09 | 10 | 0 | 0 | 0 | 0 | 0 | NA | 0 | 0 | 0 | 0 | NA | |
| Ala | C00041 | NT | C12 PARENT | 3 | 2091861.124 | 2097387.287 | 1895355.329 | 5.242978 | 0.1947 | 0.0371 | 5.45458 | 5.07136 | 5.203 | 5.24298 | 1.86E-05 | *** |
| Ala | C00041 | I3sg2 | C12 PARENT | 3 | 1529124.371 | 1739006.781 | 1288955.196 | 10.46162 | 0.3266 | 0.0312 | 10.1645 | 10.8113 | 10.409 | 10.4616 | 1.86E-05 | *** |
| Ala | C00041 | NT | C13-01 | 3 | 68709.45841 | 131187.5721 | 23296.4467 | 0.186772 | 0.1268 | 0.6789 | 0.17916 | 0.3172 | 0.06395 | 0.18677 | 0.09858 | |
| Ala | C00041 | I3sg2 | C13-01 | 3 | 10162.06178 | 0 | 0 | 0.022517 | 0.039 | 1.7321 | 0.06755 | 0 | 0 | 0.02252 | 0.09858 | |
| Ala | C00041 | NT | C13-02 | 3 | 507761.3836 | 342533.3653 | 441062.0488 | 1.120999 | 0.2598 | 0.2318 | 1.324 | 0.82822 | 1.21077 | 1.121 | 0.93212 | |
| Ala | C00041 | I3sg2 | C13-02 | 3 | 121292.0004 | 249115.0479 | 133700.5118 | 1.1449 | 0.3755 | 0.328 | 0.80626 | 1.54873 | 1.07971 | 1.1449 | 0.93212 | |
| Ala | C00041 | NT | C13-03 | 3 | 35682236.86 | 38786420.53 | 34068400.64 | 93.44925 | 0.3758 | 0.004 | 93.0423 | 93.7832 | 93.5223 | 93.4493 | 0.00033 | *** |
| Ala | C00041 | I3sg2 | C13-03 | 3 | 13383126.8 | 14097000.52 | 10960385.87 | 88.37096 | 0.6719 | 0.0076 | 88.9616 | 87.64 | 88.5113 | 88.371 | 0.00033 | *** |
| AMP | C00020 | NT | C12 PARENT | 10 | 1254212.957 | 427620.333 | 1802606.283 | 13.03607 | 0.6172 | 0.0473 | 12.7389 | 13.7457 | 12.6236 | 13.0361 | 2.79E-06 | *** |
| AMP | C00020 | I3sg2 | C12 PARENT | 10 | 2551120.193 | 5112252.331 | 4581734.591 | 31.13451 | 0.5396 | 0.0173 | 30.5446 | 31.2558 | 31.6031 | 31.1345 | 2.79E-06 | *** |
| AMP | C00020 | NT | C13-01 | 10 | 0 | 0 | 0 | 0 | 0 | NA | 0 | 0 | 0 | 0 | NA | |
| AMP | C00020 | I3sg2 | C13-01 | 10 | 0 | 0 | 0 | 0 | 0 | NA | 0 | 0 | 0 | 0 | NA | |
| AMP | C00020 | NT | C13-02 | 10 | 69820.9967 | 21881.11976 | 120476.5102 | 0.752073 | 0.0794 | 0.1056 | 0.70917 | 0.70336 | 0.84369 | 0.75207 | 0.23989 | |
| AMP | C00020 | I3sg2 | C13-02 | 10 | 63639.30566 | 139147.5195 | 157250.1952 | 0.899114 | 0.1667 | 0.1854 | 0.76195 | 0.85073 | 1.08465 | 0.89911 | 0.23989 | |
| AMP | C00020 | NT | C13-03 | 10 | 80355.18213 | 19091.79762 | 138053.2444 | 0.79888 | 0.1772 | 0.2218 | 0.81616 | 0.6137 | 0.96678 | 0.79888 | 0.00274 | ** |
| AMP | C00020 | I3sg2 | C13-03 | 10 | 129753.0234 | 281019.3412 | 218707.7685 | 1.593409 | 0.1103 | 0.0692 | 1.55354 | 1.71812 | 1.50857 | 1.59341 | 0.00274 | ** |
| AMP | C00020 | NT | C13-04 | 10 | 141595.762 | 51596.15483 | 191539.6481 | 1.479353 | 0.1626 | 0.1099 | 1.43817 | 1.65854 | 1.34135 | 1.47935 | 0.08118 | |
| AMP | C00020 | I3sg2 | C13-04 | 10 | 96076.49732 | 204131.8425 | 189994.7165 | 1.236294 | 0.0807 | 0.0653 | 1.15033 | 1.24804 | 1.31051 | 1.23629 | 0.08118 | |
| AMP | C00020 | NT | C13-05 | 10 | 6507803.713 | 2059824.748 | 9546683.042 | 66.38882 | 0.4077 | 0.0061 | 66.0991 | 66.2122 | 66.8551 | 66.3888 | 0.00496 | ** |
| AMP | C00020 | I3sg2 | C13-05 | 10 | 5385143.714 | 10395070.85 | 9139827.588 | 63.69135 | 0.7263 | 0.0114 | 64.4764 | 63.5544 | 63.0432 | 63.6913 | 0.00496 | ** |
| AMP | C00020 | NT | C13-06 | 10 | 1306447.76 | 380050.7118 | 1798695.969 | 12.69408 | 0.5332 | 0.042 | 13.2695 | 12.2166 | 12.5962 | 12.6941 | 3.04E-06 | *** |
| AMP | C00020 | I3sg2 | C13-06 | 10 | 86069.35884 | 142880.2632 | 116786.967 | 0.903207 | 0.1154 | 0.1277 | 1.03051 | 0.87356 | 0.80555 | 0.90321 | 3.04E-06 | *** |
| AMP | C00020 | NT | C13-07 | 10 | 371280.145 | 123143.2255 | 580866.1512 | 3.932411 | 0.1501 | 0.0382 | 3.77106 | 3.95839 | 4.06779 | 3.93241 | 3.80E-06 | *** |
| AMP | C00020 | I3sg2 | C13-07 | 10 | 40312.3216 | 78281.87583 | 87981.00993 | 0.52271 | 0.0729 | 0.1395 | 0.48266 | 0.47861 | 0.60686 | 0.52271 | 3.80E-06 | *** |
| AMP | C00020 | NT | C13-08 | 10 | 114003.6669 | 27734.57707 | 100741.9219 | 0.918311 | 0.2274 | 0.2476 | 1.15792 | 0.89152 | 0.70549 | 0.91831 | 0.00241 | ** |
| AMP | C00020 | I3sg2 | C13-08 | 10 | 0 | 3389.99559 | 5433.87439 | 0.019402 | 0.0188 | 0.9677 | 0 | 0.02073 | 0.03748 | 0.0194 | 0.00241 | ** |
| Arg-Succ | C03406 | NT | C12 PARENT | 10 | 154869.1345 | 121329.099 | 132188.2529 | 74.11638 | 9.4032 | 0.1269 | 65.3486 | 72.9537 | 84.0469 | 74.1164 | 0.03572 | * |
| Arg-Succ | C03406 | I3sg2 | C12 PARENT | 10 | 109555.3749 | 97272.36 | 89336.49916 | 93.76109 | 5.5617 | 0.0593 | 91.9605 | 100 | 89.3228 | 93.7611 | 0.03572 | * |
| Arg-Succ | C03406 | NT | C13-01 | 10 | 0 | 0 | 0 | 0 | 0 | NA | 0 | 0 | 0 | 0 | NA | |
| Arg-Succ | C03406 | I3sg2 | C13-01 | 10 | 0 | 0 | 0 | 0 | 0 | NA | 0 | 0 | 0 | 0 | NA | |
| Arg-Succ | C03406 | NT | C13-02 | 10 | 59486.2517 | 37155.74661 | 17774.90698 | 19.5812 | 7.302 | 0.3729 | 25.1008 | 22.3413 | 11.3015 | 19.5812 | 0.06552 | |
| Arg-Succ | C03406 | I3sg2 | C13-02 | 10 | 9577.742671 | 0 | 10678.84233 | 6.238911 | 5.5617 | 0.8915 | 8.03953 | 0 | 10.6772 | 6.23891 | 0.06552 | |
| Arg-Succ | C03406 | NT | C13-03 | 10 | 22633.90388 | 7824.963914 | 7315.981398 | 6.302415 | 2.8131 | 0.4464 | 9.5506 | 4.70505 | 4.65159 | 6.30241 | 0.01784 | * |
| Arg-Succ | C03406 | I3sg2 | C13-03 | 10 | 0 | 0 | 0 | 0 | 0 | NA | 0 | 0 | 0 | 0 | 0.01784 | * |
| Asn | C00152 | NT | C12 PARENT | 4 | 2780582.06 | 2527489.417 | 2408666.759 | 70.05004 | 0.4372 | 0.0062 | 69.8363 | 70.553 | 69.7608 | 70.05 | 0.00073 | *** |
| Asn | C00152 | I3sg2 | C12 PARENT | 4 | 1951541.419 | 2320789.327 | 2136092.36 | 84.54039 | 2.6468 | 0.0313 | 86.4596 | 81.5209 | 85.6406 | 84.5404 | 0.00073 | *** |
| Asn | C00152 | NT | C13-01 | 4 | 60378.09884 | 88608.12795 | 57054.03704 | 1.880764 | 0.5177 | 0.2753 | 1.51644 | 2.47343 | 1.65242 | 1.88076 | 0.00326 | ** |
| Asn | C00152 | I3sg2 | C13-01 | 4 | 0 | 0 | 0 | 0 | 0 | NA | 0 | 0 | 0 | 0 | 0.00326 | ** |
| Asn | C00152 | NT | C13-02 | 4 | 603494.3708 | 454848.7177 | 430201.5251 | 13.43788 | 1.4937 | 0.1112 | 15.1572 | 12.6968 | 12.4597 | 13.4379 | 0.00532 | ** |
| Asn | C00152 | I3sg2 | C13-02 | 4 | 194940.7493 | 191852.1134 | 199293.3197 | 7.788561 | 0.9646 | 0.1239 | 8.6365 | 6.73907 | 7.99011 | 7.78856 | 0.00532 | ** |
| Asn | C00152 | NT | C13-03 | 4 | 417959.7235 | 363634.1368 | 430798.4371 | 11.04163 | 1.2551 | 0.1137 | 10.4973 | 10.1506 | 12.477 | 11.0416 | 0.03321 | * |
| Asn | C00152 | I3sg2 | C13-03 | 4 | 110689.756 | 251785.4531 | 158865.06 | 6.705826 | 1.9916 | 0.297 | 4.90391 | 8.84431 | 6.36925 | 6.70583 | 0.03321 | * |
| Asn | C00152 | NT | C13-04 | 4 | 119159.5891 | 147815.4756 | 126029.1692 | 3.589683 | 0.5691 | 0.1585 | 2.99278 | 4.12616 | 3.65011 | 3.58968 | 0.06172 | |
| Asn | C00152 | I3sg2 | C13-04 | 4 | 0 | 82435.62517 | 0 | 0.965222 | 1.6718 | 1.7321 | 0 | 2.89567 | 0 | 0.96522 | 0.06172 | |
| Asp | C00049 | NT | C12 PARENT | 4 | 32154892.03 | 24542365.12 | 27877575 | 60.40205 | 0.7794 | 0.0129 | 59.5763 | 60.5049 | 61.1249 | 60.402 | 0.00176 | ** |
| Asp | C00049 | I3sg2 | C12 PARENT | 4 | 27165058.1 | 33319660 | 30769846.5 | 66.01317 | 1.0518 | 0.0159 | 64.8543 | 66.2777 | 66.9075 | 66.0132 | 0.00176 | ** |
| Asp | C00049 | NT | C13-01 | 4 | 1913663.24 | 1528256.638 | 1481734.922 | 3.520724 | 0.2603 | 0.0739 | 3.54562 | 3.76765 | 3.24888 | 3.52072 | 0.40263 | |
| Asp | C00049 | I3sg2 | C13-01 | 4 | 1540355.686 | 1914619.888 | 1631559.476 | 3.677892 | 0.1304 | 0.0354 | 3.67747 | 3.80846 | 3.54774 | 3.67789 | 0.40263 | |
| Asp | C00049 | NT | C13-02 | 4 | 9601040.366 | 6993917.662 | 7772719.579 | 17.35788 | 0.3863 | 0.0223 | 17.7887 | 17.2423 | 17.0426 | 17.3579 | 0.00937 | ** |
| Asp | C00049 | I3sg2 | C13-02 | 4 | 6813391.504 | 7879825.78 | 7094103.767 | 15.78877 | 0.4319 | 0.0274 | 16.2664 | 15.6741 | 15.4258 | 15.7888 | 0.00937 | ** |
| Asp | C00049 | NT | C13-03 | 4 | 7326864.243 | 5248642.084 | 6027509.461 | 13.2436 | 0.3187 | 0.0241 | 13.5752 | 12.9396 | 13.216 | 13.2436 | 0.00146 | ** |
| Asp | C00049 | I3sg2 | C13-03 | 4 | 4741028.854 | 5311437.832 | 4859928.939 | 10.81724 | 0.4344 | 0.0402 | 11.3188 | 10.5652 | 10.5677 | 10.8172 | 0.00146 | ** |
| Asp | C00049 | NT | C13-04 | 4 | 2976151.084 | 2249407.875 | 2448016.15 | 5.475759 | 0.095 | 0.0173 | 5.51419 | 5.54552 | 5.36757 | 5.47576 | 9.06E-05 | *** |
| Asp | C00049 | I3sg2 | C13-04 | 4 | 1626421.33 | 1847271.5 | 1633223.763 | 3.702934 | 0.1676 | 0.0453 | 3.88295 | 3.67449 | 3.55136 | 3.70293 | 9.06E-05 | *** |
| ATP | C00002 | NT | C12 PARENT | 10 | 11153490.97 | 3874541.05 | 7495432.742 | 11.56215 | 0.1926 | 0.0167 | 11.7839 | 11.4661 | 11.4364 | 11.5621 | 1.92E-06 | *** |
| ATP | C00002 | I3sg2 | C12 PARENT | 10 | 6258655.559 | 14406957.78 | 12312314.43 | 30.31252 | 0.7489 | 0.0247 | 29.7069 | 31.1499 | 30.0807 | 30.3125 | 1.92E-06 | *** |
| ATP | C00002 | NT | C13-01 | 10 | 0 | 0 | 0 | 0 | 0 | NA | 0 | 0 | 0 | 0 | NA | |
| ATP | C00002 | I3sg2 | C13-01 | 10 | 0 | 0 | 0 | 0 | 0 | NA | 0 | 0 | 0 | 0 | NA | |
| ATP | C00002 | NT | C13-02 | 10 | 699646.2902 | 256153.5216 | 402607.9838 | 0.703844 | 0.0781 | 0.111 | 0.73919 | 0.75804 | 0.61429 | 0.70384 | 0.10953 | |
| ATP | C00002 | I3sg2 | C13-02 | 10 | 249379.9813 | 407553.2853 | 330498.0517 | 0.957444 | 0.1994 | 0.2082 | 1.18369 | 0.88119 | 0.80745 | 0.95744 | 0.10953 | |
| ATP | C00002 | NT | C13-03 | 10 | 835495.2805 | 319972.2922 | 640172.0721 | 0.935464 | 0.0481 | 0.0514 | 0.88272 | 0.9469 | 0.97677 | 0.93546 | 0.001 | ** |
| ATP | C00002 | I3sg2 | C13-03 | 10 | 295501.0683 | 734766.8487 | 590388.1823 | 1.477893 | 0.098 | 0.0663 | 1.4026 | 1.58867 | 1.4424 | 1.47789 | 0.001 | ** |
| ATP | C00002 | NT | C13-04 | 10 | 1061134.932 | 426067.5685 | 534761.9677 | 1.065975 | 0.2275 | 0.2135 | 1.12112 | 1.26088 | 0.81593 | 1.06597 | 0.20709 | |
| ATP | C00002 | I3sg2 | C13-04 | 10 | 387879.6937 | 719941.0638 | 420667.9221 | 1.47515 | 0.4127 | 0.2798 | 1.84108 | 1.55662 | 1.02775 | 1.47515 | 0.20709 | |
| ATP | C00002 | NT | C13-05 | 10 | 62464830.12 | 22473777.37 | 43231154.58 | 66.15487 | 0.3058 | 0.0046 | 65.9956 | 66.5074 | 65.9616 | 66.1549 | 0.0297 | * |
| ATP | C00002 | I3sg2 | C13-05 | 10 | 13331602.21 | 28854877.07 | 26604068.2 | 63.55493 | 1.3263 | 0.0209 | 63.2788 | 62.3884 | 64.9975 | 63.5549 | 0.0297 | * |
| ATP | C00002 | NT | C13-06 | 10 | 12839718.59 | 4441562.042 | 9474774.348 | 13.72201 | 0.6701 | 0.0488 | 13.5655 | 13.1441 | 14.4565 | 13.722 | 1.13E-05 | *** |
| ATP | C00002 | I3sg2 | C13-06 | 10 | 394247.4485 | 810402.3102 | 460660.2835 | 1.582991 | 0.4007 | 0.2531 | 1.87131 | 1.75221 | 1.12546 | 1.58299 | 1.13E-05 | *** |
| ATP | C00002 | NT | C13-07 | 10 | 4685476.471 | 1653118.543 | 2985937.517 | 4.799452 | 0.2129 | 0.0444 | 4.95032 | 4.89213 | 4.55591 | 4.79945 | 8.07E-06 | *** |
| ATP | C00002 | I3sg2 | C13-07 | 10 | 150758.1683 | 310844.682 | 196851.7011 | 0.622869 | 0.1248 | 0.2004 | 0.71558 | 0.67209 | 0.48094 | 0.62287 | 8.07E-06 | *** |
| ATP | C00002 | NT | C13-08 | 10 | 910161.7893 | 346193.4559 | 775086.1753 | 1.056242 | 0.1139 | 0.1078 | 0.96161 | 1.0245 | 1.18262 | 1.05624 | 9.87E-05 | *** |
| ATP | C00002 | I3sg2 | C13-08 | 10 | 0 | 5050.091391 | 15431.66565 | 0.016207 | 0.0194 | 1.197 | 0 | 0.01092 | 0.0377 | 0.01621 | 9.87E-05 | *** |
| ATP | C00002 | NT | C13-09 | 10 | 0 | 0 | 0 | 0 | 0 | NA | 0 | 0 | 0 | 0 | NA | |
| ATP | C00002 | I3sg2 | C13-09 | 10 | 0 | 0 | 0 | 0 | 0 | NA | 0 | 0 | 0 | 0 | NA | |
| Carbamoyl-Asp | C00438 | NT | C12 PARENT | 5 | 1264941.829 | 1130655.475 | 865836.8403 | 75.48427 | 0.2678 | 0.0035 | 75.301 | 75.7916 | 75.3603 | 75.4843 | 0.03259 | * |
| Carbamoyl-Asp | C00438 | I3sg2 | C12 PARENT | 5 | 963095.9355 | 1253559.578 | 1051866.744 | 69.66539 | 3.1285 | 0.0449 | 66.2067 | 70.4917 | 72.2978 | 69.6654 | 0.03259 | * |
| Carbamoyl-Asp | C00438 | NT | C13-01 | 5 | 0 | 0 | 0 | 0 | 0 | NA | 0 | 0 | 0 | 0 | NA | |
| Carbamoyl-Asp | C00438 | I3sg2 | C13-01 | 5 | 0 | 0 | 0 | 0 | 0 | NA | 0 | 0 | 0 | 0 | NA | |
| Carbamoyl-Asp | C00438 | NT | C13-03 | 5 | 414906.0013 | 361140.7128 | 283093.0542 | 24.51573 | 0.2678 | 0.0109 | 24.699 | 24.2084 | 24.6397 | 24.5157 | 0.03259 | * |
| Carbamoyl-Asp | C00438 | I3sg2 | C13-03 | 5 | 491584.2054 | 524748.7022 | 403042.3287 | 30.33461 | 3.1285 | 0.1031 | 33.7933 | 29.5083 | 27.7022 | 30.3346 | 0.03259 | * |

| | | | | | | | | | | | | | | | | |
|---|---|---|---|---|---|---|---|---|---|---|---|---|---|---|---|---|
| CDP | C00112 | NT | C12 PARENT | 9 | 342163.5467 | 142841.224 | 410215.4535 | 21.10738 | 3.0952 | 0.1466 | 17.5984 | 23.4497 | 22.274 | 21.1074 | 5.30E-06 | *** |
| CDP | C00112 | I3sg2 | C12 PARENT | 9 | 1037198.056 | 1974211.838 | 1922720.942 | 81.62394 | 0.8824 | 0.0108 | 81.4436 | 80.8456 | 82.5826 | 81.6239 | 5.30E-06 | *** |
| CDP | C00112 | NT | C13-04 | 9 | 21175.25282 | 0 | 3391.990307 | 0.424427 | 0.5829 | 1.3735 | 1.0891 | 0 | 0.18418 | 0.42443 | 0.27582 | |
| CDP | C00112 | I3sg2 | C13-04 | 9 | 0 | 0 | 0 | 0 | 0 | NA | 0 | 0 | 0 | 0 | 0.27582 | |
| CDP | C00112 | NT | C13-05 | 9 | 972910.9486 | 324689.0263 | 916896.3911 | 51.0428 | 1.9615 | 0.0384 | 50.0395 | 53.303 | 49.7859 | 51.0428 | 2.80E-05 | *** |
| CDP | C00112 | I3sg2 | C13-05 | 9 | 226952.6521 | 429290.4146 | 332477.235 | 16.56031 | 1.9783 | 0.1195 | 17.8209 | 17.5798 | 14.2802 | 16.5603 | 2.80E-05 | *** |
| CDP | C00112 | NT | C13-06 | 9 | 142241.7651 | 24284.8352 | 105696.1001 | 5.680585 | 1.6653 | 0.2932 | 7.31589 | 3.98675 | 5.73912 | 5.68059 | 0.0047 | ** |
| CDP | C00112 | I3sg2 | C13-06 | 9 | 0 | 0 | 10053.84008 | 0.14394 | 0.2493 | 1.7321 | 0 | 0 | 0.43182 | 0.14394 | 0.0047 | ** |
| CDP | C00112 | NT | C13-07 | 9 | 296415.4183 | 92937.10611 | 247893.2693 | 14.65426 | 1.0341 | 0.0706 | 15.2455 | 15.2571 | 13.4602 | 14.6543 | 0.00011 | *** |
| CDP | C00112 | I3sg2 | C13-07 | 9 | 9365.669884 | 20596.69446 | 62987.87546 | 1.428085 | 1.1075 | 0.7755 | 0.73542 | 0.84345 | 2.70539 | 1.42809 | 0.00011 | *** |
| CDP | C00112 | NT | C13-08 | 9 | 169379.2804 | 24386.35081 | 157584.9338 | 7.090551 | 2.6747 | 0.3772 | 8.71164 | 4.00342 | 8.55659 | 7.09055 | 0.01188 | * |
| CDP | C00112 | I3sg2 | C13-08 | 9 | 0 | 17855.13451 | 0 | 0.243727 | 0.4221 | 1.7321 | 0 | 0.73118 | 0 | 0.24373 | 0.01188 | * |
| CDP-choline | C00307 | NT | C12 PARENT | 14 | 51199.42216 | 34520.61545 | 51263.66913 | 45.13635 | 4.7334 | 0.1049 | 50.316 | 41.0355 | 44.0576 | 45.1364 | 8.60E-05 | *** |
| CDP-choline | C00307 | I3sg2 | C12 PARENT | 14 | 252003.3265 | 315878.1802 | 257904.9199 | 90.22573 | 0.9872 | 0.0109 | 89.1182 | 91.0131 | 90.5459 | 90.2257 | 8.60E-05 | *** |
| CDP-choline | C00307 | NT | C13-05 | 14 | 47230.61305 | 47362.33117 | 57974.32964 | 50.84711 | 5.0212 | 0.0988 | 46.4157 | 56.3008 | 49.8249 | 50.8471 | 0.00016 | *** |
| CDP-choline | C00307 | I3sg2 | C13-05 | 14 | 30771.04361 | 31190.80575 | 26928.27598 | 9.774268 | 0.9872 | 0.101 | 10.8818 | 8.98692 | 9.45405 | 9.77427 | 0.00016 | *** |
| CDP-choline | C00307 | NT | C13-06 | 14 | 1435.212739 | 0 | 2101.795659 | 1.072265 | 0.9495 | 0.8855 | 1.41045 | 0 | 1.80635 | 1.07227 | 0.12211 | |
| CDP-choline | C00307 | I3sg2 | C13-06 | 14 | 0 | 0 | 0 | 0 | 0 | NA | 0 | 0 | 0 | 0 | 0.12211 | |
| CDP-choline | C00307 | NT | C13-07 | 14 | 1890.49908 | 2240.840415 | 5016.34876 | 2.944274 | 1.2505 | 0.4247 | 1.85788 | 2.66374 | 4.3112 | 2.94427 | 0.01512 | * |
| CDP-choline | C00307 | I3sg2 | C13-07 | 14 | 0 | 0 | 0 | 0 | 0 | NA | 0 | 0 | 0 | 0 | 0.01512 | * |
| CDP-EtA | C00570 | NT | C12 PARENT | 11 | 314941.3469 | 250842.9926 | 303881.9018 | 26.72105 | 2.6981 | 0.101 | 29.7952 | 25.6217 | 24.7462 | 26.721 | 8.21E-06 | *** |
| CDP-EtA | C00570 | I3sg2 | C12 PARENT | 11 | 1140422.223 | 1206247.884 | 1258452.954 | 86.10185 | 2.2675 | 0.0263 | 83.5117 | 87.7284 | 87.0655 | 86.1018 | 8.21E-06 | *** |
| CDP-EtA | C00570 | NT | C13-05 | 11 | 498388.0796 | 458125.7546 | 612544.8277 | 47.94205 | 1.6892 | 0.0352 | 47.1504 | 46.7941 | 49.8817 | 47.942 | 2.07E-05 | *** |
| CDP-EtA | C00570 | I3sg2 | C13-05 | 11 | 209205.8017 | 159116.0998 | 177326.611 | 13.05346 | 1.9934 | 0.1527 | 15.3199 | 11.5722 | 12.2683 | 13.0535 | 2.07E-05 | *** |
| CDP-EtA | C00570 | NT | C13-06 | 11 | 75252.28874 | 58567.64887 | 94056.45694 | 6.920297 | 0.8561 | 0.1237 | 7.1193 | 5.98224 | 7.65935 | 6.9203 | 0.00015 | *** |
| CDP-EtA | C00570 | I3sg2 | C13-06 | 11 | 0 | 0 | 0 | 0 | 0 | NA | 0 | 0 | 0 | 0 | 0.00015 | *** |
| CDP-EtA | C00570 | NT | C13-07 | 11 | 126283.0802 | 142637.1013 | 155509.2741 | 13.06002 | 1.3553 | 0.1038 | 11.9471 | 14.5693 | 12.6637 | 13.06 | 9.32E-05 | *** |
| CDP-EtA | C00570 | I3sg2 | C13-07 | 11 | 5501.689553 | 9616.189097 | 9630.352145 | 0.589507 | 0.1625 | 0.2756 | 0.40288 | 0.69937 | 0.66627 | 0.58951 | 9.32E-05 | *** |
| CDP-EtA | C00570 | NT | C13-08 | 11 | 42154.01349 | 68851.45584 | 62002.53588 | 5.356584 | 1.5454 | 0.2885 | 3.98801 | 7.03266 | 5.04909 | 5.35658 | 0.00534 | ** |
| CDP-EtA | C00570 | I3sg2 | C13-08 | 11 | 10454.15832 | 0 | 0 | 0.255182 | 0.442 | 1.7321 | 0.76554 | 0 | 0 | 0.25518 | 0.00534 | ** |
| Cit | C00158 | NT | C12 PARENT | 6 | 47066543.3 | 38357352.27 | 40888529.53 | 33.31976 | 1.553 | 0.0466 | 32.9101 | 35.0365 | 32.0127 | 33.3198 | 0.00237 | ** |
| Cit | C00158 | I3sg2 | C12 PARENT | 6 | 31344292.04 | 37067381.11 | 31842270.85 | 41.03751 | 1.1792 | 0.0287 | 41.1984 | 42.128 | 39.7862 | 41.0375 | 0.00237 | ** |
| Cit | C00158 | NT | C13-01 | 6 | 2679997.995 | 2347067.565 | 2331274.55 | 1.947667 | 0.1716 | 0.0881 | 1.87392 | 2.14386 | 1.82521 | 1.94767 | 0.03425 | * |
| Cit | C00158 | I3sg2 | C13-01 | 6 | 2286428.182 | 2289109.852 | 1861924.099 | 2.644433 | 0.3414 | 0.1291 | 3.00524 | 2.60163 | 2.32643 | 2.64443 | 0.03425 | * |
| Cit | C00158 | NT | C13-02 | 6 | 48486708.79 | 35869266.87 | 46288822.54 | 34.30254 | 1.7725 | 0.0517 | 33.9032 | 32.7638 | 36.2407 | 34.3025 | 0.15639 | |
| Cit | C00158 | I3sg2 | C13-02 | 6 | 23870961.18 | 27619505.41 | 26868667.13 | 32.11253 | 1.2638 | 0.0394 | 31.3756 | 31.3902 | 33.5718 | 32.1125 | 0.15639 | |
| Cit | C00158 | NT | C13-03 | 6 | 9568294.807 | 7308421.847 | 8268168.529 | 6.613144 | 0.1213 | 0.0183 | 6.6904 | 6.67568 | 6.47336 | 6.61314 | 0.91247 | |
| Cit | C00158 | I3sg2 | C13-03 | 6 | 5135351.296 | 5757709.101 | 5210036.706 | 6.601137 | 0.1299 | 0.0197 | 6.74981 | 6.54378 | 6.50982 | 6.60114 | 0.91247 | |
| Cit | C00158 | NT | C13-04 | 6 | 18014539.78 | 13019728.79 | 15488945.41 | 12.20514 | 0.3584 | 0.0294 | 12.5962 | 11.8925 | 12.1267 | 12.2051 | 0.0005 | *** |
| Cit | C00158 | I3sg2 | C13-04 | 6 | 7604973.012 | 8475481.755 | 7738016.032 | 9.765634 | 0.2002 | 0.0205 | 9.99584 | 9.63259 | 9.66847 | 9.76563 | 0.0005 | *** |
| Cit | C00158 | NT | C13-05 | 6 | 12970655.64 | 9698770.316 | 10829165.65 | 8.802306 | 0.2996 | 0.034 | 9.06942 | 8.85907 | 8.47843 | 8.80231 | 0.00033 | *** |
| Cit | C00158 | I3sg2 | C13-05 | 6 | 4593304.519 | 5451387.961 | 5212011.484 | 6.248427 | 0.2418 | 0.0387 | 6.03736 | 6.19563 | 6.51229 | 6.24843 | 0.00033 | *** |
| Cit | C00158 | NT | C13-06 | 6 | 4228560.597 | 2877782.661 | 3631214.289 | 2.80944 | 0.1666 | 0.0593 | 2.95672 | 2.62863 | 2.84297 | 2.80944 | 0.00031 | *** |
| Cit | C00158 | I3sg2 | C13-06 | 6 | 1246087.251 | 1326987.414 | 1300548.144 | 1.590331 | 0.0715 | 0.0449 | 1.63783 | 1.50815 | 1.62501 | 1.59033 | 0.00031 | *** |
| CMP | C00055 | NT | C12 PARENT | 9 | 20159.83598 | 12895.17299 | 21926.27547 | 20.02868 | 8.1776 | 0.4083 | 20.7754 | 27.8073 | 11.5033 | 20.0287 | 0.00019 | *** |
| CMP | C00055 | I3sg2 | C12 PARENT | 9 | 231756.445 | 257536.5691 | 338333.5722 | 90.27715 | 4.2335 | 0.0469 | 93.3647 | 85.4512 | 92.0156 | 90.2771 | 0.00019 | *** |
| CMP | C00055 | NT | C13-05 | 9 | 76877.03314 | 33478.10048 | 168682.676 | 79.97132 | 8.1776 | 0.1023 | 79.2246 | 72.1927 | 88.4967 | 79.9713 | 0.00019 | *** |
| CMP | C00055 | I3sg2 | C13-05 | 9 | 16470.69004 | 43847.90852 | 29357.97245 | 9.722853 | 4.2335 | 0.4354 | 6.63533 | 14.5488 | 7.9844 | 9.72285 | 0.00019 | *** |
| CoA | C00010 | NT | C12 PARENT | 21 | 72274.224 | 31411.856 | 51889.67099 | 24.2901 | 5.1202 | 0.2108 | 21.4372 | 30.2012 | 21.2319 | 24.2901 | 0.0031 | ** |
| CoA | C00010 | I3sg2 | C12 PARENT | 21 | 42100.59733 | 74496.73368 | 83944.1265 | 45.93972 | 2.8922 | 0.063 | 46.9564 | 42.6765 | 48.1862 | 45.9397 | 0.0031 | ** |
| CoA | C00010 | NT | C13-01 | 21 | 0 | 0 | 0 | 0 | 0 | NA | 0 | 0 | 0 | 0 | NA | |
| CoA | C00010 | I3sg2 | C13-01 | 21 | 0 | 0 | 0 | 0 | 0 | NA | 0 | 0 | 0 | 0 | NA | |
| CoA | C00010 | NT | C13-02 | 21 | 1479.812883 | 0 | 671.1277592 | 0.237845 | 0.2218 | 0.9324 | 0.43893 | 0 | 0.27461 | 0.23784 | 0.13677 | |
| CoA | C00010 | I3sg2 | C13-02 | 21 | 0 | 0 | 0 | 0 | 0 | NA | 0 | 0 | 0 | 0 | 0.13677 | |
| CoA | C00010 | NT | C13-03 | 21 | 2018.328003 | 912.6349812 | 2633.392324 | 0.851211 | 0.2405 | 0.2825 | 0.59866 | 0.87746 | 1.07752 | 0.85121 | 0.8068 | |
| CoA | C00010 | I3sg2 | C13-03 | 21 | 476.6144711 | 2407.309646 | 654.1319527 | 0.762046 | 0.54 | 0.7086 | 0.53159 | 1.37906 | 0.37549 | 0.76205 | 0.8068 | |
| CoA | C00010 | NT | C13-04 | 21 | 5022.036109 | 1502.796088 | 3348.834878 | 1.434905 | 0.0603 | 0.042 | 1.48958 | 1.44487 | 1.37026 | 1.43491 | 0.20577 | |
| CoA | C00010 | I3sg2 | C13-04 | 21 | 2458.733408 | 3415.434371 | 2419.839705 | 2.029318 | 0.6796 | 0.3349 | 2.74232 | 1.95658 | 1.38905 | 2.02932 | 0.20577 | |
| CoA | C00010 | NT | C13-05 | 21 | 209829.9316 | 60616.66312 | 152378.3528 | 60.95574 | 2.3176 | 0.038 | 62.2375 | 58.2803 | 62.3493 | 60.9557 | 0.00655 | ** |
| CoA | C00010 | I3sg2 | C13-05 | 21 | 43381.93806 | 93219.60993 | 87189.53723 | 50.6123 | 2.5553 | 0.0505 | 48.3856 | 53.4021 | 50.0492 | 50.6123 | 0.00655 | ** |
| CoA | C00010 | NT | C13-06 | 21 | 41029.79182 | 8455.921928 | 27287.60631 | 10.48841 | 2.1033 | 0.2005 | 12.1698 | 8.13001 | 11.1654 | 10.4884 | 0.00108 | ** |
| CoA | C00010 | I3sg2 | C13-06 | 21 | 0 | 847.3559085 | 0 | 0.161806 | 0.2803 | 1.7321 | 0 | 0.48542 | 0 | 0.16181 | 0.00108 | ** |
| CoA | C00010 | NT | C13-07 | 21 | 4086.45016 | 361.4863643 | 2881.432794 | 0.912881 | 0.4899 | 0.5366 | 1.21208 | 0.34755 | 1.17901 | 0.91288 | 0.03204 | * |
| CoA | C00010 | I3sg2 | C13-07 | 21 | 0 | 0 | 0 | 0 | 0 | NA | 0 | 0 | 0 | 0 | 0.03204 | * |
| CoA | C00010 | NT | C13-08 | 21 | 1403.057867 | 747.399524 | 3304.147519 | 0.828909 | 0.4776 | 0.5761 | 0.41616 | 0.71859 | 1.35197 | 0.82891 | 0.55841 | |
| CoA | C00010 | I3sg2 | C13-08 | 21 | 1240.979794 | 175.1421288 | 0 | 0.494815 | 0.7718 | 1.5597 | 1.38411 | 0.10033 | 0 | 0.49482 | 0.55841 | |
| CoA | C00010 | NT | C13-09 | 21 | 0 | 0 | 0 | 0 | 0 | NA | 0 | 0 | 0 | 0 | NA | |
| CoA | C00010 | I3sg2 | C13-09 | 21 | 0 | 0 | 0 | 0 | 0 | NA | 0 | 0 | 0 | 0 | NA | |
| Creatine | C00300 | NT | C12 PARENT | 4 | 410586112.7 | 384101393.7 | 436355417.9 | 95.10211 | 0.0773 | 8.00E-04 | 95.0265 | 95.0988 | 95.1811 | 95.1021 | 6.55E-05 | *** |
| Creatine | C00300 | I3sg2 | C12 PARENT | 4 | 188656230.8 | 455167049.5 | 217165311.9 | 98.57757 | 0.3392 | 0.0034 | 98.9544 | 98.2964 | 98.4819 | 98.5776 | 6.55E-05 | *** |
| Creatine | C00300 | NT | C13-01 | 4 | 0 | 0 | 0 | 0 | 0 | NA | 0 | 0 | 0 | 0 | NA | |
| Creatine | C00300 | I3sg2 | C13-01 | 4 | 0 | 0 | 0 | 0 | 0 | NA | 0 | 0 | 0 | 0 | NA | |
| Creatine | C00300 | NT | C13-02 | 4 | 21489261.88 | 19795914.3 | 22092292.02 | 4.897886 | 0.0773 | 0.0158 | 4.9735 | 4.90122 | 4.81893 | 4.89789 | 6.55E-05 | *** |
| Creatine | C00300 | I3sg2 | C13-02 | 4 | 1993477.484 | 7888507.789 | 3347587.543 | 1.422431 | 0.3392 | 0.2385 | 1.04562 | 1.70358 | 1.51809 | 1.42243 | 6.55E-05 | *** |
| Creatinine | C00791 | NT | C12 PARENT | 4 | 569206.588 | 937646.3849 | 932680.59 | 95.80381 | 0.3667 | 0.0038 | 95.417 | 96.1464 | 95.848 | 95.8038 | 0.05403 | |
| Creatinine | C00791 | I3sg2 | C12 PARENT | 4 | 429436.236 | 847121.2068 | 544106.5221 | 98.07871 | 1.4119 | 0.0144 | 96.9076 | 99.6465 | 97.6821 | 98.0787 | 0.05403 | |
| Creatinine | C00791 | NT | C13-01 | 4 | 8005.006191 | 12777.48972 | 14699.26516 | 1.387562 | 0.1077 | 0.0776 | 1.34189 | 1.31021 | 1.51059 | 1.38756 | 0.54946 | |
| Creatinine | C00791 | I3sg2 | C13-01 | 4 | 13703.72138 | 3005.610276 | 12911.1754 | 1.921291 | 1.4119 | 0.7348 | 3.09241 | 0.35355 | 2.31791 | 1.92129 | 0.54946 | |
| Creatinine | C00791 | NT | C13-02 | 4 | 19334.57361 | 24804.29448 | 25702.73853 | 2.808631 | 0.3777 | 0.1345 | 3.24109 | 2.54343 | 2.64137 | 2.80863 | 0.00021 | *** |
| Creatinine | C00791 | I3sg2 | C13-02 | 4 | 0 | 0 | 0 | 0 | 0 | NA | 0 | 0 | 0 | 0 | 0.00021 | *** |
| CTP | C00063 | NT | C12 PARENT | 9 | 784237.3959 | 310313.0351 | 608315.6598 | 21.28978 | 0.3438 | 0.0162 | 20.894 | 21.4602 | 21.5151 | 21.2898 | 2.72E-06 | *** |
| CTP | C00063 | I3sg2 | C12 PARENT | 9 | 1491883.861 | 2143394.161 | 2369163.03 | 79.88587 | 2.6148 | 0.0327 | 82.5801 | 79.719 | 77.3585 | 79.8859 | 2.72E-06 | *** |
| CTP | C00063 | NT | C13-01 | 9 | 0 | 0 | 0 | 0 | 0 | NA | 0 | 0 | 0 | 0 | NA | |
| CTP | C00063 | I3sg2 | C13-01 | 9 | 0 | 0 | 0 | 0 | 0 | NA | 0 | 0 | 0 | 0 | NA | |
| CTP | C00063 | NT | C13-04 | 9 | 97599.41123 | 6241.613617 | 13041.48796 | 1.164398 | 1.2436 | 1.068 | 2.60029 | 0.43165 | 0.46126 | 1.1644 | 0.18018 | |
| CTP | C00063 | I3sg2 | C13-04 | 9 | 0 | 0 | 0 | 0 | 0 | NA | 0 | 0 | 0 | 0 | 0.18018 | |
| CTP | C00063 | NT | C13-05 | 9 | 1703389.558 | 721398.1059 | 1418014.856 | 48.47493 | 2.6813 | 0.0553 | 45.3825 | 49.8894 | 50.1529 | 48.4749 | 4.96E-05 | *** |
| CTP | C00063 | I3sg2 | C13-05 | 9 | 257962.1565 | 396663.203 | 524727.685 | 15.38851 | 1.5297 | 0.0994 | 14.279 | 14.7531 | 17.1335 | 15.3885 | 4.96E-05 | *** |
| CTP | C00063 | NT | C13-06 | 9 | 373095.1535 | 133180.3865 | 258280.2043 | 9.428474 | 0.4447 | 0.0472 | 9.94018 | 9.21029 | 9.13495 | 9.42847 | 6.78E-05 | *** |
| CTP | C00063 | I3sg2 | C13-06 | 9 | 0 | 13829.32116 | 46054.38523 | 0.67271 | 0.7643 | 1.1361 | 0 | 0.51435 | 1.50378 | 0.67271 | 6.78E-05 | *** |
| CTP | C00063 | NT | C13-07 | 9 | 530223.2069 | 174742.4538 | 343611.0096 | 12.788 | 1.1596 | 0.0907 | 14.1265 | 12.0846 | 12.153 | 12.788 | 0.00046 | *** |
| CTP | C00063 | I3sg2 | C13-07 | 9 | 48479.84541 | 124081.9988 | 106294.0839 | 3.589736 | 0.9712 | 0.2706 | 2.6835 | 4.61497 | 3.47074 | 3.58974 | 0.00046 | *** |
| CTP | C00063 | NT | C13-08 | 9 | 264859.8164 | 100120.2063 | 186120.3888 | 6.854421 | 0.2444 | 0.0357 | 7.05652 | 6.92396 | 6.58278 | 6.85442 | 1.65E-05 | *** |

| | | | | | | | | | | | | | | | | |
|---|---|---|---|---|---|---|---|---|---|---|---|---|---|---|---|---|
| CTP | C00063 | I3sg2 | C13-08 | 9 | 8264.586624 | 10716.62338 | 16337.84278 | 0.463173 | 0.0676 | 0.146 | 0.45747 | 0.39858 | 0.53347 | 0.46317 | 1.65E-06 | *** |
| Cystathionine | C02291 | NT | C12 PARENT | 7 | 4756593.125 | 3967319.001 | 3415265.563 | 89.62438 | 0.4559 | 0.0051 | 89.5498 | 90.113 | 89.2104 | 89.6244 | 4.81E-05 | *** |
| Cystathionine | C02291 | I3sg2 | C12 PARENT | 7 | 455483.2297 | 479574.2859 | 443495.2845 | 99.53987 | 0.797 | 0.008 | 98.6196 | 100 | 100 | 99.5399 | 4.81E-05 | *** |
| Cystathionine | C02291 | NT | C13-01 | 7 | 146989.5337 | 135401.8643 | 137680.5827 | 3.146383 | 0.4191 | 0.1332 | 2.76729 | 3.07549 | 3.59636 | 3.14638 | 0.00666 | ** |
| Cystathionine | C02291 | I3sg2 | C13-01 | 7 | 6375.476258 | 0 | 0 | 0.460132 | 0.797 | 1.7321 | 1.3804 | 0 | 0 | 0.46013 | 0.00666 | ** |
| Cystathionine | C02291 | NT | C13-02 | 7 | 99762.45041 | 78290.00704 | 55511.03065 | 1.702149 | 0.224 | 0.1316 | 1.87817 | 1.77827 | 1.45001 | 1.70215 | 0.00019 | *** |
| Cystathionine | C02291 | I3sg2 | C13-02 | 7 | 0 | 0 | 0 | 0 | 0 | NA | 0 | 0 | 0 | 0 | 0.00019 | *** |
| Cystathionine | C02291 | NT | C13-03 | 7 | 269762.2586 | 199488.3056 | 191980.8292 | 4.874851 | 0.2994 | 0.0614 | 5.07867 | 4.53114 | 5.01474 | 4.87485 | 9.40E-06 | *** |
| Cystathionine | C02291 | I3sg2 | C13-03 | 7 | 0 | 0 | 0 | 0 | 0 | NA | 0 | 0 | 0 | 0 | 9.40E-06 | *** |
| Cystathionine | C02291 | NT | C13-04 | 7 | 38568.74879 | 22105.81386 | 27889.4185 | 0.652241 | 0.13 | 0.1994 | 0.72611 | 0.50211 | 0.7285 | 0.65224 | 0.00097 | *** |
| Cystathionine | C02291 | I3sg2 | C13-04 | 7 | 0 | 0 | 0 | 0 | 0 | NA | 0 | 0 | 0 | 0 | 0.00097 | *** |
| Cytosine | C00380 | NT | C12 PARENT | 4 | 259084.8789 | 299879.8578 | 345329.3249 | 70.00901 | 1.2287 | 0.0176 | 68.9109 | 69.7801 | 71.3361 | 70.009 | 1.87E-06 | *** |
| Cytosine | C00380 | I3sg2 | C12 PARENT | 4 | 30990.03849 | 10554.20688 | 9793.809972 | 100 | 0 | 0 | 100 | 100 | 100 | 100 | 1.87E-06 | *** |
| Cytosine | C00380 | NT | C13-02 | 4 | 73870.70082 | 81722.80387 | 81563.57589 | 18.50442 | 1.4681 | 0.0793 | 19.648 | 19.0164 | 16.8489 | 18.5044 | 2.60E-05 | *** |
| Cytosine | C00380 | I3sg2 | C13-02 | 4 | 0 | 0 | 0 | 0 | 0 | NA | 0 | 0 | 0 | 0 | 2.60E-05 | *** |
| Cytosine | C00380 | NT | C13-03 | 4 | 43015.21862 | 48147.32923 | 57195.09877 | 11.48656 | 0.3083 | 0.0268 | 11.4411 | 11.2036 | 11.815 | 11.4866 | 3.45E-07 | *** |
| Cytosine | C00380 | I3sg2 | C13-03 | 4 | 0 | 0 | 0 | 0 | 0 | NA | 0 | 0 | 0 | 0 | 3.45E-07 | *** |
| F16BP | C00354 | NT | C12 PARENT | 6 | 32502.65739 | 58308.27656 | 83321.07041 | 2.034762 | 0.5201 | 0.2556 | 1.57157 | 1.93527 | 2.59744 | 2.03476 | 0.89559 | |
| F16BP | C00354 | I3sg2 | C12 PARENT | 6 | 19928.70591 | 17726.72734 | 40901.60093 | 2.17588 | 1.6695 | 0.7673 | 0.99299 | 1.44911 | 4.08553 | 2.17588 | 0.89559 | |
| F16BP | C00354 | NT | C13-05 | 6 | 56796.367 | 88064.74024 | 40982.09591 | 2.315565 | 0.9033 | 0.3901 | 2.74622 | 2.9229 | 1.27757 | 2.31556 | 0.01187 | * |
| F16BP | C00354 | I3sg2 | C13-05 | 6 | 1049.230526 | 466.8480175 | 0 | 0.030148 | 0.027 | 0.8971 | 0.05228 | 0.03816 | 0 | 0.03015 | 0.01187 | * |
| F16BP | C00354 | NT | C13-06 | 6 | 1978867.208 | 2866546.808 | 3083507.437 | 95.64967 | 0.4924 | 0.0051 | 95.6822 | 95.1418 | 96.125 | 95.6497 | 0.09626 | |
| F16BP | C00354 | I3sg2 | C13-06 | 6 | 1985951.825 | 1205090.107 | 960230.5079 | 97.79397 | 1.6426 | 0.0168 | 98.9547 | 98.5127 | 95.9145 | 97.794 | 0.09626 | |
| G6P-F6P | C00085 | NT | C12 PARENT | 6 | 345243.0134 | 375248.3513 | 296311.9544 | 29.86638 | 5.8554 | 0.1961 | 29.8642 | 35.7228 | 24.0121 | 29.8664 | 0.02294 | * |
| G6P-F6P | C00085 | I3sg2 | C12 PARENT | 6 | 175040.5175 | 68975.68155 | 160580.1946 | 13.39372 | 5.3698 | 0.4009 | 18.1822 | 7.58808 | 14.4109 | 13.3937 | 0.02294 | * |
| G6P-F6P | C00085 | NT | C13-05 | 6 | 21723.5602 | 0 | 31392.28844 | 1.474352 | 1.3194 | 0.8949 | 1.87913 | 0 | 2.54393 | 1.47435 | 0.13297 | |
| G6P-F6P | C00085 | I3sg2 | C13-05 | 6 | 1121.911818 | 0 | 0 | 0.038846 | 0.0673 | 1.7321 | 0.11654 | 0 | 0 | 0.03885 | 0.13297 | |
| G6P-F6P | C00085 | NT | C13-06 | 6 | 789077.0977 | 675195.2287 | 906305.6442 | 68.65926 | 4.5966 | 0.0669 | 68.2567 | 64.2772 | 73.444 | 68.6593 | 0.01203 | * |
| G6P-F6P | C00085 | I3sg2 | C13-06 | 6 | 786539.1416 | 840025.2686 | 953719.9473 | 86.56744 | 5.4219 | 0.0626 | 81.7012 | 92.4119 | 85.5891 | 86.5674 | 0.01203 | * |
| GDP | C00035 | NT | C12 PARENT | 10 | 196949.8239 | 77595.60437 | 236246.0346 | 8.49439 | 0.2015 | 0.0237 | 8.2635 | 8.63472 | 8.58495 | 8.49439 | 0.00061 | *** |
| GDP | C00035 | I3sg2 | C12 PARENT | 10 | 223753.0024 | 460109.3592 | 441899.4849 | 16.57442 | 1.4172 | 0.0855 | 15.322 | 18.1129 | 16.2883 | 16.5744 | 0.00061 | *** |
| GDP | C00035 | NT | C13-04 | 10 | 53041.69674 | 27756.59614 | 23131.09541 | 2.051588 | 1.1341 | 0.5528 | 2.22549 | 3.08871 | 0.84056 | 2.05159 | 0.03795 | * |
| GDP | C00035 | I3sg2 | C13-04 | 10 | 0 | 3639.50146 | 0 | 0.047758 | 0.0827 | 1.7321 | 0 | 0.14327 | 0 | 0.04776 | 0.03795 | * |
| GDP | C00035 | NT | C13-05 | 10 | 1537423.441 | 614289.9815 | 1916444.424 | 67.50175 | 2.6725 | 0.0396 | 64.5063 | 68.3573 | 69.6417 | 67.5017 | 0.00387 | ** |
| GDP | C00035 | I3sg2 | C13-05 | 10 | 1211688.185 | 1973897.476 | 2217639.402 | 80.80686 | 2.7556 | 0.0341 | 82.9734 | 77.7054 | 81.7418 | 80.8069 | 0.00387 | ** |
| GDP | C00035 | NT | C13-06 | 10 | 414090.7536 | 132192.3424 | 406196.5155 | 15.61504 | 1.5237 | 0.0976 | 17.3742 | 14.7102 | 14.7608 | 15.615 | 0.00034 | *** |
| GDP | C00035 | I3sg2 | C13-06 | 10 | 24892.63078 | 102584.7758 | 53443.05475 | 2.570963 | 1.2777 | 0.497 | 1.70458 | 4.0384 | 1.9699 | 2.57096 | 0.00034 | *** |
| GDP | C00035 | NT | C13-07 | 10 | 180041.3572 | 45740.9924 | 153252.065 | 6.071028 | 1.3065 | 0.2152 | 7.55407 | 5.08999 | 5.56903 | 6.07103 | 0.00129 | ** |
| GDP | C00035 | I3sg2 | C13-07 | 10 | 0 | 0 | 0 | 0 | 0 | NA | 0 | 0 | 0 | 0 | 0.00129 | ** |
| GDP | C00035 | NT | C13-08 | 10 | 1822.653238 | 1070.83683 | 16593.52879 | 0.266209 | 0.2924 | 1.0985 | 0.07647 | 0.11916 | 0.60299 | 0.26621 | 0.19 | |
| GDP | C00035 | I3sg2 | C13-08 | 10 | 0 | 0 | 0 | 0 | 0 | NA | 0 | 0 | 0 | 0 | 0.19 | |
| Glc | C00031 | NT | C12 PARENT | 6 | 0 | 105757.4095 | 105917.2104 | 0.688196 | 0.602 | 0.8747 | 0 | 0.94775 | 1.11684 | 0.6882 | 0.57672 | |
| Glc | C00031 | I3sg2 | C12 PARENT | 6 | 329490.87 | 269551.3128 | 0 | 1.081933 | 0.949 | 0.8772 | 1.7736 | 1.4722 | 0 | 1.08193 | 0.57672 | |
| Glc | C00031 | NT | C13-05 | 6 | 134172.9721 | 123311.0218 | 208918.5475 | 1.762712 | 0.5803 | 0.3292 | 1.98015 | 1.10506 | 2.20293 | 1.76271 | 0.5303 | |
| Glc | C00031 | I3sg2 | C13-05 | 6 | 256832.6984 | 277902.1868 | 379171.2788 | 1.52553 | 0.1471 | 0.0964 | 1.38249 | 1.51781 | 1.67629 | 1.52553 | 0.5303 | |
| Glc | C00031 | NT | C13-06 | 6 | 6641734.685 | 10929742.1 | 9168821.883 | 97.54909 | 0.7533 | 0.0077 | 98.0199 | 97.9472 | 96.6802 | 97.5491 | 0.81849 | |
| Glc | C00031 | I3sg2 | C13-06 | 6 | 17991180.45 | 17761984.31 | 22240511.39 | 97.39254 | 0.8107 | 0.0083 | 96.8439 | 97.01 | 98.3237 | 97.3925 | 0.81849 | |
| GlcNAc-6P | C00357 | NT | C12 PARENT | 8 | 146887.9845 | 135669.4611 | 128856.9008 | 26.3888 | 2.2596 | 0.0856 | 25.0495 | 28.9977 | 25.1193 | 26.3888 | 0.8016 | |
| GlcNAc-6P | C00357 | I3sg2 | C12 PARENT | 8 | 64682.26962 | 130383.2455 | 57805.61833 | 27.8078 | 8.8709 | 0.319 | 31.0359 | 34.6127 | 17.7749 | 27.8078 | 0.8016 | |
| GlcNAc-6P | C00357 | NT | C13-06 | 8 | 196234.3376 | 136875.2726 | 202189.5442 | 34.04493 | 5.1044 | 0.1499 | 33.4647 | 29.2554 | 39.4147 | 34.0449 | 0.45703 | |
| GlcNAc-6P | C00357 | I3sg2 | C13-06 | 8 | 66931.93631 | 129321.8658 | 179986.6651 | 40.59701 | 12.8199 | 0.3158 | 32.1153 | 34.3309 | 55.3448 | 40.597 | 0.45703 | |
| GlcNAc-6P | C00357 | NT | C13-08 | 8 | 243269.3107 | 195318.6719 | 181933.8075 | 39.56627 | 3.5533 | 0.0898 | 41.4858 | 41.7469 | 35.466 | 39.5663 | 0.08775 | |
| GlcNAc-6P | C00357 | I3sg2 | C13-08 | 8 | 76797.25573 | 116987.0551 | 87417.27 | 31.59519 | 5.0061 | 0.1584 | 36.8489 | 31.0564 | 26.8803 | 31.5952 | 0.08775 | |
| Gln | C00064 | NT | C12 PARENT | 5 | 764970010.8 | 657922508.5 | 675408713.7 | 98.87122 | 0.0393 | 4.00E-04 | 98.882 | 98.8277 | 98.904 | 98.8712 | 0.0002 | *** |
| Gln | C00064 | I3sg2 | C12 PARENT | 5 | 604357980.9 | 620999834.9 | 653120321.1 | 99.55686 | 0.0828 | 8.00E-04 | 99.4906 | 99.6497 | 99.5303 | 99.5569 | 0.0002 | *** |
| Gln | C00064 | NT | C13-01 | 5 | 342615.3006 | 1241900.351 | 538678.7454 | 0.103239 | 0.0742 | 0.7186 | 0.04429 | 0.18655 | 0.07888 | 0.10324 | 0.51392 | |
| Gln | C00064 | I3sg2 | C13-01 | 5 | 717410.8897 | 0 | 483357.1542 | 0.06392 | 0.0597 | 0.9332 | 0.1181 | 0 | 0.07366 | 0.06392 | 0.51392 | |
| Gln | C00064 | NT | C13-02 | 5 | 1507498.355 | 1478846.26 | 1846762.829 | 0.229145 | 0.0383 | 0.167 | 0.19486 | 0.22214 | 0.27043 | 0.22915 | 0.00049 | *** |
| Gln | C00064 | I3sg2 | C13-02 | 5 | 0 | 0 | 0 | 0 | 0 | NA | 0 | 0 | 0 | 0 | 0.00049 | *** |
| Gln | C00064 | NT | C13-03 | 5 | 1790163.751 | 1476159.045 | 1569205.443 | 0.227642 | 0.0052 | 0.0227 | 0.2314 | 0.22174 | 0.22979 | 0.22764 | 0.00511 | ** |
| Gln | C00064 | I3sg2 | C13-03 | 5 | 877160.2682 | 628832.4014 | 1019889.882 | 0.133576 | 0.0288 | 0.2158 | 0.1444 | 0.10091 | 0.15542 | 0.13358 | 0.00511 | ** |
| Gln | C00064 | NT | C13-04 | 5 | 2990589.3 | 2144450.377 | 2127125.616 | 0.34006 | 0.0406 | 0.1195 | 0.38657 | 0.32212 | 0.31149 | 0.34006 | 0.00261 | ** |
| Gln | C00064 | I3sg2 | C13-04 | 5 | 881634.7157 | 1100133.557 | 1177645.293 | 0.167045 | 0.019 | 0.1139 | 0.14514 | 0.17653 | 0.17946 | 0.16704 | 0.00261 | ** |
| Gln | C00064 | NT | C13-05 | 5 | 2018264.809 | 1462943.701 | 1402957.698 | 0.228694 | 0.0288 | 0.1259 | 0.26089 | 0.21975 | 0.20544 | 0.22869 | 0.00186 | ** |
| Gln | C00064 | I3sg2 | C13-05 | 5 | 618163.6708 | 453947.4204 | 401573.0671 | 0.078601 | 0.0209 | 0.2657 | 0.10176 | 0.07284 | 0.0612 | 0.0786 | 0.00186 | ** |
| Glu | C00025 | NT | C12 PARENT | 5 | 286803812.2 | 242392072 | 229857793 | 58.17576 | 0.2071 | 0.0036 | 58.0374 | 58.0759 | 58.4139 | 58.1758 | 5.89E-06 | *** |
| Glu | C00025 | I3sg2 | C12 PARENT | 5 | 229311806 | 264495610.6 | 252734443.4 | 66.08472 | 0.379 | 0.0057 | 65.6946 | 66.1081 | 66.4515 | 66.0847 | 5.89E-06 | *** |
| Glu | C00025 | NT | C13-01 | 5 | 6957775.429 | 6118268.567 | 5933620.525 | 1.460597 | 0.0502 | 0.0344 | 1.40797 | 1.46591 | 1.50791 | 1.4606 | 0.00083 | *** |
| Glu | C00025 | I3sg2 | C13-01 | 5 | 6096681.803 | 7205301.087 | 6659834.768 | 1.766193 | 0.0301 | 0.0171 | 1.74661 | 1.80089 | 1.75107 | 1.76619 | 0.00083 | *** |
| Glu | C00025 | NT | C13-02 | 5 | 101874234.2 | 86177122.57 | 81372574.54 | 20.64736 | 0.032 | 0.0016 | 20.6152 | 20.6476 | 20.6793 | 20.6474 | 1.68E-06 | *** |
| Glu | C00025 | I3sg2 | C13-02 | 5 | 63746800.42 | 72347501.47 | 69262671 | 18.18545 | 0.0927 | 0.0051 | 18.2626 | 18.0825 | 18.2112 | 18.1854 | 1.68E-06 | *** |
| Glu | C00025 | NT | C13-03 | 5 | 30523234.82 | 25092087.02 | 23152860.47 | 6.024151 | 0.1468 | 0.0244 | 6.17666 | 6.01194 | 5.88385 | 6.02415 | 0.00038 | *** |
| Glu | C00025 | I3sg2 | C13-03 | 5 | 17051382.46 | 19826497.77 | 18041927.21 | 4.861395 | 0.1078 | 0.0222 | 4.88498 | 4.95544 | 4.74376 | 4.86139 | 0.00038 | *** |
| Glu | C00025 | NT | C13-04 | 5 | 41381781.59 | 34711876.56 | 32485501.2 | 8.315448 | 0.0592 | 0.0071 | 8.37399 | 8.31679 | 8.25556 | 8.31545 | 4.32E-06 | *** |
| Glu | C00025 | I3sg2 | C13-04 | 5 | 20887899.14 | 23080540.51 | 22258699.19 | 5.868441 | 0.1086 | 0.0185 | 5.98409 | 5.76875 | 5.85248 | 5.86844 | 4.32E-06 | *** |
| Glu | C00025 | NT | C13-05 | 5 | 26629438.08 | 22879436.54 | 20696246.93 | 5.376688 | 0.1116 | 0.0208 | 5.38872 | 5.4818 | 5.25955 | 5.37669 | 0.00012 | *** |
| Glu | C00025 | I3sg2 | C13-05 | 5 | 11962745.35 | 13140272.48 | 11371762.1 | 3.233806 | 0.2229 | 0.0689 | 3.42716 | 3.28428 | 2.98998 | 3.23381 | 0.00012 | *** |
| GlucA | C00191 | NT | C12 PARENT | 6 | 0 | 0 | 5650.643218 | 0.909034 | 1.5745 | 1.7321 | 0 | 0 | 2.7271 | 0.90903 | 0.4265 | |
| GlucA | C00191 | I3sg2 | C12 PARENT | 6 | 0 | 0 | 11554.28699 | 8.236723 | 14.2664 | 1.7321 | 0 | 0 | 24.7102 | 8.23672 | 0.4265 | |
| GlucA | C00191 | NT | C13-06 | 6 | 224697.1013 | 168885.8281 | 201552.5704 | 99.09097 | 1.5745 | 0.0159 | 100 | 100 | 97.2729 | 99.091 | 0.4265 | |
| GlucA | C00191 | I3sg2 | C13-06 | 6 | 65909.92254 | 58496.68386 | 35204.94952 | 91.76328 | 14.2664 | 0.1555 | 100 | 100 | 75.2898 | 91.7633 | 0.4265 | |
| Gly | C00037 | NT | C12 PARENT | 2 | 12275352.32 | 12973289.83 | 12073587.28 | 87.00422 | 0.8322 | 0.0096 | 87.4534 | 87.5154 | 86.0439 | 87.0042 | 0.00019 | *** |
| Gly | C00037 | I3sg2 | C12 PARENT | 2 | 8207910.35 | 9301678.031 | 9566382.233 | 94.78595 | 0.5956 | 0.0063 | 94.5756 | 95.4581 | 94.3241 | 94.7859 | 0.00019 | *** |
| Gly | C00037 | NT | C13-01 | 2 | 20143.80947 | 0 | 39961.95982 | 0.142768 | 0.1424 | 0.9974 | 0.14351 | 0 | 0.28479 | 0.14277 | 0.16197 | |
| Gly | C00037 | I3sg2 | C13-01 | 2 | 505.0616665 | 0 | 0 | 0.00194 | 0.0034 | 1.7321 | 0.00582 | 0 | 0 | 0.00194 | 0.16197 | |
| Gly | C00037 | NT | C13-02 | 2 | 1740960.032 | 1850721.061 | 1918340.513 | 12.85301 | 0.7098 | 0.0552 | 12.4031 | 12.4846 | 13.6713 | 12.853 | 0.00014 | *** |
| Gly | C00037 | I3sg2 | C13-02 | 2 | 470261.9671 | 442570.6449 | 575650.1074 | 5.212114 | 0.5945 | 0.1141 | 5.41859 | 4.54187 | 5.67589 | 5.21211 | 0.00014 | *** |
| Glycerol-3P | C00093 | NT | C12 PARENT | 3 | 540333.0881 | 432272.8105 | 264942.7395 | 36.64855 | 4.3213 | 0.1179 | 32.0526 | 37.2637 | 40.6293 | 36.6485 | 0.09537 | |
| Glycerol-3P | C00093 | I3sg2 | C12 PARENT | 3 | 563495.1051 | 788182.2468 | 651672.4964 | 45.53811 | 5.611 | 0.1232 | 51.7486 | 44.0315 | 40.8342 | 45.5381 | 0.09537 | |
| Glycerol-3P | C00093 | NT | C13-03 | 3 | 1145435.134 | 727764.6459 | 387154.7937 | 63.35145 | 4.3213 | 0.0682 | 67.9474 | 62.7363 | 59.3707 | 63.3515 | 0.09537 | |
| Glycerol-3P | C00093 | I3sg2 | C13-03 | 3 | 525413.5083 | 1001857.698 | 944226.8119 | 54.46189 | 5.611 | 0.103 | 48.2514 | 55.9685 | 59.1658 | 54.4619 | 0.09537 | |
| GMP | C00144 | NT | C12 PARENT | 10 | 57849.13576 | 6697.099321 | 94493.00202 | 6.726003 | 3.2123 | 0.4776 | 8.22026 | 3.0388 | 8.91894 | 6.726 | 0.01292 | * |
| GMP | C00144 | I3sg2 | C12 PARENT | 10 | 83100.29757 | 196404.9071 | 200044.7137 | 17.57842 | 3.0049 | 0.1709 | 14.4665 | 20.4633 | 17.8055 | 17.5784 | 0.01292 | * |

| | | | | | | | | | | | | | | | | |
|---|---|---|---|---|---|---|---|---|---|---|---|---|---|---|---|---|
| GMP | C00144 | NT | C13-04 | 10 | 0 | 0 | 0 | 0 | 0 | NA | 0 | 0 | 0 | 0 | NA | |
| GMP | C00144 | I3sg2 | C13-04 | 10 | 0 | 0 | 0 | 0 | 0 | NA | 0 | 0 | 0 | 0 | NA | |
| GMP | C00144 | NT | C13-05 | 10 | 495326.7091 | 198701.781 | 693487.0928 | 75.33405 | 13.0746 | 0.1736 | 70.3851 | 90.1607 | 65.4564 | 75.3341 | 0.41193 | |
| GMP | C00144 | I3sg2 | C13-05 | 10 | 491333.0855 | 763385.4547 | 923457.6823 | 82.42158 | 3.0049 | 0.0365 | 85.5335 | 79.5367 | 82.1945 | 82.4216 | 0.41193 | |
| GMP | C00144 | NT | C13-06 | 10 | 148149.9577 | 11875.27878 | 199722.75 | 15.09718 | 8.4797 | 0.5617 | 21.0519 | 5.38839 | 18.8513 | 15.0972 | 0.0368 | * |
| GMP | C00144 | I3sg2 | C13-06 | 10 | 0 | 0 | 0 | 0 | 0 | NA | 0 | 0 | 0 | 0 | 0.0368 | * |
| GMP | C00144 | NT | C13-07 | 10 | 2412.287312 | 3112.187755 | 71761.08996 | 2.842757 | 3.4457 | 1.2121 | 0.34278 | 1.41215 | 6.77334 | 2.84276 | 0.22622 | |
| GMP | C00144 | I3sg2 | C13-07 | 10 | 0 | 0 | 0 | 0 | 0 | NA | 0 | 0 | 0 | 0 | 0.22622 | |
| GSH | C00051 | NT | C12 PARENT | 10 | 66578098.05 | 64228689.09 | 62853388.44 | 64.87348 | 0.5822 | 0.009 | 64.2013 | 65.2151 | 65.2041 | 64.8735 | 0.00046 | *** |
| GSH | C00051 | I3sg2 | C12 PARENT | 10 | 44874555.48 | 43061678.59 | 44808780.66 | 68.5151 | 0.1367 | 0.002 | 68.3669 | 68.6363 | 68.5421 | 68.5151 | 0.00046 | *** |
| GSH | C00051 | NT | C13-01 | 10 | 0 | 0 | 0 | 0 | 0 | NA | 0 | 0 | 0 | 0 | 0.13597 | |
| GSH | C00051 | I3sg2 | C13-01 | 10 | 20391.09049 | 30853.4594 | 0 | 0.026748 | 0.0249 | 0.9299 | 0.03107 | 0.04918 | 0 | 0.02675 | 0.13597 | |
| GSH | C00051 | NT | C13-02 | 10 | 22234952.06 | 20870393.21 | 20362995.7 | 21.25221 | 0.167 | 0.0079 | 21.4412 | 21.1909 | 21.1246 | 21.2522 | 0.0009 | *** |
| GSH | C00051 | I3sg2 | C13-02 | 10 | 12911514.05 | 11970646.4 | 12786863.56 | 19.43682 | 0.3139 | 0.0162 | 19.6708 | 19.0801 | 19.5595 | 19.4368 | 0.0009 | *** |
| GSH | C00051 | NT | C13-03 | 10 | 3711113.182 | 3338587.539 | 3433781.849 | 3.51023 | 0.1046 | 0.0298 | 3.57863 | 3.38986 | 3.5622 | 3.51023 | 0.01726 | * |
| GSH | C00051 | I3sg2 | C13-03 | 10 | 2487569.463 | 2593936.707 | 2599036.897 | 3.966657 | 0.1725 | 0.0435 | 3.78984 | 4.13449 | 3.97564 | 3.96666 | 0.01726 | * |
| GSH | C00051 | NT | C13-04 | 10 | 7194302.459 | 6332206.78 | 6177755.029 | 6.591907 | 0.2994 | 0.0454 | 6.93747 | 6.42946 | 6.4088 | 6.59191 | 0.00271 | ** |
| GSH | C00051 | I3sg2 | C13-04 | 10 | 3545321.658 | 3341241.965 | 3589750.473 | 5.406019 | 0.0828 | 0.0153 | 5.40134 | 5.32563 | 5.49109 | 5.40602 | 0.00271 | ** |
| GSH | C00051 | NT | C13-05 | 10 | 3511877.562 | 3261333.394 | 3127134.606 | 3.314004 | 0.0712 | 0.0215 | 3.3865 | 3.31142 | 3.24409 | 3.314 | 0.00431 | ** |
| GSH | C00051 | I3sg2 | C13-05 | 10 | 1731477.578 | 1709073.531 | 1517666.178 | 2.56118 | 0.212 | 0.0828 | 2.63793 | 2.7241 | 2.32151 | 2.56118 | 0.00431 | ** |
| GSH | C00051 | NT | C13-06 | 10 | 263641.282 | 302587.4578 | 258821.0793 | 0.276655 | 0.0274 | 0.0991 | 0.25423 | 0.30723 | 0.2685 | 0.27665 | 0.00247 | ** |
| GSH | C00051 | I3sg2 | C13-06 | 10 | 48619.34824 | 12087.90792 | 71984.85143 | 0.067817 | 0.0457 | 0.6745 | 0.07407 | 0.01927 | 0.11011 | 0.06782 | 0.00247 | ** |
| GSH | C00051 | NT | C13-07 | 10 | 208179.8237 | 153623.4868 | 181029.1353 | 0.18151 | 0.023 | 0.1269 | 0.20075 | 0.15598 | 0.1878 | 0.18151 | 0.00061 | *** |
| GSH | C00051 | I3sg2 | C13-07 | 10 | 18380.72078 | 19428.27897 | 0 | 0.019657 | 0.0171 | 0.8693 | 0.028 | 0.03097 | 0 | 0.01966 | 0.00061 | *** |
| GSSG | C00127 | NT | C12 PARENT | 20 | 4182559.518 | 5165982.957 | 3134540.483 | 45.39624 | 1.2856 | 0.0283 | 44.3316 | 46.8245 | 45.0326 | 45.3962 | 0.00945 | ** |
| GSSG | C00127 | I3sg2 | C12 PARENT | 20 | 3494537.609 | 3718002.871 | 4297458.096 | 49.10202 | 0.4775 | 0.0097 | 48.5533 | 49.4233 | 49.3295 | 49.102 | 0.00945 | ** |
| GSSG | C00127 | NT | C13-01 | 20 | 111902.7343 | 120094.8402 | 31669.58799 | 0.909866 | 0.3969 | 0.4363 | 1.18607 | 1.08854 | 0.45498 | 0.90987 | 0.0141 | * |
| GSSG | C00127 | I3sg2 | C13-01 | 20 | 125807.1816 | 160768.187 | 182606.8991 | 1.993719 | 0.2138 | 0.1072 | 1.74797 | 2.13709 | 2.0961 | 1.99372 | 0.0141 | * |
| GSSG | C00127 | NT | C13-02 | 20 | 2306800.329 | 2780750.532 | 1878889.91 | 25.54937 | 1.3061 | 0.0511 | 24.4501 | 25.2048 | 26.9932 | 25.5494 | 0.03872 | * |
| GSSG | C00127 | I3sg2 | C13-02 | 20 | 1681765.013 | 1745567.649 | 2021304.931 | 23.25746 | 0.0945 | 0.0041 | 23.3665 | 23.2038 | 23.2021 | 23.2575 | 0.03872 | * |
| GSSG | C00127 | NT | C13-03 | 20 | 646507.269 | 445286.6565 | 407639.2099 | 5.581631 | 1.4281 | 0.2559 | 6.85243 | 4.03608 | 5.85638 | 5.58163 | 0.1701 | |
| GSSG | C00127 | I3sg2 | C13-03 | 20 | 502572.6906 | 503636.8259 | 634938.1145 | 6.988633 | 0.2968 | 0.0425 | 6.98277 | 6.69483 | 7.2883 | 6.98863 | 0.1701 | |
| GSSG | C00127 | NT | C13-04 | 20 | 1066949.111 | 1309673.822 | 749983.5969 | 11.31812 | 0.5482 | 0.0484 | 11.3088 | 11.8709 | 10.7747 | 11.3181 | 0.02691 | * |
| GSSG | C00127 | I3sg2 | C13-04 | 20 | 751513.2637 | 731949.3848 | 845603.1784 | 9.959272 | 0.4178 | 0.042 | 10.4416 | 9.72978 | 9.70648 | 9.95927 | 0.02691 | * |
| GSSG | C00127 | NT | C13-05 | 20 | 585283.398 | 725408.0767 | 405334.409 | 6.200638 | 0.3759 | 0.0606 | 6.20351 | 6.57511 | 5.82327 | 6.20063 | 0.01318 | * |
| GSSG | C00127 | I3sg2 | C13-05 | 20 | 372704.1466 | 398217.4035 | 462842.3027 | 5.261572 | 0.0727 | 0.0138 | 5.17837 | 5.29349 | 5.31286 | 5.26157 | 0.01318 | * |
| GSSG | C00127 | NT | C13-06 | 20 | 228234.1249 | 192046.3081 | 196906.4899 | 2.329557 | 0.5496 | 0.2359 | 2.41909 | 1.74071 | 2.82887 | 2.32956 | 0.26306 | |
| GSSG | C00127 | I3sg2 | C13-06 | 20 | 145395.5317 | 152341.4804 | 137145.3921 | 1.873154 | 0.2589 | 0.1382 | 2.02013 | 2.02507 | 1.57426 | 1.87315 | 0.26306 | |
| GSSG | C00127 | NT | C13-07 | 20 | 209775.5425 | 177332.9955 | 118327.9407 | 1.843586 | 0.3322 | 0.1802 | 2.22344 | 1.60735 | 1.69997 | 1.84359 | 0.03491 | * |
| GSSG | C00127 | I3sg2 | C13-07 | 20 | 78041.01741 | 93606.89103 | 113078.9325 | 1.208875 | 0.1112 | 0.092 | 1.08431 | 1.24431 | 1.29801 | 1.20888 | 0.03491 | * |
| GSSG | C00127 | NT | C13-08 | 20 | 60586.2008 | 85051.42687 | 37311.87081 | 0.649704 | 0.1176 | 0.181 | 0.64216 | 0.77091 | 0.53604 | 0.6497 | 0.06899 | |
| GSSG | C00127 | I3sg2 | C13-08 | 20 | 38515.60912 | 0 | 0 | 0.178379 | 0.309 | 1.7321 | 0.53514 | 0 | 0 | 0.17838 | 0.06899 | |
| GSSG | C00127 | NT | C13-09 | 20 | 36114.35517 | 31014.19889 | 0 | 0.221298 | 0.1983 | 0.896 | 0.38278 | 0.28111 | 0 | 0.2213 | 0.73751 | |
| GSSG | C00127 | I3sg2 | C13-09 | 20 | 6476.184676 | 18683.62319 | 16762.04799 | 0.176916 | 0.0803 | 0.454 | 0.08998 | 0.24836 | 0.19241 | 0.17692 | 0.73751 | |
| GTP | C00044 | NT | C12 PARENT | 10 | 4161898.013 | 781236.399 | 2480318.23 | 8.350766 | 0.5071 | 0.0607 | 8.6677 | 7.76595 | 8.61865 | 8.35077 | 3.15E-05 | *** |
| GTP | C00044 | I3sg2 | C12 PARENT | 10 | 1698211.611 | 4166092.398 | 3847606.243 | 18.33284 | 0.6583 | 0.0359 | 19.0005 | 17.6842 | 18.3138 | 18.3328 | 3.15E-05 | *** |
| GTP | C00044 | NT | C13-01 | 10 | 0 | 0 | 0 | 0 | 0 | NA | 0 | 0 | 0 | 0 | NA | |
| GTP | C00044 | I3sg2 | C13-01 | 10 | 0 | 0 | 0 | 0 | 0 | NA | 0 | 0 | 0 | 0 | NA | |
| GTP | C00044 | NT | C13-02 | 10 | 178195.1836 | 17844.28373 | 173594.5549 | 0.383902 | 0.2132 | 0.5554 | 0.37111 | 0.17738 | 0.60321 | 0.3839 | 0.34271 | |
| GTP | C00044 | I3sg2 | C13-02 | 10 | 29067.28765 | 131529.6494 | 255037.0437 | 0.699154 | 0.4608 | 0.6591 | 0.32522 | 0.55832 | 1.21393 | 0.69915 | 0.34271 | |
| GTP | C00044 | NT | C13-03 | 10 | 434677.187 | 7666.176014 | 246232.6116 | 0.612364 | 0.465 | 0.7593 | 0.90527 | 0.07621 | 0.85561 | 0.61236 | 0.39842 | |
| GTP | C00044 | I3sg2 | C13-03 | 10 | 62925.40406 | 239129.0121 | 193751.6203 | 0.880438 | 0.1597 | 0.1813 | 0.70404 | 1.01505 | 0.92222 | 0.88044 | 0.39842 | |
| GTP | C00044 | NT | C13-04 | 10 | 476133.4473 | 80799.88451 | 219125.6706 | 0.852077 | 0.1226 | 0.1439 | 0.99161 | 0.8032 | 0.76142 | 0.85208 | 0.01248 | * |
| GTP | C00044 | I3sg2 | C13-04 | 10 | 278414.5039 | 466166.2858 | 741035.9358 | 2.87367 | 0.8019 | 0.2791 | 3.11505 | 1.97878 | 3.52718 | 2.87367 | 0.01248 | * |
| GTP | C00044 | NT | C13-05 | 10 | 31836699.38 | 6967581.562 | 19293883.25 | 67.53622 | 1.5394 | 0.0228 | 66.3041 | 69.2619 | 67.0427 | 67.5362 | 0.00894 | ** |
| GTP | C00044 | I3sg2 | C13-05 | 10 | 6677848.863 | 17858443.75 | 15154901.98 | 74.21831 | 1.8853 | 0.0254 | 74.7152 | 75.8054 | 72.1343 | 74.2183 | 0.00894 | ** |
| GTP | C00044 | NT | C13-06 | 10 | 8094078.64 | 1551817.784 | 4429895.833 | 15.89201 | 0.8359 | 0.0526 | 16.857 | 15.426 | 15.3931 | 15.892 | 5.21E-05 | *** |
| GTP | C00044 | I3sg2 | C13-06 | 10 | 153948.2269 | 445920.4828 | 730435.6679 | 2.364007 | 0.9674 | 0.4092 | 1.72245 | 1.89284 | 3.47673 | 2.36401 | 5.21E-05 | *** |
| GTP | C00044 | NT | C13-07 | 10 | 2226783.565 | 616918.6037 | 1625640.803 | 5.472969 | 0.7628 | 0.1394 | 4.63757 | 6.13254 | 5.6488 | 5.47297 | 0.00059 | *** |
| GTP | C00044 | I3sg2 | C13-07 | 10 | 37317.37196 | 251006.8083 | 86506.42959 | 0.631584 | 0.3758 | 0.595 | 0.41753 | 1.06547 | 0.41175 | 0.63158 | 0.00059 | *** |
| GTP | C00044 | NT | C13-08 | 10 | 607700.6976 | 35898.72399 | 309825.3923 | 0.899686 | 0.4795 | 0.533 | 1.26562 | 0.35685 | 1.07659 | 0.89969 | 0.03138 | * |
| GTP | C00044 | I3sg2 | C13-08 | 10 | 0 | 0 | 0 | 0 | 0 | NA | 0 | 0 | 0 | 0 | 0.03138 | * |
| IMP | C00130 | NT | C12 PARENT | 10 | 0 | 7884.944085 | 11699.61921 | 3.366155 | 2.9486 | 0.8759 | 0 | 5.49175 | 4.60671 | 3.36616 | 0.0081 | ** |
| IMP | C00130 | I3sg2 | C12 PARENT | 10 | 140709.8423 | 83087.43323 | 151480.4965 | 25.98876 | 7.4513 | 0.2867 | 30.2266 | 17.3851 | 30.3546 | 25.9888 | 0.0081 | ** |
| IMP | C00130 | NT | C13-04 | 10 | 0 | 0 | 0 | 0 | 0 | NA | 0 | 0 | 0 | 0 | NA | |
| IMP | C00130 | I3sg2 | C13-04 | 10 | 0 | 0 | 0 | 0 | 0 | NA | 0 | 0 | 0 | 0 | NA | |
| IMP | C00130 | NT | C13-05 | 10 | 148235.6662 | 135692.9902 | 242269.2975 | 96.63384 | 2.9486 | 0.0305 | 100 | 94.5082 | 95.3933 | 96.6338 | 0.0081 | ** |
| IMP | C00130 | I3sg2 | C13-05 | 10 | 324807.3551 | 394835.6041 | 347555.7095 | 74.01124 | 7.4513 | 0.1007 | 69.7734 | 82.6149 | 69.6454 | 74.0112 | 0.0081 | ** |
| Lac | C00186 | NT | C12 PARENT | 3 | 1609134.49 | 1663379.287 | 1719100.361 | 10.05818 | 0.5498 | 0.0547 | 10.5925 | 10.088 | 9.4941 | 10.0582 | 0.00711 | ** |
| Lac | C00186 | I3sg2 | C12 PARENT | 3 | 1690414.821 | 2017037.651 | 1562774.454 | 26.41955 | 5.5583 | 0.2104 | 27.1367 | 31.5844 | 20.5375 | 26.4196 | 0.00711 | ** |
| Lac | C00186 | NT | C13-01 | 3 | 0 | 78026.37228 | 68947.45039 | 0.284662 | 0.2508 | 0.8811 | 0 | 0.47321 | 0.38078 | 0.28466 | 0.12076 | |
| Lac | C00186 | I3sg2 | C13-01 | 3 | 0 | 0 | 0 | 0 | 0 | NA | 0 | 0 | 0 | 0 | 0.12076 | |
| Lac | C00186 | NT | C13-02 | 3 | 285461.6486 | 150292.9701 | 289020.6916 | 1.462261 | 0.4975 | 0.3402 | 1.87912 | 0.91149 | 1.59618 | 1.46226 | 0.00773 | ** |
| Lac | C00186 | I3sg2 | C13-02 | 3 | 5765.571481 | 0 | 0 | 0.030852 | 0.0534 | 1.7321 | 0.09256 | 0 | 0 | 0.03085 | 0.00773 | ** |
| Lac | C00186 | NT | C13-03 | 3 | 13296669.22 | 14597069.25 | 16029968.5 | 88.19489 | 0.5772 | 0.0065 | 87.5284 | 88.5273 | 88.5289 | 88.1949 | 0.01054 | * |
| Lac | C00186 | I3sg2 | C13-03 | 3 | 4533066.479 | 4369135.77 | 6046610.54 | 73.5496 | 5.5645 | 0.0757 | 72.7707 | 68.4156 | 79.4625 | 73.5496 | 0.01054 | * |
| Mal | C00149 | NT | C12 PARENT | 4 | 44067116.22 | 37319931.67 | 34013435.11 | 61.87487 | 0.9184 | 0.0148 | 60.8254 | 62.2675 | 62.5316 | 61.8749 | 0.00035 | *** |
| Mal | C00149 | I3sg2 | C12 PARENT | 4 | 32379189.73 | 35360720.62 | 30854341.33 | 68.11724 | 0.2707 | 0.004 | 68.3872 | 68.1186 | 67.8459 | 68.1172 | 0.00035 | *** |
| Mal | C00149 | NT | C13-01 | 4 | 2280766.967 | 1645339.086 | 1451097.692 | 2.853697 | 0.2579 | 0.0904 | 3.14812 | 2.74521 | 2.66776 | 2.8537 | 0.3861 | |
| Mal | C00149 | I3sg2 | C13-01 | 4 | 1212376.053 | 1405287.911 | 1278106.498 | 2.692734 | 0.1255 | 0.0466 | 2.56063 | 2.70714 | 2.81044 | 2.69273 | 0.3861 | |
| Mal | C00149 | NT | C13-02 | 4 | 12297014.46 | 10102967.54 | 8915271.947 | 16.74008 | 0.3086 | 0.0184 | 16.9735 | 16.8566 | 16.3902 | 16.7401 | 0.00125 | ** |
| Mal | C00149 | I3sg2 | C13-02 | 4 | 7163046.768 | 7929933.679 | 6808317.58 | 15.1253 | 0.1527 | 0.0101 | 15.1289 | 15.2762 | 14.9709 | 15.1253 | 0.00125 | ** |
| Mal | C00149 | NT | C13-03 | 4 | 9565313.552 | 7635496.38 | 7029130.03 | 12.95507 | 0.2333 | 0.018 | 13.2029 | 12.7397 | 12.9226 | 12.9551 | 0.00032 | *** |
| Mal | C00149 | I3sg2 | C13-03 | 4 | 4695815.763 | 5177668.88 | 4798157.244 | 10.14761 | 0.3502 | 0.0345 | 9.91791 | 9.97422 | 10.5507 | 10.1476 | 0.00032 | *** |
| Mal | C00149 | NT | C13-04 | 4 | 4238298.772 | 3231084.897 | 2985022.607 | 5.576289 | 0.242 | 0.0434 | 5.85008 | 5.391 | 5.48778 | 5.57629 | 0.00037 | *** |
| Mal | C00149 | I3sg2 | C13-04 | 4 | 1896410.678 | 2036886.487 | 1738185.223 | 3.917104 | 0.0918 | 0.0234 | 4.00536 | 3.92384 | 3.82211 | 3.9171 | 0.00037 | *** |
| Met | C00073 | NT | C12 PARENT | 5 | 11949241.36 | 20763994.22 | 22588755.62 | 97.80162 | 0.5893 | 0.006 | 98.3734 | 97.8352 | 97.1962 | 97.8016 | 0.01464 | * |
| Met | C00073 | I3sg2 | C12 PARENT | 5 | 11934867.9 | 23913757.28 | 19759933.86 | 99.39207 | 0.3165 | 0.0032 | 99.0277 | 99.5489 | 99.5996 | 99.3921 | 0.01464 | * |
| Met | C00073 | NT | C13-01 | 5 | 0 | 0 | 0 | 0 | 0 | NA | 0 | 0 | 0 | 0 | NA | |
| Met | C00073 | I3sg2 | C13-01 | 5 | 0 | 0 | 0 | 0 | 0 | NA | 0 | 0 | 0 | 0 | NA | |
| Met | C00073 | NT | C13-04 | 5 | 197576.7918 | 459439.0553 | 651610.8869 | 2.198378 | 0.5893 | 0.2681 | 1.62657 | 2.16477 | 2.80379 | 2.19838 | 0.01464 | * |
| Met | C00073 | I3sg2 | C13-04 | 5 | 117176.079 | 108370.4596 | 79436.36931 | 0.607925 | 0.3165 | 0.5207 | 0.97225 | 0.45113 | 0.4004 | 0.60793 | 0.01464 | * |
| MTA | C00170 | NT | C12 PARENT | 11 | 66458.8887 | 690770.1711 | 500522.2476 | 10.4351 | 0.2103 | 0.0202 | 10.3137 | 10.3136 | 10.6779 | 10.4351 | 3.21E-05 | *** |

| | | | | | | | | | | | | | | | | |
|---|---|---|---|---|---|---|---|---|---|---|---|---|---|---|---|---|
| MTA | C00170 | I3sg2 | C12 PARENT | 11 | 494754.9977 | 245386.0702 | 147472.67 | 28.11933 | 1.4642 | 0.0521 | 26.4611 | 28.6629 | 29.234 | 28.1193 | 3.21E-05 | *** |
| MTA | C00170 | NT | C13-01 | 11 | 0 | 0 | 0 | 0 | 0 | NA | 0 | 0 | 0 | 0 | NA | |
| MTA | C00170 | I3sg2 | C13-01 | 11 | 0 | 0 | 0 | 0 | 0 | NA | 0 | 0 | 0 | 0 | NA | |
| MTA | C00170 | NT | C13-02 | 11 | 1346.837799 | 38517.7856 | 41918.23727 | 0.559459 | 0.3429 | 0.6129 | 0.20902 | 0.57509 | 0.89427 | 0.55946 | 0.80689 | |
| MTA | C00170 | I3sg2 | C13-02 | 11 | 24384.85836 | 0 | 0 | 0.434727 | 0.753 | 1.7321 | 1.30418 | 0 | 0 | 0.43473 | 0.80689 | |
| MTA | C00170 | NT | C13-03 | 11 | 0 | 56044.72264 | 43625.49116 | 0.589157 | 0.5124 | 0.8697 | 0 | 0.83678 | 0.93069 | 0.58916 | 0.60296 | |
| MTA | C00170 | I3sg2 | C13-03 | 11 | 15569.66153 | 6808.224729 | 3273.058414 | 0.758932 | 0.0972 | 0.128 | 0.83272 | 0.79525 | 0.64883 | 0.75893 | 0.60296 | |
| MTA | C00170 | NT | C13-04 | 11 | 0 | 89206.18664 | 65825.84868 | 0.912068 | 0.7907 | 0.8669 | 0 | 1.3319 | 1.4043 | 0.91207 | 0.26091 | |
| MTA | C00170 | I3sg2 | C13-04 | 11 | 9330.9749 | 3026.356096 | 0 | 0.284184 | 0.2566 | 0.9031 | 0.49905 | 0.3535 | 0 | 0.28418 | 0.26091 | |
| MTA | C00170 | NT | C13-05 | 11 | 464212.5647 | 4615728.548 | 3234062.809 | 69.98364 | 1.7822 | 0.0255 | 72.0411 | 68.9157 | 68.9941 | 69.9836 | 0.87814 | |
| MTA | C00170 | I3sg2 | C13-05 | 11 | 1311621.981 | 600889.4735 | 353710.8598 | 70.15179 | 0.0356 | 5.00E-04 | 70.1498 | 70.1883 | 70.1172 | 70.1518 | 0.87814 | |
| MTA | C00170 | NT | C13-06 | 11 | 89868.22534 | 857388.7429 | 556183.6612 | 12.87113 | 1.0424 | 0.081 | 13.9466 | 12.8014 | 11.8654 | 12.8711 | 4.20E-05 | *** |
| MTA | C00170 | I3sg2 | C13-06 | 11 | 14081.0032 | 0 | 0 | 0.251033 | 0.4348 | 1.7321 | 0.7531 | 0 | 0 | 0.25103 | 4.20E-05 | *** |
| MTA | C00170 | NT | C13-07 | 11 | 22485.55268 | 291280.8773 | 205673.7298 | 4.075431 | 0.5078 | 0.1246 | 3.48953 | 4.34901 | 4.38776 | 4.07543 | 0.00016 | *** |
| MTA | C00170 | I3sg2 | C13-07 | 11 | 0 | 0 | 0 | 0 | 0 | NA | 0 | 0 | 0 | 0 | 0.00016 | *** |
| MTA | C00170 | NT | C13-08 | 11 | 0 | 58705.71201 | 39633.82419 | 0.574015 | 0.4974 | 0.8664 | 0 | 0.87651 | 0.84553 | 0.57401 | 0.11625 | |
| MTA | C00170 | I3sg2 | C13-08 | 11 | 0 | 0 | 0 | 0 | 0 | NA | 0 | 0 | 0 | 0 | 0.11625 | |
| NAD+ | C00003 | NT | C12 PARENT | 21 | 303730.8281 | 274333.5146 | 245315.7587 | 3.517316 | 0.0802 | 0.0228 | 3.47981 | 3.46276 | 3.60938 | 3.51732 | 0.00049 | *** |
| NAD+ | C00003 | I3sg2 | C12 PARENT | 21 | 224896.6408 | 38199.3709 | 34526.99424 | 6.943981 | 0.5672 | 0.0817 | 6.7687 | 6.48514 | 7.5781 | 6.94398 | 0.00049 | *** |
| NAD+ | C00003 | NT | C13-04 | 21 | 34484.85254 | 34335.15927 | 30490.39976 | 0.425698 | 0.0276 | 0.0648 | 0.39509 | 0.43339 | 0.44861 | 0.4257 | 0.80224 | |
| NAD+ | C00003 | I3sg2 | C13-04 | 21 | 57944.42641 | 0 | 0 | 0.581317 | 1.0069 | 1.7321 | 1.74395 | 0 | 0 | 0.58132 | 0.80224 | |
| NAD+ | C00003 | NT | C13-05 | 21 | 1703257.379 | 1555179.266 | 1287179.868 | 19.36091 | 0.3704 | 0.0191 | 19.514 | 19.6302 | 18.9385 | 19.3609 | 0.00073 | *** |
| NAD+ | C00003 | I3sg2 | C13-05 | 21 | 1216926.6 | 192154.8777 | 175903.5787 | 35.952 | 3.0492 | 0.0848 | 36.6258 | 32.6223 | 38.6079 | 35.952 | 0.00073 | *** |
| NAD+ | C00003 | NT | C13-06 | 21 | 0 | 18961.37842 | 2078.901175 | 0.089975 | 0.1303 | 1.4477 | 0 | 0.23934 | 0.03059 | 0.08998 | 0.69522 | |
| NAD+ | C00003 | I3sg2 | C13-06 | 21 | 5139.808954 | 0 | 0 | 0.051564 | 0.0893 | 1.7321 | 0.15469 | 0 | 0 | 0.05156 | 0.69522 | |
| NAD+ | C00003 | NT | C13-07 | 21 | 138719.1735 | 104894.1186 | 107107.1045 | 1.496399 | 0.1494 | 0.0999 | 1.58929 | 1.32402 | 1.57589 | 1.4964 | 0.04744 | * |
| NAD+ | C00003 | I3sg2 | C13-07 | 21 | 39754.70447 | 1088.788535 | 1473.774223 | 0.56827 | 0.5485 | 0.9651 | 1.1965 | 0.18484 | 0.32347 | 0.56827 | 0.04744 | * |
| NAD+ | C00003 | NT | C13-08 | 21 | 151389.186 | 132588.4739 | 112196.8728 | 1.686271 | 0.0433 | 0.0257 | 1.73445 | 1.67359 | 1.65077 | 1.68627 | 0.46349 | |
| NAD+ | C00003 | I3sg2 | C13-08 | 21 | 81325.28907 | 0 | 3963.223201 | 1.105835 | 1.2408 | 1.122 | 2.44764 | 0 | 0.86986 | 1.10583 | 0.46349 | |
| NAD+ | C00003 | NT | C13-09 | 21 | 234473.2712 | 171744.6571 | 184405.1432 | 2.522453 | 0.3074 | 0.1219 | 2.68633 | 2.16784 | 2.71319 | 2.52245 | 0.94585 | |
| NAD+ | C00003 | I3sg2 | C13-09 | 21 | 90873.03329 | 25868.45854 | 782.4153056 | 2.432813 | 2.1262 | 0.874 | 2.735 | 4.39171 | 0.17173 | 2.43281 | 0.94585 | |
| NAD+ | C00003 | NT | C13-10 | 21 | 4977614.648 | 4625650.275 | 3969148.889 | 57.93795 | 0.7881 | 0.0136 | 57.0279 | 58.3871 | 58.3988 | 57.9379 | 0.05622 | |
| NAD+ | C00003 | I3sg2 | C13-10 | 21 | 1557238.705 | 329049.2939 | 224855.2103 | 50.69437 | 4.6453 | 0.0916 | 46.8681 | 55.863 | 49.352 | 50.6944 | 0.05622 | |
| NAD+ | C00003 | NT | C13-11 | 21 | 839987.8241 | 719582.4931 | 599258.6269 | 9.174515 | 0.411 | 0.0448 | 9.62364 | 9.0829 | 8.817 | 9.17452 | 0.00079 | *** |
| NAD+ | C00003 | I3sg2 | C13-11 | 21 | 35550.97704 | 2668.305577 | 14110.30229 | 1.539984 | 1.3832 | 0.8982 | 1.06998 | 0.453 | 3.09698 | 1.53998 | 0.00079 | *** |
| NAD+ | C00003 | NT | C13-12 | 21 | 305889.8427 | 233206.2792 | 223367.6471 | 3.244877 | 0.2828 | 0.0871 | 3.50454 | 2.94364 | 3.28645 | 3.24488 | 0.00012 | *** |
| NAD+ | C00003 | I3sg2 | C13-12 | 21 | 12945.29477 | 0 | 0 | 0.129871 | 0.2249 | 1.7321 | 0.38961 | 0 | 0 | 0.12987 | 0.00012 | *** |
| NAD+ | C00003 | NT | C13-13 | 21 | 38830.5185 | 51912.08224 | 36075.04397 | 0.543638 | 0.1058 | 0.1946 | 0.44488 | 0.65526 | 0.53078 | 0.54364 | 0.00088 | *** |
| NAD+ | C00003 | I3sg2 | C13-13 | 21 | 0 | 0 | 0 | 0 | 0 | NA | 0 | 0 | 0 | 0 | 0.00088 | *** |
| NAD+ | C00003 | NT | C13-14 | 21 | 0 | 0 | 0 | 0 | 0 | NA | 0 | 0 | 0 | 0 | NA | |
| NAD+ | C00003 | I3sg2 | C13-14 | 21 | 0 | 0 | 0 | 0 | 0 | NA | 0 | 0 | 0 | 0 | NA | |
| NADH | C00004 | NT | C12 PARENT | 21 | 12377.96966 | 3927.043978 | 5172.943273 | 1.936284 | 0.2146 | 0.1108 | 2.17868 | 1.85947 | 1.77071 | 1.93628 | 0.81387 | |
| NADH | C00004 | I3sg2 | C12 PARENT | 21 | 28498.25872 | 0 | 0 | 2.587647 | 4.4819 | 1.7321 | 7.76294 | 0 | 0 | 2.58765 | 0.81387 | |
| NADH | C00004 | NT | C13-04 | 21 | 736.3550367 | 0 | 3150.273536 | 0.402651 | 0.5887 | 1.4622 | 0.12961 | 0 | 1.07835 | 0.40265 | 0.97422 | |
| NADH | C00004 | I3sg2 | C13-04 | 21 | 0 | 1448.966055 | 0 | 0.384991 | 0.6668 | 1.7321 | 0 | 1.15497 | 0 | 0.38499 | 0.97422 | |
| NADH | C00004 | NT | C13-05 | 21 | 93845.96316 | 43792.68171 | 65604.69894 | 19.90355 | 3.0556 | 0.1535 | 16.518 | 20.736 | 22.4567 | 19.9036 | 0.00093 | *** |
| NADH | C00004 | I3sg2 | C13-05 | 21 | 138085.4297 | 53815.04762 | 52846.89212 | 40.46067 | 2.6646 | 0.0659 | 37.6145 | 42.896 | 40.8714 | 40.4607 | 0.00093 | *** |
| NADH | C00004 | NT | C13-06 | 21 | 0 | 0 | 0 | 0 | 0 | NA | 0 | 0 | 0 | 0 | NA | |
| NADH | C00004 | I3sg2 | C13-06 | 21 | 0 | 0 | 0 | 0 | 0 | NA | 0 | 0 | 0 | 0 | NA | |
| NADH | C00004 | NT | C13-07 | 21 | 3334.406578 | 0 | 415.9177419 | 0.243089 | 0.3061 | 1.2594 | 0.5869 | 0 | 0.14237 | 0.24309 | 0.57841 | |
| NADH | C00004 | I3sg2 | C13-07 | 21 | 7157.658044 | 0 | 0 | 0.649917 | 1.1257 | 1.7321 | 1.94975 | 0 | 0 | 0.64992 | 0.57841 | |
| NADH | C00004 | NT | C13-08 | 21 | 10328.98913 | 5480.487155 | 1954.309969 | 1.694007 | 0.969 | 0.572 | 1.81803 | 2.59503 | 0.66897 | 1.69401 | 0.50342 | |
| NADH | C00004 | I3sg2 | C13-08 | 21 | 10018.51218 | 0 | 0 | 0.909683 | 1.5756 | 1.7321 | 2.72905 | 0 | 0 | 0.90968 | 0.50342 | |
| NADH | C00004 | NT | C13-09 | 21 | 0 | 977.1455693 | 0 | 0.154227 | 0.2671 | 1.7321 | 0 | 0.46268 | 0 | 0.15423 | 0.49312 | |
| NADH | C00004 | I3sg2 | C13-09 | 21 | 5609.609164 | 0 | 132.5405938 | 0.543522 | 0.8542 | 1.5716 | 1.52806 | 0 | 0.10251 | 0.54352 | 0.49312 | |
| NADH | C00004 | NT | C13-10 | 21 | 408719.6142 | 131697.3467 | 182436.4044 | 65.58239 | 5.5058 | 0.084 | 71.9397 | 62.359 | 62.4484 | 65.5824 | 0.03863 | * |
| NADH | C00004 | I3sg2 | C13-10 | 21 | 177092.7842 | 70190.60656 | 71940.90808 | 53.27594 | 4.3639 | 0.0819 | 48.2402 | 55.949 | 55.6386 | 53.2759 | 0.03863 | * |
| NADH | C00004 | NT | C13-11 | 21 | 26671.97196 | 23260.14652 | 25590.0142 | 8.155955 | 3.2025 | 0.3927 | 4.6946 | 11.0137 | 8.75953 | 8.15595 | 0.0119 | * |
| NADH | C00004 | I3sg2 | C13-11 | 21 | 644.2092097 | 0 | 0 | 0.058494 | 0.1013 | 1.7321 | 0.17548 | 0 | 0 | 0.05849 | 0.0119 | * |
| NADH | C00004 | NT | C13-12 | 21 | 12126.67411 | 2057.189079 | 7814.716677 | 1.927842 | 0.8691 | 0.4508 | 2.13444 | 0.97408 | 2.675 | 1.92784 | 0.55323 | |
| NADH | C00004 | I3sg2 | C13-12 | 21 | 0 | 0 | 4379.935545 | 1.129138 | 1.9557 | 1.7321 | 0 | 0 | 3.38741 | 1.12914 | 0.55323 | |
| NADH | C00004 | NT | C13-13 | 21 | 0 | 0 | 0 | 0 | 0 | NA | 0 | 0 | 0 | 0 | NA | |
| NADH | C00004 | I3sg2 | C13-13 | 21 | 0 | 0 | 0 | 0 | 0 | NA | 0 | 0 | 0 | 0 | NA | |
| NADP+ | C00006 | NT | C12 PARENT | 21 | 45102.28408 | 14966.78807 | 30033.34009 | 3.896477 | 0.4632 | 0.1189 | 4.06124 | 3.37339 | 4.2548 | 3.89648 | 0.00395 | ** |
| NADP+ | C00006 | I3sg2 | C12 PARENT | 21 | 24456.46294 | 6383.555441 | 3783.749261 | 7.905163 | 1.0661 | 0.1349 | 7.87016 | 8.98834 | 6.85699 | 7.90516 | 0.00395 | ** |
| NADP+ | C00006 | NT | C13-04 | 21 | 5430.189272 | 2021.900264 | 3738.493186 | 0.491437 | 0.037 | 0.0753 | 0.48896 | 0.45572 | 0.52963 | 0.49144 | 0.17008 | |
| NADP+ | C00006 | I3sg2 | C13-04 | 21 | 1108.281886 | 1855.606115 | 1003.448555 | 1.595966 | 1.1444 | 0.7171 | 0.35665 | 2.61278 | 1.81847 | 1.59597 | 0.17008 | |
| NADP+ | C00006 | NT | C13-05 | 21 | 212050.4372 | 85682.62868 | 136193.4918 | 19.23356 | 0.1211 | 0.0063 | 19.0941 | 19.3121 | 19.2944 | 19.2336 | 9.74E-07 | *** |
| NADP+ | C00006 | I3sg2 | C13-05 | 21 | 115396.0997 | 25871.60252 | 19846.32274 | 36.50971 | 0.5887 | 0.0161 | 37.1348 | 36.4284 | 35.9659 | 36.5097 | 9.74E-07 | *** |
| NADP+ | C00006 | NT | C13-06 | 21 | 9707.074576 | 798.8753448 | 2981.233928 | 0.492161 | 0.3522 | 0.7157 | 0.87407 | 0.18006 | 0.42235 | 0.49216 | 0.17559 | |
| NADP+ | C00006 | I3sg2 | C13-06 | 21 | 1036.621105 | 0 | 0 | 0.111196 | 0.1926 | 1.7321 | 0.33359 | 0 | 0 | 0.1112 | 0.17559 | |
| NADP+ | C00006 | NT | C13-07 | 21 | 11671.01771 | 5684.762181 | 11789.29091 | 1.334132 | 0.313 | 0.2346 | 1.05092 | 1.2813 | 1.67018 | 1.33413 | 0.08197 | |
| NADP+ | C00006 | I3sg2 | C13-07 | 21 | 2259.784188 | 667.0948461 | 0 | 0.555502 | 0.4926 | 0.8868 | 0.72721 | 0.9393 | 0 | 0.5555 | 0.08197 | |
| NADP+ | C00006 | NT | C13-08 | 21 | 18653.52584 | 7813.760737 | 12766.45056 | 1.74981 | 0.0652 | 0.0373 | 1.67966 | 1.76115 | 1.80861 | 1.74981 | 0.88681 | |
| NADP+ | C00006 | I3sg2 | C13-08 | 21 | 8241.310839 | 239.4525492 | 1439.330596 | 1.865874 | 1.3241 | 0.7096 | 2.65208 | 0.33716 | 2.60838 | 1.86587 | 0.88681 | |
| NADP+ | C00006 | NT | C13-09 | 21 | 25539.27638 | 9576.822576 | 15713.59242 | 2.228119 | 0.0706 | 0.0317 | 2.29969 | 2.15853 | 2.22613 | 2.22812 | 0.00908 | ** |
| NADP+ | C00006 | I3sg2 | C13-09 | 21 | 3606.348883 | 0 | 0 | 0.386845 | 0.67 | 1.7321 | 1.16053 | 0 | 0 | 0.38684 | 0.00908 | ** |
| NADP+ | C00006 | NT | C13-10 | 21 | 641085.5419 | 258215.1465 | 406701.8295 | 57.84777 | 0.3095 | 0.0053 | 57.7266 | 58.1995 | 57.6172 | 57.8478 | 0.00572 | ** |
| NADP+ | C00006 | I3sg2 | C13-10 | 21 | 149434.6038 | 36003.09191 | 29108.09484 | 50.51093 | 2.3363 | 0.0463 | 48.0885 | 50.694 | 52.7503 | 50.5109 | 0.00572 | ** |
| NADP+ | C00006 | NT | C13-11 | 21 | 95162.22257 | 42552.92171 | 55551.49404 | 8.676634 | 0.8656 | 0.0998 | 8.5689 | 9.59106 | 7.86994 | 8.67663 | 0.00041 | *** |
| NADP+ | C00006 | I3sg2 | C13-11 | 21 | 5209.604408 | 0 | 0 | 0.558822 | 0.9679 | 1.7321 | 1.67647 | 0 | 0 | 0.55882 | 0.00041 | *** |
| NADP+ | C00006 | NT | C13-12 | 21 | 39622.78362 | 14529.55286 | 25271.63798 | 3.474297 | 0.1728 | 0.0498 | 3.56784 | 3.27484 | 3.58022 | 3.4743 | 4.06E-06 | *** |
| NADP+ | C00006 | I3sg2 | C13-12 | 21 | 0 | 0 | 0 | 0 | 0 | NA | 0 | 0 | 0 | 0 | 4.06E-06 | *** |
| NADP+ | C00006 | NT | C13-13 | 21 | 6529.750479 | 1829.433464 | 5128.203485 | 0.575607 | 0.1574 | 0.2735 | 0.58797 | 0.41234 | 0.72651 | 0.57561 | 0.00318 | ** |
| NADP+ | C00006 | I3sg2 | C13-13 | 21 | 0 | 0 | 0 | 0 | 0 | NA | 0 | 0 | 0 | 0 | 0.00318 | ** |
| NADPH | C00005 | NT | C12 PARENT | 21 | 0 | 3883.534515 | 1906.190995 | 4.091291 | 1.0713 | 0.2618 | 0 | 4.8488 | 3.33378 | 4.09129 | 0.8816 | |
| NADPH | C00005 | I3sg2 | C12 PARENT | 21 | 4940.373644 | 0 | 0 | 9.894468 | NA | NA | 9.89447 | 0 | 0 | 9.89447 | 0.8816 | |
| NADPH | C00005 | NT | C13-05 | 21 | 0 | 16770.65143 | 11667.4769 | 20.67229 | 0.3772 | 0.0182 | 0 | 20.939 | 20.4055 | 20.6723 | 0.96634 | |
| NADPH | C00005 | I3sg2 | C13-05 | 21 | 19647.10461 | 0 | 0 | 39.34877 | NA | NA | 39.3488 | 0 | 0 | 39.3488 | 0.96634 | |
| NADPH | C00005 | NT | C13-06 | 21 | 0 | 0 | 0 | 0 | 0 | NA | 0 | 0 | 0 | 0 | NA | |
| NADPH | C00005 | I3sg2 | C13-06 | 21 | 0 | 0 | 0 | 0 | NA | NA | 0 | 0 | 0 | 0 | NA | |
| NADPH | C00005 | NT | C13-07 | 21 | 0 | 1598.756185 | 868.88835 | 1.757876 | 0.3369 | 0.1917 | 0 | 1.99613 | 1.51962 | 1.75788 | 0.12337 | |
| NADPH | C00005 | I3sg2 | C13-07 | 21 | 0 | 0 | 0 | 0 | NA | NA | 0 | 0 | 0 | 0 | 0.12337 | |

| | | | | | | | | | | | | | | | | |
|---|---|---|---|---|---|---|---|---|---|---|---|---|---|---|---|---|
| NADPH | C00005 | NT | C13-09 | 21 | 0 | 2623.333238 | 839.5728335 | 2.37186 | 1.2778 | 0.5387 | 0 | 3.27537 | 1.46835 | 2.37186 | 0.53259 | |
| NADPH | C00005 | I3sg2 | C13-09 | 21 | 1130.865587 | 0 | 0 | 2.264872 | NA | NA | 2.26487 | 0 | 0 | 2.26487 | 0.53259 | |
| NADPH | C00005 | NT | C13-10 | 21 | 0 | 46683.65436 | 34501.02016 | 59.31335 | 1.4515 | 0.0245 | 0 | 58.287 | 60.3397 | 59.3133 | 0.40635 | |
| NADPH | C00005 | I3sg2 | C13-10 | 21 | 23925.0093 | 0 | 0 | 47.91646 | NA | NA | 47.9165 | 0 | 0 | 47.9165 | 0.40635 | |
| NADPH | C00005 | NT | C13-11 | 21 | 0 | 6217.220338 | 5357.206417 | 8.565939 | 1.1362 | 0.1326 | 0 | 7.76253 | 9.36935 | 8.56594 | 0.12969 | |
| NADPH | C00005 | I3sg2 | C13-11 | 21 | 287.312622 | 0 | 0 | 0.575423 | NA | NA | 0.57542 | 0 | 0 | 0.57542 | 0.12969 | |
| NADPH | C00005 | NT | C13-12 | 21 | 0 | 1869.107706 | 1841.080877 | 2.776796 | 0.6267 | 0.2257 | 0 | 2.33368 | 3.21991 | 2.7768 | 0.12615 | |
| NADPH | C00005 | I3sg2 | C13-12 | 21 | 0 | 0 | 0 | 0 | NA | NA | 0 | 0 | 0 | 0 | 0.12615 | |
| NADPH | C00005 | NT | C13-13 | 21 | 0 | 446.4640993 | 196.5598637 | 0.450601 | 0.1511 | 0.3353 | 0 | 0.55743 | 0.34377 | 0.4506 | 0.13798 | |
| NADPH | C00005 | I3sg2 | C13-13 | 21 | 0 | 0 | 0 | 0 | NA | NA | 0 | 0 | 0 | 0 | 0.13798 | |
| Orotate | C00295 | NT | C12 PARENT | 5 | 549053.1506 | 93763.2654 | 797753.3236 | 80.10405 | 9.5865 | 0.1197 | 75.5027 | 91.1237 | 73.6857 | 80.104 | 0.33198 | |
| Orotate | C00295 | I3sg2 | C12 PARENT | 5 | 149319.7257 | 376259.7277 | 523965.75 | 73.90617 | 1.688 | 0.0228 | 75.3118 | 74.3727 | 72.034 | 73.9062 | 0.33198 | |
| Orotate | C00295 | NT | C13-01 | 5 | 0 | 0 | 0 | 0 | 0 | NA | 0 | 0 | 0 | 0 | NA | |
| Orotate | C00295 | I3sg2 | C13-01 | 5 | 0 | 0 | 0 | 0 | 0 | NA | 0 | 0 | 0 | 0 | NA | |
| Orotate | C00295 | NT | C13-02 | 5 | 8855.434837 | 0 | 4098.618383 | 0.532108 | 0.6232 | 1.1712 | 1.21775 | 0 | 0.37858 | 0.53211 | 0.50977 | |
| Orotate | C00295 | I3sg2 | C13-02 | 5 | 0 | 3418.684209 | 0 | 0.225249 | 0.3901 | 1.7321 | 0 | 0.67575 | 0 | 0.22525 | 0.50977 | |
| Orotate | C00295 | NT | C13-03 | 5 | 169287.9846 | 9133.410385 | 280791.1604 | 19.36385 | 9.1791 | 0.474 | 23.2795 | 8.87629 | 25.9357 | 19.3638 | 0.29498 | |
| Orotate | C00295 | I3sg2 | C13-03 | 5 | 48948.98402 | 126232.6314 | 203421.013 | 25.86859 | 1.8212 | 0.0704 | 24.6882 | 24.9515 | 27.966 | 25.8686 | 0.29498 | |
| Orotidine | X00006 | NT | C12 PARENT | 10 | 3930276.423 | 3077075.681 | 3049312.442 | 39.97892 | 0.8086 | 0.0202 | 39.0499 | 40.3625 | 40.5244 | 39.9789 | 7.33E-06 | *** |
| Orotidine | X00006 | I3sg2 | C12 PARENT | 10 | 1381921.676 | 1527798.735 | 1514755.417 | 61.15057 | 0.9154 | 0.015 | 61.304 | 60.1681 | 61.9796 | 61.1506 | 7.33E-06 | *** |
| Orotidine | X00006 | NT | C13-01 | 10 | 0 | 0 | 0 | 0 | 0 | NA | 0 | 0 | 0 | 0 | NA | |
| Orotidine | X00006 | I3sg2 | C13-01 | 10 | 0 | 0 | 0 | 0 | 0 | NA | 0 | 0 | 0 | 0 | NA | |
| Orotidine | X00006 | NT | C13-04 | 10 | 99733.59091 | 60438.6763 | 34402.32023 | 0.746966 | 0.2698 | 0.3612 | 0.99092 | 0.79278 | 0.4572 | 0.74697 | 0.22699 | |
| Orotidine | X00006 | I3sg2 | C13-04 | 10 | 0 | 0 | 20856.48616 | 0.284463 | 0.4927 | 1.7321 | 0 | 0 | 0.85339 | 0.28446 | 0.22699 | |
| Orotidine | X00006 | NT | C13-05 | 10 | 3858315.461 | 2897830.19 | 2856949.189 | 38.10471 | 0.2005 | 0.0053 | 38.3349 | 38.0113 | 37.9679 | 38.1047 | 3.19E-05 | *** |
| Orotidine | X00006 | I3sg2 | C13-05 | 10 | 599490.8418 | 719250.0324 | 667351.6106 | 27.4087 | 0.8702 | 0.0318 | 26.5943 | 28.3257 | 27.3062 | 27.4087 | 3.19E-05 | *** |
| Orotidine | X00006 | NT | C13-06 | 10 | 174688.5001 | 168613.2685 | 114479.9233 | 1.822925 | 0.3533 | 0.1938 | 1.73564 | 2.21173 | 1.5214 | 1.82292 | 0.00173 | ** |
| Orotidine | X00006 | I3sg2 | C13-06 | 10 | 268.0560385 | 4425.382245 | 7705.677307 | 0.167156 | 0.1518 | 0.9083 | 0.01189 | 0.17428 | 0.31529 | 0.16716 | 0.00173 | ** |
| Orotidine | X00006 | NT | C13-07 | 10 | 914242.1919 | 652848.402 | 658288.5372 | 8.798519 | 0.2636 | 0.03 | 9.08359 | 8.56352 | 8.74844 | 8.79852 | 0.01039 | * |
| Orotidine | X00006 | I3sg2 | C13-07 | 10 | 147083.5637 | 154085.3646 | 105055.8974 | 5.630552 | 1.1759 | 0.2088 | 6.52484 | 6.06822 | 4.29859 | 5.63055 | 0.01039 | * |
| Orotidine | X00006 | NT | C13-08 | 10 | 855951.971 | 580199.5158 | 614526.142 | 8.093957 | 0.4514 | 0.0558 | 8.50444 | 7.61057 | 8.16685 | 8.09396 | 0.00062 | *** |
| Orotidine | X00006 | I3sg2 | C13-08 | 10 | 125445.8806 | 133657.7961 | 128233.9549 | 5.358559 | 0.1789 | 0.0334 | 5.56496 | 5.26374 | 5.24698 | 5.35856 | 0.00062 | *** |
| Orotidine | X00006 | NT | C13-09 | 10 | 231554.5152 | 186592.1556 | 196678.9136 | 2.454002 | 0.1567 | 0.0638 | 2.30065 | 2.44756 | 2.6138 | 2.454 | 1.10E-05 | *** |
| Orotidine | X00006 | I3sg2 | C13-09 | 10 | 0 | 0 | 0 | 0 | 0 | NA | 0 | 0 | 0 | 0 | 1.10E-05 | *** |
| P-Creatine | C02305 | NT | C12 PARENT | 4 | 7719823.298 | 5598284.523 | 5267442.245 | 96.64185 | 0.1248 | 0.0013 | 96.7277 | 96.4987 | 96.6991 | 96.6419 | 0.00034 | *** |
| P-Creatine | C02305 | I3sg2 | C12 PARENT | 4 | 5568167.243 | 7424056.15 | 7773537.674 | 99.06916 | 0.3479 | 0.0035 | 98.6705 | 99.3116 | 99.2254 | 99.0692 | 0.00034 | *** |
| P-Creatine | C02305 | NT | C13-01 | 4 | 0 | 0 | 0 | 0 | 0 | NA | 0 | 0 | 0 | 0 | NA | |
| P-Creatine | C02305 | I3sg2 | C13-01 | 4 | 0 | 0 | 0 | 0 | 0 | NA | 0 | 0 | 0 | 0 | NA | |
| P-Creatine | C02305 | NT | C13-02 | 4 | 261163.1878 | 203123.0336 | 179805.6743 | 3.358147 | 0.1248 | 0.0372 | 3.27232 | 3.50127 | 3.30085 | 3.35815 | 0.00034 | *** |
| P-Creatine | C02305 | I3sg2 | C13-02 | 4 | 75025.31809 | 51462.01924 | 60685.48562 | 0.930837 | 0.3479 | 0.3738 | 1.32948 | 0.68841 | 0.77462 | 0.93084 | 0.00034 | *** |
| P-Ser | C01005 | NT | C12 PARENT | 3 | 333869.2473 | 271392.7644 | 275905.2986 | 100 | 0 | 0 | 100 | 100 | 100 | 100 | NA | |
| P-Ser | C01005 | I3sg2 | C12 PARENT | 3 | 115925.328 | 121519.5251 | 135731.1334 | 100 | 0 | 0 | 100 | 100 | 100 | 100 | NA | |
| PEP | C00074 | NT | C12 PARENT | 3 | 17207.28415 | 57845.30121 | 24400.04044 | 49.6151 | 38.9623 | 0.7853 | 4.64847 | 73.3496 | 70.8473 | 49.6151 | 0.58165 | |
| PEP | C00074 | I3sg2 | C12 PARENT | 3 | 39780.09323 | 0 | 23704.63179 | 29.10512 | 44.7569 | 1.5378 | 80.6422 | 0 | 6.6732 | 29.1051 | 0.58165 | |
| PEP | C00074 | NT | C13-03 | 3 | 352963.8667 | 21017.18412 | 10040.30403 | 50.3849 | 38.9623 | 0.7733 | 95.3515 | 26.6504 | 29.1527 | 50.3849 | 0.58165 | |
| PEP | C00074 | I3sg2 | C13-03 | 3 | 9549.052243 | 481936.8486 | 331516.6039 | 70.89488 | 44.7569 | 0.6313 | 19.3578 | 100 | 93.3268 | 70.8949 | 0.58165 | |
| Pro | C00148 | NT | C12 PARENT | 5 | 136010322.5 | 281272965.6 | 262603013.7 | 77.69581 | 0.4528 | 0.0058 | 77.2045 | 77.7866 | 78.0964 | 77.6958 | 0.00266 | ** |
| Pro | C00148 | I3sg2 | C12 PARENT | 5 | 72771935.93 | 146915559.2 | 134546210.8 | 84.48372 | 1.7103 | 0.0202 | 82.8981 | 84.2571 | 86.296 | 84.4837 | 0.00266 | ** |
| Pro | C00148 | NT | C13-01 | 5 | 3475451.141 | 6783876.627 | 6697931.762 | 1.946936 | 0.0621 | 0.0319 | 1.97279 | 1.87609 | 1.99192 | 1.94694 | 0.52871 | |
| Pro | C00148 | I3sg2 | C13-01 | 5 | 1765665.919 | 3774321.435 | 1097921.8 | 1.626716 | 0.8026 | 0.4934 | 2.01136 | 2.1646 | 0.70419 | 1.62672 | 0.52871 | |
| Pro | C00148 | NT | C13-03 | 5 | 10961552.78 | 23367701.97 | 21334228.93 | 6.343074 | 0.1201 | 0.0189 | 6.22218 | 6.46238 | 6.34466 | 6.34307 | 0.00382 | ** |
| Pro | C00148 | I3sg2 | C13-03 | 5 | 4653706.671 | 8120389.84 | 7036443.303 | 4.823813 | 0.4197 | 0.087 | 5.30126 | 4.6571 | 4.51307 | 4.82381 | 0.00382 | ** |
| Pro | C00148 | NT | C13-04 | 5 | 15975067.84 | 30530107.16 | 27813718.86 | 8.594271 | 0.4192 | 0.0488 | 9.06804 | 8.44316 | 8.27161 | 8.59427 | 0.00094 | *** |
| Pro | C00148 | I3sg2 | C13-04 | 5 | 5508836.026 | 10174627.84 | 8794990.541 | 5.917195 | 0.325 | 0.0549 | 6.27538 | 5.83522 | 5.64098 | 5.91719 | 0.00094 | *** |
| Pro | C00148 | NT | C13-05 | 5 | 9746587.857 | 19641031.57 | 17806165.96 | 5.419908 | 0.119 | 0.022 | 5.53252 | 5.43177 | 5.29543 | 5.41991 | 0.00039 | *** |
| Pro | C00148 | I3sg2 | C13-05 | 5 | 3084692.553 | 5380889.787 | 4436895.721 | 3.148554 | 0.3384 | 0.1075 | 3.51392 | 3.08598 | 2.84576 | 3.14855 | 0.00039 | *** |
| Propanoyl-CoA | C00100 | NT | C12 PARENT | 24 | 3955.164866 | 841.9121385 | 3175.947621 | 15.9683 | 2.3343 | 0.1462 | 17.4158 | 13.2754 | 17.2137 | 15.9683 | 0.04359 | * |
| Propanoyl-CoA | C00100 | I3sg2 | C12 PARENT | 24 | 3202.020546 | 4512.53792 | 1636.673633 | 34.482 | 10.7613 | 0.3121 | 41.2604 | 40.112 | 22.0736 | 34.482 | 0.04359 | * |
| Propanoyl-CoA | C00100 | NT | C13-05 | 24 | 15977.98551 | 4856.920527 | 13521.8923 | 73.40987 | 3.1162 | 0.0424 | 70.356 | 76.5848 | 73.2889 | 73.4099 | 0.2631 | |
| Propanoyl-CoA | C00100 | I3sg2 | C13-05 | 24 | 4558.499084 | 6170.00084 | 5777.958048 | 63.8371 | 12.3561 | 0.1936 | 58.7396 | 54.8453 | 77.9264 | 63.8371 | 0.2631 | |
| Propanoyl-CoA | C00100 | NT | C13-06 | 24 | 2777.06153 | 643.054555 | 1752.287872 | 10.62183 | 1.4278 | 0.1344 | 12.2283 | 10.1398 | 9.49743 | 10.6218 | 0.0088 | ** |
| Propanoyl-CoA | C00100 | I3sg2 | C13-06 | 24 | 0 | 567.2949549 | 0 | 1.680899 | 2.9114 | 1.7321 | 0 | 5.0427 | 0 | 1.6809 | 0.0088 | ** |
| S7P | C05382 | NT | C12 PARENT | 7 | 0 | 16712.42352 | 0 | 1.713823 | 2.9684 | 1.7321 | 0 | 5.14147 | 0 | 1.71382 | 0.02755 | * |
| S7P | C05382 | I3sg2 | C12 PARENT | 7 | 63357.14482 | 21710.69007 | 73571.00172 | 21.17172 | 9.49 | 0.4482 | 25.1384 | 10.342 | 28.0347 | 21.1717 | 0.02755 | * |
| S7P | C05382 | NT | C13-07 | 7 | 337053.7786 | 308339.0636 | 415717.0208 | 98.28618 | 2.9684 | 0.0302 | 100 | 94.8585 | 100 | 98.2862 | 0.02755 | * |
| S7P | C05382 | I3sg2 | C13-07 | 7 | 188675.8189 | 188216.5074 | 188857.0853 | 78.82828 | 9.49 | 0.1204 | 74.8616 | 89.658 | 71.9653 | 78.8283 | 0.02755 | * |
| SAM | C00019 | NT | C12 PARENT | 15 | 1103039.072 | 827596.9461 | 919892.1433 | 11.34093 | 0.2787 | 0.0246 | 11.0327 | 11.5754 | 11.4147 | 11.3409 | 7.86E-07 | *** |
| SAM | C00019 | I3sg2 | C12 PARENT | 15 | 910304.2508 | 1473277.91 | 1512450.178 | 25.96704 | 0.3935 | 0.0152 | 25.6778 | 25.8082 | 26.4152 | 25.967 | 7.86E-07 | *** |
| SAM | C00019 | NT | C13-01 | 15 | 0 | 0 | 0 | 0 | 0 | NA | 0 | 0 | 0 | 0 | NA | |
| SAM | C00019 | I3sg2 | C13-01 | 15 | 0 | 0 | 0 | 0 | 0 | NA | 0 | 0 | 0 | 0 | NA | |
| SAM | C00019 | NT | C13-02 | 15 | 11521.89906 | 11748.02326 | 6540.168714 | 0.120238 | 0.0418 | 0.3477 | 0.11524 | 0.16432 | 0.08116 | 0.12024 | 0.00759 | ** |
| SAM | C00019 | I3sg2 | C13-02 | 15 | 0 | 0 | 0 | 0 | 0 | NA | 0 | 0 | 0 | 0 | 0.00759 | ** |
| SAM | C00019 | NT | C13-03 | 15 | 95567.00488 | 73332.62102 | 85169.1669 | 1.012799 | 0.0517 | 0.051 | 0.95587 | 1.02568 | 1.05684 | 1.0128 | 0.09563 | |
| SAM | C00019 | I3sg2 | C13-03 | 15 | 61048.26206 | 61298.57509 | 85599.50554 | 1.430284 | 0.3289 | 0.23 | 1.72204 | 1.0738 | 1.49501 | 1.43028 | 0.09563 | |
| SAM | C00019 | NT | C13-04 | 15 | 192159.6864 | 81739.91431 | 138428.4949 | 1.594333 | 0.4038 | 0.2532 | 1.922 | 1.14327 | 1.71772 | 1.59433 | 0.31617 | |
| SAM | C00019 | I3sg2 | C13-04 | 15 | 70511.04268 | 111690.1735 | 97230.05038 | 1.881214 | 0.1594 | 0.0847 | 1.98897 | 1.95654 | 1.69814 | 1.88121 | 0.31617 | |
| SAM | C00019 | NT | C13-05 | 15 | 6912435.433 | 4962593.894 | 5666672.583 | 69.62186 | 0.6164 | 0.0089 | 69.139 | 69.4104 | 70.3162 | 69.6219 | 0.29676 | |
| SAM | C00019 | I3sg2 | C13-05 | 15 | 2487592.077 | 4062303.506 | 3985583.048 | 70.31335 | 0.7862 | 0.0112 | 70.1698 | 71.1615 | 69.6088 | 70.3133 | 0.29676 | |
| SAM | C00019 | NT | C13-06 | 15 | 1537591.918 | 1081364.032 | 1127265.48 | 14.83061 | 0.7408 | 0.05 | 15.3792 | 15.1247 | 13.9879 | 14.8306 | 7.55E-06 | *** |
| SAM | C00019 | I3sg2 | C13-06 | 15 | 15650.15903 | 0 | 44825.25258 | 0.408113 | 0.3925 | 0.9618 | 0.44146 | 0 | 0.78288 | 0.40811 | 7.55E-06 | *** |
| SAM | C00019 | NT | C13-07 | 15 | 109366.1305 | 86290.95905 | 79526.13993 | 1.095879 | 0.1101 | 0.1004 | 1.09389 | 1.20693 | 0.98682 | 1.09588 | 6.63E-05 | *** |
| SAM | C00019 | I3sg2 | C13-07 | 15 | 0 | 0 | 0 | 0 | 0 | NA | 0 | 0 | 0 | 0 | 6.63E-05 | *** |
| SAM | C00019 | NT | C13-08 | 15 | 36198.18285 | 24973.85989 | 35352.85464 | 0.383348 | 0.0483 | 0.1261 | 0.36206 | 0.3493 | 0.43868 | 0.38335 | 0.00016 | *** |
| SAM | C00019 | I3sg2 | C13-08 | 15 | 0 | 0 | 0 | 0 | 0 | NA | 0 | 0 | 0 | 0 | 0.00016 | *** |
| Ser | C00065 | NT | C12 PARENT | 3 | 42302088.38 | 43136166.12 | 40503713.32 | 75.59225 | 0.3195 | 0.0042 | 75.2705 | 75.9095 | 75.5967 | 75.5922 | 3.74E-07 | *** |
| Ser | C00065 | I3sg2 | C12 PARENT | 3 | 37800909.68 | 40855340.27 | 41053481.06 | 90.14795 | 0.2383 | 0.0026 | 90.2241 | 89.8809 | 90.3389 | 90.1479 | 3.74E-07 | *** |
| Ser | C00065 | NT | C13-01 | 3 | 5351203.233 | 5580943.266 | 5111462.474 | 9.627652 | 0.1678 | 0.0174 | 9.5217 | 9.82115 | 9.5401 | 9.62765 | 9.21E-07 | *** |
| Ser | C00065 | I3sg2 | C13-01 | 3 | 1641861.529 | 1791916.224 | 1704723.468 | 3.870761 | 0.1041 | 0.0269 | 3.91883 | 3.94218 | 3.75127 | 3.87076 | 9.21E-07 | *** |
| Ser | C00065 | NT | C13-02 | 3 | 3225609.526 | 3014892.031 | 2961465.055 | 5.524111 | 0.217 | 0.0393 | 5.73951 | 5.3055 | 5.52732 | 5.52411 | 0.00018 | *** |
| Ser | C00065 | I3sg2 | C13-02 | 3 | 1136219.998 | 1376236.232 | 1114459.158 | 2.730678 | 0.2881 | 0.1055 | 2.71196 | 3.02769 | 2.45239 | 2.73068 | 0.00018 | *** |
| Ser | C00065 | NT | C13-03 | 3 | 5321152.661 | 5093754.443 | 5002071.006 | 9.255992 | 0.2615 | 0.0283 | 9.46823 | 8.96381 | 9.33593 | 9.25599 | 5.14E-06 | *** |
| Ser | C00065 | I3sg2 | C13-03 | 3 | 1317698.36 | 1431498.976 | 1571205.344 | 3.250615 | 0.1791 | 0.0551 | 3.14511 | 3.14927 | 3.45746 | 3.25061 | 5.14E-06 | *** |
| Sorbitol | C00794 | NT | C12 PARENT | 6 | 64634312.02 | 53724675.1 | 37724761.73 | 99.01188 | 0.3189 | 0.0032 | 99.2996 | 99.067 | 98.6689 | 99.0119 | 0.08155 | |

| | | | | | | | | | | | | | | | | |
|---|---|---|---|---|---|---|---|---|---|---|---|---|---|---|---|---|
| Sorbitol | C00794 | I3sg2 | C12 PARENT | 6 | 56007017.4 | 51552694.95 | 43401682 | 99.48357 | 0.1509 | 0.0015 | 99.6486 | 99.3525 | 99.4497 | 99.4836 | 0.08155 | |
| Sorbitol | C00794 | NT | C13-01 | 6 | 0 | 0 | 0 | 0 | 0 | NA | 0 | 0 | 0 | 0 | NA | |
| Sorbitol | C00794 | I3sg2 | C13-01 | 6 | 0 | 0 | 0 | 0 | 0 | NA | 0 | 0 | 0 | 0 | NA | |
| Sorbitol | C00794 | NT | C13-06 | 6 | 455863.4205 | 505944.7302 | 508909.4352 | 0.988119 | 0.3189 | 0.3228 | 0.70036 | 0.93295 | 1.33105 | 0.98812 | 0.08155 | |
| Sorbitol | C00794 | I3sg2 | C13-06 | 6 | 197523.5273 | 336003.2056 | 240169.5857 | 0.516434 | 0.1509 | 0.2923 | 0.35144 | 0.64755 | 0.55032 | 0.51643 | 0.08155 | |
| Succ | C00042 | NT | C12 PARENT | 4 | 1856443.661 | 1749518.206 | 1568777.467 | 69.107 | 2.7779 | 0.0402 | 70.1427 | 65.9601 | 71.2183 | 69.107 | 0.0105 | * |
| Succ | C00042 | I3sg2 | C12 PARENT | 4 | 1748343.847 | 1731113.494 | 2141830.114 | 81.9678 | 4.0451 | 0.0494 | 79.2061 | 80.0863 | 86.611 | 81.9678 | 0.0105 | * |
| Succ | C00042 | NT | C13-01 | 4 | 0 | 12866.03266 | 44981.46022 | 0.84237 | 1.0669 | 1.2665 | 0 | 0.48507 | 2.04204 | 0.84237 | 0.67176 | |
| Succ | C00042 | I3sg2 | C13-01 | 4 | 3865.199813 | 28805.44495 | 0 | 0.502577 | 0.7242 | 1.4409 | 0.17511 | 1.33262 | 0 | 0.50258 | 0.67176 | |
| Succ | C00042 | NT | C13-02 | 4 | 429515.2905 | 508843.5269 | 334193.2741 | 16.86145 | 2.08 | 0.1234 | 16.2285 | 19.1843 | 15.1715 | 16.8614 | 0.04787 | * |
| Succ | C00042 | I3sg2 | C13-02 | 4 | 289787.6908 | 257495.0815 | 178279.6496 | 10.75004 | 3.1261 | 0.2908 | 13.1284 | 11.9125 | 7.20925 | 10.75 | 0.04787 | * |
| Succ | C00042 | NT | C13-03 | 4 | 240346.8043 | 216848.5124 | 143854.8815 | 7.929107 | 1.293 | 0.1631 | 9.08111 | 8.17559 | 6.53062 | 7.92911 | 0.00906 | ** |
| Succ | C00042 | I3sg2 | C13-03 | 4 | 94427.8056 | 99563.44664 | 95495.95623 | 4.248551 | 0.3731 | 0.0878 | 4.27791 | 4.60609 | 3.86165 | 4.24855 | 0.00906 | ** |
| Succ | C00042 | NT | C13-04 | 4 | 120362.3732 | 164313.6332 | 110967.1204 | 5.260076 | 0.8459 | 0.1608 | 4.54769 | 6.19493 | 5.03761 | 5.26008 | 0.01043 | * |
| Succ | C00042 | I3sg2 | C13-04 | 4 | 70910.14675 | 44582.65878 | 57325.01078 | 2.531034 | 0.6038 | 0.2386 | 3.21248 | 2.06252 | 2.3181 | 2.53103 | 0.01043 | * |
| Succinyl-CoA | C00091 | NT | C12 PARENT | 25 | 3683.568186 | 398.0889308 | 2173.385311 | 12.51194 | 1.2084 | 0.0966 | 12.3903 | 13.7765 | 11.369 | 12.5119 | 0.00036 | *** |
| Succinyl-CoA | C00091 | I3sg2 | C12 PARENT | 25 | 1956.935273 | 4772.094458 | 6001.986216 | 33.77916 | 3.0467 | 0.0902 | 31.3904 | 32.7367 | 37.2103 | 33.7792 | 0.00036 | *** |
| Succinyl-CoA | C00091 | NT | C13-05 | 25 | 17128.29141 | 2425.628231 | 13249.32285 | 70.28802 | 13.1918 | 0.1877 | 57.614 | 83.9428 | 69.3073 | 70.288 | 0.25152 | |
| Succinyl-CoA | C00091 | I3sg2 | C13-05 | 25 | 3997.308476 | 8629.093293 | 8913.902111 | 59.52613 | 4.4373 | 0.0745 | 64.1193 | 59.1959 | 55.2632 | 59.5261 | 0.25152 | |
| Succinyl-CoA | C00091 | NT | C13-06 | 25 | 5372.424205 | 65.90290746 | 2697.025721 | 11.48664 | 8.2151 | 0.7152 | 18.0711 | 2.28068 | 14.1082 | 11.4866 | 0.23155 | |
| Succinyl-CoA | C00091 | I3sg2 | C13-06 | 25 | 0 | 703.5553261 | 1214.018016 | 4.11764 | 3.813 | 0.926 | 0 | 4.82642 | 7.5265 | 4.11764 | 0.23155 | |
| Succinyl-CoA | C00091 | NT | C13-07 | 25 | 3545.130005 | 0 | 997.0466144 | 5.713404 | 5.9779 | 1.0463 | 11.9247 | 0 | 5.21556 | 5.7134 | 0.44456 | |
| Succinyl-CoA | C00091 | I3sg2 | C13-07 | 25 | 279.9306369 | 472.4358359 | 0 | 2.577063 | 2.3176 | 0.8993 | 4.49026 | 3.24093 | 0 | 2.57706 | 0.44456 | |
| UDP | C00015 | NT | C12 PARENT | 9 | 389420.254 | 123100.1436 | 347294.6949 | 5.824984 | 0.3776 | 0.0648 | 5.87419 | 5.42522 | 6.17555 | 5.82498 | 8.11E-06 | *** |
| UDP | C00015 | I3sg2 | C12 PARENT | 9 | 1464429.654 | 2593732.489 | 2492555.409 | 39.72346 | 1.97 | 0.0496 | 37.4533 | 40.983 | 40.7341 | 39.7235 | 8.11E-06 | *** |
| UDP | C00015 | NT | C13-01 | 9 | 646.4970974 | 0 | 11015.84304 | 0.068545 | 0.1104 | 1.6104 | 0.00975 | 0 | 0.19588 | 0.06854 | 0.34268 | |
| UDP | C00015 | I3sg2 | C13-01 | 9 | 0 | 0 | 0 | 0 | 0 | NA | 0 | 0 | 0 | 0 | 0.34268 | |
| UDP | C00015 | NT | C13-02 | 9 | 10982.51509 | 0 | 23273.86814 | 0.193173 | 0.2083 | 1.0783 | 0.16567 | 0 | 0.41385 | 0.19317 | 0.18348 | |
| UDP | C00015 | I3sg2 | C13-02 | 9 | 0 | 0 | 0 | 0 | 0 | NA | 0 | 0 | 0 | 0 | 0.18348 | |
| UDP | C00015 | NT | C13-03 | 9 | 51907.36787 | 8516.483797 | 21616.12595 | 0.514235 | 0.2328 | 0.4527 | 0.78299 | 0.37533 | 0.38437 | 0.51423 | 0.58841 | |
| UDP | C00015 | I3sg2 | C13-03 | 9 | 11101.69646 | 9873.433133 | 44048.41109 | 0.386597 | 0.2956 | 0.7646 | 0.28393 | 0.15601 | 0.71985 | 0.3866 | 0.58841 | |
| UDP | C00015 | NT | C13-04 | 9 | 63404.84758 | 36224.56336 | 118983.8489 | 1.556218 | 0.5807 | 0.3732 | 0.95643 | 1.59647 | 2.11575 | 1.55622 | 0.08214 | |
| UDP | C00015 | I3sg2 | C13-04 | 9 | 2198.726075 | 68065.40472 | 26424.45766 | 0.521185 | 0.5155 | 0.989 | 0.05623 | 1.07549 | 0.43184 | 0.52119 | 0.08214 | |
| UDP | C00015 | NT | C13-05 | 9 | 3791799.358 | 1357695.841 | 3180586.112 | 57.86324 | 1.738 | 0.03 | 57.1972 | 59.8358 | 56.5567 | 57.8632 | 0.00051 | *** |
| UDP | C00015 | I3sg2 | C13-05 | 9 | 1696480.704 | 2463506.511 | 2496269.356 | 41.03606 | 2.2411 | 0.0546 | 43.3881 | 38.9253 | 40.7948 | 41.0361 | 0.00051 | *** |
| UDP | C00015 | NT | C13-06 | 9 | 603470.6813 | 183771.5756 | 587137.9256 | 9.214176 | 1.1746 | 0.1275 | 9.10302 | 8.0991 | 10.4404 | 9.21418 | 0.00398 | ** |
| UDP | C00015 | I3sg2 | C13-06 | 9 | 208883.4525 | 320556.6945 | 296582.1903 | 5.084715 | 0.2483 | 0.0488 | 5.34226 | 5.06505 | 4.84684 | 5.08472 | 0.00398 | ** |
| UDP | C00015 | NT | C13-07 | 9 | 1114111.969 | 311346.0764 | 838913.3304 | 15.14824 | 1.555 | 0.1027 | 16.8058 | 13.7215 | 14.9174 | 15.1482 | 0.00242 | ** |
| UDP | C00015 | I3sg2 | C13-07 | 9 | 340874.025 | 583803.7531 | 503671.062 | 8.724556 | 0.4967 | 0.0569 | 8.71797 | 9.22456 | 8.23114 | 8.72456 | 0.00242 | ** |
| UDP | C00015 | NT | C13-08 | 9 | 603601.5689 | 248381.3035 | 494886.9998 | 9.617188 | 1.1613 | 0.1208 | 9.105 | 10.9466 | 8.80001 | 9.61719 | 0.00177 | ** |
| UDP | C00015 | I3sg2 | C13-08 | 9 | 186049.4533 | 289261.8087 | 259537.8135 | 4.523428 | 0.2616 | 0.0578 | 4.75828 | 4.57056 | 4.24145 | 4.52343 | 0.00177 | ** |
| UDP-GlcNAc | C00043 | NT | C12 PARENT | 17 | 485555.1709 | 454148.147 | 563643.345 | 2.237314 | 0.3861 | 0.1726 | 2.08049 | 1.95426 | 2.67719 | 2.23731 | 4.68E-05 | *** |
| UDP-GlcNAc | C00043 | I3sg2 | C12 PARENT | 17 | 1104831.857 | 1121655.172 | 1131998.421 | 8.76409 | 0.4594 | 0.0524 | 9.28987 | 8.44061 | 8.56179 | 8.76409 | 4.68E-05 | *** |
| UDP-GlcNAc | C00043 | NT | C13-01 | 17 | 0 | 0 | 0 | 0 | 0 | NA | 0 | 0 | 0 | 0 | NA | |
| UDP-GlcNAc | C00043 | I3sg2 | C13-01 | 17 | 0 | 0 | 0 | 0 | 0 | NA | 0 | 0 | 0 | 0 | NA | |
| UDP-GlcNAc | C00043 | NT | C13-02 | 17 | 42417.21968 | 66664.94688 | 58242.73432 | 0.248419 | 0.058 | 0.2333 | 0.18175 | 0.28687 | 0.27664 | 0.24842 | 0.00529 | ** |
| UDP-GlcNAc | C00043 | I3sg2 | C13-02 | 17 | 148842.3967 | 119595.1102 | 109425.6345 | 0.993043 | 0.2268 | 0.2283 | 1.25153 | 0.89997 | 0.82763 | 0.99304 | 0.00529 | ** |
| UDP-GlcNAc | C00043 | NT | C13-04 | 17 | 101076.7857 | 46946.10529 | 33890.50389 | 0.26536 | 0.1467 | 0.5528 | 0.43309 | 0.20202 | 0.16097 | 0.26536 | 0.67093 | |
| UDP-GlcNAc | C00043 | I3sg2 | C13-04 | 17 | 62785.02915 | 0 | 0 | 0.175974 | 0.3048 | 1.7321 | 0.52792 | 0 | 0 | 0.17597 | 0.67093 | |
| UDP-GlcNAc | C00043 | NT | C13-05 | 17 | 2251487.246 | 2456076.207 | 1994467.825 | 9.896418 | 0.5888 | 0.0595 | 9.64711 | 10.5688 | 9.4733 | 9.89642 | 0.00235 | ** |
| UDP-GlcNAc | C00043 | I3sg2 | C13-05 | 17 | 755551.9129 | 851575.9596 | 969696.8506 | 6.698481 | 0.5513 | 0.0823 | 6.35299 | 6.40823 | 7.33423 | 6.69848 | 0.00235 | ** |
| UDP-GlcNAc | C00043 | NT | C13-06 | 17 | 928262.3478 | 945296.5074 | 1026430.746 | 4.306822 | 0.4944 | 0.1148 | 3.97739 | 4.06774 | 4.87533 | 4.30682 | 0.00035 | *** |
| UDP-GlcNAc | C00043 | I3sg2 | C13-06 | 17 | 1998165.949 | 2688456.853 | 2199971.436 | 17.89057 | 2.0285 | 0.1134 | 16.8014 | 20.231 | 16.6393 | 17.8906 | 0.00035 | *** |
| UDP-GlcNAc | C00043 | NT | C13-07 | 17 | 922403.2113 | 991871.6458 | 722534.8907 | 3.884113 | 0.4223 | 0.1087 | 3.95229 | 4.26816 | 3.43189 | 3.88411 | 0.12583 | |
| UDP-GlcNAc | C00043 | I3sg2 | C13-07 | 17 | 269954.1737 | 230061.3818 | 502220.6696 | 2.599879 | 1.0724 | 0.4125 | 2.26988 | 1.73124 | 3.79851 | 2.59988 | 0.12583 | |
| UDP-GlcNAc | C00043 | NT | C13-08 | 17 | 1077562.134 | 1068718.904 | 950924.7636 | 4.577549 | 0.0535 | 0.0117 | 4.61711 | 4.59885 | 4.51669 | 4.57755 | 4.86E-06 | *** |
| UDP-GlcNAc | C00043 | I3sg2 | C13-08 | 17 | 1954525.123 | 2136946.621 | 2286454.698 | 16.6029 | 0.6236 | 0.0376 | 16.4344 | 16.0808 | 17.2934 | 16.6029 | 4.86E-06 | *** |
| UDP-GlcNAc | C00043 | NT | C13-09 | 17 | 253296.8874 | 290605.6502 | 239063.2639 | 1.157112 | 0.0847 | 0.0732 | 1.08532 | 1.25052 | 1.1355 | 1.15711 | 0.10658 | |
| UDP-GlcNAc | C00043 | I3sg2 | C13-09 | 17 | 36536.09985 | 84175.50108 | 143876.2751 | 0.67628 | 0.3923 | 0.58 | 0.30721 | 0.63343 | 1.0882 | 0.67628 | 0.10658 | |
| UDP-GlcNAc | C00043 | NT | C13-10 | 17 | 383896.1753 | 437731.1176 | 394536.3251 | 1.80083 | 0.1351 | 0.075 | 1.64491 | 1.88362 | 1.87396 | 1.80083 | 0.0067 | ** |
| UDP-GlcNAc | C00043 | I3sg2 | C13-10 | 17 | 106980.995 | 90747.10888 | 19215.06923 | 0.575919 | 0.3883 | 0.6743 | 0.89954 | 0.68288 | 0.14533 | 0.57592 | 0.0067 | ** |
| UDP-GlcNAc | C00043 | NT | C13-11 | 17 | 4706521.595 | 4990949.517 | 4546592.485 | 21.07949 | 0.793 | 0.0376 | 20.1664 | 21.4768 | 21.5953 | 21.0795 | 0.00359 | ** |
| UDP-GlcNAc | C00043 | I3sg2 | C13-11 | 17 | 2037395.823 | 2368894.083 | 2402080.477 | 17.70849 | 0.5283 | 0.0298 | 17.1312 | 17.8263 | 18.168 | 17.7085 | 0.00359 | ** |
| UDP-GlcNAc | C00043 | NT | C13-12 | 17 | 1228419.029 | 814766.534 | 912690.2735 | 4.368212 | 0.8792 | 0.2013 | 5.2635 | 3.50606 | 4.33508 | 4.36821 | 0.04536 | * |
| UDP-GlcNAc | C00043 | I3sg2 | C13-12 | 17 | 373898.4772 | 369873.3797 | 340391.8733 | 2.833922 | 0.288 | 0.1016 | 3.14389 | 2.78335 | 2.57453 | 2.83392 | 0.04536 | * |
| UDP-GlcNAc | C00043 | NT | C13-13 | 17 | 6284950.038 | 6375121.793 | 5231027.786 | 26.40297 | 1.3714 | 0.0519 | 26.9296 | 27.433 | 24.8463 | 26.403 | 0.00029 | *** |
| UDP-GlcNAc | C00043 | I3sg2 | C13-13 | 17 | 1806706.73 | 2129070.338 | 1876370.978 | 15.13495 | 0.9162 | 0.0605 | 15.1915 | 16.0215 | 14.1918 | 15.135 | 0.00029 | *** |
| UDP-GlcNAc | C00043 | NT | C13-14 | 17 | 1765901.981 | 1556230.944 | 1771334.966 | 7.558878 | 0.8584 | 0.1136 | 7.56649 | 6.69668 | 8.41346 | 7.55888 | 0.00648 | ** |
| UDP-GlcNAc | C00043 | I3sg2 | C13-14 | 17 | 462580.2985 | 403976.6588 | 630139.9914 | 3.89852 | 0.8631 | 0.2214 | 3.88956 | 3.03998 | 4.76602 | 3.89852 | 0.00648 | ** |
| UDP-GlcNAc | C00043 | NT | C13-15 | 17 | 1749331.437 | 1756831.181 | 1630263.095 | 7.599595 | 0.1286 | 0.0169 | 7.49549 | 7.55989 | 7.7434 | 7.59959 | 0.00025 | *** |
| UDP-GlcNAc | C00043 | I3sg2 | C13-15 | 17 | 475260.3601 | 494611.0837 | 387591.4034 | 3.549906 | 0.5528 | 0.1557 | 3.99618 | 3.72202 | 2.93152 | 3.54991 | 0.00025 | *** |
| UDP-GlcNAc | C00043 | NT | C13-16 | 17 | 1157378.938 | 986878.0367 | 977930.6929 | 4.616915 | 0.357 | 0.0773 | 4.95911 | 4.24668 | 4.64496 | 4.61691 | 0.0019 | ** |
| UDP-GlcNAc | C00043 | I3sg2 | C13-16 | 17 | 298847.8151 | 199154.0196 | 222086.2653 | 1.897076 | 0.5409 | 0.2851 | 2.51283 | 1.49866 | 1.67973 | 1.89708 | 0.0019 | ** |
| UMP | C00105 | NT | C12 PARENT | 9 | 14474.29165 | 22585.54691 | 97100.36439 | 4.628948 | 1.9894 | 0.4298 | 3.0244 | 4.00752 | 6.85493 | 4.62895 | 0.00053 | *** |
| UMP | C00105 | I3sg2 | C12 PARENT | 9 | 377728.293 | 307539.6874 | 404523.863 | 38.2252 | 5.3645 | 0.1403 | 42.274 | 32.1408 | 40.2607 | 38.2252 | 0.00053 | *** |
| UMP | C00105 | NT | C13-05 | 9 | 373781.0928 | 451833.813 | 878486.3592 | 73.43051 | 9.9377 | 0.1353 | 78.1015 | 80.1722 | 62.0179 | 73.4305 | 0.02284 | * |
| UMP | C00105 | I3sg2 | C13-05 | 9 | 378017.4688 | 530130.6631 | 486685.1709 | 48.71601 | 6.5531 | 0.1345 | 42.3064 | 55.4037 | 48.4379 | 48.716 | 0.02284 | * |
| UMP | C00105 | NT | C13-06 | 9 | 19723.46955 | 11007.17172 | 139085.9116 | 5.297749 | 4.0628 | 0.7669 | 4.12122 | 1.95308 | 9.81895 | 5.29775 | 0.30434 | |
| UMP | C00105 | I3sg2 | C13-06 | 9 | 0 | 51951.21603 | 0 | 1.809799 | 3.1347 | 1.7321 | 0 | 5.4294 | 0 | 1.8098 | 0.30434 | |
| UMP | C00105 | NT | C13-07 | 9 | 31808.38895 | 27115.36161 | 193695.2946 | 8.377267 | 4.6781 | 0.5584 | 6.64636 | 4.81128 | 13.6742 | 8.37727 | 0.87922 | |
| UMP | C00105 | I3sg2 | C13-07 | 9 | 89631.64925 | 51442.09872 | 113551.1634 | 8.902923 | 3.1196 | 0.3504 | 10.0313 | 5.37619 | 11.3013 | 8.90292 | 0.87922 | |
| UMP | C00105 | NT | C13-08 | 9 | 38796.54316 | 51037.49852 | 108137.1757 | 8.265523 | 0.7241 | 0.0876 | 8.10653 | 9.05596 | 7.63408 | 8.26552 | 0.02292 | * |
| UMP | C00105 | I3sg2 | C13-08 | 9 | 48145.81727 | 15787.07071 | 0 | 2.34607 | 2.7608 | 1.1768 | 5.38831 | 1.6499 | 0 | 2.34607 | 0.02292 | * |
| UTP | C00075 | NT | C12 PARENT | 9 | 643914.6666 | 381653.9582 | 436727.9556 | 6.136502 | 0.9988 | 0.1628 | 5.33237 | 7.25454 | 5.8226 | 6.1365 | 1.83E-06 | *** |
| UTP | C00075 | I3sg2 | C12 PARENT | 9 | 1613276.303 | 2586940.413 | 2583119.91 | 39.4883 | 0.9206 | 0.0233 | 39.1863 | 38.7566 | 40.522 | 39.4883 | 1.83E-06 | *** |
| UTP | C00075 | NT | C13-01 | 9 | 0 | 0 | 0 | 0 | 0 | NA | 0 | 0 | 0 | 0 | NA | |
| UTP | C00075 | I3sg2 | C13-01 | 9 | 0 | 0 | 0 | 0 | 0 | NA | 0 | 0 | 0 | 0 | NA | |
| UTP | C00075 | NT | C13-02 | 9 | 80809.24955 | 11248.67979 | 11723.9039 | 0.34644 | 0.281 | 0.8111 | 0.6692 | 0.21382 | 0.15631 | 0.34644 | 0.35076 | |
| UTP | C00075 | I3sg2 | C13-02 | 9 | 0 | 0 | 24517.77924 | 0.128205 | 0.2221 | 1.7321 | 0 | 0 | 0.38462 | 0.12821 | 0.35076 | |
| UTP | C00075 | NT | C13-03 | 9 | 49006.6853 | 1803.035636 | 51281.8942 | 0.374604 | 0.3258 | 0.8698 | 0.40583 | 0.03427 | 0.68371 | 0.3746 | 0.77193 | |
| UTP | C00075 | I3sg2 | C13-03 | 9 | 0 | 34170.00093 | 24586.35938 | 0.299205 | 0.2667 | 0.8913 | 0 | 0.51192 | 0.38569 | 0.2992 | 0.77193 | |
| UTP | C00075 | NT | C13-04 | 9 | 127409.6261 | 62645.02915 | 24719.91978 | 0.858481 | 0.463 | 0.5394 | 1.0551 | 1.19077 | 0.32957 | 0.85848 | 0.28108 | |
| UTP | C00075 | I3sg2 | C13-04 | 9 | 36877.23637 | 83566.84025 | 158680.9479 | 1.545658 | 0.8364 | 0.5411 | 0.89574 | 1.25197 | 2.48926 | 1.54566 | 0.28108 | |

| | | | | | | | | | | | | | | | | |
|---|---|---|---|---|---|---|---|---|---|---|---|---|---|---|---|---|
| UTP | C00075 | NT | C13-05 | 9 | 6852619.649 | 2989903.58 | 4550828.829 | 58.08447 | 2.2422 | 0.0386 | 56.7477 | 56.8326 | 60.6731 | 58.0845 | 0.00047 | *** |
| UTP | C00075 | I3sg2 | C13-05 | 9 | 1677815.88 | 2710819.829 | 2359285.9 | 39.45904 | 2.1216 | 0.0538 | 40.7539 | 40.6126 | 37.0106 | 39.459 | 0.00047 | *** |
| UTP | C00075 | NT | C13-06 | 9 | 1177755.966 | 544624.6482 | 724069.1742 | 9.91968 | 0.378 | 0.0381 | 9.7532 | 10.3523 | 9.65352 | 9.91968 | 0.00013 | *** |
| UTP | C00075 | I3sg2 | C13-06 | 9 | 225414.8834 | 397993.9155 | 385774.2355 | 5.829877 | 0.3103 | 0.0532 | 5.4753 | 5.96261 | 6.05172 | 5.82988 | 0.00013 | *** |
| UTP | C00075 | NT | C13-07 | 9 | 1955449.048 | 808024.1969 | 1169166.815 | 15.71339 | 0.4311 | 0.0274 | 16.1934 | 15.3591 | 15.5877 | 15.7134 | 1.57E-05 | *** |
| UTP | C00075 | I3sg2 | C13-07 | 9 | 364979.767 | 570818.5829 | 534107.1696 | 8.598592 | 0.2467 | 0.0287 | 8.86531 | 8.55181 | 8.37865 | 8.59859 | 1.57E-05 | *** |
| UTP | C00075 | NT | C13-08 | 9 | 1188617.809 | 460994.989 | 532050.1869 | 8.566428 | 1.3853 | 0.1617 | 9.84315 | 8.76267 | 7.09346 | 8.56643 | 0.00858 | ** |
| UTP | C00075 | I3sg2 | C13-08 | 9 | 198576.8019 | 290521.5567 | 304545.2068 | 4.651122 | 0.2596 | 0.0558 | 4.82341 | 4.35249 | 4.77747 | 4.65112 | 0.00858 | ** |

| | | | | | | | Abundance | | | | | | | |
|---|---|---|---|---|---|---|---|---|---|---|---|---|---|---|
| **Name** | **KEGG.ID** | **Condition** | **Av** | **Std** | **CV** | **Exp001** | **Exp002** | **Exp003** | **ANOVA** | **Sig** | **Av2** | **denom** | **Norm_Av** | **Norm_Std** |
| ADP | C00008 | NT2 | 29687814.67 | 2747385.312 | 0.09254 | 31571368 | 30956761 | 26535315 | 0.00567 | ** | 29687814.67 | 17474110 | 1.69896 | 0.15723 |
| ADP | C00008 | I3sg5 | 17474110 | 2787412.287 | 0.15952 | 17741290 | 20118312 | 14562728 | 0.00567 | ** | 17474110 | 17474110 | 1 | 0.15952 |
| Ala | C00041 | NT2 | 20530081.67 | 3352270.196 | 0.16329 | 22902725 | 21992465 | 16695055 | 0.02415 | * | 20530081.67 | 12259519.33 | 1.6746237 | 0.27344 |
| Ala | C00041 | I3sg5 | 12259519.33 | 2279303.992 | 0.18592 | 10058225 | 14609548 | 12110785 | 0.02415 | * | 12259519.33 | 12259519.33 | 1 | 0.18592 |
| AMP | C00020 | NT2 | 22652352.33 | 2011770.744 | 0.08881 | 23928430 | 23695367 | 20333260 | 0.03864 | * | 22652352.33 | 13814123.67 | 1.6397966 | 0.14563 |
| AMP | C00020 | I3sg5 | 13814123.67 | 4627504.037 | 0.33498 | 11210192 | 19156937 | 11075242 | 0.03864 | * | 13814123.67 | 13814123.67 | 1 | 0.33498 |
| Asp | C00049 | NT2 | 47838155.67 | 5689769.478 | 0.11894 | 52393660 | 49660283 | 41460524 | 0.00624 | ** | 47838155.67 | 27266563 | 1.7544623 | 0.20867 |
| Asp | C00049 | I3sg5 | 27266563 | 3670691.569 | 0.13462 | 23802892 | 31114105 | 26882692 | 0.00624 | ** | 27266563 | 27266563 | 1 | 0.13462 |
| ATP | C00002 | NT2 | 60887045 | 6077458.213 | 0.09982 | 67466606 | 55483611 | 59710918 | 0.00446 | ** | 60887045 | 32744685.33 | 1.8594482 | 0.1856 |
| ATP | C00002 | I3sg5 | 32744685.33 | 5854785.413 | 0.1788 | 35604530 | 36619900 | 26009626 | 0.00446 | ** | 32744685.33 | 32744685.33 | 1 | 0.1788 |
| Carnitine | C00318 | NT2 | 4201218.333 | 443087.1715 | 0.10547 | 4512842 | 4396824 | 3693989 | 0.02801 | * | 4201218.333 | 2472379.667 | 1.699261 | 0.17921 |
| Carnitine | C00318 | I3sg5 | 2472379.667 | 769774.7345 | 0.31135 | 1795014 | 3309495 | 2312630 | 0.02801 | * | 2472379.667 | 2472379.667 | 1 | 0.31135 |
| CDP | C00112 | NT2 | 2391516.333 | 155556.8143 | 0.06505 | 2570124 | 2285708 | 2318717 | 0.00078 | *** | 2391516.333 | 1405581.667 | 1.7014425 | 0.11067 |
| CDP | C00112 | I3sg5 | 1405581.667 | 101731.1533 | 0.07238 | 1429989 | 1492889 | 1293867 | 0.00078 | *** | 1405581.667 | 1405581.667 | 1 | 0.07238 |
| Cit | C00158 | NT2 | 111904613.3 | 11694657.29 | 0.10451 | 125395932 | 104655715 | 105662193 | 0.01512 | * | 111904613.3 | 77565258 | 1.4427157 | 0.15077 |
| Cit | C00158 | I3sg5 | 77565258 | 8711735.455 | 0.11231 | 77049712 | 86523318 | 69122744 | 0.01512 | * | 77565258 | 77565258 | 1 | 0.11231 |
| CoA | C00010 | NT2 | 297851.3333 | 34540.56057 | 0.11597 | 323707 | 311223 | 258624 | 0.01409 | * | 297851.3333 | 176716 | 1.6854803 | 0.19546 |
| CoA | C00010 | I3sg5 | 176716 | 36674.50148 | 0.20753 | 162794 | 218313 | 149041 | 0.01409 | * | 176716 | 176716 | 1 | 0.20753 |
| CTP | C00063 | NT2 | 2558552.667 | 192417.2111 | 0.07521 | 2780586 | 2454627 | 2440445 | 0.01719 | * | 2558552.667 | 1589672.333 | 1.6094843 | 0.12104 |
| CTP | C00063 | I3sg5 | 1589672.333 | 381919.4769 | 0.24025 | 1791551 | 1828286 | 1149180 | 0.01719 | * | 1589672.333 | 1589672.333 | 1 | 0.24025 |
| Cystathionine | C02291 | NT2 | 1020551 | 221656.1002 | 0.21719 | 1218156 | 1062623 | 780874 | 0.0495 | * | 1020551 | 662197.6667 | 1.5411577 | 0.33473 |
| Cystathionine | C02291 | I3sg5 | 662197.6667 | 22195.64111 | 0.03352 | 636786 | 672017 | 677790 | 0.0495 | * | 662197.6667 | 662197.6667 | 1 | 0.03352 |
| F16BP | C00354 | NT2 | 1171676.333 | 236517.525 | 0.20186 | 1396353 | 1193803 | 924873 | 0.01606 | * | 1171676.333 | 579557 | 2.0216758 | 0.4081 |
| F16BP | C00354 | I3sg5 | 579557 | 98143.33663 | 0.16934 | 552969 | 688255 | 497447 | 0.01606 | * | 579557 | 579557 | 1 | 0.16934 |
| GABA | C00334 | NT2 | 4237106 | 382806.3826 | 0.09035 | 4583614 | 3826176 | 4301528 | 0.00971 | ** | 4237106 | 3001807 | 1.4115185 | 0.12753 |
| GABA | C00334 | I3sg5 | 3001807 | 256515.5015 | 0.08545 | 3023017 | 3247059 | 2735345 | 0.00971 | ** | 3001807 | 3001807 | 1 | 0.08545 |
| GDP | C00035 | NT2 | 3240595.333 | 519604.7114 | 0.16034 | 3795023 | 3161993 | 2764770 | 0.00938 | ** | 3240595.333 | 1654241.333 | 1.9589617 | 0.3141 |
| GDP | C00035 | I3sg5 | 1654241.333 | 270867.1706 | 0.16374 | 1669527 | 1917142 | 1376055 | 0.00938 | ** | 1654241.333 | 1654241.333 | 1 | 0.16374 |
| GlcNAc-6P | C00357 | NT2 | 472874.6667 | 75410.93378 | 0.15947 | 548068 | 473308 | 397248 | 0.01118 | * | 472874.6667 | 264473 | 1.7879884 | 0.28514 |
| GlcNAc-6P | C00357 | I3sg5 | 264473 | 29507.44359 | 0.11157 | 297615 | 254750 | 241054 | 0.01118 | * | 264473 | 264473 | 1 | 0.11157 |
| Gln | C00064 | NT2 | 741289960.7 | 26325017.79 | 0.03551 | 711051114 | 753723183 | 759095585 | 0.02614 | * | 741289960.7 | 525835826.3 | 1.4097365 | 0.05006 |
| Gln | C00064 | I3sg5 | 525835826.3 | 105035551.7 | 0.19975 | 509626354 | 638033823 | 429847302 | 0.02614 | * | 525835826.3 | 525835826.3 | 1 | 0.19975 |
| Gly | C00037 | NT2 | 14088756.33 | 875840.6699 | 0.06217 | 15037086 | 13918894 | 13310289 | 0.00849 | ** | 14088756.33 | 9263137.667 | 1.5209486 | 0.09455 |
| Gly | C00037 | I3sg5 | 9263137.667 | 1494718.604 | 0.16136 | 8226445 | 10976530 | 8586438 | 0.00849 | ** | 9263137.667 | 9263137.667 | 1 | 0.16136 |
| GMP | C00144 | NT2 | 1552161.333 | 29315.47333 | 0.01889 | 1583046 | 1524719 | 1548719 | 0.01161 | * | 1552161.333 | 852451.6667 | 1.8208203 | 0.03439 |
| GMP | C00144 | I3sg5 | 852451.6667 | 273329.8772 | 0.32064 | 671871 | 1166912 | 718572 | 0.01161 | * | 852451.6667 | 852451.6667 | 1 | 0.32064 |
| GSH | C00051 | NT2 | 99566167.33 | 10036631.49 | 0.1008 | 108177556 | 101977389 | 88543557 | 0.00925 | ** | 99566167.33 | 62824450.33 | 1.5848315 | 0.15976 |
| GSH | C00051 | I3sg5 | 62824450.33 | 9054845.555 | 0.14413 | 63796470 | 71354072 | 53322809 | 0.00925 | ** | 62824450.33 | 62824450.33 | 1 | 0.14413 |
| GSSG | C00127 | NT2 | 9952964.667 | 929198.7724 | 0.09336 | 8915971 | 10709985 | 10232938 | 0.00421 | ** | 9952964.667 | 6340444.667 | 1.5697581 | 0.14655 |
| GSSG | C00127 | I3sg5 | 6340444.667 | 522926.7378 | 0.08247 | 6411607 | 6824146 | 5785581 | 0.00421 | ** | 6340444.667 | 6340444.667 | 1 | 0.08247 |
| GTP | C00044 | NT2 | 34529720 | 4765738.285 | 0.13802 | 37109090 | 29030237 | 37449833 | 0.0207 | * | 34529720 | 20568904 | 1.6787341 | 0.2317 |
| GTP | C00044 | I3sg5 | 20568904 | 4452187.554 | 0.21645 | 23160921 | 23117770 | 15428021 | 0.0207 | * | 20568904 | 20568904 | 1 | 0.21645 |
| His | C00135 | NT2 | 37494460.67 | 4703395.137 | 0.12544 | 40099648 | 32064922 | 40318812 | 0.0373 | * | 37494460.67 | 26739034 | 1.4022369 | 0.1759 |
| His | C00135 | I3sg5 | 26739034 | 3834942.05 | 0.14342 | 26911274 | 30484954 | 22820874 | 0.0373 | * | 26739034 | 26739034 | 1 | 0.14342 |
| Lys | C00047 | NT2 | 3195490.667 | 282205.8335 | 0.08831 | 2909693 | 3202817 | 3473962 | 0.01271 | * | 3195490.667 | 2243890.333 | 1.424085 | 0.12577 |
| Lys | C00047 | I3sg5 | 2243890.333 | 260239.9959 | 0.11598 | 2437165 | 2346523 | 1947983 | 0.01271 | * | 2243890.333 | 2243890.333 | 1 | 0.11598 |
| Mal | C00149 | NT2 | 66252472.67 | 5661976.686 | 0.08546 | 71564175 | 66897734 | 60295509 | 0.00575 | ** | 66252472.67 | 43583657.67 | 1.5201219 | 0.12991 |
| Mal | C00149 | I3sg5 | 43583657.67 | 4595969.134 | 0.10545 | 45185891 | 47164044 | 38401038 | 0.00575 | ** | 43583657.67 | 43583657.67 | 1 | 0.10545 |
| NAM | C00153 | NT2 | 147956365.3 | 16967788.32 | 0.11468 | 155800704 | 159582886 | 128485704 | 0.04767 | * | 147956365.3 | 112385117.3 | 1.3165121 | 0.15098 |
| NAM | C00153 | I3sg5 | 112385117.3 | 13721903.75 | 0.1221 | 97705256 | 114561048 | 124889048 | 0.04767 | * | 112385117.3 | 112385117.3 | 1 | 0.1221 |
| Ornithine | C00077 | NT2 | 246775 | 11052.75653 | 0.04479 | 259194 | 238018 | 243113 | 0.00792 | ** | 246775 | 180932 | 1.3639102 | 0.06109 |
| Ornithine | C00077 | I3sg5 | 180932 | 20361.3433 | 0.11254 | 203971 | 173473 | 165352 | 0.00792 | ** | 180932 | 180932 | 1 | 0.11254 |
| Orotidine | X00006 | NT2 | 4002345.333 | 268779.8036 | 0.06716 | 4311679 | 3869521 | 3825836 | 0.00513 | ** | 4002345.333 | 2496873.667 | 1.6029427 | 0.10765 |
| Orotidine | X00006 | I3sg5 | 2496873.667 | 384406.1457 | 0.15395 | 2467613 | 2895074 | 2127934 | 0.00513 | ** | 2496873.667 | 2496873.667 | 1 | 0.15395 |
| P-Choline | C00588 | NT2 | 79721416 | 13161855.23 | 0.1651 | 93982720 | 77140032 | 68041496 | 0.0139 | * | 79721416 | 44373289.33 | 1.7966082 | 0.29662 |
| P-Choline | C00588 | I3sg5 | 44373289.33 | 6410635.471 | 0.14447 | 41541244 | 51712224 | 39866400 | 0.0139 | * | 44373289.33 | 44373289.33 | 1 | 0.14447 |
| Palmitate | C00249 | NT2 | 34748899.33 | 8587824.169 | 0.24714 | 33886178 | 26624998 | 43735522 | 0.01769 | * | 34748899.33 | 11021330.67 | 3.1528769 | 0.7792 |
| Palmitate | C00249 | I3sg5 | 11021330.67 | 6151909.142 | 0.55818 | 10670246 | 17341264 | 5052482 | 0.01769 | * | 11021330.67 | 11021330.67 | 1 | 0.55818 |
| PPi | C00013 | NT2 | 261988.3333 | 33859.29033 | 0.12924 | 236943 | 300511 | 248511 | 0.01131 | * | 261988.3333 | 115839.3333 | 2.2616526 | 0.2923 |
| PPi | C00013 | I3sg5 | 115839.3333 | 45817.58729 | 0.39553 | 128692 | 153858 | 64968 | 0.01131 | * | 115839.3333 | 115839.3333 | 1 | 0.39553 |
| Pro | C00148 | NT2 | 299319356.7 | 58560030.67 | 0.19564 | 358925707 | 297167392 | 241864971 | 0.01936 | * | 299319356.7 | 164119571.3 | 1.8237883 | 0.35681 |
| Pro | C00148 | I3sg5 | 164119571.3 | 19996639.91 | 0.12184 | 144965264 | 184863640 | 162529810 | 0.01936 | * | 164119571.3 | 164119571.3 | 1 | 0.12184 |
| PRPP | C00119 | NT2 | 652263.3333 | 103628.5494 | 0.15888 | 708821 | 532662 | 715307 | 0.00664 | ** | 652263.3333 | 260317 | 2.5056502 | 0.39809 |
| PRPP | C00119 | I3sg5 | 260317 | 80568.87501 | 0.3095 | 312542 | 167528 | 300881 | 0.00664 | ** | 260317 | 260317 | 1 | 0.3095 |
| SAM | C00019 | NT2 | 8324709.667 | 704831.8896 | 0.08467 | 8302210 | 7631397 | 9040522 | 0.00401 | ** | 8324709.667 | 4564173 | 1.8239251 | 0.15443 |
| SAM | C00019 | I3sg5 | 4564173 | 838630.3057 | 0.18374 | 5062640 | 5033931 | 3595948 | 0.00401 | ** | 4564173 | 4564173 | 1 | 0.18374 |
| Ser | C00065 | NT2 | 52075303 | 4317479.693 | 0.08291 | 50441421 | 56971270 | 48813218 | 0.01363 | * | 52075303 | 34199572 | 1.5226887 | 0.12624 |
| Ser | C00065 | I3sg5 | 34199572 | 5962888.109 | 0.17436 | 32982528 | 40677092 | 28939096 | 0.01363 | * | 34199572 | 34199572 | 1 | 0.17436 |
| Succinyl-CoA | C00091 | NT2 | 20263 | 3522.649429 | 0.17385 | 22773 | 16236 | 21780 | 0.01253 | * | 20263 | 8524 | 2.3771703 | 0.41326 |
| Succinyl-CoA | C00091 | I3sg5 | 8524 | 3134.583864 | 0.36774 | 11484 | 8848 | 5240 | 0.01253 | * | 8524 | 8524 | 1 | 0.36774 |
| Thr | C00188 | NT2 | 111169640 | 7252335.308 | 0.06524 | 118295320 | 111416640 | 103796960 | 0.01068 | * | 111169640 | 76898821.33 | 1.4456612 | 0.09431 |
| Thr | C00188 | I3sg5 | 76898821.33 | 10958126.2 | 0.1425 | 68810776 | 89370072 | 72515616 | 0.01068 | * | 76898821.33 | 76898821.33 | 1 | 0.1425 |
| Trp | C00078 | NT2 | 28660116.67 | 1742795.111 | 0.06081 | 30671166 | 27718586 | 27590598 | 0.01156 | * | 28660116.67 | 20010652.67 | 1.432243 | 0.08709 |
| Trp | C00078 | I3sg5 | 20010652.67 | 2911670.378 | 0.14551 | 17290858 | 23082234 | 19658866 | 0.01156 | * | 20010652.67 | 20010652.67 | 1 | 0.14551 |
| UDP | C00015 | NT2 | 7702028.333 | 1197800.459 | 0.15552 | 8644629 | 8107286 | 6354170 | 0.01547 | * | 7702028.333 | 4542612.333 | 1.6955064 | 0.26368 |
| UDP | C00015 | I3sg5 | 4542612.333 | 624977.3193 | 0.13758 | 5004133 | 4792316 | 3831388 | 0.01547 | * | 4542612.333 | 4542612.333 | 1 | 0.13758 |
| UDP-GlcNAc | C00043 | NT2 | 22200515 | 1215510.967 | 0.05475 | 20830578 | 23149910 | 22621057 | 0.00547 | ** | 22200515 | 15244714.33 | 1.4562762 | 0.07973 |
| UDP-GlcNAc | C00043 | I3sg5 | 15244714.33 | 1841608.796 | 0.1208 | 16957058 | 15480528 | 13296557 | 0.00547 | ** | 15244714.33 | 15244714.33 | 1 | 0.1208 |
| UMP | C00105 | NT2 | 1414392.667 | 350945.1089 | 0.24812 | 1297799 | 1808795 | 1136584 | 0.04772 | * | 1414392.667 | 796514 | 1.7757286 | 0.4406 |
| UMP | C00105 | I3sg5 | 796514 | 143612.1622 | 0.1803 | 745780 | 958607 | 685155 | 0.04772 | * | 796514 | 796514 | 1 | 0.1803 |
| UTP | C00075 | NT2 | 7838009.333 | 194050.1177 | 0.02476 | 8057887 | 7690710 | 7765431 | 0.00756 | ** | 7838009.333 | 4658482.667 | 1.6825241 | 0.04166 |
| UTP | C00075 | I3sg5 | 4658482.667 | 1087125.453 | 0.23336 | 5528324 | 5007382 | 3439742 | 0.00756 | ** | 4658482.667 | 4658482.667 | 1 | 0.23336 |
| Xanthine | C00385 | NT2 | 59006.33333 | 18505.75879 | 0.31362 | 77823 | 58368 | 40828 | 0.04865 | * | 59006.33333 | 29040.66667 | 2.0318519 | 0.63724 |
| Xanthine | C00385 | I3sg5 | 29040.66667 | 577.4221448 | 0.01988 | 28483 | 29636 | 29003 | 0.04865 | * | 29040.66667 | 29040.66667 | 1 | 0.01988 |

**Isotopologue Distribution**

| Name | KEGG.ID | Condition | Iso | Nr.C | Exp001 | Exp002 | Exp003 | Norm_Av | Norm_Std | CV | MID001 | MID002 | MID003 | Av | ANOVA | Sig |
|---|---|---|---|---|---|---|---|---|---|---|---|---|---|---|---|---|
| 2-HG | C02630 | NT2 | C12 PARENT | 5 | 2343834.88 | 2154520.372 | 2348404.775 | 85.648782 | 1.7802 | 0.0208 | 83.6085 | 86.452 | 86.8859 | 85.6488 | 0.00651 | ** |
| 2-HG | C02630 | I3sg5 | C12 PARENT | 5 | 2131700.76 | 2396061.267 | 1951072.9 | 91.219231 | 0.522 | 0.0057 | 91.2428 | 90.6859 | 91.7291 | 91.2192 | 0.00651 | ** |
| 2-HG | C02630 | NT2 | C13-02 | 5 | 246545.8235 | 210178.7515 | 228738.678 | 8.5637122 | 0.2006 | 0.0234 | 8.7947 | 8.4336 | 8.46284 | 8.56371 | 0.00626 | ** |
| 2-HG | C02630 | I3sg5 | C13-02 | 5 | 145910.104 | 185790.0729 | 118901.8488 | 6.2890852 | 0.7218 | 0.1148 | 6.24536 | 7.03176 | 5.59013 | 6.28909 | 0.00626 | ** |
| 2-HG | C02630 | NT2 | C13-03 | 5 | 30643.05482 | 60620.1114 | 61010.97511 | 1.9275989 | 0.728 | 0.3777 | 1.09309 | 2.43243 | 2.25727 | 1.9276 | 0.03301 | * |
| 2-HG | C02630 | I3sg5 | C13-03 | 5 | 0 | 5675.687175 | 18100.30268 | 0.3552642 | 0.4425 | 1.2456 | 0 | 0.21481 | 0.85098 | 0.35526 | 0.03301 | * |
| 2-HG | C02630 | NT2 | C13-04 | 5 | 98808.67433 | 66840.12783 | 13219.45528 | 2.2319256 | 1.567 | 0.7021 | 3.52467 | 2.68202 | 0.48909 | 2.23193 | 0.92286 | |
| 2-HG | C02630 | I3sg5 | C13-04 | 5 | 58684.48285 | 54628.50871 | 38920.31759 | 2.1364201 | 0.3462 | 0.162 | 2.51186 | 2.06757 | 1.82983 | 2.13642 | 0.92286 | |
| 2-HG | C02630 | NT2 | C13-05 | 5 | 83513.90385 | 0 | 51485.81653 | 1.6279812 | 1.5087 | 0.9267 | 2.97908 | 0 | 1.90486 | 1.62798 | 0.13499 | |
| 2-HG | C02630 | I3sg5 | C13-05 | 5 | 0 | 0 | 0 | 0 | 0 | NA | 0 | 0 | 0 | 0 | 0.13499 | |
| 3PG | C00197 | NT2 | C12 PARENT | 3 | 52195.45967 | 115314.6225 | 161052.4924 | 6.5299052 | 2.8654 | 0.4388 | 3.4464 | 7.03257 | 9.11074 | 6.52991 | 0.65569 | |
| 3PG | C00197 | I3sg5 | C12 PARENT | 3 | 141904.4427 | 89467.30189 | 93277.41158 | 7.768724 | 3.4197 | 0.4402 | 11.2746 | 4.44218 | 7.58937 | 7.76872 | 0.65569 | |
| 3PG | C00197 | NT2 | C13-03 | 3 | 1462296.792 | 1524406.818 | 1606668.477 | 93.470095 | 2.8654 | 0.0307 | 96.5536 | 92.9674 | 90.8893 | 93.4701 | 0.65569 | |
| 3PG | C00197 | I3sg5 | C13-03 | 3 | 1116713.846 | 1924570.905 | 1135776.669 | 92.231276 | 3.4197 | 0.0371 | 88.7254 | 95.5578 | 92.4106 | 92.2313 | 0.65569 | |
| 5-Oxoproline | C01879 | NT2 | C12 PARENT | 5 | 15914934.95 | 14748059.43 | 12696871.18 | 92.456833 | 1.7122 | 0.0185 | 92.8362 | 90.5868 | 93.9475 | 92.4568 | 0.22821 | |
| 5-Oxoproline | C01879 | I3sg5 | C12 PARENT | 5 | 10927359.66 | 14375548.32 | 10835106.52 | 93.986719 | 0.737 | 0.0078 | 94.5221 | 93.1461 | 94.292 | 93.9867 | 0.22821 | |
| 5-Oxoproline | C01879 | NT2 | C13-01 | 5 | 1963.284442 | 199562.2387 | 75758.99429 | 0.5992604 | 0.6081 | 1.0147 | 0.01145 | 1.22577 | 0.56056 | 0.59926 | 0.5125 | |
| 5-Oxoproline | C01879 | I3sg5 | C13-01 | 5 | 0 | 128636.817 | 0 | 0.2778333 | 0.4812 | 1.7321 | 0 | 0.8335 | 0 | 0.27783 | 0.5125 | |
| 5-Oxoproline | C01879 | NT2 | C13-02 | 5 | 778180.9319 | 841223.1433 | 588887.8272 | 4.6879044 | 0.4248 | 0.0906 | 4.53935 | 5.16703 | 4.35734 | 4.6879 | 0.05309 | |
| 5-Oxoproline | C01879 | I3sg5 | C13-02 | 5 | 405928.6941 | 584701.121 | 481857.965 | 3.831068 | 0.343 | 0.0895 | 3.5113 | 3.78856 | 4.19335 | 3.83107 | 0.05309 | |
| 5-Oxoproline | C01879 | NT2 | C13-04 | 5 | 354829.9308 | 344918.635 | 112244.3178 | 1.6729784 | 0.73 | 0.4363 | 2.06982 | 2.11859 | 0.83053 | 1.67298 | 0.5511 | |
| 5-Oxoproline | C01879 | I3sg5 | C13-04 | 5 | 134356.2896 | 256446.5932 | 152207.0489 | 1.3828008 | 0.2548 | 0.1842 | 1.16219 | 1.66164 | 1.32457 | 1.3828 | 0.5511 | |
| 5-Oxoproline | C01879 | NT2 | C13-05 | 5 | 93113.09291 | 146822.5298 | 41097.37218 | 0.5830235 | 0.3009 | 0.516 | 0.54315 | 0.90183 | 0.30409 | 0.58302 | 0.81755 | |
| 5-Oxoproline | C01879 | I3sg5 | C13-05 | 5 | 93000.49731 | 87999.33176 | 21843.14466 | 0.5215789 | 0.3101 | 0.5945 | 0.80446 | 0.57019 | 0.19009 | 0.52158 | 0.81755 | |
| A | C00212 | NT2 | C12 PARENT | 10 | 232912.7924 | 189368.8492 | 201244.6027 | 29.642607 | 3.8527 | 0.13 | 29.6257 | 25.7984 | 33.5037 | 29.6426 | 0.35127 | |
| A | C00212 | I3sg5 | C12 PARENT | 10 | 114440.5201 | 184611.2438 | 168734.2359 | 33.53696 | 5.1087 | 0.1523 | 39.4088 | 30.1114 | 31.0907 | 33.537 | 0.35127 | |
| A | C00212 | NT2 | C13-04 | 10 | 16096.05285 | 23773.19731 | 0 | 1.7620246 | 1.6381 | 0.9297 | 2.04737 | 3.23871 | 0 | 1.76202 | 0.39618 | |
| A | C00212 | I3sg5 | C13-04 | 10 | 0 | 3415.306344 | 9155.589706 | 0.7480179 | 0.8596 | 1.1491 | 0 | 0.55706 | 1.68699 | 0.74802 | 0.39618 | |
| A | C00212 | NT2 | C13-05 | 10 | 485925.7493 | 480830.2896 | 374533.6731 | 63.222219 | 1.9959 | 0.0316 | 61.8082 | 65.5052 | 62.3532 | 63.2222 | 0.79598 | |
| A | C00212 | I3sg5 | C13-05 | 10 | 175952.7941 | 405067.5534 | 350974.2282 | 63.77683 | 2.8462 | 0.0446 | 60.5912 | 66.0694 | 64.6699 | 63.7768 | 0.79598 | |
| A | C00212 | NT2 | C13-06 | 10 | 51249.07693 | 40060.96922 | 24886.01998 | 5.3731495 | 1.1901 | 0.2215 | 6.51872 | 5.45765 | 4.14308 | 5.37315 | 0.04642 | * |
| A | C00212 | I3sg5 | C13-06 | 10 | 0 | 20000.02236 | 13852.46547 | 1.9381922 | 1.7156 | 0.8852 | 0 | 3.26215 | 2.55243 | 1.93819 | 0.04642 | * |
| Ac-carnitine | C02571 | NT2 | C12 PARENT | 9 | 9198687.045 | 8603879.979 | 5904976.485 | 65.348949 | 1.3952 | 0.0214 | 65.8984 | 66.3858 | 63.7626 | 65.3489 | 0.02917 | * |
| Ac-carnitine | C02571 | I3sg5 | C12 PARENT | 9 | 3959574.588 | 7078004.506 | 4438011.666 | 60.64518 | 2.012 | 0.0332 | 59.8135 | 59.1824 | 62.9397 | 60.6452 | 0.02917 | * |
| Ac-carnitine | C02571 | NT2 | C13-01 | 9 | 0 | 0 | 0 | 0 | 0 | NA | 0 | 0 | 0 | 0 | NA | |
| Ac-carnitine | C02571 | I3sg5 | C13-01 | 9 | 0 | 0 | 0 | 0 | 0 | NA | 0 | 0 | 0 | 0 | NA | |
| Ac-carnitine | C02571 | NT2 | C13-02 | 9 | 4760201.828 | 4356545.756 | 3355894.728 | 34.651051 | 1.3952 | 0.0403 | 34.1016 | 33.6142 | 36.2374 | 34.6511 | 0.02917 | * |
| Ac-carnitine | C02571 | I3sg5 | C13-02 | 9 | 2660297.215 | 4881635.853 | 2613204.976 | 39.35482 | 2.012 | 0.0511 | 40.1865 | 40.8176 | 37.0603 | 39.3548 | 0.02917 | * |
| Ac-carnitine | C02571 | NT2 | C13-03 | 9 | 0 | 0 | 0 | 0 | 0 | NA | 0 | 0 | 0 | 0 | NA | |
| Ac-carnitine | C02571 | I3sg5 | C13-03 | 9 | 0 | 0 | 0 | 0 | 0 | NA | 0 | 0 | 0 | 0 | NA | |
| Acetyl-CoA | C00024 | NT2 | C12 PARENT | 23 | 6946.123343 | 7112.645581 | 17920.71294 | 11.925446 | 1.5227 | 0.1277 | 10.7879 | 11.3331 | 13.6553 | 11.9254 | 0.20886 | |
| Acetyl-CoA | C00024 | I3sg5 | C12 PARENT | 23 | 16697.64761 | 5849.868831 | 8432.881503 | 13.577532 | 1.1565 | 0.0852 | 14.7498 | 12.4375 | 13.5453 | 13.5775 | 0.20886 | |
| Acetyl-CoA | C00024 | NT2 | C13-01 | 23 | 0 | 54.76352522 | 439.4611594 | 0.1407073 | 0.1737 | 1.2346 | 0 | 0.08726 | 0.33486 | 0.14071 | 0.23328 | |
| Acetyl-CoA | C00024 | I3sg5 | C13-01 | 23 | 0 | 0 | 0 | 0 | 0 | NA | 0 | 0 | 0 | 0 | 0.23328 | |
| Acetyl-CoA | C00024 | NT2 | C13-02 | 23 | 8629.146431 | 9883.661704 | 17800.59272 | 14.23798 | 1.3105 | 0.092 | 13.4018 | 15.7483 | 13.5638 | 14.238 | 0.97327 | |
| Acetyl-CoA | C00024 | I3sg5 | C13-02 | 23 | 15753.16567 | 7662.370296 | 7695.463933 | 14.189128 | 1.9793 | 0.1395 | 13.9155 | 16.291 | 12.3608 | 14.1891 | 0.97327 | |
| Acetyl-CoA | C00024 | NT2 | C13-03 | 23 | 0 | 0 | 263.7339446 | 0.0669872 | 0.116 | 1.7321 | 0 | 0 | 0.20096 | 0.06699 | 0.18617 | |
| Acetyl-CoA | C00024 | I3sg5 | C13-03 | 23 | 1316.114503 | 0 | 425.1421716 | 0.615157 | 0.5842 | 0.9497 | 1.16259 | 0 | 0.68288 | 0.61516 | 0.18617 | |
| Acetyl-CoA | C00024 | NT2 | C13-04 | 23 | 0 | 958.1863354 | 2063.7996 | 1.0331102 | 0.895 | 0.8663 | 0 | 1.52674 | 1.57259 | 1.03311 | 0.48127 | |
| Acetyl-CoA | C00024 | I3sg5 | C13-04 | 23 | 134.721198 | 509.1916119 | 330.6772493 | 0.5775841 | 0.4835 | 0.8371 | 0.11901 | 1.0826 | 0.53115 | 0.57758 | 0.48127 | |
| Acetyl-CoA | C00024 | NT2 | C13-05 | 23 | 19253.05449 | 17703.0764 | 44446.76354 | 30.659007 | 2.9052 | 0.0948 | 29.9017 | 28.2075 | 33.8678 | 30.659 | 0.40345 | |
| Acetyl-CoA | C00024 | I3sg5 | C13-05 | 23 | 36464.90214 | 15392.00134 | 19795.99474 | 32.244518 | 0.4648 | 0.0144 | 32.2112 | 32.7251 | 31.7973 | 32.2445 | 0.40345 | |
| Acetyl-CoA | C00024 | NT2 | C13-06 | 23 | 3340.616294 | 3353.675707 | 6023.554301 | 5.0405918 | 0.398 | 0.079 | 5.18827 | 5.34364 | 4.58987 | 5.04059 | 0.16452 | |
| Acetyl-CoA | C00024 | I3sg5 | C13-06 | 23 | 3008.036417 | 2423.912361 | 2076.922156 | 3.7155657 | 1.2907 | 0.3474 | 2.65714 | 5.1535 | 3.33606 | 3.71557 | 0.16452 | |
| Acetyl-CoA | C00024 | NT2 | C13-07 | 23 | 24410.22699 | 23305.03177 | 41630.65451 | 35.588901 | 3.3713 | 0.0947 | 37.9112 | 37.1335 | 31.722 | 35.5889 | 0.55862 | |
| Acetyl-CoA | C00024 | I3sg5 | C13-07 | 23 | 39093.528 | 14957.51397 | 22427.39442 | 34.119488 | 2.1415 | 0.0628 | 34.5332 | 31.8013 | 36.024 | 34.1195 | 0.55862 | |
| Acetyl-CoA | C00024 | NT2 | C13-08 | 23 | 1808.700174 | 373.2560965 | 0 | 1.1346014 | 1.4803 | 1.3047 | 2.80907 | 0.59473 | 0 | 1.1346 | 0.86198 | |
| Acetyl-CoA | C00024 | I3sg5 | C13-08 | 23 | 737.6249013 | 239.443179 | 1072.323154 | 0.9610268 | 0.6632 | 0.6901 | 0.65158 | 0.50908 | 1.72242 | 0.96103 | 0.86198 | |
| Acetyl-CoA | C00024 | NT2 | C13-09 | 23 | 0 | 15.84204293 | 646.6801574 | 0.1726679 | 0.2775 | 1.6071 | 0 | 0.02524 | 0.49276 | 0.17267 | 0.34181 | |
| Acetyl-CoA | C00024 | I3sg5 | C13-09 | 23 | 0 | 0 | 0 | 0 | 0 | NA | 0 | 0 | 0 | 0 | 0.34181 | |
| Aconitate | C00417 | NT2 | C12 PARENT | 6 | 179971.3443 | 113908.5318 | 231782.0085 | 40.806264 | 9.065 | 0.2221 | 38.9875 | 32.7885 | 50.6428 | 40.8063 | 0.95967 | |
| Aconitate | C00417 | I3sg5 | C12 PARENT | 6 | 122337.0465 | 139782.6742 | 193284.409 | 40.504403 | 3.5004 | 0.0864 | 37.0345 | 40.4442 | 44.0346 | 40.5044 | 0.95967 | |
| Aconitate | C00417 | NT2 | C13-02 | 6 | 175632.6465 | 173100.7404 | 129225.4574 | 38.703144 | 10.8109 | 0.2793 | 38.0476 | 49.8269 | 28.2349 | 38.7031 | 0.19088 | |
| Aconitate | C00417 | I3sg5 | C13-02 | 6 | 114106.4925 | 62342.18609 | 116221.2477 | 26.352853 | 8.2532 | 0.3132 | 34.5429 | 18.0378 | 26.4778 | 26.3529 | 0.19088 | |
| Aconitate | C00417 | NT2 | C13-03 | 6 | 0 | 1672.63098 | 34250.97121 | 2.6550215 | 4.1886 | 1.5776 | 0 | 0.48147 | 7.4836 | 2.65502 | 0.31934 | |
| Aconitate | C00417 | I3sg5 | C13-03 | 6 | 0 | 62464.38061 | 41697.7121 | 9.1909588 | 9.0405 | 0.9836 | 0 | 18.0732 | 9.49969 | 9.19096 | 0.31934 | |
| Aconitate | C00417 | NT2 | C13-04 | 6 | 59849.9481 | 43961.60177 | 35599.46603 | 11.132654 | 2.9092 | 0.2613 | 12.9654 | 12.6543 | 7.77824 | 11.1327 | 0.02467 | * |
| Aconitate | C00417 | I3sg5 | C13-04 | 6 | 85068.52462 | 68500.25085 | 79249.90932 | 21.20896 | 4.0324 | 0.1901 | 25.7524 | 19.8196 | 18.0549 | 21.209 | 0.02467 | * |
| Aconitate | C00417 | NT2 | C13-05 | 6 | 46158.40975 | 14760.55538 | 26822.5813 | 6.7029167 | 2.9664 | 0.4426 | 9.99939 | 4.24881 | 5.86055 | 6.70292 | 0.0903 | |
| Aconitate | C00417 | I3sg5 | C13-05 | 6 | 8820.780715 | 12529.51767 | 8484.501937 | 2.7428242 | 0.8485 | 0.3093 | 2.67027 | 3.62524 | 1.93296 | 2.74282 | 0.0903 | |
| Adenine | C00147 | NT2 | C12 PARENT | 5 | 626329.3085 | 412474.2127 | 396499.9778 | 100 | 0 | 0 | 100 | 100 | 100 | 100 | 0.57733 | |
| Adenine | C00147 | I3sg5 | C12 PARENT | 5 | 190094.9679 | 484823.6149 | 236892.5897 | 100 | 0 | 0 | 100 | 100 | 100 | 100 | 0.57733 | |
| Adenine | C00147 | NT2 | C13-01 | 5 | 0 | 0 | 0 | 0 | 0 | NA | 0 | 0 | 0 | 0 | NA | |
| Adenine | C00147 | I3sg5 | C13-01 | 5 | 0 | 0 | 0 | 0 | 0 | NA | 0 | 0 | 0 | 0 | NA | |
| ADP | C00008 | NT2 | C12 PARENT | 10 | 5961141.676 | 6068920.412 | 5431949.632 | 19.584657 | 0.8213 | 0.0419 | 18.811 | 19.4965 | 20.4465 | 19.5847 | 0.0196 | * |
| ADP | C00008 | I3sg5 | C12 PARENT | 10 | 3843903.167 | 4381971.544 | 3100020.61 | 21.460666 | 0.2612 | 0.0122 | 21.5742 | 21.6459 | 21.1619 | 21.4607 | 0.0196 | * |
| ADP | C00008 | NT2 | C13-01 | 10 | 0 | 0 | 0 | 0 | 0 | NA | 0 | 0 | 0 | 0 | NA | |
| ADP | C00008 | I3sg5 | C13-01 | 10 | 0 | 0 | 0 | 0 | 0 | NA | 0 | 0 | 0 | 0 | NA | |
| ADP | C00008 | NT2 | C13-02 | 10 | 243896.6853 | 222555.9664 | 223387.6615 | 0.775154 | 0.0631 | 0.0814 | 0.76964 | 0.71497 | 0.84086 | 0.77515 | 0.59888 | |
| ADP | C00008 | I3sg5 | C13-02 | 10 | 92656.64765 | 173901.9774 | 111749.617 | 0.7139738 | 0.1747 | 0.2447 | 0.52004 | 0.85903 | 0.76284 | 0.71397 | 0.59888 | |
| ADP | C00008 | NT2 | C13-03 | 10 | 257605.151 | 370340.5461 | 271391.6271 | 1.0080579 | 0.1888 | 0.1873 | 0.8129 | 1.18973 | 1.02155 | 1.00806 | 0.18688 | |
| ADP | C00008 | I3sg5 | C13-03 | 10 | 229599.6458 | 274009.5061 | 154805.4625 | 1.2329824 | 0.156 | 0.1265 | 1.28865 | 1.35354 | 1.05676 | 1.23298 | 0.18688 | |
| ADP | C00008 | NT2 | C13-04 | 10 | 481722.3284 | 473797.9252 | 405050.3815 | 1.5222877 | 0.0023 | 0.0015 | 1.52012 | 1.52209 | 1.52466 | 1.52229 | 0.46903 | |
| ADP | C00008 | I3sg5 | C13-04 | 10 | 100323.6341 | 210953.2666 | 288736.0741 | 1.1920509 | 0.7159 | 0.6005 | 0.56307 | 1.04206 | 1.97102 | 1.19205 | 0.46903 | |
| ADP | C00008 | NT2 | C13-05 | 10 | 22129955.64 | 21630952.13 | 18523932.06 | 69.683098 | 0.1757 | 0.0025 | 69.8333 | 69.4899 | 69.7262 | 69.6831 | 0.44996 | |
| ADP | C00008 | I3sg5 | C13-05 | 10 | 12802914.46 | 14151563.27 | 10146163.65 | 70.341401 | 1.3518 | 0.0192 | 71.8574 | 69.9053 | 69.2615 | 70.3414 | 0.44996 | |
| ADP | C00008 | NT2 | C13-06 | 10 | 1878580.553 | 1782959.434 | 1357005.972 | 5.5879209 | 0.4276 | 0.0765 | 5.92805 | 5.72779 | 5.10792 | 5.58792 | 0.02403 | * |
| ADP | C00008 | I3sg5 | C13-06 | 10 | 605168.1618 | 723886.7108 | 689557.2083 | 3.8931853 | 0.7106 | 0.1825 | 3.39656 | 3.57582 | 4.70717 | 3.89319 | 0.02403 | * |
| ADP | C00008 | NT2 | C13-07 | 10 | 655604.0149 | 578682.9284 | 350057.9545 | 1.7485038 | 0.3876 | 0.2217 | 2.06882 | 1.85903 | 1.31766 | 1.7485 | 0.15053 | |
| ADP | C00008 | I3sg5 | C13-07 | 10 | 142543.233 | 327622.4 | 158035.5643 | 1.1657402 | 0.416 | 0.3569 | 0.80004 | 1.61838 | 1.07881 | 1.16574 | 0.15053 | |
| ADP | C00008 | NT2 | C13-08 | 10 | 81200.97252 | 0 | 3911.878848 | 0.0903208 | 0.1439 | 1.593 | 0.25624 | 0 | 0.01472 | 0.09032 | 0.33803 | |
| ADP | C00008 | I3sg5 | C13-08 | 10 | 0 | 0 | 0 | 0 | 0 | NA | 0 | 0 | 0 | 0 | 0.33803 | |
| Ala | C00041 | NT2 | C12 PARENT | 3 | 2004462.27 | 1910419.311 | 1396335.319 | 8.5180426 | 0.2126 | 0.025 | 8.67177 | 8.60694 | 8.27542 | 8.51804 | 0.13401 | |
| Ala | C00041 | I3sg5 | C12 PARENT | 3 | 918449.2615 | 1269435.138 | 1141095.657 | 8.9759088 | 0.3656 | 0.0407 | 9.00636 | 8.59607 | 9.32529 | 8.97591 | 0.13401 | |
| Ala | C00041 | NT2 | C13-01 | 3 | 15697.05102 | 2227.36587 | 0 | 0.0259813 | 0.0367 | 1.4108 | 0.06791 | 0.01003 | 0 | 0.02598 | 0.28688 | |
| Ala | C00041 | I3sg5 | C13-01 | 3 | 0 | 0 | 0 | 0 | 0 | NA | 0 | 0 | 0 | 0 | 0.28688 | |
| Ala | C00041 | NT2 | C13-02 | 3 | 398048.113 | 217643.9287 | 239732.7575 | 1.374458 | 0.3729 | 0.2713 | 1.72205 | 0.98054 | 1.42078 | 1.37446 | 0.13668 | |

| | | | | | | | | | | | | | | | | |
|---|---|---|---|---|---|---|---|---|---|---|---|---|---|---|---|---|
| Ala | C00041 | I3sg5 | C13-02 | 3 | 0 | 134106.3193 | 126819.7039 | 0.64817 | 0.565 | 0.8717 | 0 | 0.90811 | 1.0364 | 0.64817 | 0.13668 | |
| Ala | C00041 | NT2 | C13-03 | 3 | 20696604.4 | 20065972.48 | 15237215.85 | 90.081518 | 0.473 | 0.0053 | 89.5383 | 90.4025 | 90.3038 | 90.0815 | 0.5735 | |
| Ala | C00041 | I3sg5 | C13-03 | 3 | 9279333.415 | 13364082.19 | 10968648.36 | 90.375921 | 0.6856 | 0.0076 | 90.9936 | 90.4958 | 89.6383 | 90.3759 | 0.5735 | |
| AMP | C00020 | NT2 | C12 PARENT | 10 | 4736954.325 | 4865386.693 | 4310876.32 | 20.389953 | 0.6757 | 0.0331 | 19.6958 | 20.4284 | 21.0456 | 20.39 | 0.01751 | * |
| AMP | C00020 | I3sg5 | C12 PARENT | 10 | 2539661.105 | 4218472.381 | 2610611.297 | 22.65773 | 0.7462 | 0.0329 | 22.5812 | 21.9527 | 23.4392 | 22.6577 | 0.01751 | * |
| AMP | C00020 | NT2 | C13-01 | 10 | 0 | 0 | 0 | 0 | 0 | NA | 0 | 0 | 0 | 0 | NA | |
| AMP | C00020 | I3sg5 | C13-01 | 10 | 0 | 0 | 0 | 0 | 0 | NA | 0 | 0 | 0 | 0 | NA | |
| AMP | C00020 | NT2 | C13-02 | 10 | 159981.6853 | 197750.5256 | 154038.064 | 0.7491668 | 0.0826 | 0.1102 | 0.66519 | 0.8303 | 0.75201 | 0.74917 | 0.0756 | |
| AMP | C00020 | I3sg5 | C13-02 | 10 | 108765.5164 | 157866.3024 | 117802.5273 | 0.9487646 | 0.1191 | 0.1256 | 0.96708 | 0.82153 | 1.05768 | 0.94876 | 0.0756 | |
| AMP | C00020 | NT2 | C13-03 | 10 | 237781.0896 | 208624.2609 | 148033.0259 | 0.862441 | 0.1335 | 0.1548 | 0.98867 | 0.87596 | 0.72269 | 0.86244 | 0.16402 | |
| AMP | C00020 | I3sg5 | C13-03 | 10 | 126108.2727 | 196454.6098 | 103448.2176 | 1.0241427 | 0.0963 | 0.094 | 1.12128 | 1.02234 | 0.9288 | 1.02414 | 0.16402 | |
| AMP | C00020 | NT2 | C13-04 | 10 | 382545.0338 | 389515.6395 | 258990.8723 | 1.4968152 | 0.2025 | 0.1353 | 1.59059 | 1.63547 | 1.26439 | 1.49682 | 0.87958 | |
| AMP | C00020 | I3sg5 | C13-04 | 10 | 146606.1529 | 243277.5503 | 229224.6774 | 1.5425418 | 0.4469 | 0.2897 | 1.30354 | 1.266 | 2.05808 | 1.54254 | 0.87958 | |
| AMP | C00020 | NT2 | C13-05 | 10 | 16688830.12 | 16534954.68 | 14427319.26 | 69.750087 | 0.5923 | 0.0085 | 69.3907 | 69.4258 | 70.4338 | 69.7501 | 0.50786 | |
| AMP | C00020 | I3sg5 | C13-05 | 10 | 7833813.799 | 13497708.12 | 7482197.293 | 69.02459 | 1.6255 | 0.0235 | 69.6538 | 70.2414 | 67.1785 | 69.0246 | 0.50786 | |
| AMP | C00020 | NT2 | C13-06 | 10 | 1375672.561 | 1216329.605 | 948727.6669 | 5.1528756 | 0.5456 | 0.1059 | 5.71993 | 5.10704 | 4.63166 | 5.15288 | 0.01873 | * |
| AMP | C00020 | I3sg5 | C13-06 | 10 | 382495.1196 | 705185.9068 | 456136.8922 | 3.7220282 | 0.3502 | 0.0941 | 3.40093 | 3.66975 | 4.0954 | 3.72203 | 0.01873 | * |
| AMP | C00020 | NT2 | C13-07 | 10 | 426752.1533 | 337607.563 | 235545.2759 | 1.4472821 | 0.3133 | 0.2165 | 1.7744 | 1.41752 | 1.14993 | 1.44728 | 0.13858 | |
| AMP | C00020 | I3sg5 | C13-07 | 10 | 109329.7629 | 197209.362 | 138358.1007 | 1.0802024 | 0.1429 | 0.1323 | 0.9721 | 1.02627 | 1.24224 | 1.0802 | 0.13858 | |
| AMP | C00020 | NT2 | C13-08 | 10 | 42009.57479 | 66559.1073 | 0 | 0.1513786 | 0.1412 | 0.9326 | 0.17467 | 0.27946 | 0 | 0.15138 | 0.13685 | |
| AMP | C00020 | I3sg5 | C13-08 | 10 | 0 | 0 | 0 | 0 | 0 | NA | 0 | 0 | 0 | 0 | 0.13685 | |
| Arg-Succ | C03406 | NT2 | C12 PARENT | 10 | 144235.188 | 93192.2119 | 102910.6968 | 86.074386 | 7.2799 | 0.0846 | 93.2036 | 78.6527 | 86.3668 | 86.0744 | 0.17904 | |
| Arg-Succ | C03406 | I3sg5 | C12 PARENT | 10 | 81337.45273 | 119740.3274 | 14705.31558 | 94.614803 | 5.4455 | 0.0576 | 89.1109 | 94.7335 | 100 | 94.6148 | 0.17904 | |
| Arg-Succ | C03406 | NT2 | C13-02 | 10 | 0 | 19133.66647 | 8214.499468 | 7.6808138 | 8.103 | 1.055 | 0 | 16.1485 | 6.89394 | 7.68081 | 0.70466 | |
| Arg-Succ | C03406 | I3sg5 | C13-02 | 10 | 9939.190955 | 6656.706727 | 0 | 5.3851967 | 5.4455 | 1.0112 | 10.8891 | 5.26651 | 0 | 5.3852 | 0.70466 | |
| Arg-Succ | C03406 | NT2 | C13-03 | 10 | 10517.57777 | 6159.846269 | 8030.134947 | 6.2447998 | 0.9063 | 0.1451 | 6.79637 | 5.19881 | 6.73922 | 6.2448 | 0.00028 | *** |
| Arg-Succ | C03406 | I3sg5 | C13-03 | 10 | 0 | 0 | 0 | 0 | 0 | NA | 0 | 0 | 0 | 0 | 0.00028 | *** |
| Asn | C00152 | NT2 | C12 PARENT | 4 | 2193066.117 | 2117512.489 | 1927887.164 | 77.95279 | 8.2876 | 0.1063 | 72.2819 | 74.1126 | 87.4639 | 77.9528 | 0.94238 | |
| Asn | C00152 | I3sg5 | C12 PARENT | 4 | 1527369.537 | 1877092.971 | 1252089.27 | 78.503293 | 9.2168 | 0.1174 | 88.73 | 75.9414 | 70.8385 | 78.5033 | 0.94238 | |
| Asn | C00152 | NT2 | C13-01 | 4 | 128932.2728 | 4274.923981 | 0 | 1.4663783 | 2.4114 | 1.6445 | 4.24951 | 0.14962 | 0 | 1.46638 | 0.95745 | |
| Asn | C00152 | I3sg5 | C13-01 | 4 | 12586.29683 | 34546.533 | 44605.09641 | 1.5508045 | 0.906 | 0.5842 | 0.73118 | 1.39765 | 2.52359 | 1.5508 | 0.95745 | |
| Asn | C00152 | NT2 | C13-02 | 4 | 318033.7701 | 330590.6791 | 175296.5058 | 10.001864 | 1.8561 | 0.1856 | 10.4822 | 11.5706 | 7.95281 | 10.0019 | 0.76295 | |
| Asn | C00152 | I3sg5 | C13-02 | 4 | 0 | 272252.7958 | 258221.5636 | 8.5412345 | 7.6122 | 0.8912 | 0 | 11.0145 | 14.6092 | 8.54123 | 0.76295 | |
| Asn | C00152 | NT2 | C13-03 | 4 | 371794.9796 | 363126.0835 | 101025.4844 | 9.8489158 | 4.5658 | 0.4636 | 12.2541 | 12.7094 | 4.5833 | 9.84892 | 0.65469 | |
| Asn | C00152 | I3sg5 | C13-03 | 4 | 181411.1566 | 287873.4757 | 198056.093 | 11.130173 | 0.5576 | 0.0501 | 10.5388 | 11.6465 | 11.2053 | 11.1302 | 0.65469 | |
| Asn | C00152 | NT2 | C13-04 | 4 | 22220.83765 | 41650.87783 | 0 | 0.7300523 | 0.7289 | 0.9984 | 0.73238 | 1.45777 | 0 | 0.73005 | 0.41585 | |
| Asn | C00152 | I3sg5 | C13-04 | 4 | 0 | 0 | 14555.35689 | 0.2744956 | 0.4754 | 1.7321 | 0 | 0 | 0.82349 | 0.2745 | 0.41585 | |
| Asp | C00049 | NT2 | C12 PARENT | 4 | 33247027.67 | 32009841.61 | 27000262.92 | 63.962103 | 0.8359 | 0.0131 | 63.0765 | 64.0724 | 64.7373 | 63.9621 | 0.80636 | |
| Asp | C00049 | I3sg5 | C12 PARENT | 4 | 15042944.19 | 20009742.13 | 17467867.3 | 63.776595 | 0.8983 | 0.0141 | 62.8129 | 63.9263 | 64.5906 | 63.7766 | 0.80636 | |
| Asp | C00049 | NT2 | C13-01 | 4 | 1714620.872 | 1348818.57 | 977062.1631 | 2.7651705 | 0.4587 | 0.1659 | 3.25299 | 2.69986 | 2.34266 | 2.76517 | 0.46247 | |
| Asp | C00049 | I3sg5 | C13-01 | 4 | 491500.1796 | 938902.8705 | 621579.4786 | 2.4500882 | 0.4915 | 0.2006 | 2.05229 | 2.99957 | 2.2984 | 2.45009 | 0.46247 | |
| Asp | C00049 | NT2 | C13-02 | 4 | 8367980.843 | 7724329.893 | 6547056.214 | 15.67826 | 0.2079 | 0.0133 | 15.8758 | 15.4614 | 15.6976 | 15.6783 | 0.62503 | |
| Asp | C00049 | I3sg5 | C13-02 | 4 | 3820629.501 | 4771345.098 | 4179329 | 15.550144 | 0.3647 | 0.0235 | 15.9533 | 15.2433 | 15.4538 | 15.5501 | 0.62503 | |
| Asp | C00049 | NT2 | C13-03 | 4 | 6599225.527 | 6368406.671 | 5443877.181 | 12.773319 | 0.2672 | 0.0209 | 12.5201 | 12.7473 | 13.0525 | 12.7733 | 0.31607 | |
| Asp | C00049 | I3sg5 | C13-03 | 4 | 3327092.07 | 4033694.619 | 3471339.557 | 13.205032 | 0.5959 | 0.0451 | 13.8925 | 12.8867 | 12.8359 | 13.205 | 0.31607 | |
| Asp | C00049 | NT2 | C13-04 | 4 | 2780173.652 | 2507436.975 | 1739144.137 | 4.8211479 | 0.5783 | 0.12 | 5.27457 | 5.01901 | 4.16987 | 4.82115 | 0.61528 | |
| Asp | C00049 | I3sg5 | C13-04 | 4 | 1266648.859 | 1547582.982 | 1303867.121 | 5.0181405 | 0.2425 | 0.0483 | 5.28898 | 4.94415 | 4.82128 | 5.01814 | 0.61528 | |
| ATP | C00002 | NT2 | C12 PARENT | 10 | 12560540.38 | 10768324.18 | 11946929.47 | 19.285598 | 0.6955 | 0.0361 | 18.5626 | 19.3442 | 19.95 | 19.2856 | 0.11004 | |
| ATP | C00002 | I3sg5 | C12 PARENT | 10 | 7315920.476 | 7967076.399 | 5176762.463 | 20.676605 | 0.9493 | 0.0459 | 20.5191 | 21.6948 | 19.8159 | 20.6766 | 0.11004 | |
| ATP | C00002 | NT2 | C13-01 | 10 | 0 | 0 | 0 | 0 | 0 | NA | 0 | 0 | 0 | 0 | NA | |
| ATP | C00002 | I3sg5 | C13-01 | 10 | 0 | 0 | 0 | 0 | 0 | NA | 0 | 0 | 0 | 0 | NA | |
| ATP | C00002 | NT2 | C13-02 | 10 | 592262.3517 | 377943.5297 | 645852.5026 | 0.8775703 | 0.1998 | 0.2277 | 0.87527 | 0.67894 | 1.0785 | 0.87757 | 0.2096 | |
| ATP | C00002 | I3sg5 | C13-02 | 10 | 270279.0901 | 260006.9336 | 161534.6732 | 0.6948008 | 0.0708 | 0.1019 | 0.75806 | 0.70801 | 0.61833 | 0.6948 | 0.2096 | |
| ATP | C00002 | NT2 | C13-03 | 10 | 618964.7947 | 513707.9112 | 616068.3096 | 0.9554414 | 0.0636 | 0.0666 | 0.91474 | 0.92283 | 1.02876 | 0.95544 | 0.17209 | |
| ATP | C00002 | I3sg5 | C13-03 | 10 | 491007.1108 | 375356.9854 | 276442.664 | 1.1524793 | 0.1954 | 0.1695 | 1.37713 | 1.02212 | 1.05818 | 1.15248 | 0.17209 | |
| ATP | C00002 | NT2 | C13-04 | 10 | 1163767.906 | 852082.4252 | 742612.8141 | 1.4968774 | 0.2417 | 0.1615 | 1.71987 | 1.53068 | 1.24008 | 1.49688 | 0.87127 | |
| ATP | C00002 | I3sg5 | C13-04 | 10 | 439698.1359 | 581114.8243 | 466445.0651 | 1.5337081 | 0.2793 | 0.1821 | 1.23323 | 1.58241 | 1.78549 | 1.53371 | 0.87127 | |
| ATP | C00002 | NT2 | C13-05 | 10 | 46903559.45 | 38882429.38 | 41720365.5 | 69.610991 | 0.2706 | 0.0039 | 69.3164 | 69.8484 | 69.6681 | 69.611 | 0.71154 | |
| ATP | C00002 | I3sg5 | C13-05 | 10 | 25250002.82 | 25519935.06 | 18066168.17 | 69.82204 | 0.8798 | 0.0126 | 70.819 | 69.4924 | 69.1547 | 69.822 | 0.71154 | |
| ATP | C00002 | NT2 | C13-06 | 10 | 4548159.611 | 3326235.355 | 3304659.03 | 6.0717142 | 0.6073 | 0.1 | 6.7215 | 5.97525 | 5.51839 | 6.07171 | 0.10425 | |
| ATP | C00002 | I3sg5 | C13-06 | 10 | 1423429.7 | 1393506.095 | 1520056.771 | 4.5351591 | 1.1158 | 0.246 | 3.99231 | 3.7946 | 5.81856 | 4.53516 | 0.10425 | |
| ATP | C00002 | NT2 | C13-07 | 10 | 1192634.479 | 763333.9525 | 819942.2845 | 1.5009987 | 0.2265 | 0.1509 | 1.76253 | 1.37125 | 1.36921 | 1.501 | 0.99356 | |
| ATP | C00002 | I3sg5 | C13-07 | 10 | 454347.2918 | 572885.9658 | 434751.2378 | 1.4994946 | 0.2018 | 0.1346 | 1.27431 | 1.56 | 1.66417 | 1.49949 | 0.99356 | |
| ATP | C00002 | NT2 | C13-08 | 10 | 85995.64044 | 182792.2825 | 88012.28905 | 0.200809 | 0.1109 | 0.5523 | 0.12709 | 0.32837 | 0.14697 | 0.20081 | 0.18829 | |
| ATP | C00002 | I3sg5 | C13-08 | 10 | 9568.853656 | 53489.41287 | 22113.245 | 0.0857131 | 0.0594 | 0.6932 | 0.02684 | 0.14565 | 0.08465 | 0.08571 | 0.18829 | |
| Carbamoyl-Asp | C00438 | NT2 | C12 PARENT | 5 | 6529014.76 | 5545414.936 | 7033159.171 | 72.692384 | 0.9553 | 0.0131 | 73.2872 | 73.1995 | 71.5904 | 72.6924 | 0.11346 | |
| Carbamoyl-Asp | C00438 | I3sg5 | C12 PARENT | 5 | 8361222.126 | 8127675.226 | 5694983.676 | 71.317581 | 0.6903 | 0.0097 | 70.5342 | 71.5817 | 71.8368 | 71.3176 | 0.11346 | |
| Carbamoyl-Asp | C00438 | NT2 | C13-01 | 5 | 15206.37913 | 0 | 0 | 0.0568965 | 0.0985 | 1.7321 | 0.17069 | 0 | 0 | 0.0569 | 0.69167 | |
| Carbamoyl-Asp | C00438 | I3sg5 | C13-01 | 5 | 3509.038569 | 0 | 21711.33828 | 0.1011566 | 0.1503 | 1.4858 | 0.0296 | 0 | 0.27387 | 0.10116 | 0.69167 | |
| Carbamoyl-Asp | C00438 | NT2 | C13-02 | 5 | 0 | 0 | 0 | 0 | 0 | NA | 0 | 0 | 0 | 0 | NA | |
| Carbamoyl-Asp | C00438 | I3sg5 | C13-02 | 5 | 0 | 0 | 0 | 0 | 0 | NA | 0 | 0 | 0 | 0 | NA | |
| Carbamoyl-Asp | C00438 | NT2 | C13-03 | 5 | 2347089.412 | 2030341.293 | 2791003.285 | 27.185276 | 1.0844 | 0.0399 | 26.3457 | 26.8005 | 28.4096 | 27.1853 | 0.17142 | |
| Carbamoyl-Asp | C00438 | I3sg5 | C13-03 | 5 | 3447920.966 | 3226729.776 | 2198688.289 | 28.41297 | 0.6759 | 0.0238 | 29.0862 | 28.4183 | 27.7344 | 28.413 | 0.17142 | |
| Carbamoyl-Asp | C00438 | NT2 | C13-04 | 5 | 17490.64389 | 0 | 0 | 0.0654433 | 0.1134 | 1.7321 | 0.19633 | 0 | 0 | 0.06544 | 0.44165 | |
| Carbamoyl-Asp | C00438 | I3sg5 | C13-04 | 5 | 41483.18397 | 0 | 12282.32674 | 0.1682923 | 0.1754 | 1.042 | 0.34995 | 0 | 0.15493 | 0.16829 | 0.44165 | |
| CDP | C00112 | NT2 | C12 PARENT | 9 | 1065986.24 | 961927.991 | 906641.2083 | 38.757314 | 1.4048 | 0.0362 | 39.2988 | 39.8109 | 37.1623 | 38.7573 | 0.05982 | |
| CDP | C00112 | I3sg5 | C12 PARENT | 9 | 685152.7104 | 640986.3958 | 598070.6393 | 42.886118 | 2.3603 | 0.055 | 45.0192 | 40.3505 | 43.2887 | 42.8861 | 0.05982 | |
| CDP | C00112 | NT2 | C13-04 | 9 | 0 | 0 | 0 | 0 | 0 | NA | 0 | 0 | 0 | 0 | NA | |
| CDP | C00112 | I3sg5 | C13-04 | 9 | 0 | 0 | 0 | 0 | 0 | NA | 0 | 0 | 0 | 0 | NA | |
| CDP | C00112 | NT2 | C13-05 | 9 | 1114666.064 | 991605.8754 | 1077932.959 | 42.105307 | 1.7999 | 0.0427 | 41.0934 | 41.0391 | 44.1834 | 42.1053 | 0.30391 | |
| CDP | C00112 | I3sg5 | C13-05 | 9 | 624010.0053 | 660500.3967 | 544292.1733 | 40.658922 | 1.131 | 0.0278 | 41.0017 | 41.5789 | 39.3961 | 40.6589 | 0.30391 | |
| CDP | C00112 | NT2 | C13-06 | 9 | 183363.437 | 109537.1672 | 126513.4462 | 5.492973 | 1.1446 | 0.2084 | 6.7599 | 4.53336 | 5.18566 | 5.49297 | 0.12019 | |
| CDP | C00112 | I3sg5 | C13-06 | 9 | 0 | 73654.92257 | 40579.90439 | 2.5246052 | 2.3457 | 0.9291 | 0 | 4.63662 | 2.93719 | 2.52461 | 0.12019 | |
| CDP | C00112 | NT2 | C13-07 | 9 | 218319.6137 | 200971.5321 | 257216.0043 | 8.9697119 | 1.3691 | 0.1526 | 8.04859 | 8.31751 | 10.543 | 8.96971 | 0.71261 | |
| CDP | C00112 | I3sg5 | C13-07 | 9 | 132453.9333 | 140755.5998 | 145157.344 | 9.3567804 | 0.9989 | 0.1068 | 8.70313 | 8.86065 | 10.5066 | 9.35678 | 0.71261 | |
| CDP | C00112 | NT2 | C13-08 | 9 | 130183.4187 | 152202.4066 | 71375.18796 | 4.6746943 | 1.6902 | 0.3616 | 4.79936 | 6.29913 | 2.9256 | 4.67469 | 0.92837 | |
| CDP | C00112 | I3sg5 | C13-08 | 9 | 80294.90957 | 72650.03526 | 53487.25977 | 4.5735744 | 0.7022 | 0.1535 | 5.27592 | 4.57336 | 3.87144 | 4.57357 | 0.92837 | |
| CDP-choline | C00307 | NT2 | C12 PARENT | 14 | 72279.46216 | 71387.53505 | 80579.9398 | 54.179768 | 1.9399 | 0.0358 | 54.9711 | 55.5989 | 51.9693 | 54.1798 | 0.69517 | |
| CDP-choline | C00307 | I3sg5 | C12 PARENT | 14 | 95822.57364 | 136381.1292 | 68196.10656 | 55.906966 | 6.8303 | 0.1222 | 48.8108 | 56.4741 | 62.436 | 55.907 | 0.69517 | |
| CDP-choline | C00307 | NT2 | C13-05 | 14 | 57617.63029 | 57009.90009 | 72241.93566 | 44.937726 | 1.4616 | 0.0325 | 43.8203 | 44.4011 | 46.5918 | 44.9377 | 0.40851 | |
| CDP-choline | C00307 | I3sg5 | C13-05 | 14 | 94594.74406 | 96173.93517 | 41029.55751 | 41.858033 | 5.595 | 0.1337 | 48.1854 | 39.8247 | 37.564 | 41.858 | 0.40851 | |
| CDP-choline | C00307 | NT2 | C13-06 | 14 | 0 | 0 | 0 | 0 | 0 | NA | 0 | 0 | 0 | 0 | NA | |
| CDP-choline | C00307 | I3sg5 | C13-06 | 14 | 0 | 0 | 0 | 0 | 0 | NA | 0 | 0 | 0 | 0 | NA | |
| CDP-choline | C00307 | NT2 | C13-07 | 14 | 1589.174846 | 0 | 2231.04676 | 0.882506 | 0.7729 | 0.8758 | 1.20862 | 0 | 1.43889 | 0.88251 | 0.32977 | |
| CDP-choline | C00307 | I3sg5 | C13-07 | 14 | 5896.929107 | 8938.098782 | 0 | 2.2350007 | 1.9667 | 0.88 | 3.00382 | 3.70118 | 0 | 2.235 | 0.32977 | |
| CDP-EtA | C00570 | NT2 | C12 PARENT | 11 | 464925.9679 | 475639.0581 | 454581.9064 | 45.412629 | 1.556 | 0.0343 | 46.2337 | 46.3862 | 43.618 | 45.4126 | 0.11823 | |
| CDP-EtA | C00570 | I3sg5 | C12 PARENT | 11 | 452739.2387 | 588837.3079 | 316156.1012 | 47.535565 | 1.0064 | 0.0212 | 47.4167 | 46.5939 | 48.5961 | 47.5356 | 0.11823 | |
| CDP-EtA | C00570 | NT2 | C13-05 | 11 | 424060.6623 | 421224.848 | 413632.1367 | 40.979406 | 1.2436 | 0.0303 | 42.1699 | 41.0795 | 39.6888 | 40.9794 | 0.26857 | |
| CDP-EtA | C00570 | I3sg5 | C13-05 | 11 | 389407.5619 | 544806.3929 | 280515.9854 | 42.337155 | 1.3453 | 0.0318 | 40.7838 | 43.1098 | 43.1179 | 42.3372 | 0.26857 | |

| | | | | | | | | | | | | | | | | |
|---|---|---|---|---|---|---|---|---|---|---|---|---|---|---|---|---|
| CDP-EtA | C00570 | NT2 | C13-06 | 11 | 45205.28205 | 30806.57452 | 18677.56516 | 3.0972931 | 1.354 | 0.4372 | 4.49535 | 3.00438 | 1.79215 | 3.09729 | 0.15951 | |
| CDP-EtA | C00570 | I3sg5 | C13-06 | 11 | 12655.85062 | 31135.45273 | 3121.054663 | 1.4229751 | 0.9956 | 0.6996 | 1.32548 | 2.46371 | 0.47973 | 1.42298 | 0.15951 | |
| CDP-EtA | C00570 | NT2 | C13-07 | 11 | 41390.66054 | 79322.63367 | 112007.581 | 7.533073 | 3.3203 | 0.4408 | 4.11601 | 7.73586 | 10.7473 | 7.53307 | 0.72376 | |
| CDP-EtA | C00570 | I3sg5 | C13-07 | 11 | 76595.58503 | 72812.18572 | 42332.31827 | 6.7634912 | 1.1519 | 0.1703 | 8.02207 | 5.76153 | 6.50687 | 6.76349 | 0.72376 | |
| CDP-EtA | C00570 | NT2 | C13-08 | 11 | 30018.07706 | 18396.15119 | 43288.75059 | 2.9775986 | 1.1798 | 0.3962 | 2.98509 | 1.79407 | 4.15364 | 2.9776 | 0.24467 | |
| CDP-EtA | C00570 | I3sg5 | C13-08 | 11 | 23411.9628 | 26173.02127 | 8453.63088 | 1.9408131 | 0.5872 | 0.3026 | 2.452 | 2.07104 | 1.2994 | 1.94081 | 0.24467 | |
| Cit | C00158 | NT2 | C12 PARENT | 6 | 40435219.2 | 32855413.39 | 35352424.84 | 32.304049 | 1.0362 | 0.0321 | 32.1825 | 31.334 | 33.3957 | 32.304 | 0.03479 | * |
| Cit | C00158 | I3sg5 | C12 PARENT | 6 | 27197054.13 | 30701755.79 | 23379102.11 | 34.804696 | 0.9092 | 0.0261 | 35.2336 | 35.4202 | 33.7603 | 34.8047 | 0.03479 | * |
| Cit | C00158 | NT2 | C13-01 | 6 | 1760910.248 | 1744938.334 | 1857030.902 | 1.6066311 | 0.1833 | 0.1141 | 1.40151 | 1.66414 | 1.75425 | 1.60663 | 0.8325 | |
| Cit | C00158 | I3sg5 | C13-01 | 6 | 1214118.97 | 1409166.877 | 1071974.618 | 1.5821969 | 0.0397 | 0.0251 | 1.57288 | 1.62574 | 1.54797 | 1.5822 | 0.8325 | |
| Cit | C00158 | NT2 | C13-02 | 6 | 44076753.79 | 37571323.52 | 36830109.91 | 35.234627 | 0.5368 | 0.0152 | 35.0808 | 35.8315 | 34.7916 | 35.2346 | 0.07053 | |
| Cit | C00158 | I3sg5 | C13-02 | 6 | 25632146.3 | 29253855.74 | 24057508.39 | 33.898662 | 0.7776 | 0.0229 | 33.2062 | 33.7498 | 34.74 | 33.8987 | 0.07053 | |
| Cit | C00158 | NT2 | C13-03 | 6 | 7647603.701 | 6658655.482 | 6431144.19 | 6.1707474 | 0.1556 | 0.0252 | 6.08674 | 6.35031 | 6.07518 | 6.17075 | 0.3132 | |
| Cit | C00158 | I3sg5 | C13-03 | 6 | 4957340.619 | 5413857.288 | 4309693.535 | 6.2971506 | 0.1089 | 0.0173 | 6.42219 | 6.24589 | 6.22337 | 6.29715 | 0.3132 | |
| Cit | C00158 | NT2 | C13-04 | 6 | 15891207.41 | 13232599.41 | 12767945.02 | 12.442977 | 0.3309 | 0.0266 | 12.6478 | 12.6198 | 12.0612 | 12.443 | 0.0334 | * |
| Cit | C00158 | I3sg5 | C13-04 | 6 | 9092936.133 | 9870962.572 | 8214732.659 | 11.676737 | 0.2534 | 0.0217 | 11.7798 | 11.388 | 11.8624 | 11.6767 | 0.0334 | * |
| Cit | C00158 | NT2 | C13-05 | 6 | 11529232.97 | 9504529.134 | 9425684.469 | 9.0481745 | 0.1368 | 0.0151 | 9.17614 | 9.0644 | 8.90398 | 9.04817 | 0.49238 | |
| Cit | C00158 | I3sg5 | C13-05 | 6 | 7051019.874 | 7836766.607 | 6348175.286 | 9.1142401 | 0.0653 | 0.0072 | 9.13454 | 9.04117 | 9.16701 | 9.11424 | 0.49238 | |
| Cit | C00158 | NT2 | C13-06 | 6 | 4302678.722 | 3288066.433 | 3194900.041 | 3.1927939 | 0.2091 | 0.0655 | 3.42451 | 3.13581 | 3.01806 | 3.19279 | 0.01237 | * |
| Cit | C00158 | I3sg5 | C13-06 | 6 | 2046161.717 | 2192306.781 | 1869017.603 | 2.626318 | 0.0875 | 0.0333 | 2.65079 | 2.52923 | 2.69893 | 2.62632 | 0.01237 | * |
| CMP | C00055 | NT2 | C12 PARENT | 9 | 137666.5794 | 141783.7434 | 228789.1758 | 60.277907 | 17.1123 | 0.2839 | 65.9401 | 41.0522 | 73.8414 | 60.2779 | 0.74184 | |
| CMP | C00055 | I3sg5 | C12 PARENT | 9 | 191014.6187 | 138034.3999 | 48527.82669 | 53.971137 | 25.7733 | 0.4775 | 77.7718 | 57.5437 | 26.5979 | 53.9711 | 0.74184 | |
| CMP | C00055 | NT2 | C13-05 | 9 | 71108.56315 | 203590.8459 | 81049.29429 | 39.722093 | 17.1123 | 0.4308 | 34.0599 | 58.9478 | 26.1586 | 39.7221 | 0.74184 | |
| CMP | C00055 | I3sg5 | C13-05 | 9 | 54594.49 | 101842.9515 | 133922.2032 | 46.028863 | 25.7733 | 0.5599 | 22.2282 | 42.4563 | 73.4021 | 46.0289 | 0.74184 | |
| CoA | C00010 | NT2 | C12 PARENT | 21 | 103303.0562 | 97997.87305 | 83816.9873 | 31.723587 | 0.3952 | 0.0125 | 31.6396 | 31.3771 | 32.1541 | 31.7236 | 0.00318 | ** |
| CoA | C00010 | I3sg5 | C12 PARENT | 21 | 60147.09046 | 77772.6126 | 52827.74306 | 35.392861 | 0.9221 | 0.0261 | 36.4576 | 34.85 | 34.871 | 35.3929 | 0.00318 | ** |
| CoA | C00010 | NT2 | C13-01 | 21 | 0 | 0 | 0 | 0 | 0 | NA | 0 | 0 | 0 | 0 | NA | |
| CoA | C00010 | I3sg5 | C13-01 | 21 | 0 | 0 | 0 | 0 | 0 | NA | 0 | 0 | 0 | 0 | NA | |
| CoA | C00010 | NT2 | C13-02 | 21 | 0 | 0 | 0 | 0 | 0 | NA | 0 | 0 | 0 | 0 | NA | |
| CoA | C00010 | I3sg5 | C13-02 | 21 | 0 | 0 | 0 | 0 | 0 | NA | 0 | 0 | 0 | 0 | NA | |
| CoA | C00010 | NT2 | C13-03 | 21 | 3205.07413 | 3886.622776 | 2184.797583 | 1.0214032 | 0.206 | 0.2017 | 0.98165 | 1.24442 | 0.83814 | 1.0214 | 0.03548 | * |
| CoA | C00010 | I3sg5 | C13-03 | 21 | 1080.644241 | 992.8734239 | 1006.124419 | 0.5880205 | 0.124 | 0.2109 | 0.65502 | 0.44491 | 0.66413 | 0.58802 | 0.03548 | * |
| CoA | C00010 | NT2 | C13-04 | 21 | 6017.665869 | 4668.940252 | 4222.447894 | 1.6526067 | 0.1764 | 0.1067 | 1.84309 | 1.49491 | 1.61983 | 1.65261 | 0.32929 | |
| CoA | C00010 | I3sg5 | C13-04 | 21 | 3829.76521 | 3177.260597 | 3354.559244 | 1.9864724 | 0.4903 | 0.2468 | 2.32137 | 1.42373 | 2.21431 | 1.98647 | 0.32929 | |
| CoA | C00010 | NT2 | C13-05 | 21 | 196290.3688 | 195234.2223 | 164013.7111 | 61.849794 | 1.5122 | 0.0244 | 60.1197 | 62.5104 | 62.9193 | 61.8498 | 0.22576 | |
| CoA | C00010 | I3sg5 | C13-05 | 21 | 97176.19147 | 137637.166 | 90668.01178 | 60.142244 | 1.4095 | 0.0234 | 58.9024 | 61.6753 | 59.849 | 60.1422 | 0.22576 | |
| CoA | C00010 | NT2 | C13-06 | 21 | 17573.49576 | 8723.113671 | 5027.970309 | 3.3680732 | 1.7972 | 0.5336 | 5.3824 | 2.79298 | 1.92884 | 3.36807 | 0.15061 | |
| CoA | C00010 | I3sg5 | C13-06 | 21 | 2288.742464 | 3001.155444 | 2719.667451 | 1.5091144 | 0.2487 | 0.1648 | 1.3873 | 1.34482 | 1.79522 | 1.50911 | 0.15061 | |
| CoA | C00010 | NT2 | C13-07 | 21 | 0 | 0 | 0 | 0 | 0 | NA | 0 | 0 | 0 | 0 | NA | |
| CoA | C00010 | I3sg5 | C13-07 | 21 | 0 | 0 | 0 | 0 | 0 | NA | 0 | 0 | 0 | 0 | NA | |
| CoA | C00010 | NT2 | C13-08 | 21 | 109.6562377 | 1812.202904 | 1407.088221 | 0.3845365 | 0.3046 | 0.7921 | 0.03359 | 0.58023 | 0.53979 | 0.38454 | 0.98833 | |
| CoA | C00010 | I3sg5 | C13-08 | 21 | 455.9239134 | 582.9806906 | 918.4751468 | 0.3812879 | 0.195 | 0.5114 | 0.27635 | 0.26123 | 0.60628 | 0.38129 | 0.98833 | |
| CoA | C00010 | NT2 | C13-09 | 21 | 0 | 0 | 0 | 0 | 0 | NA | 0 | 0 | 0 | 0 | NA | |
| CoA | C00010 | I3sg5 | C13-09 | 21 | 0 | 0 | 0 | 0 | 0 | NA | 0 | 0 | 0 | 0 | NA | |
| Creatine | C00300 | NT2 | C12 PARENT | 4 | 369744217 | 358686421.4 | 212381653.5 | 97.727609 | 0.3017 | 0.0031 | 97.4662 | 97.6589 | 98.0577 | 97.7276 | 0.13067 | |
| Creatine | C00300 | I3sg5 | C12 PARENT | 4 | 222119613.4 | 241482179.1 | 210034931.1 | 98.115455 | 0.1854 | 0.0019 | 98.2235 | 98.2214 | 97.9014 | 98.1155 | 0.13067 | |
| Creatine | C00300 | NT2 | C13-01 | 4 | 0 | 0 | 0 | 0 | 0 | NA | 0 | 0 | 0 | 0 | NA | |
| Creatine | C00300 | I3sg5 | C13-01 | 4 | 0 | 0 | 0 | 0 | 0 | NA | 0 | 0 | 0 | 0 | NA | |
| Creatine | C00300 | NT2 | C13-02 | 4 | 9612270.265 | 8598366.617 | 4206738.922 | 2.2723906 | 0.3017 | 0.1328 | 2.53384 | 2.34106 | 1.94227 | 2.27239 | 0.13067 | |
| Creatine | C00300 | I3sg5 | C13-02 | 4 | 4017227.402 | 4372756.045 | 4502241.058 | 1.8845446 | 0.1854 | 0.0984 | 1.77646 | 1.77859 | 2.09858 | 1.88454 | 0.13067 | |
| Creatinine | C00791 | NT2 | C12 PARENT | 4 | 912631.6023 | 840591.8736 | 578311.292 | 98.831438 | 1.0028 | 0.0101 | 99.7586 | 97.767 | 98.9687 | 98.8314 | 0.62773 | |
| Creatinine | C00791 | I3sg5 | C12 PARENT | 4 | 266641.5869 | 700156.4136 | 402312.555 | 98.072246 | 2.2984 | 0.0234 | 95.5286 | 100 | 98.6881 | 98.0722 | 0.62773 | |
| Creatinine | C00791 | NT2 | C13-01 | 4 | 2208.810739 | 19198.70702 | 6026.221244 | 1.1685616 | 1.0028 | 0.8582 | 0.24144 | 2.23295 | 1.03129 | 1.16856 | 0.62773 | |
| Creatinine | C00791 | I3sg5 | C13-01 | 4 | 12480.6234 | 0 | 5348.014913 | 1.9277542 | 2.2984 | 1.1923 | 4.47138 | 0 | 1.31188 | 1.92775 | 0.62773 | |
| CTP | C00063 | NT2 | C12 PARENT | 9 | 1223032.741 | 1041918.343 | 1061645.142 | 41.429129 | 0.967 | 0.0233 | 42.2265 | 40.3535 | 41.7073 | 41.4291 | 0.08621 | |
| CTP | C00063 | I3sg5 | C12 PARENT | 9 | 750548.0477 | 772116.5957 | 493698.2106 | 40.08786 | 0.3421 | 0.0085 | 39.8178 | 39.9733 | 40.4725 | 40.0879 | 0.08621 | |
| CTP | C00063 | NT2 | C13-01 | 9 | 0 | 0 | 0 | 0 | 0 | NA | 0 | 0 | 0 | 0 | NA | |
| CTP | C00063 | I3sg5 | C13-01 | 9 | 0 | 0 | 0 | 0 | 0 | NA | 0 | 0 | 0 | 0 | NA | |
| CTP | C00063 | NT2 | C13-04 | 9 | 0 | 0 | 0 | 0 | 0 | NA | 0 | 0 | 0 | 0 | NA | |
| CTP | C00063 | I3sg5 | C13-04 | 9 | 0 | 0 | 0 | 0 | 0 | NA | 0 | 0 | 0 | 0 | NA | |
| CTP | C00063 | NT2 | C13-05 | 9 | 1153918.527 | 1084420.408 | 1028514.502 | 40.748565 | 1.1198 | 0.0275 | 39.8403 | 41.9996 | 40.4058 | 40.7486 | 0.26128 | |
| CTP | C00063 | I3sg5 | C13-05 | 9 | 802798.1866 | 780795.3552 | 547289.8434 | 42.626062 | 2.2218 | 0.0521 | 42.5897 | 40.4227 | 44.8658 | 42.6261 | 0.26128 | |
| CTP | C00063 | NT2 | C13-06 | 9 | 145684.062 | 124075.185 | 113427.0294 | 4.7637935 | 0.2892 | 0.0607 | 5.0299 | 4.80544 | 4.45605 | 4.76379 | 0.83205 | |
| CTP | C00063 | I3sg5 | C13-06 | 9 | 113485.8699 | 73391.7253 | 60028.50756 | 4.9137346 | 1.1105 | 0.226 | 6.0206 | 3.79957 | 4.92103 | 4.91373 | 0.83205 | |
| CTP | C00063 | NT2 | C13-07 | 9 | 279830.881 | 190921.3794 | 214198.5402 | 8.490255 | 1.1354 | 0.1337 | 9.66146 | 7.39439 | 8.41491 | 8.49025 | 0.81781 | |
| CTP | C00063 | I3sg5 | C13-07 | 9 | 142141.7366 | 203739.7973 | 77332.08247 | 8.1427414 | 2.1678 | 0.2662 | 7.54084 | 10.5478 | 6.33954 | 8.14274 | 0.81781 | |
| CTP | C00063 | NT2 | C13-08 | 9 | 93895.85807 | 140639.7345 | 127678.7648 | 4.5682569 | 1.1687 | 0.2558 | 3.24185 | 5.44698 | 5.01593 | 4.56826 | 0.71607 | |
| CTP | C00063 | I3sg5 | C13-08 | 9 | 75984.58117 | 101535.2881 | 41487.99893 | 4.2296027 | 0.9435 | 0.2231 | 4.0311 | 5.2566 | 3.40111 | 4.2296 | 0.71607 | |
| Cystathionine | C02291 | NT2 | C12 PARENT | 7 | 1266176.239 | 1122762.697 | 805596.9876 | 98.523166 | 1.5731 | 0.016 | 98.7556 | 99.9672 | 96.8467 | 98.5232 | 0.17927 | |
| Cystathionine | C02291 | I3sg5 | C12 PARENT | 7 | 694193.1886 | 724643.6058 | 734288.3905 | 100 | 0 | 0 | 100 | 100 | 100 | 100 | 0.17927 | |
| Cystathionine | C02291 | NT2 | C13-01 | 7 | 15954.85351 | 368.9238088 | 26229.59614 | 1.4768339 | 1.5731 | 1.0652 | 1.2444 | 0.03285 | 3.15325 | 1.47683 | 0.17927 | |
| Cystathionine | C02291 | I3sg5 | C13-01 | 7 | 0 | 0 | 0 | 0 | 0 | NA | 0 | 0 | 0 | 0 | 0.17927 | |
| Cystathionine | C02291 | NT2 | C13-02 | 7 | 0 | 0 | 0 | 0 | 0 | NA | 0 | 0 | 0 | 0 | NA | |
| Cystathionine | C02291 | I3sg5 | C13-02 | 7 | 0 | 0 | 0 | 0 | 0 | NA | 0 | 0 | 0 | 0 | NA | |
| Cytosine | C00380 | NT2 | C12 PARENT | 4 | 235258.5368 | 208010.4225 | 107355.753 | 100 | 0 | 0 | 100 | 100 | 100 | 100 | 0.28312 | |
| Cytosine | C00380 | I3sg5 | C12 PARENT | 4 | 75531.41785 | 105900.7874 | 4011.985809 | 100 | 0 | 0 | 100 | 100 | 100 | 100 | 0.28312 | |
| F16BP | C00354 | NT2 | C12 PARENT | 6 | 68594.58793 | 39643.62912 | 28791.69551 | 3.6480892 | 0.949 | 0.2601 | 4.7345 | 3.22928 | 2.98049 | 3.64809 | 0.17377 | |
| F16BP | C00354 | I3sg5 | C12 PARENT | 6 | 0 | 17979.87363 | 16369.28115 | 1.8455861 | 1.6336 | 0.8851 | 0 | 2.43078 | 3.10597 | 1.84559 | 0.17377 | |
| F16BP | C00354 | NT2 | C13-05 | 6 | 0 | 0 | 0 | 0 | 0 | NA | 0 | 0 | 0 | 0 | NA | |
| F16BP | C00354 | I3sg5 | C13-05 | 6 | 0 | 0 | 0 | 0 | 0 | NA | 0 | 0 | 0 | 0 | NA | |
| F16BP | C00354 | NT2 | C13-06 | 6 | 1380231 | 1187986.597 | 937213.5216 | 96.351911 | 0.949 | 0.0098 | 95.2655 | 96.7707 | 97.0195 | 96.3519 | 0.17377 | |
| F16BP | C00354 | I3sg5 | C13-06 | 6 | 587693.973 | 721693.7306 | 510656.4863 | 98.154414 | 1.6336 | 0.0166 | 100 | 97.5692 | 96.894 | 98.1544 | 0.17377 | |
| G6P-F6P | C00085 | NT2 | C12 PARENT | 6 | 322049.0401 | 268590.5654 | 361015.1542 | 27.957913 | 0.3933 | 0.0141 | 28.0691 | 27.521 | 28.2837 | 27.9579 | 0.39772 | |
| G6P-F6P | C00085 | I3sg5 | C12 PARENT | 6 | 281057.2928 | 261539.1386 | 153528.8863 | 25.205685 | 5.0238 | 0.1993 | 26.5869 | 29.3944 | 19.6358 | 25.2057 | 0.39772 | |
| G6P-F6P | C00085 | NT2 | C13-05 | 6 | 0 | 6423.513607 | 30467.14753 | 1.015042 | 1.2328 | 1.2146 | 0 | 0.65818 | 2.38694 | 1.01504 | 0.59934 | |
| G6P-F6P | C00085 | I3sg5 | C13-05 | 6 | 0 | 13737.03434 | 0 | 0.5146346 | 0.8914 | 1.7321 | 0 | 1.5439 | 0 | 0.51463 | 0.59934 | |
| G6P-F6P | C00085 | NT2 | C13-06 | 6 | 825296.6216 | 700932.8643 | 884926.2619 | 71.027045 | 1.4712 | 0.0207 | 71.9309 | 71.8208 | 69.3294 | 71.027 | 0.39282 | |
| G6P-F6P | C00085 | I3sg5 | C13-06 | 6 | 776068.9251 | 614483.6046 | 628354.3173 | 74.27968 | 5.7009 | 0.0767 | 73.4131 | 69.0617 | 80.3642 | 74.2797 | 0.39282 | |
| GDP | C00035 | NT2 | C12 PARENT | 10 | 403325.4259 | 356579.3862 | 344821.9468 | 11.244208 | 0.9128 | 0.0812 | 10.434 | 11.0655 | 12.2331 | 11.2442 | 0.01245 | * |
| GDP | C00035 | I3sg5 | C12 PARENT | 10 | 260272.3289 | 282199.9577 | 190050.2348 | 14.319936 | 0.8294 | 0.0579 | 15.1521 | 14.3143 | 13.4934 | 14.3199 | 0.01245 | * |
| GDP | C00035 | NT2 | C13-04 | 10 | 19654.02894 | 0 | 3574.621963 | 0.211755 | 0.2647 | 1.2498 | 0.50845 | 0 | 0.12682 | 0.21175 | 0.51308 | |
| GDP | C00035 | I3sg5 | C13-04 | 10 | 0 | 46441.29639 | 0 | 0.7852307 | 1.3601 | 1.7321 | 0 | 2.35569 | 0 | 0.78523 | 0.51308 | |
| GDP | C00035 | NT2 | C13-05 | 10 | 3092739.198 | 2574081.249 | 2314546.143 | 80.66703 | 1.2533 | 0.0155 | 80.0091 | 79.8797 | 82.1123 | 80.667 | 0.62505 | |
| GDP | C00035 | I3sg5 | C13-05 | 10 | 1367731.204 | 1607784.203 | 1163802.189 | 81.268814 | 1.5223 | 0.0187 | 79.6243 | 81.5534 | 82.6288 | 81.2688 | 0.62505 | |
| GDP | C00035 | NT2 | C13-06 | 10 | 277388.1696 | 248594.6568 | 134927.1665 | 6.5590853 | 1.5583 | 0.2376 | 7.17603 | 7.71447 | 4.78676 | 6.55909 | 0.06593 | |
| GDP | C00035 | I3sg5 | C13-06 | 10 | 80301.0818 | 35024.73988 | 54086.98413 | 3.4305176 | 1.4919 | 0.4349 | 4.67483 | 1.7766 | 3.84012 | 3.43052 | 0.06593 | |
| GDP | C00035 | NT2 | C13-07 | 10 | 72376.30116 | 43191.65725 | 20888.53178 | 1.3179219 | 0.566 | 0.4295 | 1.87237 | 1.34034 | 0.74105 | 1.31792 | 0.03915 | * |
| GDP | C00035 | I3sg5 | C13-07 | 10 | 9426.909701 | 0 | 531.0704986 | 0.1955018 | 0.3065 | 1.568 | 0.5488 | 0 | 0.03771 | 0.1955 | 0.03915 | * |
| Glc | C00031 | NT2 | C12 PARENT | 6 | 260594.2228 | 206230.4152 | 336148.715 | 1.3297531 | 0.1559 | 0.1172 | 1.49451 | 1.18465 | 1.3101 | 1.32975 | 0.22536 | |

| | | | | | | | | | | | | | | | | |
|---|---|---|---|---|---|---|---|---|---|---|---|---|---|---|---|---|
| Glc | C00031 | I3sg5 | C12 PARENT | 6 | 215678.6398 | 313356.2146 | 318272.3475 | 1.5646368 | 0.2375 | 0.1518 | 1.29042 | 1.7004 | 1.70309 | 1.56464 | 0.22536 | |
| Glc | C00031 | NT2 | C13-05 | 6 | 288190.4801 | 259402.1027 | 523452.4569 | 1.7276505 | 0.2825 | 0.1635 | 1.65278 | 1.49009 | 2.04009 | 1.72765 | 0.77255 | |
| Glc | C00031 | I3sg5 | C13-05 | 6 | 270119.681 | 360728.6335 | 333797.8858 | 1.7865915 | 0.1707 | 0.0955 | 1.61614 | 1.95747 | 1.78617 | 1.78659 | 0.77255 | |
| Glc | C00031 | NT2 | C13-06 | 6 | 16887968.5 | 16942910.26 | 24798694.84 | 96.942596 | 0.3466 | 0.0036 | 96.8527 | 97.3253 | 96.6498 | 96.9426 | 0.38718 | |
| Glc | C00031 | I3sg5 | C13-06 | 6 | 16228092.67 | 17754265.15 | 18035844.2 | 96.648772 | 0.3942 | 0.0041 | 97.0934 | 96.3421 | 96.5107 | 96.6488 | 0.38718 | |
| GlcNAc-6P | C00357 | NT2 | C12 PARENT | 8 | 211322.9568 | 118462.8045 | 156081.0557 | 31.221382 | 7.2932 | 0.2336 | 35.0747 | 22.8098 | 35.7796 | 31.2214 | 0.5976 | |
| GlcNAc-6P | C00357 | I3sg5 | C12 PARENT | 8 | 114714.0613 | 83456.25815 | 44831.76296 | 27.312749 | 9.3075 | 0.3408 | 35.0563 | 29.8953 | 16.9866 | 27.3127 | 0.5976 | |
| GlcNAc-6P | C00357 | NT2 | C13-06 | 8 | 141043.8294 | 149117.4483 | 138580.7852 | 27.963407 | 4.229 | 0.1512 | 23.41 | 28.7123 | 31.7679 | 27.9634 | 0.35457 | |
| GlcNAc-6P | C00357 | I3sg5 | C13-06 | 8 | 72260.11636 | 120736.634 | 108089.8745 | 35.42906 | 11.6153 | 0.3278 | 22.0825 | 43.2497 | 40.955 | 35.4291 | 0.35457 | |
| GlcNAc-6P | C00357 | NT2 | C13-08 | 8 | 250126.9339 | 251770.0788 | 141567.023 | 40.815211 | 8.0356 | 0.1969 | 41.5153 | 48.4779 | 32.4525 | 40.8152 | 0.63687 | |
| GlcNAc-6P | C00357 | I3sg5 | C13-08 | 8 | 140253.9239 | 74968.84351 | 111002.0016 | 37.258191 | 9.0184 | 0.2421 | 42.8612 | 26.855 | 42.0584 | 37.2582 | 0.63687 | |
| Gln | C00064 | NT2 | C12 PARENT | 5 | 713428254.4 | 756114748.8 | 763240843.5 | 99.56109 | 0.0874 | 0.0009 | 99.5161 | 99.5053 | 99.6618 | 99.5611 | 0.83951 | |
| Gln | C00064 | I3sg5 | C12 PARENT | 5 | 512958964.5 | 641060554.7 | 431880066.1 | 99.577498 | 0.0983 | 0.001 | 99.6695 | 99.4739 | 99.5891 | 99.5775 | 0.83951 | |
| Gln | C00064 | NT2 | C13-01 | 5 | 270255.9304 | 153316.856 | 237477.5036 | 0.0296279 | 0.0088 | 0.2984 | 0.0377 | 0.02018 | 0.03101 | 0.02963 | 0.70901 | |
| Gln | C00064 | I3sg5 | C13-01 | 5 | 0 | 959441.5284 | 0 | 0.0496258 | 0.086 | 1.7321 | 0 | 0.14888 | 0 | 0.04963 | 0.70901 | |
| Gln | C00064 | NT2 | C13-02 | 5 | 0 | 0 | 0 | 0 | 0 | NA | 0 | 0 | 0 | 0 | NA | |
| Gln | C00064 | I3sg5 | C13-02 | 5 | 0 | 0 | 0 | 0 | 0 | NA | 0 | 0 | 0 | 0 | NA | |
| Gln | C00064 | NT2 | C13-03 | 5 | 811779.0268 | 1231560.505 | 565771.35 | 0.1163954 | 0.0442 | 0.3796 | 0.11324 | 0.16207 | 0.07388 | 0.1164 | 0.34867 | |
| Gln | C00064 | I3sg5 | C13-03 | 5 | 217910.0287 | 669499.5145 | 433724.315 | 0.0820806 | 0.0345 | 0.42 | 0.04234 | 0.10389 | 0.10001 | 0.08208 | 0.34867 | |
| Gln | C00064 | NT2 | C13-04 | 5 | 1439610.478 | 1293671.176 | 1281589.185 | 0.1794686 | 0.0185 | 0.1033 | 0.20081 | 0.17025 | 0.16735 | 0.17947 | 0.34218 | |
| Gln | C00064 | I3sg5 | C13-04 | 5 | 832122.4791 | 1060110.189 | 759698.5438 | 0.1671214 | 0.0071 | 0.0426 | 0.16168 | 0.1645 | 0.17518 | 0.16712 | 0.34218 | |
| Gln | C00064 | NT2 | C13-05 | 5 | 947290.0144 | 1080191.547 | 505158.9378 | 0.1134179 | 0.0414 | 0.365 | 0.13214 | 0.14215 | 0.06596 | 0.11342 | 0.70449 | |
| Gln | C00064 | I3sg5 | C13-05 | 5 | 651094.4127 | 701392.1853 | 588383.432 | 0.1236744 | 0.0136 | 0.1103 | 0.12651 | 0.10884 | 0.13568 | 0.12367 | 0.70449 | |
| Glu | C00025 | NT2 | C12 PARENT | 5 | 339456146.7 | 269539866.9 | 293720932.9 | 63.037251 | 1.1437 | 0.0181 | 62.4004 | 62.3537 | 64.3576 | 63.0373 | 0.03388 | * |
| Glu | C00025 | I3sg5 | C12 PARENT | 5 | 237804638.8 | 293724588.1 | 199666638 | 65.31507 | 0.4915 | 0.0075 | 65.6468 | 65.548 | 64.7505 | 65.3151 | 0.03388 | * |
| Glu | C00025 | NT2 | C13-01 | 5 | 6752858.665 | 6630254.264 | 5175992.006 | 1.303088 | 0.2069 | 0.1588 | 1.24134 | 1.5338 | 1.13412 | 1.30309 | 0.29542 | |
| Glu | C00025 | I3sg5 | C13-01 | 5 | 4106498.242 | 5179008.983 | 3655794.465 | 1.1583051 | 0.0261 | 0.0225 | 1.13361 | 1.15575 | 1.18555 | 1.15831 | 0.29542 | |
| Glu | C00025 | NT2 | C13-02 | 5 | 100206934 | 79682977.75 | 82003006.76 | 18.273894 | 0.2652 | 0.0145 | 18.4205 | 18.4334 | 17.9678 | 18.2739 | 0.03181 | * |
| Glu | C00025 | I3sg5 | C13-02 | 5 | 61407043.9 | 77702191.11 | 54820312.07 | 17.356528 | 0.4133 | 0.0238 | 16.9516 | 17.3401 | 17.7778 | 17.3565 | 0.03181 | * |
| Glu | C00025 | NT2 | C13-03 | 5 | 29014394.05 | 22574821.28 | 22346648.23 | 5.1507634 | 0.2272 | 0.0441 | 5.33356 | 5.22232 | 4.89641 | 5.15076 | 0.08346 | |
| Glu | C00025 | I3sg5 | C13-03 | 5 | 17407610.16 | 21610755.91 | 15108631.77 | 4.8425787 | 0.0501 | 0.0104 | 4.80543 | 4.82269 | 4.89962 | 4.84258 | 0.08346 | |
| Glu | C00025 | NT2 | C13-04 | 5 | 41878392.24 | 33331254.67 | 32656732.39 | 7.5214654 | 0.317 | 0.0421 | 7.69828 | 7.71065 | 7.15547 | 7.52147 | 0.02545 | * |
| Glu | C00025 | I3sg5 | C13-04 | 5 | 25004303.75 | 29553089.69 | 21245779.16 | 6.795828 | 0.1739 | 0.0256 | 6.90252 | 6.59511 | 6.88985 | 6.79583 | 0.02545 | * |
| Glu | C00025 | NT2 | C13-05 | 5 | 26688235.97 | 20516153.89 | 20485352.58 | 4.713538 | 0.2106 | 0.0447 | 4.90595 | 4.74608 | 4.48858 | 4.71354 | 0.21335 | |
| Glu | C00025 | I3sg5 | C13-05 | 5 | 16518655.96 | 20336669.19 | 13866114.31 | 4.5316903 | 0.0322 | 0.0071 | 4.56003 | 4.53836 | 4.49668 | 4.53169 | 0.21335 | |
| GlucA | C00191 | NT2 | C12 PARENT | 6 | 9660.636463 | 9622.806175 | 0 | 12.192148 | 10.5935 | 0.8689 | 19.1457 | 17.4307 | 0 | 12.1921 | 0.81347 | |
| GlucA | C00191 | I3sg5 | C12 PARENT | 6 | 0 | 17496.80148 | 0 | 9.3720597 | 16.2329 | 1.7321 | 0 | 28.1162 | 0 | 9.37206 | 0.81347 | |
| GlucA | C00191 | NT2 | C13-06 | 6 | 40797.8572 | 45583.16345 | 58997.95886 | 87.807852 | 10.5935 | 0.1206 | 80.8543 | 82.5693 | 100 | 87.8079 | 0.81347 | |
| GlucA | C00191 | I3sg5 | C13-06 | 6 | 20219.10337 | 44733.56597 | 17435.57127 | 90.62794 | 16.2329 | 0.1791 | 100 | 71.8838 | 100 | 90.6279 | 0.81347 | |
| Gly | C00037 | NT2 | C12 PARENT | 2 | 14284660.67 | 13306448.2 | 12722724.82 | 94.468679 | 0.3734 | 0.004 | 94.0664 | 94.8043 | 94.5353 | 94.4687 | 0.13508 | |
| Gly | C00037 | I3sg5 | C12 PARENT | 2 | 8011783.522 | 10514590.14 | 8279618.795 | 95.142253 | 0.5004 | 0.0053 | 95.7201 | 94.8533 | 94.8533 | 95.1423 | 0.13508 | |
| Gly | C00037 | NT2 | C13-01 | 2 | 0 | 0 | 0 | 0 | 0 | NA | 0 | 0 | 0 | 0 | NA | |
| Gly | C00037 | I3sg5 | C13-01 | 2 | 0 | 0 | 0 | 0 | 0 | NA | 0 | 0 | 0 | 0 | NA | |
| Gly | C00037 | NT2 | C13-02 | 2 | 901059.5874 | 729252.4625 | 735444.1267 | 5.5313206 | 0.3734 | 0.0675 | 5.9336 | 5.1957 | 5.46467 | 5.53132 | 0.13508 | |
| Gly | C00037 | I3sg5 | C13-02 | 2 | 358226.4102 | 570513.5886 | 449247.8433 | 4.8577469 | 0.5004 | 0.103 | 4.27988 | 5.14667 | 5.14669 | 4.85775 | 0.13508 | |
| Glycerol-3P | C00093 | NT2 | C12 PARENT | 3 | 563828.2995 | 433593.3172 | 998476.5108 | 37.897788 | 2.2544 | 0.0595 | 38.4718 | 35.4118 | 39.8098 | 37.8978 | 0.64117 | |
| Glycerol-3P | C00093 | I3sg5 | C12 PARENT | 3 | 1031047.71 | 920848.0701 | 661340.7024 | 37.035546 | 1.9289 | 0.0521 | 35.8801 | 35.9642 | 39.2623 | 37.0355 | 0.64117 | |
| Glycerol-3P | C00093 | NT2 | C13-03 | 3 | 901735.8594 | 790836.8757 | 1509643.307 | 62.102212 | 2.2544 | 0.0363 | 61.5282 | 64.5882 | 60.1902 | 62.1022 | 0.64117 | |
| Glycerol-3P | C00093 | I3sg5 | C13-03 | 3 | 1842543.435 | 1639609.45 | 1023074.799 | 62.964454 | 1.9289 | 0.0306 | 64.1199 | 64.0358 | 60.7377 | 62.9645 | 0.64117 | |
| GMP | C00144 | NT2 | C12 PARENT | 10 | 136007.471 | 196444.1846 | 202531.6769 | 11.24924 | 2.402 | 0.2135 | 8.47565 | 12.6362 | 12.6359 | 11.2492 | 0.60084 | |
| GMP | C00144 | I3sg5 | C12 PARENT | 10 | 114356.4045 | 198696.0546 | 51610.9794 | 13.263334 | 5.6609 | 0.4268 | 16.6282 | 16.4341 | 6.72769 | 13.2633 | 0.60084 | |
| GMP | C00144 | NT2 | C13-04 | 10 | 2146.260335 | 52128.29197 | 58173.39864 | 2.3720995 | 1.9434 | 0.8193 | 0.13375 | 3.35312 | 3.62943 | 2.3721 | 0.40856 | |
| GMP | C00144 | I3sg5 | C13-04 | 10 | 18148.39716 | 8204.809271 | 0 | 1.1058404 | 1.3703 | 1.2392 | 2.6389 | 0.67862 | 0 | 1.10584 | 0.40856 | |
| GMP | C00144 | NT2 | C13-05 | 10 | 1331488.97 | 1233788.719 | 1263244.072 | 80.383797 | 2.2609 | 0.0281 | 82.9751 | 79.3628 | 78.8135 | 80.3838 | 0.32645 | |
| GMP | C00144 | I3sg5 | C13-05 | 10 | 549667.5645 | 995207.0608 | 691368.4922 | 84.120446 | 5.3333 | 0.0634 | 79.9255 | 82.3133 | 90.1226 | 84.1204 | 0.32645 | |
| GMP | C00144 | NT2 | C13-06 | 10 | 133722.0733 | 60691.0127 | 78877.23613 | 5.719426 | 2.3201 | 0.4056 | 8.33323 | 3.90392 | 4.92113 | 5.71943 | 0.05398 | |
| GMP | C00144 | I3sg5 | C13-06 | 10 | 2865.225468 | 6940.362344 | 24163.02807 | 1.3801345 | 1.5345 | 1.1119 | 0.41662 | 0.57404 | 3.14974 | 1.38013 | 0.05398 | |
| GMP | C00144 | NT2 | C13-07 | 10 | 1320.239703 | 11566.99131 | 0 | 0.2754381 | 0.4079 | 1.4809 | 0.08227 | 0.74404 | 0 | 0.27544 | 0.61816 | |
| GMP | C00144 | I3sg5 | C13-07 | 10 | 2687.180592 | 0 | 0 | 0.130245 | 0.2256 | 1.7321 | 0.39073 | 0 | 0 | 0.13024 | 0.61816 | |
| GSH | C00051 | NT2 | C12 PARENT | 10 | 77573593.43 | 73623339.68 | 66116659.54 | 69.359623 | 1.511 | 0.0218 | 68.2856 | 68.7058 | 71.0875 | 69.3596 | 0.08177 | |
| GSH | C00051 | I3sg5 | C12 PARENT | 10 | 49241308.67 | 55333446.54 | 39496041.63 | 72.510974 | 1.8127 | 0.025 | 73.2109 | 73.8694 | 70.4526 | 72.511 | 0.08177 | |
| GSH | C00051 | NT2 | C13-01 | 10 | 0 | 0 | 0 | 0 | 0 | NA | 0 | 0 | 0 | 0 | NA | |
| GSH | C00051 | I3sg5 | C13-01 | 10 | 0 | 0 | 0 | 0 | 0 | NA | 0 | 0 | 0 | 0 | NA | |
| GSH | C00051 | NT2 | C13-02 | 10 | 21518585.86 | 20392768.7 | 16543176.99 | 18.58658 | 0.6939 | 0.0373 | 18.9421 | 19.0307 | 17.7869 | 18.5866 | 0.09115 | |
| GSH | C00051 | I3sg5 | C13-02 | 10 | 10901076.66 | 11521551.65 | 10165995.05 | 16.574197 | 1.4126 | 0.0852 | 16.2075 | 15.3811 | 18.134 | 16.5742 | 0.09115 | |
| GSH | C00051 | NT2 | C13-03 | 10 | 3989767.426 | 3549190.253 | 2914544.402 | 3.3192875 | 0.1893 | 0.057 | 3.51207 | 3.31213 | 3.13367 | 3.31929 | 0.31993 | |
| GSH | C00051 | I3sg5 | C13-03 | 10 | 1936476.487 | 2352681.765 | 1875929.589 | 3.1220575 | 0.2341 | 0.075 | 2.87911 | 3.1408 | 3.34626 | 3.12206 | 0.31993 | |
| GSH | C00051 | NT2 | C13-04 | 10 | 6736884.181 | 6350297.732 | 4935399.907 | 5.7209546 | 0.359 | 0.0627 | 5.93027 | 5.92614 | 5.30646 | 5.72095 | 0.04936 | * |
| GSH | C00051 | I3sg5 | C13-04 | 10 | 3405504.11 | 3665331.43 | 2948452.953 | 5.0719386 | 0.1833 | 0.0361 | 5.06323 | 4.89317 | 5.25942 | 5.07194 | 0.04936 | * |
| GSH | C00051 | NT2 | C13-05 | 10 | 3624723.988 | 3108408.298 | 2404082.336 | 2.8921152 | 0.303 | 0.1048 | 3.19073 | 2.90079 | 2.58483 | 2.89212 | 0.28668 | |
| GSH | C00051 | I3sg5 | C13-05 | 10 | 1761086.666 | 2006247.568 | 1527663.37 | 2.6738961 | 0.0535 | 0.02 | 2.61834 | 2.67831 | 2.72503 | 2.6739 | 0.28668 | |
| GSH | C00051 | NT2 | C13-06 | 10 | 47707.38398 | 66035.78698 | 28660.09879 | 0.0448117 | 0.0156 | 0.3481 | 0.042 | 0.06163 | 0.03081 | 0.04481 | 0.23723 | |
| GSH | C00051 | I3sg5 | C13-06 | 10 | 0 | 0 | 29416.15615 | 0.0174907 | 0.0303 | 1.7321 | 0 | 0 | 0.05247 | 0.01749 | 0.23723 | |
| GSH | C00051 | NT2 | C13-07 | 10 | 110406.3657 | 67366.39309 | 64948.34011 | 0.0766285 | 0.0181 | 0.2368 | 0.09719 | 0.06287 | 0.06983 | 0.07663 | 0.0147 | * |
| GSH | C00051 | I3sg5 | C13-07 | 10 | 14115.7394 | 27831.96059 | 16927.12092 | 0.0294456 | 0.0081 | 0.2754 | 0.02099 | 0.03716 | 0.03019 | 0.02945 | 0.0147 | * |
| GSSG | C00127 | NT2 | C12 PARENT | 20 | 4630265.064 | 5439227.35 | 5617878.515 | 52.490691 | 2.1404 | 0.0408 | 51.8869 | 50.7171 | 54.8681 | 52.4907 | 0.06379 | |
| GSSG | C00127 | I3sg5 | C12 PARENT | 20 | 3667991.205 | 3922287.378 | 3160731.278 | 56.382241 | 1.5637 | 0.0277 | 57.162 | 57.4027 | 54.582 | 56.3822 | 0.06379 | |
| GSSG | C00127 | NT2 | C13-01 | 20 | 71714.3864 | 107605.8502 | 81413.65656 | 0.8673756 | 0.1178 | 0.1359 | 0.80363 | 1.00335 | 0.79514 | 0.86738 | 0.01739 | * |
| GSSG | C00127 | I3sg5 | C13-01 | 20 | 34322.8142 | 43532.95645 | 29369.76667 | 0.5597242 | 0.0684 | 0.1223 | 0.53489 | 0.63711 | 0.50718 | 0.55972 | 0.01739 | * |
| GSSG | C00127 | NT2 | C13-02 | 20 | 2048006.844 | 2441747.538 | 2275978.07 | 22.648812 | 0.375 | 0.0166 | 22.95 | 22.7676 | 22.2288 | 22.6488 | 0.10462 | |
| GSSG | C00127 | I3sg5 | C13-02 | 20 | 1349456.787 | 1413164.222 | 1302598.526 | 21.401973 | 0.9619 | 0.0449 | 21.0299 | 20.6817 | 22.4943 | 21.402 | 0.10462 | |
| GSSG | C00127 | NT2 | C13-03 | 20 | 480748.0328 | 630738.511 | 534220.9437 | 5.4953517 | 0.3448 | 0.0627 | 5.38728 | 5.8812 | 5.21757 | 5.49535 | 0.10814 | |
| GSSG | C00127 | I3sg5 | C13-03 | 20 | 312540.6904 | 325472.0266 | 305801.7762 | 4.9715832 | 0.2731 | 0.0549 | 4.87063 | 4.76329 | 5.28083 | 4.97158 | 0.10814 | |
| GSSG | C00127 | NT2 | C13-04 | 20 | 848985.0778 | 1065726.469 | 933571.9541 | 9.522945 | 0.4097 | 0.043 | 9.51375 | 9.93717 | 9.11791 | 9.52295 | 0.19359 | |
| GSSG | C00127 | I3sg5 | C13-04 | 20 | 533468.3542 | 630050.0168 | 534527.4878 | 8.9216688 | 0.5267 | 0.059 | 8.31357 | 9.22079 | 9.23065 | 8.92167 | 0.19359 | |
| GSSG | C00127 | NT2 | C13-05 | 20 | 537652.2397 | 680177.2335 | 536012.4032 | 5.8674014 | 0.5701 | 0.0972 | 6.02495 | 6.34219 | 5.23507 | 5.8674 | 0.11566 | |
| GSSG | C00127 | I3sg5 | C13-05 | 20 | 348928.524 | 346466.0184 | 283379.6007 | 5.1339537 | 0.2775 | 0.0541 | 5.4377 | 5.07053 | 4.89362 | 5.13395 | 0.11566 | |
| GSSG | C00127 | NT2 | C13-06 | 20 | 169680.7588 | 165903.8494 | 94020.40672 | 1.4555521 | 0.4979 | 0.3421 | 1.90145 | 1.54694 | 0.91827 | 1.45555 | 0.51078 | |
| GSSG | C00127 | I3sg5 | C13-06 | 20 | 76768.34163 | 87669.63051 | 73128.23849 | 1.2474135 | 0.0454 | 0.0364 | 1.19636 | 1.28305 | 1.26284 | 1.24741 | 0.51078 | |
| GSSG | C00127 | NT2 | C13-07 | 20 | 98987.12936 | 111821.784 | 153602.5063 | 1.2173678 | 0.2472 | 0.203 | 1.10925 | 1.04266 | 1.50019 | 1.21737 | 0.34128 | |
| GSSG | C00127 | I3sg5 | C13-07 | 20 | 62763.51731 | 59640.25861 | 71074.2537 | 1.026102 | 0.1821 | 0.1774 | 0.97811 | 0.87283 | 1.22737 | 1.0261 | 0.34128 | |
| GSSG | C00127 | NT2 | C13-08 | 20 | 23628.96051 | 41471.1727 | 12178.44122 | 0.2568067 | 0.1341 | 0.522 | 0.26479 | 0.38669 | 0.11894 | 0.25681 | 0.99756 | |
| GSSG | C00127 | I3sg5 | C13-08 | 20 | 23530.52263 | 0 | 23463.67042 | 0.2572963 | 0.2237 | 0.8692 | 0.3667 | 0 | 0.40519 | 0.2573 | 0.99756 | |
| GSSG | C00127 | NT2 | C13-09 | 20 | 14096.29774 | 40231.18424 | 0 | 0.1776972 | 0.1883 | 1.0599 | 0.15796 | 0.37513 | 0 | 0.1777 | 0.5083 | |
| GSSG | C00127 | I3sg5 | C13-09 | 20 | 7066.722509 | 4646.592364 | 6717.543052 | 0.0980449 | 0.0262 | 0.267 | 0.11013 | 0.068 | 0.116 | 0.09804 | 0.5083 | |
| GTP | C00044 | NT2 | C12 PARENT | 10 | 3976994.639 | 3429731.912 | 4203467.33 | 11.210515 | 0.4873 | 0.0435 | 10.7277 | 11.7021 | 11.2017 | 11.2105 | 0.07306 | |
| GTP | C00044 | I3sg5 | C12 PARENT | 10 | 3096735.085 | 3482070.711 | 1870059.122 | 13.352284 | 1.456 | 0.109 | 13.2008 | 14.8781 | 11.978 | 13.3523 | 0.07306 | |
| GTP | C00044 | NT2 | C13-01 | 10 | 0 | 0 | 0 | 0 | 0 | NA | 0 | 0 | 0 | 0 | NA | |
| GTP | C00044 | I3sg5 | C13-01 | 10 | 0 | 0 | 0 | 0 | 0 | NA | 0 | 0 | 0 | 0 | NA | |

| | | | | | | | | | | | | | | | | |
|---|---|---|---|---|---|---|---|---|---|---|---|---|---|---|---|---|
| GTP | C00044 | NT2 | C13-02 | 10 | 249028.8823 | 180283.3377 | 172241.6384 | 0.5819543 | 0.1102 | 0.1893 | 0.67174 | 0.61512 | 0.459 | 0.58195 | 0.37272 | |
| GTP | C00044 | I3sg5 | C13-02 | 10 | 40520.66532 | 93776.67252 | 107146.6205 | 0.4199026 | 0.2573 | 0.6128 | 0.17273 | 0.40069 | 0.68629 | 0.4199 | 0.37272 | |
| GTP | C00044 | NT2 | C13-03 | 10 | 201523.3889 | 258187.7229 | 377216.0705 | 0.8099198 | 0.2389 | 0.2949 | 0.5436 | 0.88093 | 1.00524 | 0.80992 | 0.27494 | |
| GTP | C00044 | I3sg5 | C13-03 | 10 | 293425.4572 | 352098.1005 | 110481.4694 | 1.1542992 | 0.4071 | 0.3527 | 1.25081 | 1.50443 | 0.70765 | 1.1543 | 0.27494 | |
| GTP | C00044 | NT2 | C13-04 | 10 | 770490.5615 | 671407.4617 | 469377.2087 | 1.8733336 | 0.5495 | 0.2933 | 2.07835 | 2.29082 | 1.25083 | 1.87333 | 0.0588 | |
| GTP | C00044 | I3sg5 | C13-04 | 10 | 163713.9135 | 302017.5293 | 62944.06035 | 0.7971655 | 0.4519 | 0.5669 | 0.69788 | 1.29045 | 0.40317 | 0.79717 | 0.0588 | |
| GTP | C00044 | NT2 | C13-05 | 10 | 27998367.24 | 22447335.72 | 29312290.99 | 76.742347 | 1.3017 | 0.017 | 75.5238 | 76.5895 | 78.1138 | 76.7423 | 0.429 | |
| GTP | C00044 | I3sg5 | C13-05 | 10 | 18711892.22 | 17744742.38 | 12219197.13 | 77.950004 | 1.9918 | 0.0256 | 79.765 | 75.8192 | 78.2658 | 77.95 | 0.429 | |
| GTP | C00044 | NT2 | C13-06 | 10 | 2976464.199 | 1789908.797 | 2136438.595 | 6.6097592 | 1.2462 | 0.1885 | 8.02882 | 6.1071 | 5.69335 | 6.60976 | 0.217 | |
| GTP | C00044 | I3sg5 | C13-06 | 10 | 747470.535 | 977553.8293 | 1059691.07 | 4.7168866 | 1.8603 | 0.3944 | 3.18632 | 4.17686 | 6.78748 | 4.71689 | 0.217 | |
| GTP | C00044 | NT2 | C13-07 | 10 | 890476.6279 | 531776.0261 | 790928.7718 | 2.1080448 | 0.2938 | 0.1394 | 2.402 | 1.8144 | 2.10773 | 2.10804 | 0.15306 | |
| GTP | C00044 | I3sg5 | C13-07 | 10 | 405007.5764 | 451775.6377 | 182911.5156 | 1.6094581 | 0.3927 | 0.244 | 1.72647 | 1.93033 | 1.17158 | 1.60946 | 0.15306 | |
| GTP | C00044 | NT2 | C13-08 | 10 | 8907.160807 | 0 | 63174.49994 | 0.0641263 | 0.0911 | 1.42 | 0.02403 | 0 | 0.16835 | 0.06413 | 0.28955 | |
| GTP | C00044 | I3sg5 | C13-08 | 10 | 0 | 0 | 0 | 0 | 0 | NA | 0 | 0 | 0 | 0 | 0.28955 | |
| IMP | C00130 | NT2 | C12 PARENT | 10 | 59940.84125 | 98449.25236 | 75310.96383 | 17.40287 | 2.6025 | 0.1495 | 14.6039 | 19.7497 | 17.8551 | 17.4029 | 0.44135 | |
| IMP | C00130 | I3sg5 | C12 PARENT | 10 | 38253.50786 | 108771.741 | 22277.88623 | 13.740766 | 6.9587 | 0.5064 | 10.3847 | 21.7415 | 9.09613 | 13.7408 | 0.44135 | |
| IMP | C00130 | NT2 | C13-05 | 10 | 350504.5348 | 400036.7579 | 346478.5487 | 82.59713 | 2.6025 | 0.0315 | 85.3961 | 80.2503 | 82.1449 | 82.5971 | 0.44135 | |
| IMP | C00130 | I3sg5 | C13-05 | 10 | 330110.9036 | 391524.4088 | 222638.0915 | 86.259234 | 6.9587 | 0.0807 | 89.6153 | 78.2585 | 90.9039 | 86.2592 | 0.44135 | |
| Lac | C00186 | NT2 | C12 PARENT | 3 | 2282324.339 | 1544608.714 | 1982256.606 | 19.378653 | 2.8295 | 0.146 | 22.3165 | 19.1478 | 16.6716 | 19.3787 | 0.96021 | |
| Lac | C00186 | I3sg5 | C12 PARENT | 3 | 1214854.225 | 1982800.644 | 1114513.847 | 19.260836 | 2.6016 | 0.1351 | 20.2828 | 21.1963 | 16.3034 | 19.2608 | 0.96021 | |
| Lac | C00186 | NT2 | C13-02 | 3 | 92415.53561 | 42155.69952 | 204090.1093 | 1.0475686 | 0.6098 | 0.5821 | 0.90364 | 0.52259 | 1.71648 | 1.04757 | 0.37858 | |
| Lac | C00186 | I3sg5 | C13-02 | 3 | 0 | 131379.2728 | 0 | 0.4681523 | 0.8109 | 1.7321 | 0 | 1.40446 | 0 | 0.46815 | 0.37858 | |
| Lac | C00186 | NT2 | C13-03 | 3 | 7852324.683 | 6479987.999 | 9703670.05 | 79.573779 | 2.5031 | 0.0315 | 76.7798 | 80.3296 | 81.6119 | 79.5738 | 0.78046 | |
| Lac | C00186 | I3sg5 | C13-03 | 3 | 4774734.42 | 7240274.613 | 5721561.673 | 80.271012 | 3.185 | 0.0397 | 79.7172 | 77.3992 | 83.6966 | 80.271 | 0.78046 | |
| Mal | C00149 | NT2 | C12 PARENT | 4 | 46151175.87 | 44124817.66 | 39924937.9 | 65.385934 | 0.9308 | 0.0142 | 64.3213 | 65.7906 | 66.0459 | 65.3859 | 0.5587 | |
| Mal | C00149 | I3sg5 | C12 PARENT | 4 | 30295339.12 | 31169693.36 | 24975234.64 | 65.887955 | 0.9982 | 0.0151 | 66.871 | 65.9176 | 64.8753 | 65.888 | 0.5587 | |
| Mal | C00149 | NT2 | C13-01 | 4 | 1594239.12 | 1207377.697 | 1168132.937 | 1.9848349 | 0.2157 | 0.1087 | 2.22191 | 1.80021 | 1.93238 | 1.98483 | 0.98382 | |
| Mal | C00149 | I3sg5 | C13-01 | 4 | 717947.0608 | 1071263.992 | 804131.5617 | 1.9796776 | 0.3533 | 0.1784 | 1.58473 | 2.26551 | 2.0888 | 1.97968 | 0.98382 | |
| Mal | C00149 | NT2 | C13-02 | 4 | 11441310.16 | 10336683.72 | 8998244.963 | 15.414443 | 0.5303 | 0.0344 | 15.9459 | 15.4121 | 14.8854 | 15.4144 | 0.60997 | |
| Mal | C00149 | I3sg5 | C13-02 | 4 | 6315938.727 | 6863661.534 | 6327744.235 | 14.964432 | 1.3071 | 0.0873 | 13.9412 | 14.5153 | 16.4369 | 14.9644 | 0.60997 | |
| Mal | C00149 | NT2 | C13-03 | 4 | 8782247.813 | 8026344.563 | 7261689.461 | 12.07331 | 0.146 | 0.0121 | 12.2399 | 11.9674 | 12.0127 | 12.0733 | 0.54879 | |
| Mal | C00149 | I3sg5 | C13-03 | 4 | 5538524.542 | 5770557.647 | 4621946.867 | 12.144887 | 0.1209 | 0.01 | 12.2252 | 12.2036 | 12.0059 | 12.1449 | 0.54879 | |
| Mal | C00149 | NT2 | C13-04 | 4 | 3782022.227 | 3373308.038 | 3097324.886 | 5.1414778 | 0.1217 | 0.0237 | 5.27104 | 5.02964 | 5.12375 | 5.14148 | 0.64771 | |
| Mal | C00149 | I3sg5 | C13-04 | 4 | 2436401.154 | 2410680.195 | 1768246.796 | 5.0230489 | 0.3977 | 0.0792 | 5.37788 | 5.0981 | 4.59317 | 5.02305 | 0.64771 | |
| Met | C00073 | NT2 | C12 PARENT | 5 | 33880300.87 | 34575815.07 | 24790073.89 | 99.066426 | 0.309 | 0.0031 | 98.9959 | 98.7988 | 99.4046 | 99.0664 | 0.72249 | |
| Met | C00073 | I3sg5 | C12 PARENT | 5 | 28414811.23 | 25204192.14 | 30656477.28 | 99.156643 | 0.2695 | 0.0027 | 99.4608 | 99.0616 | 98.9475 | 99.1566 | 0.72249 | |
| Met | C00073 | NT2 | C13-01 | 5 | 0 | 0 | 0 | 0 | 0 | NA | 0 | 0 | 0 | 0 | NA | |
| Met | C00073 | I3sg5 | C13-01 | 5 | 0 | 0 | 0 | 0 | 0 | NA | 0 | 0 | 0 | 0 | NA | |
| Met | C00073 | NT2 | C13-04 | 5 | 343648.1611 | 420365.4189 | 148492.1942 | 0.9335744 | 0.309 | 0.331 | 1.00412 | 1.20118 | 0.59543 | 0.93357 | 0.72249 | |
| Met | C00073 | I3sg5 | C13-04 | 5 | 154040.9848 | 238759.1569 | 326081.5886 | 0.8433567 | 0.2695 | 0.3196 | 0.53919 | 0.93841 | 1.05247 | 0.84336 | 0.72249 | |
| MTA | C00170 | NT2 | C12 PARENT | 11 | 393772.183 | 234272.9379 | 185897.0001 | 18.892694 | 1.159 | 0.0613 | 19.7648 | 17.5775 | 19.3358 | 18.8927 | 0.34102 | |
| MTA | C00170 | I3sg5 | C12 PARENT | 11 | 104044.2899 | 287757.2968 | 169077.8313 | 20.21451 | 1.7756 | 0.0878 | 19.3889 | 22.2526 | 19.0021 | 20.2145 | 0.34102 | |
| MTA | C00170 | NT2 | C13-01 | 11 | 0 | 0 | 0 | 0 | 0 | NA | 0 | 0 | 0 | 0 | NA | |
| MTA | C00170 | I3sg5 | C13-01 | 11 | 0 | 0 | 0 | 0 | 0 | NA | 0 | 0 | 0 | 0 | NA | |
| MTA | C00170 | NT2 | C13-02 | 11 | 11691.12485 | 5861.809752 | 5143.356656 | 0.520536 | 0.0746 | 0.1432 | 0.58682 | 0.43981 | 0.53498 | 0.52054 | 0.00046 | *** |
| MTA | C00170 | I3sg5 | C13-02 | 11 | 0 | 772.832118 | 0 | 0.0199213 | 0.0345 | 1.7321 | 0 | 0.05976 | 0 | 0.01992 | 0.00046 | *** |
| MTA | C00170 | NT2 | C13-03 | 11 | 4386.901268 | 3531.827039 | 3210.637575 | 0.2730454 | 0.0573 | 0.2099 | 0.22019 | 0.26499 | 0.33395 | 0.27305 | 0.14715 | |
| MTA | C00170 | I3sg5 | C13-03 | 11 | 0 | 3649.61256 | 0 | 0.0940761 | 0.1629 | 1.7321 | 0 | 0.28223 | 0 | 0.09408 | 0.14715 | |
| MTA | C00170 | NT2 | C13-04 | 11 | 30730.38913 | 12785.20604 | 16616.83315 | 1.4100377 | 0.4013 | 0.2846 | 1.54247 | 0.95927 | 1.72837 | 1.41004 | 0.76659 | |
| MTA | C00170 | I3sg5 | C13-04 | 11 | 0 | 12045.30625 | 22724.30392 | 1.161794 | 1.2924 | 1.1124 | 0 | 0.93148 | 2.55391 | 1.16179 | 0.76659 | |
| MTA | C00170 | NT2 | C13-05 | 11 | 1442334.018 | 994644.5394 | 693105.4187 | 73.038737 | 1.3848 | 0.019 | 72.3959 | 74.6281 | 72.0922 | 73.0387 | 0.94985 | |
| MTA | C00170 | I3sg5 | C13-05 | 11 | 406241.1834 | 911696.7855 | 645685.6302 | 72.924264 | 2.6191 | 0.0359 | 75.704 | 70.5025 | 72.5664 | 72.9243 | 0.94985 | |
| MTA | C00170 | NT2 | C13-06 | 11 | 109372.1729 | 81704.82381 | 57442.27659 | 5.86495 | 0.3341 | 0.057 | 5.48978 | 6.13031 | 5.97476 | 5.86495 | 0.51429 | |
| MTA | C00170 | I3sg5 | C13-06 | 11 | 26332.68658 | 77219.86093 | 52298.56023 | 5.5854339 | 0.5893 | 0.1055 | 4.90716 | 5.97149 | 5.87765 | 5.58543 | 0.51429 | |
| NAD+ | C00003 | NT2 | C12 PARENT | 21 | 103488.936 | 114924.2041 | 87115.3312 | 3.1952744 | 0.1773 | 0.0555 | 3.0225 | 3.37678 | 3.18654 | 3.19527 | 0.64793 | |
| NAD+ | C00003 | I3sg5 | C12 PARENT | 21 | 70568.12653 | 101208.3373 | 47281.44665 | 3.0906975 | 0.3219 | 0.1042 | 2.77327 | 3.4169 | 3.08192 | 3.0907 | 0.64793 | |
| NAD+ | C00003 | NT2 | C13-04 | 21 | 42137.05214 | 28481.57464 | 14176.89116 | 0.8620298 | 0.3567 | 0.4138 | 1.23066 | 0.83686 | 0.51857 | 0.86203 | 0.72354 | |
| NAD+ | C00003 | I3sg5 | C13-04 | 21 | 23218.26753 | 31405.1793 | 3733.022587 | 0.7386861 | 0.4353 | 0.5893 | 0.91246 | 1.06027 | 0.24333 | 0.73869 | 0.72354 | |
| NAD+ | C00003 | NT2 | C13-05 | 21 | 949338.593 | 916750.0395 | 796259.7941 | 27.929637 | 1.1087 | 0.0397 | 27.7264 | 26.9366 | 29.1259 | 27.9296 | 0.64897 | |
| NAD+ | C00003 | I3sg5 | C13-05 | 21 | 741323.8656 | 885243.1422 | 407116.6651 | 28.519014 | 1.7574 | 0.0616 | 29.1334 | 29.8867 | 26.5369 | 28.519 | 0.64897 | |
| NAD+ | C00003 | NT2 | C13-06 | 21 | 0 | 0 | 0 | 0 | 0 | NA | 0 | 0 | 0 | 0 | 0.04129 | * |
| NAD+ | C00003 | I3sg5 | C13-06 | 21 | 5205.643024 | 8712.77012 | 9474.762046 | 0.3721062 | 0.2173 | 0.5839 | 0.20458 | 0.29415 | 0.61759 | 0.37211 | 0.04129 | * |
| NAD+ | C00003 | NT2 | C13-07 | 21 | 58604.99468 | 55662.28063 | 52495.328 | 1.7557741 | 0.1474 | 0.0839 | 1.71162 | 1.63551 | 1.9202 | 1.75577 | 0.009 | ** |
| NAD+ | C00003 | I3sg5 | C13-07 | 21 | 29225.46432 | 40391.10018 | 17146.38219 | 1.2099423 | 0.134 | 0.1108 | 1.14854 | 1.36365 | 1.11764 | 1.20994 | 0.009 | ** |
| NAD+ | C00003 | NT2 | C13-08 | 21 | 40863.6729 | 59975.28974 | 20808.9137 | 1.2389525 | 0.5021 | 0.4053 | 1.19347 | 1.76223 | 0.76116 | 1.23895 | 0.13175 | |
| NAD+ | C00003 | I3sg5 | C13-08 | 21 | 60808.08525 | 59131.05701 | 23221.86641 | 1.9665646 | 0.4388 | 0.2231 | 2.38971 | 1.99633 | 1.51366 | 1.96656 | 0.13175 | |
| NAD+ | C00003 | NT2 | C13-09 | 21 | 95583.94302 | 91652.57496 | 88554.77628 | 2.9079399 | 0.2911 | 0.1001 | 2.79163 | 2.693 | 3.23919 | 2.90794 | 0.1892 | |
| NAD+ | C00003 | I3sg5 | C13-09 | 21 | 46912.03448 | 83336.4089 | 38293.83693 | 2.3844058 | 0.4945 | 0.2074 | 1.8436 | 2.81352 | 2.49609 | 2.38441 | 0.1892 | |
| NAD+ | C00003 | NT2 | C13-10 | 21 | 1909805.045 | 1970349.524 | 1553471.102 | 56.83183 | 1.0582 | 0.0186 | 55.7778 | 57.8941 | 56.8235 | 56.8318 | 0.69283 | |
| NAD+ | C00003 | I3sg5 | C13-10 | 21 | 1470684.842 | 1637598.891 | 905327.2102 | 57.365089 | 1.8994 | 0.0331 | 57.7967 | 55.287 | 59.0115 | 57.3651 | 0.69283 | |
| NAD+ | C00003 | NT2 | C13-11 | 21 | 186479.41 | 141377.6359 | 101784.3817 | 4.4411631 | 0.8968 | 0.2019 | 5.44632 | 4.15405 | 3.72311 | 4.44116 | 0.16312 | |
| NAD+ | C00003 | I3sg5 | C13-11 | 21 | 65594.82167 | 85330.30718 | 64344.75651 | 3.2176055 | 0.8592 | 0.267 | 2.57782 | 2.88084 | 4.19415 | 3.21761 | 0.16312 | |
| NAD+ | C00003 | NT2 | C13-12 | 21 | 37647.88595 | 24193.4687 | 19185.7702 | 0.8373998 | 0.2271 | 0.2712 | 1.09955 | 0.71087 | 0.70179 | 0.8374 | 0.1136 | |
| NAD+ | C00003 | I3sg5 | C13-12 | 21 | 31040.85731 | 29637.39448 | 18213.39056 | 1.1358881 | 0.1183 | 0.1042 | 1.21988 | 1.00059 | 1.18719 | 1.13589 | 0.1136 | |
| NADH | C00004 | NT2 | C12 PARENT | 21 | 31442.30729 | 0 | 17416.4559 | 2.233793 | 2.0492 | 0.9174 | 4.02653 | 0 | 2.67485 | 2.23379 | 0.81209 | |
| NADH | C00004 | I3sg5 | C12 PARENT | 21 | 20027.16583 | 23248.22295 | 0 | 2.6907034 | 2.3487 | 0.8729 | 4.33015 | 3.74196 | 0 | 2.6907 | 0.81209 | |
| NADH | C00004 | NT2 | C13-04 | 21 | 7720.088256 | 10107.33088 | 8927.094018 | 1.3787484 | 0.394 | 0.2858 | 0.98864 | 1.77657 | 1.37104 | 1.37875 | 0.04804 | * |
| NADH | C00004 | I3sg5 | C13-04 | 21 | 4242.14599 | 0 | 0 | 0.3057371 | 0.5296 | 1.7321 | 0.91721 | 0 | 0 | 0.30574 | 0.04804 | * |
| NADH | C00004 | NT2 | C13-05 | 21 | 227230.0711 | 179466.9215 | 185292.6677 | 29.700584 | 1.6291 | 0.0549 | 29.0993 | 31.5449 | 28.4575 | 29.7006 | 0.74022 | |
| NADH | C00004 | I3sg5 | C13-05 | 21 | 124447.9534 | 187378.1607 | 88185.4492 | 30.562707 | 3.8725 | 0.1267 | 26.9074 | 30.1598 | 34.621 | 30.5627 | 0.74022 | |
| NADH | C00004 | NT2 | C13-06 | 21 | 0 | 0 | 0 | 0 | 0 | NA | 0 | 0 | 0 | 0 | NA | |
| NADH | C00004 | I3sg5 | C13-06 | 21 | 0 | 0 | 0 | 0 | 0 | NA | 0 | 0 | 0 | 0 | NA | |
| NADH | C00004 | NT2 | C13-07 | 21 | 2779.38839 | 0 | 17339.46668 | 1.006318 | 1.4457 | 1.4367 | 0.35593 | 0 | 2.66302 | 1.00632 | 0.99974 | |
| NADH | C00004 | I3sg5 | C13-07 | 21 | 4125.42635 | 13207.81007 | 0 | 1.0059535 | 1.0675 | 1.0612 | 0.89197 | 2.12589 | 0 | 1.00595 | 0.99974 | |
| NADH | C00004 | NT2 | C13-08 | 21 | 21869.91704 | 8303.294493 | 10991.58233 | 1.9827526 | 0.7175 | 0.3619 | 2.80068 | 1.45947 | 1.6881 | 1.98275 | 0.8352 | |
| NADH | C00004 | I3sg5 | C13-08 | 21 | 7002.922634 | 14204.69932 | 4671.91621 | 1.8782105 | 0.388 | 0.2066 | 1.51413 | 2.28634 | 1.83416 | 1.87821 | 0.8352 | |
| NADH | C00004 | NT2 | C13-09 | 21 | 16742.09497 | 19525.73503 | 27064.5458 | 3.2442213 | 1.0194 | 0.3142 | 2.14401 | 3.43204 | 4.15662 | 3.24422 | 0.11949 | |
| NADH | C00004 | I3sg5 | C13-09 | 21 | 3361.461897 | 14888.68062 | 5131.260015 | 1.7125744 | 0.8748 | 0.5108 | 0.72679 | 2.39643 | 2.0145 | 1.71257 | 0.11949 | |
| NADH | C00004 | NT2 | C13-10 | 21 | 435387.1211 | 321643.7601 | 332580.7152 | 54.456568 | 2.9515 | 0.0542 | 55.7561 | 56.5353 | 51.0783 | 54.4566 | 0.06553 | |
| NADH | C00004 | I3sg5 | C13-10 | 21 | 284175.0987 | 355543.6983 | 155862.8381 | 59.95346 | 2.3644 | 0.0394 | 61.4426 | 57.2271 | 61.1906 | 59.9535 | 0.06553 | |
| NADH | C00004 | NT2 | C13-11 | 21 | 28737.88712 | 20571.51114 | 45790.32508 | 4.7762026 | 1.9543 | 0.4092 | 3.6802 | 3.61586 | 7.03255 | 4.7762 | 0.07445 | |
| NADH | C00004 | I3sg5 | C13-11 | 21 | 11618.20034 | 12813.79658 | 865.4511281 | 1.6380845 | 1.1466 | 0.7 | 2.51202 | 2.06247 | 0.33977 | 1.63808 | 0.07445 | |
| NADH | C00004 | NT2 | C13-12 | 21 | 8969.218483 | 9306.614353 | 5716.856737 | 1.2208116 | 0.384 | 0.3146 | 1.14861 | 1.63582 | 0.878 | 1.22081 | 0.04497 | * |
| NADH | C00004 | I3sg5 | C13-12 | 21 | 3504.444703 | 0 | 0 | 0.25257 | 0.4375 | 1.7321 | 0.75771 | 0 | 0 | 0.25257 | 0.04497 | * |
| NADP+ | C00006 | NT2 | C12 PARENT | 21 | 16095.69471 | 15428.07437 | 10347.92706 | 3.838931 | 0.2256 | 0.0588 | 3.64779 | 4.08777 | 3.78124 | 3.83893 | 0.20362 | |
| NADP+ | C00006 | I3sg5 | C12 PARENT | 21 | 9209.551011 | 11633.30535 | 2944.741167 | 4.4221995 | 0.6261 | 0.1416 | 5.08245 | 3.837 | 4.34715 | 4.4222 | 0.20362 | |
| NADP+ | C00006 | NT2 | C13-04 | 21 | 4738.531701 | 2886.407065 | 2766.227481 | 0.9498265 | 0.1633 | 0.172 | 1.0739 | 0.76477 | 1.01081 | 0.94983 | 0.01128 | * |
| NADP+ | C00006 | I3sg5 | C13-04 | 21 | 0 | 1387.33111 | 0 | 0.1525272 | 0.2642 | 1.7321 | 0 | 0.45758 | 0 | 0.15253 | 0.01128 | * |
| NADP+ | C00006 | NT2 | C13-05 | 21 | 126565.7498 | 115645.4857 | 79146.41974 | 29.415227 | 1.0682 | 0.0363 | 28.6838 | 30.641 | 28.9209 | 29.4152 | 0.54182 | |

| | | | | | | | | | | | | | | | | |
|---|---|---|---|---|---|---|---|---|---|---|---|---|---|---|---|---|
| NADP+ | C00006 | I3sg5 | C13-05 | 21 | 57863.38276 | 92543.75107 | 19141.69671 | 30.23809 | 1.8541 | 0.0613 | 31.9329 | 30.5236 | 28.2578 | 30.2381 | 0.54182 | |
| NADP+ | C00006 | NT2 | C13-06 | 21 | 0 | 0 | 0 | 0 | 0 | NA | 0 | 0 | 0 | 0 | NA | |
| NADP+ | C00006 | I3sg5 | C13-06 | 21 | 0 | 0 | 0 | 0 | 0 | NA | 0 | 0 | 0 | 0 | NA | |
| NADP+ | C00006 | NT2 | C13-07 | 21 | 6188.203552 | 3568.646971 | 3141.007597 | 1.1652443 | 0.229 | 0.1965 | 1.40244 | 0.94554 | 1.14776 | 1.16524 | 0.82078 | |
| NADP+ | C00006 | I3sg5 | C13-07 | 21 | 2068.037899 | 4599.120952 | 424.9154325 | 1.0951609 | 0.4466 | 0.4078 | 1.14128 | 1.51692 | 0.62728 | 1.09516 | 0.82078 | |
| NADP+ | C00006 | NT2 | C13-08 | 21 | 8548.416299 | 8118.981842 | 4816.762071 | 1.9495363 | 0.1958 | 0.1004 | 1.93734 | 2.15118 | 1.76009 | 1.94954 | 0.46403 | |
| NADP+ | C00006 | I3sg5 | C13-08 | 21 | 3478.503182 | 5550.550629 | 2186.052127 | 2.3258484 | 0.7818 | 0.3361 | 1.91967 | 1.83073 | 3.22714 | 2.32585 | 0.46403 | |
| NADP+ | C00006 | NT2 | C13-09 | 21 | 10424.60947 | 8821.80329 | 7539.066973 | 2.4849292 | 0.2341 | 0.0942 | 2.36254 | 2.33739 | 2.75485 | 2.48493 | 0.26471 | |
| NADP+ | C00006 | I3sg5 | C13-09 | 21 | 4709.108096 | 6108.318295 | 1013.753466 | 2.0366823 | 0.5515 | 0.2708 | 2.5988 | 2.0147 | 1.49655 | 2.03668 | 0.26471 | |
| NADP+ | C00006 | NT2 | C13-10 | 21 | 251112.1069 | 204676.8245 | 157673.0065 | 56.251874 | 1.7858 | 0.0317 | 56.9099 | 54.2304 | 57.6153 | 56.2519 | 0.92332 | |
| NADP+ | C00006 | I3sg5 | C13-10 | 21 | 96129.1382 | 169601.072 | 40985.32519 | 56.49803 | 3.7581 | 0.0665 | 53.0505 | 55.9393 | 60.5043 | 56.498 | 0.92332 | |
| NADP+ | C00006 | NT2 | C13-11 | 21 | 8683.395159 | 13753.12201 | 5477.345582 | 2.5377948 | 0.9581 | 0.3775 | 1.96793 | 3.64398 | 2.00148 | 2.53779 | 0.85762 | |
| NADP+ | C00006 | I3sg5 | C13-11 | 21 | 5803.991027 | 10099.18156 | 307.7357919 | 2.3294418 | 1.6252 | 0.6977 | 3.20303 | 3.331 | 0.45429 | 2.32944 | 0.85762 | |
| NADP+ | C00006 | NT2 | C13-12 | 21 | 8888.219171 | 4521.314718 | 2757.479064 | 1.4066373 | 0.5348 | 0.3802 | 2.01435 | 1.19795 | 1.00761 | 1.40664 | 0.22894 | |
| NADP+ | C00006 | I3sg5 | C13-12 | 21 | 1941.292391 | 1665.018997 | 735.3496411 | 0.9020203 | 0.3057 | 0.3389 | 1.07134 | 0.54917 | 1.08555 | 0.90202 | 0.22894 | |
| Orotate | C00295 | NT2 | C12 PARENT | 5 | 2005618.673 | 1783282.924 | 1043646.18 | 72.628266 | 1.5418 | 0.0212 | 72.98 | 70.941 | 73.9638 | 72.6283 | 0.06835 | |
| Orotate | C00295 | I3sg5 | C12 PARENT | 5 | 691725.658 | 1508821.48 | 586996.7178 | 68.837512 | 2.1546 | 0.0313 | 68.6952 | 71.0598 | 66.7576 | 68.8375 | 0.06835 | |
| Orotate | C00295 | NT2 | C13-01 | 5 | 0 | 16024.92297 | 3601.577509 | 0.2975784 | 0.3208 | 1.0782 | 0 | 0.63749 | 0.25525 | 0.29758 | 0.73446 | |
| Orotate | C00295 | I3sg5 | C13-01 | 5 | 6003.994871 | 0 | 0 | 0.198752 | 0.3442 | 1.7321 | 0.59626 | 0 | 0 | 0.19875 | 0.73446 | |
| Orotate | C00295 | NT2 | C13-02 | 5 | 46407.15738 | 15382.61945 | 0 | 0.7668636 | 0.8549 | 1.1148 | 1.68865 | 0.61194 | 0 | 0.76686 | 0.32084 | |
| Orotate | C00295 | I3sg5 | C13-02 | 5 | 0 | 11099.67665 | 0 | 0.1742509 | 0.3018 | 1.7321 | 0 | 0.52275 | 0 | 0.17425 | 0.32084 | |
| Orotate | C00295 | NT2 | C13-03 | 5 | 696150.2551 | 699065.3532 | 363774.6065 | 26.307292 | 1.3203 | 0.0502 | 25.3314 | 27.8096 | 25.7809 | 26.3073 | 0.04773 | * |
| Orotate | C00295 | I3sg5 | C13-03 | 5 | 309219.351 | 603392.098 | 292299.4522 | 30.789485 | 2.4135 | 0.0784 | 30.7085 | 28.4175 | 33.2424 | 30.7895 | 0.04773 | * |
| Orotate | C00295 | NT2 | C13-04 | 5 | 0 | 0 | 0 | 0 | 0 | NA | 0 | 0 | 0 | 0 | NA | |
| Orotate | C00295 | I3sg5 | C13-04 | 5 | 0 | 0 | 0 | 0 | 0 | NA | 0 | 0 | 0 | 0 | NA | |
| Orotidine | X00006 | NT2 | C12 PARENT | 10 | 2120443.295 | 1963117.326 | 1919799.36 | 48.982368 | 0.7307 | 0.0149 | 48.1601 | 49.5573 | 49.2297 | 48.9824 | 0.07079 | |
| Orotidine | X00006 | I3sg5 | C12 PARENT | 10 | 1355207.826 | 1567248.358 | 1090114.497 | 52.062932 | 2.0558 | 0.0395 | 53.3872 | 53.107 | 49.6946 | 52.0629 | 0.07079 | |
| Orotidine | X00006 | NT2 | C13-01 | 10 | 0 | 0 | 0 | 0 | 0 | NA | 0 | 0 | 0 | 0 | NA | |
| Orotidine | X00006 | I3sg5 | C13-01 | 10 | 0 | 0 | 0 | 0 | 0 | NA | 0 | 0 | 0 | 0 | NA | |
| Orotidine | X00006 | NT2 | C13-04 | 10 | 27470.25015 | 16838.61785 | 21520.82774 | 0.533617 | 0.1007 | 0.1886 | 0.62391 | 0.42508 | 0.55186 | 0.53362 | 0.1599 | |
| Orotidine | X00006 | I3sg5 | C13-04 | 10 | 14503.1557 | 0 | 0 | 0.1904462 | 0.3299 | 1.7321 | 0.57134 | 0 | 0 | 0.19045 | 0.1599 | |
| Orotidine | X00006 | NT2 | C13-05 | 10 | 1641741.934 | 1418280.763 | 1374900.41 | 36.115941 | 1.0509 | 0.0291 | 37.2877 | 35.8033 | 35.2568 | 36.1159 | 0.36193 | |
| Orotidine | X00006 | I3sg5 | C13-05 | 10 | 859008.9007 | 941802.0626 | 819209.9672 | 34.366101 | 2.7537 | 0.0801 | 33.8399 | 31.9135 | 37.345 | 34.3661 | 0.36193 | |
| Orotidine | X00006 | NT2 | C13-06 | 10 | 57727.96304 | 24900.24832 | 47368.49058 | 1.0514659 | 0.3694 | 0.3513 | 1.31113 | 0.62859 | 1.21468 | 1.05147 | 0.18279 | |
| Orotidine | X00006 | I3sg5 | C13-06 | 10 | 0 | 28942.63266 | 10124.4542 | 0.4807585 | 0.4907 | 1.0206 | 0 | 0.98074 | 0.46154 | 0.48076 | 0.18279 | |
| Orotidine | X00006 | NT2 | C13-07 | 10 | 257395.6178 | 265160.61 | 259268.7133 | 6.3960905 | 0.4769 | 0.0746 | 5.84604 | 6.69376 | 6.64847 | 6.39609 | 0.78403 | |
| Orotidine | X00006 | I3sg5 | C13-07 | 10 | 148736.4412 | 190903.6275 | 143708.3992 | 6.2931264 | 0.3779 | 0.0601 | 5.85934 | 6.46887 | 6.55117 | 6.29313 | 0.78403 | |
| Orotidine | X00006 | NT2 | C13-08 | 10 | 298125.698 | 273012.8062 | 276816.7732 | 6.9205181 | 0.1655 | 0.0239 | 6.77111 | 6.89198 | 7.09846 | 6.92052 | 0.55276 | |
| Orotidine | X00006 | I3sg5 | C13-08 | 10 | 160995.1389 | 222215.0362 | 130472.1785 | 6.6066366 | 0.8235 | 0.1247 | 6.34226 | 7.52988 | 5.94778 | 6.60664 | 0.55276 | |
| P-Creatine | C02305 | NT2 | C12 PARENT | 4 | 2384468.35 | 2573297.767 | 3106549.009 | 99.040987 | 0.831 | 0.0084 | 98.5348 | 98.5881 | 100 | 99.041 | 0.56009 | |
| P-Creatine | C02305 | I3sg5 | C12 PARENT | 4 | 2579081.367 | 2493737.794 | 1521268.285 | 99.423596 | 0.6321 | 0.0064 | 99.5232 | 100 | 98.7476 | 99.4236 | 0.56009 | |
| P-Creatine | C02305 | NT2 | C13-01 | 4 | 11573.99242 | 11026.31895 | 0 | 0.3002397 | 0.2615 | 0.871 | 0.47828 | 0.42244 | 0 | 0.30024 | 0.43244 | |
| P-Creatine | C02305 | I3sg5 | C13-01 | 4 | 9933.423924 | 0 | 0 | 0.1277724 | 0.2213 | 1.7321 | 0.38332 | 0 | 0 | 0.12777 | 0.43244 | |
| P-Creatine | C02305 | NT2 | C13-02 | 4 | 23881.56412 | 25826.03419 | 0 | 0.6587729 | 0.5705 | 0.866 | 0.98687 | 0.98945 | 0 | 0.65877 | 0.70697 | |
| P-Creatine | C02305 | I3sg5 | C13-02 | 4 | 2423.04855 | 0 | 19293.88628 | 0.4486316 | 0.6976 | 1.5551 | 0.0935 | 0 | 1.25239 | 0.44863 | 0.70697 | |
| PEP | C00074 | NT2 | C12 PARENT | 3 | 0 | 0 | 13021.08884 | 0.8668154 | 1.5014 | 1.7321 | 0 | 0 | 2.60045 | 0.86682 | 0.15979 | |
| PEP | C00074 | I3sg5 | C12 PARENT | 3 | 42256.11615 | 15642.03556 | 11163.89226 | 4.6262745 | 3.4656 | 0.7491 | 8.6044 | 2.26136 | 3.01307 | 4.62627 | 0.15979 | |
| PEP | C00074 | NT2 | C13-03 | 3 | 562633.6168 | 466720.8494 | 487704.0722 | 99.133185 | 1.5014 | 0.0151 | 100 | 100 | 97.3996 | 99.1332 | 0.15979 | |
| PEP | C00074 | I3sg5 | C13-03 | 3 | 448842.7565 | 676068.515 | 359352.0127 | 95.373725 | 3.4656 | 0.0363 | 91.3956 | 97.7386 | 96.9869 | 95.3737 | 0.15979 | |
| Pro | C00148 | NT2 | C12 PARENT | 5 | 294753519.1 | 245530755.7 | 200559820.2 | 81.60877 | 0.4064 | 0.005 | 81.1747 | 81.6713 | 81.9802 | 81.6088 | 0.05431 | |
| Pro | C00148 | I3sg5 | C12 PARENT | 5 | 126955831.7 | 156060900.2 | 137215015.9 | 84.528164 | 1.8309 | 0.0217 | 86.6423 | 83.4653 | 83.4769 | 84.5282 | 0.05431 | |
| Pro | C00148 | NT2 | C13-01 | 5 | 3976745.138 | 4185429.677 | 2049423.222 | 1.1083712 | 0.2775 | 0.2503 | 1.09519 | 1.39221 | 0.83772 | 1.10837 | 0.33986 | |
| Pro | C00148 | I3sg5 | C13-01 | 5 | 52782.25395 | 2101456.159 | 1564025.377 | 0.7038109 | 0.5847 | 0.8308 | 0.03602 | 1.12391 | 0.9515 | 0.70381 | 0.33986 | |
| Pro | C00148 | NT2 | C13-03 | 5 | 19832716.1 | 15116279.4 | 12694269.62 | 5.2263105 | 0.2193 | 0.042 | 5.4619 | 5.02816 | 5.18887 | 5.22631 | 0.0338 | * |
| Pro | C00148 | I3sg5 | C13-03 | 5 | 5260970.473 | 8447775.392 | 7339500.009 | 4.191195 | 0.521 | 0.1243 | 3.5904 | 4.51808 | 4.4651 | 4.1912 | 0.0338 | * |
| Pro | C00148 | NT2 | C13-04 | 5 | 27290174.99 | 22195440.87 | 17773928.75 | 7.3879356 | 0.1253 | 0.017 | 7.51568 | 7.38291 | 7.26522 | 7.38794 | 0.0722 | |
| Pro | C00148 | I3sg5 | C13-04 | 5 | 8114402.395 | 12300551.02 | 11408608.54 | 6.3523353 | 0.7283 | 0.1146 | 5.53776 | 6.57864 | 6.94061 | 6.35234 | 0.0722 | |
| Pro | C00148 | NT2 | C13-05 | 5 | 17256784 | 13604762.82 | 11566685.64 | 4.6686123 | 0.1247 | 0.0267 | 4.7525 | 4.52538 | 4.72796 | 4.66861 | 0.00645 | ** |
| Pro | C00148 | I3sg5 | C13-05 | 5 | 6144710.718 | 8066388.838 | 6847619.189 | 4.2244947 | 0.0788 | 0.0187 | 4.19352 | 4.31411 | 4.16586 | 4.22449 | 0.00645 | ** |
| Propanoyl-CoA | C00100 | NT2 | C12 PARENT | 24 | 1453.322549 | 1852.811813 | 3384.146769 | 21.848135 | 1.119 | 0.0512 | 22.75 | 20.5959 | 22.1985 | 21.8481 | 0.4484 | |
| Propanoyl-CoA | C00100 | I3sg5 | C12 PARENT | 24 | 3063.239657 | 865.0843734 | 2028.384174 | 24.427858 | 5.203 | 0.213 | 30.4343 | 21.3074 | 21.5419 | 24.4279 | 0.4484 | |
| Propanoyl-CoA | C00100 | NT2 | C13-05 | 24 | 4934.914654 | 6636.097745 | 11091.31605 | 74.590411 | 2.3583 | 0.0316 | 77.25 | 73.767 | 72.7542 | 74.5904 | 0.37823 | |
| Propanoyl-CoA | C00100 | I3sg5 | C13-05 | 24 | 7001.859489 | 3097.720756 | 6632.69546 | 72.101551 | 3.6606 | 0.0508 | 69.5657 | 76.2982 | 70.4407 | 72.1016 | 0.37823 | |
| Propanoyl-CoA | C00100 | NT2 | C13-06 | 24 | 0 | 507.115888 | 769.4489392 | 3.5614538 | 3.0984 | 0.87 | 0 | 5.63711 | 5.04725 | 3.56145 | 0.97709 | |
| Propanoyl-CoA | C00100 | I3sg5 | C13-06 | 24 | 0 | 97.21375939 | 754.9139112 | 3.4705915 | 4.1156 | 1.1858 | 0 | 2.39442 | 8.01736 | 3.47059 | 0.97709 | |
| R5P | C00117 | NT2 | C12 PARENT | 5 | 20649.28217 | 18903.57978 | 0 | 2.0654608 | 1.7898 | 0.8665 | 3.0365 | 3.15988 | 0 | 2.06546 | 0.11627 | |
| R5P | C00117 | I3sg5 | C12 PARENT | 5 | 0 | 0 | 0 | 0 | 0 | NA | 0 | 0 | 0 | 0 | 0.11627 | |
| R5P | C00117 | NT2 | C13-05 | 5 | 659385.8791 | 579333.6178 | 747973.2826 | 97.934539 | 1.7898 | 0.0183 | 96.9635 | 96.8401 | 100 | 97.9345 | 0.11627 | |
| R5P | C00117 | I3sg5 | C13-05 | 5 | 573417.5413 | 529424.1485 | 627469.5018 | 100 | 0 | 0 | 100 | 100 | 100 | 100 | 0.11627 | |
| S7P | C05382 | NT2 | C12 PARENT | 7 | 43525.05266 | 70818.4658 | 96197.7364 | 20.289895 | 2.885 | 0.1422 | 16.9752 | 22.2352 | 21.6593 | 20.2899 | 0.20263 | |
| S7P | C05382 | I3sg5 | C12 PARENT | 7 | 23477.18186 | 94544.0211 | 54507.66551 | 13.198715 | 7.5356 | 0.5709 | 5.82136 | 20.8831 | 12.8917 | 13.1987 | 0.20263 | |
| S7P | C05382 | NT2 | C13-07 | 7 | 212879.0631 | 247677.9128 | 347943.6647 | 79.710105 | 2.885 | 0.0362 | 83.0248 | 77.7648 | 78.3407 | 79.7101 | 0.20263 | |
| S7P | C05382 | I3sg5 | C13-07 | 7 | 379816.8214 | 358185.3189 | 368305.4882 | 86.801285 | 7.5356 | 0.0868 | 94.1786 | 79.1169 | 87.1083 | 86.8013 | 0.20263 | |
| SAM | C00019 | NT2 | C12 PARENT | 15 | 1549210.144 | 1395929.958 | 1737333.037 | 18.327188 | 0.4618 | 0.0252 | 18.2364 | 17.9175 | 18.8277 | 18.3272 | 0.00184 | ** |
| SAM | C00019 | I3sg5 | C12 PARENT | 15 | 1074756.501 | 1060677.464 | 748478.5094 | 20.488096 | 0.2172 | 0.0106 | 20.7087 | 20.4812 | 20.2744 | 20.4881 | 0.00184 | ** |
| SAM | C00019 | NT2 | C13-01 | 15 | 0 | 0 | 0 | 0 | 0 | NA | 0 | 0 | 0 | 0 | NA | |
| SAM | C00019 | I3sg5 | C13-01 | 15 | 0 | 0 | 0 | 0 | 0 | NA | 0 | 0 | 0 | 0 | NA | |
| SAM | C00019 | NT2 | C13-02 | 15 | 0 | 0 | 0 | 0 | 0 | NA | 0 | 0 | 0 | 0 | NA | |
| SAM | C00019 | I3sg5 | C13-02 | 15 | 0 | 0 | 0 | 0 | 0 | NA | 0 | 0 | 0 | 0 | NA | |
| SAM | C00019 | NT2 | C13-03 | 15 | 80742.56848 | 77346.90396 | 108406.4089 | 1.0393517 | 0.1192 | 0.1147 | 0.95045 | 0.99279 | 1.17481 | 1.03935 | 0.86961 | |
| SAM | C00019 | I3sg5 | C13-03 | 15 | 57374.54613 | 60021.28914 | 33504.83068 | 1.057351 | 0.1324 | 0.1253 | 1.10551 | 1.15898 | 0.90756 | 1.05735 | 0.86961 | |
| SAM | C00019 | NT2 | C13-04 | 15 | 156846.3285 | 118274.2148 | 143794.2483 | 1.6409096 | 0.179 | 0.1091 | 1.8463 | 1.51811 | 1.55832 | 1.64091 | 0.02456 | * |
| SAM | C00019 | I3sg5 | C13-04 | 15 | 66442.60414 | 63600.21849 | 48024.44382 | 1.2697287 | 0.0375 | 0.0295 | 1.28023 | 1.22809 | 1.30086 | 1.26973 | 0.02456 | * |
| SAM | C00019 | NT2 | C13-05 | 15 | 6252779.289 | 5799441.569 | 6804472.689 | 73.927925 | 0.4478 | 0.0061 | 73.6039 | 74.439 | 73.7409 | 73.9279 | 0.4552 | |
| SAM | C00019 | I3sg5 | C13-05 | 15 | 3924655.425 | 3836359.418 | 2720196.082 | 74.460968 | 1.024 | 0.0138 | 75.6212 | 74.0784 | 73.6833 | 74.461 | 0.4552 | |
| SAM | C00019 | NT2 | C13-06 | 15 | 455592.0661 | 399874.7492 | 433539.3197 | 5.0646265 | 0.3375 | 0.0666 | 5.36295 | 5.13261 | 4.69832 | 5.06463 | 0.03974 | * |
| SAM | C00019 | I3sg5 | C13-06 | 15 | 66658.73624 | 158123.0996 | 141537.1636 | 2.7238569 | 1.3063 | 0.4796 | 1.2844 | 3.05329 | 3.83389 | 2.72386 | 0.03974 | * |
| SAM | C00019 | NT2 | C13-07 | 15 | 0 | 0 | 0 | 0 | 0 | NA | 0 | 0 | 0 | 0 | NA | |
| SAM | C00019 | I3sg5 | C13-07 | 15 | 0 | 0 | 0 | 0 | 0 | NA | 0 | 0 | 0 | 0 | NA | |
| SAM | C00019 | NT2 | C13-08 | 15 | 0 | 0 | 0 | 0 | 0 | NA | 0 | 0 | 0 | 0 | NA | |
| SAM | C00019 | I3sg5 | C13-08 | 15 | 0 | 0 | 0 | 0 | 0 | NA | 0 | 0 | 0 | 0 | NA | |
| Ser | C00065 | NT2 | C12 PARENT | 3 | 43957831.83 | 50912459.49 | 43676148.47 | 87.723796 | 1.3005 | 0.0148 | 86.2235 | 88.4187 | 88.5292 | 87.7238 | 0.03626 | * |
| Ser | C00065 | I3sg5 | C12 PARENT | 3 | 30382581.56 | 37604947.24 | 26182386.84 | 90.709909 | 1.0462 | 0.0115 | 91.1428 | 91.4701 | 89.5168 | 90.7099 | 0.03626 | * |
| Ser | C00065 | NT2 | C13-01 | 3 | 3034488.49 | 2844145.718 | 2571863.304 | 5.3681882 | 0.5239 | 0.0976 | 5.95216 | 4.93937 | 5.21303 | 5.36819 | 0.02527 | * |
| Ser | C00065 | I3sg5 | C13-01 | 3 | 994927.1738 | 1544031.176 | 1255427.486 | 3.677528 | 0.6573 | 0.1787 | 2.98462 | 3.7557 | 4.29227 | 3.67753 | 0.02527 | * |
| Ser | C00065 | NT2 | C13-02 | 3 | 1887584.322 | 1378135.716 | 1376023.344 | 2.9616707 | 0.6714 | 0.2267 | 3.7025 | 2.39338 | 2.78913 | 2.96167 | 0.44765 | |
| Ser | C00065 | I3sg5 | C13-02 | 3 | 926636.1776 | 912313.1822 | 818104.6291 | 2.5986467 | 0.3288 | 0.1265 | 2.77976 | 2.21911 | 2.79707 | 2.59865 | 0.44765 | |
| Ser | C00065 | NT2 | C13-03 | 3 | 2101389.055 | 2446350.161 | 1711255.53 | 3.9463455 | 0.4185 | 0.1061 | 4.12188 | 4.24853 | 3.46862 | 3.94635 | 0.05366 | |
| Ser | C00065 | I3sg5 | C13-03 | 3 | 1030994.485 | 1050424.108 | 992662.825 | 3.0139159 | 0.4249 | 0.141 | 3.09282 | 2.55505 | 3.39388 | 3.01392 | 0.05366 | |

| | | | | | | | | | | | | | | | | |
|---|---|---|---|---|---|---|---|---|---|---|---|---|---|---|---|---|
| Sorbitol | C00794 | NT2 | C12 PARENT | 6 | 24815356.36 | 61663364.95 | 32849229.66 | 98.112811 | 0.8681 | 0.0088 | 97.1826 | 98.9014 | 98.2544 | 98.1128 | 0.98154 | |
| Sorbitol | C00794 | I3sg5 | C12 PARENT | 6 | 41200255.31 | 21046537.03 | 24231548.76 | 98.09196 | 1.1824 | 0.0121 | 99.0803 | 96.782 | 98.4136 | 98.092 | 0.98154 | |
| Sorbitol | C00794 | NT2 | C13-01 | 6 | 0 | 0 | 0 | 0 | 0 | NA | 0 | 0 | 0 | 0 | NA | |
| Sorbitol | C00794 | I3sg5 | C13-01 | 6 | 0 | 0 | 0 | 0 | 0 | NA | 0 | 0 | 0 | 0 | NA | |
| Sorbitol | C00794 | NT2 | C13-06 | 6 | 719416.4797 | 684947.5826 | 583598.9495 | 1.8871894 | 0.8681 | 0.46 | 2.8174 | 1.09858 | 1.74559 | 1.88719 | 0.98154 | |
| Sorbitol | C00794 | I3sg5 | C13-06 | 6 | 382453.8929 | 699799.7259 | 390597.4873 | 1.9080404 | 1.1824 | 0.6197 | 0.91974 | 3.21801 | 1.58637 | 1.90804 | 0.98154 | |
| Succ | C00042 | NT2 | C12 PARENT | 4 | 2591204.804 | 2203844.459 | 3249629.093 | 78.146132 | 0.9458 | 0.0121 | 78.708 | 77.0542 | 78.6762 | 78.1461 | 0.00375 | ** |
| Succ | C00042 | I3sg5 | C12 PARENT | 4 | 2528543.586 | 2654748.873 | 2009667.698 | 82.19056 | 0.6657 | 0.0081 | 82.3998 | 81.4453 | 82.7265 | 82.1906 | 0.00375 | ** |
| Succ | C00042 | NT2 | C13-01 | 4 | 0 | 0 | 0 | 0 | 0 | NA | 0 | 0 | 0 | 0 | NA | |
| Succ | C00042 | I3sg5 | C13-01 | 4 | 0 | 0 | 0 | 0 | 0 | NA | 0 | 0 | 0 | 0 | NA | |
| Succ | C00042 | NT2 | C13-02 | 4 | 439382.0956 | 291429.4174 | 461058.8362 | 11.566091 | 1.6166 | 0.1398 | 13.3463 | 10.1894 | 11.1626 | 11.5661 | 0.11233 | |
| Succ | C00042 | I3sg5 | C13-02 | 4 | 309429.1888 | 298432.6206 | 231904.277 | 9.5951515 | 0.4659 | 0.0486 | 10.0836 | 9.15565 | 9.54617 | 9.59515 | 0.11233 | |
| Succ | C00042 | NT2 | C13-03 | 4 | 140434.8828 | 224093.6482 | 267474.9302 | 6.1922048 | 1.8015 | 0.2909 | 4.26572 | 7.83511 | 6.47579 | 6.1922 | 0.28057 | |
| Succ | C00042 | I3sg5 | C13-03 | 4 | 146400.4475 | 177926.7524 | 102995.314 | 4.8230798 | 0.6111 | 0.1267 | 4.77088 | 5.45863 | 4.23973 | 4.82308 | 0.28057 | |
| Succ | C00042 | NT2 | C13-04 | 4 | 121154.1285 | 140754.0136 | 152220.8473 | 4.0955716 | 0.7151 | 0.1746 | 3.68006 | 4.92126 | 3.68539 | 4.09557 | 0.26217 | |
| Succ | C00042 | I3sg5 | C13-04 | 4 | 84254.63437 | 128438.8144 | 84722.95772 | 3.3912088 | 0.6032 | 0.1779 | 2.74568 | 3.94039 | 3.48756 | 3.39121 | 0.26217 | |
| Succinyl-CoA | C00091 | NT2 | C12 PARENT | 25 | 5720.871171 | 4820.536982 | 5920.928823 | 24.458081 | 2.0245 | 0.0828 | 22.473 | 26.5198 | 24.3814 | 24.4581 | 0.03758 | * |
| Succinyl-CoA | C00091 | I3sg5 | C12 PARENT | 25 | 2624.679324 | 1965.743981 | 1328.438063 | 20.585365 | 0.8365 | 0.0406 | 20.4534 | 19.8227 | 21.48 | 20.5854 | 0.03758 | * |
| Succinyl-CoA | C00091 | NT2 | C13-05 | 25 | 14396.14515 | 10840.43327 | 16700.50184 | 61.653139 | 6.3536 | 0.1031 | 56.5517 | 59.6377 | 68.77 | 61.6531 | 0.33289 | |
| Succinyl-CoA | C00091 | I3sg5 | C13-05 | 25 | 8495.102687 | 6714.309775 | 3942.502176 | 65.885101 | 1.9987 | 0.0303 | 66.2 | 67.7076 | 63.7477 | 65.8851 | 0.33289 | |
| Succinyl-CoA | C00091 | NT2 | C13-06 | 25 | 4074.586153 | 2516.165276 | 866.1159288 | 11.138333 | 6.646 | 0.5967 | 16.006 | 13.8425 | 3.56653 | 11.1383 | 0.5725 | |
| Succinyl-CoA | C00091 | I3sg5 | C13-06 | 25 | 1712.698229 | 1236.569731 | 913.6023348 | 13.529535 | 1.1622 | 0.0859 | 13.3466 | 12.4697 | 14.7724 | 13.5295 | 0.5725 | |
| Succinyl-CoA | C00091 | NT2 | C13-07 | 25 | 1265.018348 | 0 | 797.0277289 | 2.7504476 | 2.5269 | 0.9187 | 4.96931 | 0 | 3.28203 | 2.75045 | 0.13248 | |
| Succinyl-CoA | C00091 | I3sg5 | C13-07 | 25 | 0 | 0 | 0 | 0 | 0 | NA | 0 | 0 | 0 | 0 | 0.13248 | |
| UDP | C00015 | NT2 | C12 PARENT | 9 | 1169622.375 | 1125800.059 | 904563.3619 | 13.725836 | 0.3483 | 0.0254 | 13.3854 | 13.7105 | 14.0815 | 13.7258 | 0.08696 | |
| UDP | C00015 | I3sg5 | C12 PARENT | 9 | 727098.0705 | 753870.4435 | 552127.7876 | 14.732332 | 0.6894 | 0.0468 | 14.4875 | 15.5107 | 14.1988 | 14.7323 | 0.08696 | |
| UDP | C00015 | NT2 | C13-01 | 9 | 0 | 0 | 0 | 0 | 0 | NA | 0 | 0 | 0 | 0 | NA | |
| UDP | C00015 | I3sg5 | C13-01 | 9 | 0 | 0 | 0 | 0 | 0 | NA | 0 | 0 | 0 | 0 | NA | |
| UDP | C00015 | NT2 | C13-02 | 9 | 0 | 0 | 0 | 0 | 0 | NA | 0 | 0 | 0 | 0 | NA | |
| UDP | C00015 | I3sg5 | C13-02 | 9 | 0 | 0 | 0 | 0 | 0 | NA | 0 | 0 | 0 | 0 | NA | |
| UDP | C00015 | NT2 | C13-03 | 9 | 70796.95061 | 74610.46108 | 29753.33627 | 0.7273454 | 0.234 | 0.3217 | 0.81022 | 0.90864 | 0.46318 | 0.72735 | 0.03955 | * |
| UDP | C00015 | I3sg5 | C13-03 | 9 | 16249.99213 | 12165.62535 | 13835.77418 | 0.309965 | 0.0541 | 0.1745 | 0.32378 | 0.25031 | 0.35581 | 0.30996 | 0.03955 | * |
| UDP | C00015 | NT2 | C13-04 | 9 | 166224.2327 | 90733.2518 | 122609.2866 | 1.6386627 | 0.4622 | 0.282 | 1.90231 | 1.10499 | 1.90868 | 1.63866 | 0.14603 | |
| UDP | C00015 | I3sg5 | C13-04 | 9 | 28589.73588 | 47072.3883 | 55180.34355 | 0.9857343 | 0.425 | 0.4311 | 0.56965 | 0.9685 | 1.41905 | 0.98573 | 0.14603 | |
| UDP | C00015 | NT2 | C13-05 | 9 | 4778871.53 | 4590443.958 | 3634238.77 | 55.72339 | 0.9552 | 0.0171 | 54.6905 | 55.9047 | 56.575 | 55.7234 | 0.60349 | |
| UDP | C00015 | I3sg5 | C13-05 | 9 | 2812854.939 | 2590765.261 | 2178454.151 | 55.124327 | 1.5761 | 0.0286 | 56.0463 | 53.3045 | 56.0223 | 55.1243 | 0.60349 | |
| UDP | C00015 | NT2 | C13-06 | 9 | 642722.6932 | 570209.2744 | 421382.3892 | 6.9531688 | 0.3979 | 0.0572 | 7.35547 | 6.94429 | 6.55975 | 6.95317 | 0.17809 | |
| UDP | C00015 | I3sg5 | C13-06 | 9 | 342913.3592 | 407409.9146 | 331724.2199 | 7.9152439 | 0.9406 | 0.1188 | 6.83256 | 8.38238 | 8.53079 | 7.91524 | 0.17809 | |
| UDP | C00015 | NT2 | C13-07 | 9 | 1170416.215 | 1125163.431 | 867598.8588 | 13.534468 | 0.1561 | 0.0115 | 13.3945 | 13.7028 | 13.5061 | 13.5345 | 0.62185 | |
| UDP | C00015 | I3sg5 | C13-07 | 9 | 698443.1208 | 663433.0198 | 472043.9996 | 13.235279 | 0.9584 | 0.0724 | 13.9165 | 13.65 | 12.1393 | 13.2353 | 0.62185 | |
| UDP | C00015 | NT2 | C13-08 | 9 | 739370.2684 | 634238.3124 | 443611.5084 | 7.6971288 | 0.7782 | 0.1011 | 8.46153 | 7.72406 | 6.90579 | 7.69713 | 0.99998 | |
| UDP | C00015 | I3sg5 | C13-08 | 9 | 392660.5563 | 385598.2257 | 285185.2942 | 7.6971189 | 0.3193 | 0.0415 | 7.82378 | 7.93361 | 7.33397 | 7.69712 | 0.99998 | |
| UDP-GlcNAc | C00043 | NT2 | C12 PARENT | 17 | 846607.5769 | 913110.3755 | 1074176.315 | 4.2284197 | 0.4349 | 0.1029 | 4.03827 | 3.92098 | 4.72601 | 4.22842 | 0.03833 | * |
| UDP-GlcNAc | C00043 | I3sg5 | C12 PARENT | 17 | 821801.709 | 853990.3916 | 737076.5427 | 5.2627827 | 0.3972 | 0.0755 | 4.80463 | 5.47456 | 5.50916 | 5.26278 | 0.03833 | * |
| UDP-GlcNAc | C00043 | NT2 | C13-01 | 17 | 0 | 0 | 0 | 0 | 0 | NA | 0 | 0 | 0 | 0 | NA | |
| UDP-GlcNAc | C00043 | I3sg5 | C13-01 | 17 | 0 | 0 | 0 | 0 | 0 | NA | 0 | 0 | 0 | 0 | NA | |
| UDP-GlcNAc | C00043 | NT2 | C13-02 | 17 | 212331.3946 | 129026.3191 | 226479.435 | 0.8544307 | 0.2603 | 0.3046 | 1.01281 | 0.55405 | 0.99643 | 0.85443 | 0.33214 | |
| UDP-GlcNAc | C00043 | I3sg5 | C13-02 | 17 | 119205.9511 | 94125.11375 | 99928.30065 | 0.6824085 | 0.0728 | 0.1067 | 0.69693 | 0.60339 | 0.7469 | 0.68241 | 0.33214 | |
| UDP-GlcNAc | C00043 | NT2 | C13-04 | 17 | 36908.02564 | 27194.27243 | 95953.46184 | 0.2383288 | 0.1619 | 0.6795 | 0.17605 | 0.11677 | 0.42216 | 0.23833 | 0.13048 | |
| UDP-GlcNAc | C00043 | I3sg5 | C13-04 | 17 | 116676.0908 | 105268.6291 | 37628.67611 | 0.5460741 | 0.2294 | 0.42 | 0.68214 | 0.67483 | 0.28125 | 0.54607 | 0.13048 | |
| UDP-GlcNAc | C00043 | NT2 | C13-05 | 17 | 2016538.721 | 2290385.929 | 2589276.178 | 10.281944 | 0.9673 | 0.0941 | 9.61878 | 9.83512 | 11.3919 | 10.2819 | 0.0197 | * |
| UDP-GlcNAc | C00043 | I3sg5 | C13-05 | 17 | 2298017.659 | 2111245.042 | 1617515.453 | 13.019797 | 0.8069 | 0.062 | 13.4353 | 13.5343 | 12.0899 | 13.0198 | 0.0197 | * |
| UDP-GlcNAc | C00043 | NT2 | C13-06 | 17 | 1273220.319 | 1833316.177 | 1884431.212 | 7.4121562 | 1.1783 | 0.159 | 6.07319 | 7.87242 | 8.29085 | 7.41216 | 0.74008 | |
| UDP-GlcNAc | C00043 | I3sg5 | C13-06 | 17 | 1372858.364 | 1205324.582 | 969757.4371 | 7.6671543 | 0.3924 | 0.0512 | 8.02636 | 7.72681 | 7.2483 | 7.66715 | 0.74008 | |
| UDP-GlcNAc | C00043 | NT2 | C13-07 | 17 | 888535.2822 | 970082.3179 | 1010710.582 | 4.2835556 | 0.146 | 0.0341 | 4.23827 | 4.16562 | 4.44678 | 4.28356 | 0.00375 | ** |
| UDP-GlcNAc | C00043 | I3sg5 | C13-07 | 17 | 1034156.914 | 932302.1721 | 711411.3775 | 5.780022 | 0.4022 | 0.0696 | 6.04615 | 5.97658 | 5.31733 | 5.78002 | 0.00375 | ** |
| UDP-GlcNAc | C00043 | NT2 | C13-08 | 17 | 1626275.684 | 2032888.542 | 1575252.313 | 7.8057412 | 0.9004 | 0.1154 | 7.75725 | 8.7294 | 6.93057 | 7.80574 | 0.6588 | |
| UDP-GlcNAc | C00043 | I3sg5 | C13-08 | 17 | 1325325.588 | 1162687.384 | 998054.9381 | 7.5539138 | 0.1685 | 0.0223 | 7.74846 | 7.45348 | 7.4598 | 7.55391 | 0.6588 | |
| UDP-GlcNAc | C00043 | NT2 | C13-09 | 17 | 248607.2024 | 330190.7756 | 225805.7978 | 1.1990728 | 0.2125 | 0.1772 | 1.18584 | 1.41791 | 0.99347 | 1.19907 | 0.9753 | |
| UDP-GlcNAc | C00043 | I3sg5 | C13-09 | 17 | 132043.0086 | 240219.9201 | 175378.9589 | 1.2075902 | 0.3943 | 0.3265 | 0.77198 | 1.53994 | 1.31084 | 1.20759 | 0.9753 | |
| UDP-GlcNAc | C00043 | NT2 | C13-10 | 17 | 395305.5886 | 379691.3314 | 325477.935 | 1.6493355 | 0.2274 | 0.1379 | 1.88559 | 1.63043 | 1.43199 | 1.64934 | 0.42612 | |
| UDP-GlcNAc | C00043 | I3sg5 | C13-10 | 17 | 260223.6235 | 281434.7025 | 317598.4066 | 1.8997932 | 0.4342 | 0.2286 | 1.52139 | 1.80415 | 2.37384 | 1.89979 | 0.42612 | |
| UDP-GlcNAc | C00043 | NT2 | C13-11 | 17 | 4134236.951 | 4796454.034 | 4827090.211 | 20.518012 | 0.7618 | 0.0371 | 19.7201 | 20.5964 | 21.2376 | 20.518 | 0.26993 | |
| UDP-GlcNAc | C00043 | I3sg5 | C13-11 | 17 | 3563540.076 | 2945954.63 | 2507943.66 | 19.488178 | 1.1677 | 0.0599 | 20.8341 | 18.8852 | 18.7452 | 19.4882 | 0.26993 | |
| UDP-GlcNAc | C00043 | NT2 | C13-12 | 17 | 948436.1617 | 900652.422 | 717349.7478 | 3.8491884 | 0.6841 | 0.1777 | 4.52399 | 3.86748 | 3.15609 | 3.84919 | 0.52056 | |
| UDP-GlcNAc | C00043 | I3sg5 | C13-12 | 17 | 497708.8371 | 534360.7017 | 549108.546 | 3.47987 | 0.599 | 0.1721 | 2.90983 | 3.42555 | 4.10422 | 3.47987 | 0.52056 | |
| UDP-GlcNAc | C00043 | NT2 | C13-13 | 17 | 4660569.786 | 4943626.198 | 4811264.254 | 21.542319 | 0.5969 | 0.0277 | 22.2307 | 21.2284 | 21.1679 | 21.5423 | 0.02421 | * |
| UDP-GlcNAc | C00043 | I3sg5 | C13-13 | 17 | 3246266.995 | 2988068.235 | 2733312.323 | 19.521355 | 0.7916 | 0.0405 | 18.9792 | 19.1552 | 20.4297 | 19.5214 | 0.02421 | * |
| UDP-GlcNAc | C00043 | NT2 | C13-14 | 17 | 1433532.669 | 1420952.49 | 1367492.492 | 6.3186902 | 0.4516 | 0.0715 | 6.83788 | 6.10169 | 6.0165 | 6.31869 | 0.04209 | * |
| UDP-GlcNAc | C00043 | I3sg5 | C13-14 | 17 | 959030.1384 | 864364.0216 | 733160.6664 | 5.5426267 | 0.0635 | 0.0115 | 5.60693 | 5.54106 | 5.47989 | 5.54263 | 0.04209 | * |
| UDP-GlcNAc | C00043 | NT2 | C13-15 | 17 | 1516278.795 | 1715315.729 | 1260785.727 | 6.7151044 | 1.0138 | 0.151 | 7.23257 | 7.36572 | 5.54703 | 6.7151 | 0.13261 | |
| UDP-GlcNAc | C00043 | I3sg5 | C13-15 | 17 | 785603.8212 | 907314.8446 | 768140.1228 | 5.3835794 | 0.6857 | 0.1274 | 4.593 | 5.8164 | 5.74134 | 5.38358 | 0.13261 | |
| UDP-GlcNAc | C00043 | NT2 | C13-16 | 17 | 727206.2049 | 604940.6757 | 737488.238 | 3.1037001 | 0.4523 | 0.1457 | 3.46874 | 2.59767 | 3.2447 | 3.1037 | 0.74133 | |
| UDP-GlcNAc | C00043 | I3sg5 | C13-16 | 17 | 571918.7132 | 372593.5508 | 423092.1323 | 2.9648556 | 0.5073 | 0.1711 | 3.3437 | 2.38853 | 3.16233 | 2.96486 | 0.74133 | |
| UMP | C00105 | NT2 | C12 PARENT | 9 | 124876.1477 | 315040.1115 | 95422.22118 | 11.418093 | 4.7206 | 0.4134 | 9.38004 | 16.8153 | 8.05894 | 11.4181 | 0.96829 | |
| UMP | C00105 | I3sg5 | C12 PARENT | 9 | 98820.21511 | 114060.6052 | 75488.73796 | 11.536025 | 1.0171 | 0.0882 | 12.5678 | 11.5061 | 10.5342 | 11.536 | 0.96829 | |
| UMP | C00105 | NT2 | C13-04 | 9 | 0 | 0 | 0 | 0 | 0 | NA | 0 | 0 | 0 | 0 | NA | |
| UMP | C00105 | I3sg5 | C13-04 | 9 | 0 | 0 | 0 | 0 | 0 | NA | 0 | 0 | 0 | 0 | NA | |
| UMP | C00105 | NT2 | C13-05 | 9 | 875525.5732 | 1284027.079 | 896586.5008 | 70.007241 | 5.1391 | 0.0734 | 65.7649 | 68.5351 | 75.7218 | 70.0072 | 0.25013 | |
| UMP | C00105 | I3sg5 | C13-05 | 9 | 512046.944 | 683715.7378 | 430934.6085 | 64.74263 | 4.4299 | 0.0684 | 65.1213 | 68.971 | 60.1356 | 64.7426 | 0.25013 | |
| UMP | C00105 | NT2 | C13-06 | 9 | 102280.5809 | 41745.00908 | 0 | 3.3036417 | 3.9527 | 1.1965 | 7.68278 | 2.22814 | 0 | 3.30364 | 0.70593 | |
| UMP | C00105 | I3sg5 | C13-06 | 9 | 0 | 98720.83543 | 31688.10933 | 4.793536 | 4.9897 | 1.0409 | 0 | 9.95863 | 4.42198 | 4.79354 | 0.70593 | |
| UMP | C00105 | NT2 | C13-07 | 9 | 146128.8477 | 223131.5686 | 95395.77197 | 10.31427 | 2.01 | 0.1949 | 10.9764 | 11.9097 | 8.05671 | 10.3143 | 0.53672 | |
| UMP | C00105 | I3sg5 | C13-07 | 9 | 139315.6796 | 61298.48056 | 102935.7882 | 12.755302 | 5.9331 | 0.4652 | 17.7179 | 6.18359 | 14.3644 | 12.7553 | 0.53672 | |
| UMP | C00105 | NT2 | C13-08 | 9 | 82485.0807 | 9589.334197 | 96649.37704 | 4.9567539 | 3.973 | 0.8015 | 6.19585 | 0.51183 | 8.16258 | 4.95675 | 0.72231 | |
| UMP | C00105 | I3sg5 | C13-08 | 9 | 36114.01629 | 33513.6319 | 75557.78882 | 6.1725074 | 3.8339 | 0.6211 | 4.59292 | 3.38074 | 10.5439 | 6.17251 | 0.72231 | |
| UTP | C00075 | NT2 | C12 PARENT | 9 | 1190778.889 | 1046043.424 | 1205681.45 | 14.474212 | 0.9169 | 0.0633 | 14.6225 | 13.4922 | 15.3079 | 14.4742 | 0.26237 | |
| UTP | C00075 | I3sg5 | C12 PARENT | 9 | 798740.6701 | 706686.6637 | 419608.1471 | 13.362047 | 1.159 | 0.0867 | 14.1859 | 13.8635 | 12.0368 | 13.362 | 0.26237 | |
| UTP | C00075 | NT2 | C13-01 | 9 | 0 | 0 | 0 | 0 | 0 | NA | 0 | 0 | 0 | 0 | NA | |
| UTP | C00075 | I3sg5 | C13-01 | 9 | 0 | 0 | 0 | 0 | 0 | NA | 0 | 0 | 0 | 0 | NA | |
| UTP | C00075 | NT2 | C13-02 | 9 | 13884.36167 | 33907.9518 | 17199.72155 | 0.2754096 | 0.1423 | 0.5166 | 0.1705 | 0.43736 | 0.21838 | 0.27541 | 0.63136 | |
| UTP | C00075 | I3sg5 | C13-02 | 9 | 0 | 0 | 18311.61629 | 0.175094 | 0.3033 | 1.7321 | 0 | 0 | 0.52528 | 0.17509 | 0.63136 | |
| UTP | C00075 | NT2 | C13-03 | 9 | 32879.07233 | 24739.5556 | 66046.67655 | 0.5204692 | 0.2787 | 0.5355 | 0.40375 | 0.3191 | 0.83856 | 0.52047 | 0.36097 | |
| UTP | C00075 | I3sg5 | C13-03 | 9 | 6150.161395 | 33310.87026 | 0 | 0.2542355 | 0.35 | 1.3768 | 0.10923 | 0.65348 | 0 | 0.25424 | 0.36097 | |
| UTP | C00075 | NT2 | C13-04 | 9 | 62760.54349 | 134198.514 | 13094.84204 | 0.8892936 | 0.7891 | 0.8873 | 0.77069 | 1.73094 | 0.16626 | 0.88929 | 0.50071 | |
| UTP | C00075 | I3sg5 | C13-04 | 9 | 64231.35328 | 104276.7608 | 27512.52944 | 1.3252143 | 0.6482 | 0.4891 | 1.14077 | 2.04565 | 0.78922 | 1.32521 | 0.50071 | |
| UTP | C00075 | NT2 | C13-05 | 9 | 4571041.606 | 4271432.64 | 4535825.507 | 56.271596 | 1.2532 | 0.0223 | 56.1314 | 55.0943 | 57.589 | 56.2716 | 0.62891 | |
| UTP | C00075 | I3sg5 | C13-05 | 9 | 3219817.826 | 2808190.168 | 2037557.306 | 56.907907 | 1.6966 | 0.0298 | 57.1851 | 55.0898 | 58.4488 | 56.9079 | 0.62891 | |
| UTP | C00075 | NT2 | C13-06 | 9 | 532636.8567 | 688871.997 | 615134.8537 | 7.7453377 | 1.1737 | 0.1515 | 6.54067 | 8.8853 | 7.81005 | 7.74534 | 0.45242 | |

| | | | | | | | | | | | | | | | |
|---|---|---|---|---|---|---|---|---|---|---|---|---|---|---|---|
| UTP | C00075 | I3sg5 | C13-06 | 9 | 321193.5581 | 326214.9716 | 295009.1287 | 6.8555331 | 1.4344 | 0.2092 | 5.70451 | 6.39954 | 8.46255 | 6.85553 | 0.45242 |
| UTP | C00075 | NT2 | C13-07 | 9 | 1049026.972 | 1036008.149 | 889911.7487 | 12.51445 | 1.0799 | 0.0863 | 12.8818 | 13.3628 | 11.2988 | 12.5144 | 0.47557 |
| UTP | C00075 | I3sg5 | C13-07 | 9 | 772920.4697 | 730726.7989 | 411526.9955 | 13.28912 | 1.3208 | 0.0994 | 13.7273 | 14.3351 | 11.8049 | 13.2891 | 0.47557 |
| UTP | C00075 | NT2 | C13-08 | 9 | 690455.0006 | 517742.0606 | 533301.0718 | 7.3092318 | 1.0138 | 0.1387 | 8.47864 | 6.67801 | 6.77105 | 7.30923 | 0.43044 |
| UTP | C00075 | I3sg5 | C13-08 | 9 | 447462.367 | 388071.153 | 276529.5859 | 7.8308488 | 0.1888 | 0.0241 | 7.94709 | 7.613 | 7.93245 | 7.83085 | 0.43044 |
