## Supplementary Table 2 for "Metabolic regulation of RNA methylation by the m^6^A-reader IGF2BP3"

| NAME | SEQUENCE |
|---|---|
| MAT2A_F | gaccagggcttaatgtttggc |
| MAT2A_R | tagaatcagggcgtaaccaagg |
| MAT2B_F | gagaacaatctaggagctgctg |
| MAT2B_R | tgccagtgatccatgtttgc |
| MTHFR_F | atctgtgtggcaggttaccc |
| MTHFR_R | aagcggaagaatgtgtcagc |
| PKM_F | tgtaccattggcccagcttc |
| PKM_R | tgtgcgcacattcttgatgg |
| PSAT1_F | tccattccgcattggcaatg |
| PSAT1_R | atgcctcccacagacctatg |
| SHMT1_F | agtcacaggtggttctgacaac |
| SHMT1_R | aacaggcttctagcaccttctc |
