## Supplementary Table 3 for "Metabolic regulation of RNA methylation by the m^6^A-reader IGF2BP3"

| Pathway | Proteins | Catalogue number | Vendor |
|---|---|---|---|
| Glycolysis | PKM2 | 15822-1-AP | Proteintech |
| Serine/Glycine Biosynthesis | PHGDH | PA5-80896 | Thermo Fisher |
| | PSAT1 | PA5-121028 | Thermo Fisher |
| | SHMT1 | 30192-1-AP | Proteintech |
| | SHMT2 | 11099-1-AP | Proteintech |
| METHYL | MTHFS | PA5-118092 | Thermo Fisher |
| | MTHFR | PA5-140282 | Thermo Fisher |
| | MAT2A | 55309-1-AP | Proteintech |
| | MAT2B | 15952-1-AP | Proteintech |
| RNA Methylation | METTL3 | 15073-1-AP | Proteintech |
| | METTL14 | 26158-1-AP | Proteintech |
| | METTL16 | 19924-1-AP | Proteintech |
| | FTO | 27226-1-AP | Proteintech |
| | $m^6A$ | MABE-1006 | Sigma Aldrich |
| Histone Methylation | Histone H3 | D1H2 mAb 4499 | Thermo Fisher |
| | H3K4ME3 | MA511199 | Thermo Fisher |
| | H3K4me1 | 710795 | Thermo Fisher |
| Regulatory | IGF2BP3 | 14642-1-AP | Proteintech |
| | ACTIN | A5441 | Sigma Aldrich |
| | VINCULIN | sc-73614 | Santa Cruz |
| | Puromycin | MABE343, clone12D1 | Sigma Aldrich |

| Antibody | Catalogue | Vendor |
|---|---|---|
| APC anti-mouse CD3ε Antibody | 100311 | Biolegend |
| APC/Cyanine7 anti-mouse CD117 (c-kit) Antibody | 105826 | Biolegend |
| PE/Cyanine7 anti-mouse/human CD11b Antibody | 101215 | Biolegend |
| PerCP/Cyanine5.5 anti-mouse Ly-6A/E (Sca-1) Antibody | 108124 | Biolegend |
| BD Horizon™ BV786 Mouse Anti-Mouse CD45.2 | 563686 | BD Biosciences |
| Biotin anti-mouse CD4 Antibody 500 ug | 100404 | Biolegend |
| Biotin anti-mouse TCR γ/δ Antibody 50 ug | 118103 | Biolegend |
| Biotin anti-mouse/human CD11b Antibody | 101203 | Biolegend |
| IgM Monoclonal Antibody (II/41), Biotin, eBioscience™, Invitrogen™ | 13-579-082 | Fisher |
| Biotin anti-human CD45 Antibody | 368534 | Biolegend |
| PE/Cyanine7 anti-human CD19 Antibody | 302215 | Biolegend |
| PerCP/Cyanine5.5 anti-human CD34 Antibody | 343521 | Biolegend |
| Biotin anti-mouse/human CD45R/B220 Antibody, 500 ug | 103204 | Biolegend |
| Biotin anti-mouse Ly-6G/Ly-6C (Gr-1) Antibody, 500ug | 108404 | Biolegend |
| Biotin anti-mouse NK-1.1 Antibody, 500 ug | 108704 | Biolegend |
| Biotin anti-mouse TER-119/Erythroid Cells Antibody, 500ug | 116204 | Biolegend |
| Biotin anti-mouse TCR β chain Antibody, 500ug | 109204 | Biolegend |
| Biotin anti-mouse CD8a Antibody, 500ug | 100704 | Biolegend |
| PE/Cyanine7 anti-mouse CD16/32 Antibody, 100ug | 101318 | Biolegend |
| eBioscience™ Streptavidin eFluor™ 450 Conjugate | 48-4317-82 | LifeTech |
| CD34 Monoclonal Antibody (RAM34), Alexa Fluor 700, eBioscience™, 100ug | 56-0341-82 | LifeTech |
| BD Horizon™ BV605 Mouse Anti-Mouse CD45.2 | 563051 | BD Biosciences |
