## Supplementary Images for "Metabolic regulation of RNA methylation by the m^6^A-reader IGF2BP3"

### Supplementary Figure 1

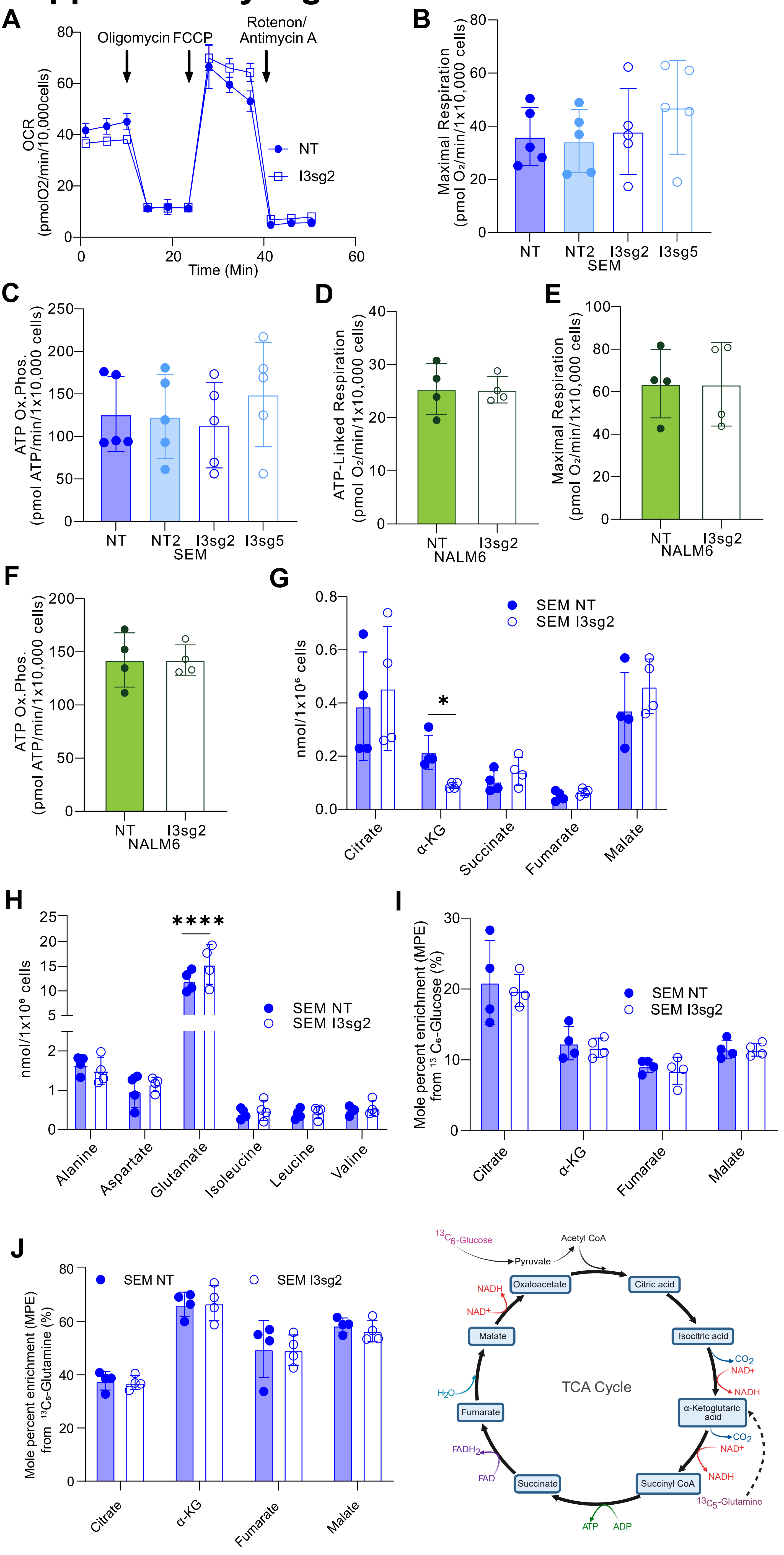

### Supplementary Figure 2

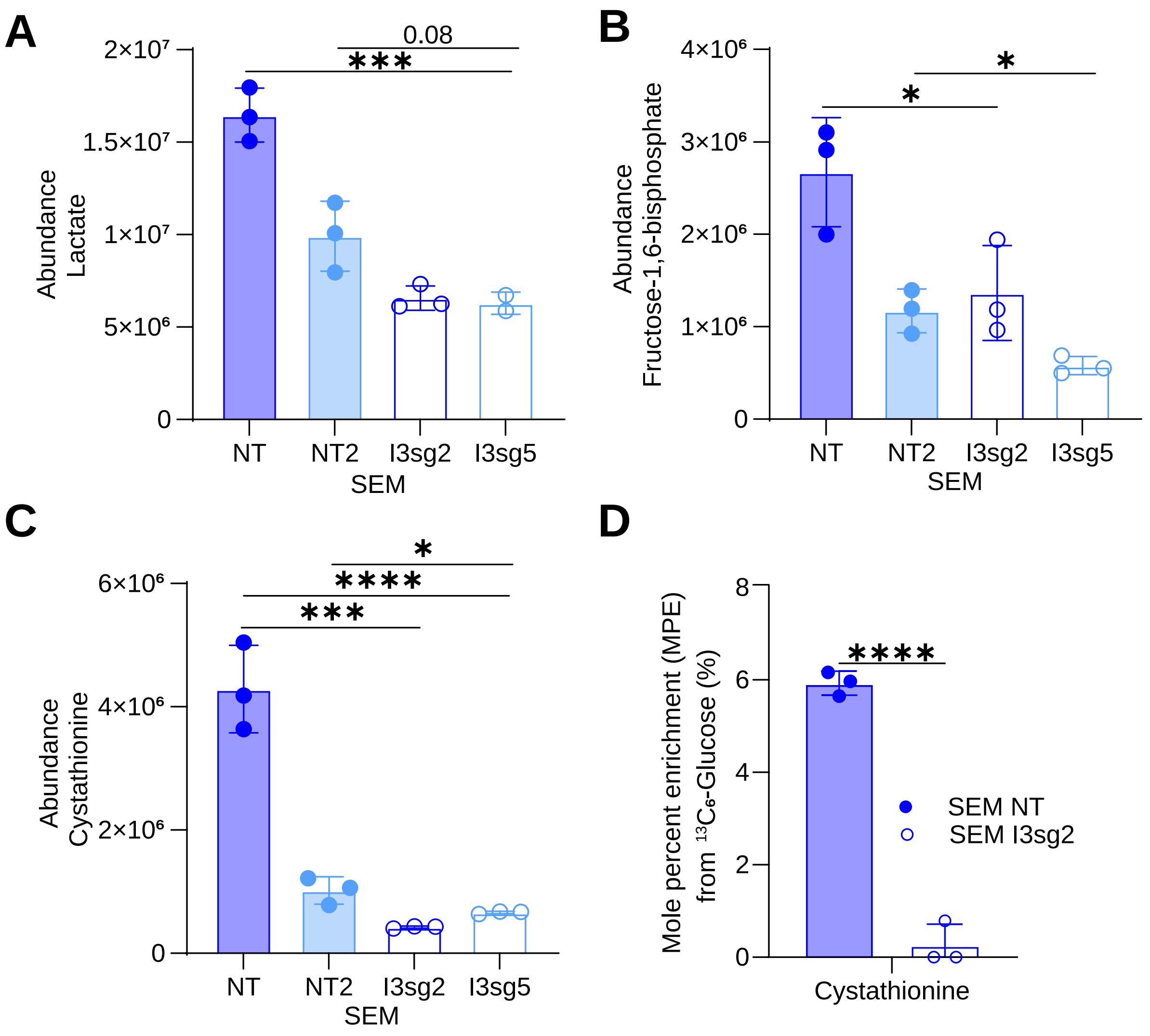

### Supplementary Figure 3

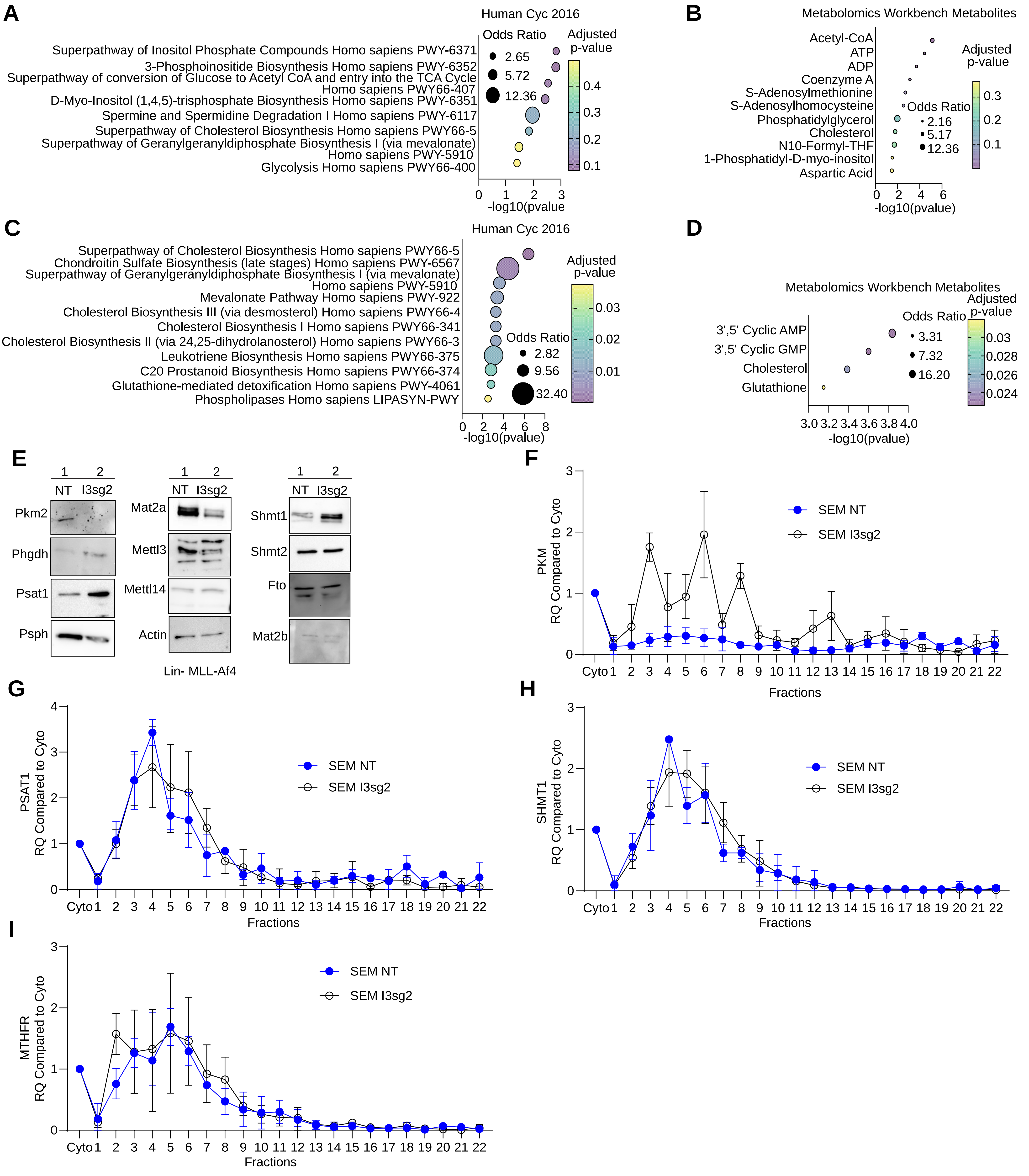

### Supplementary Figure 4

A

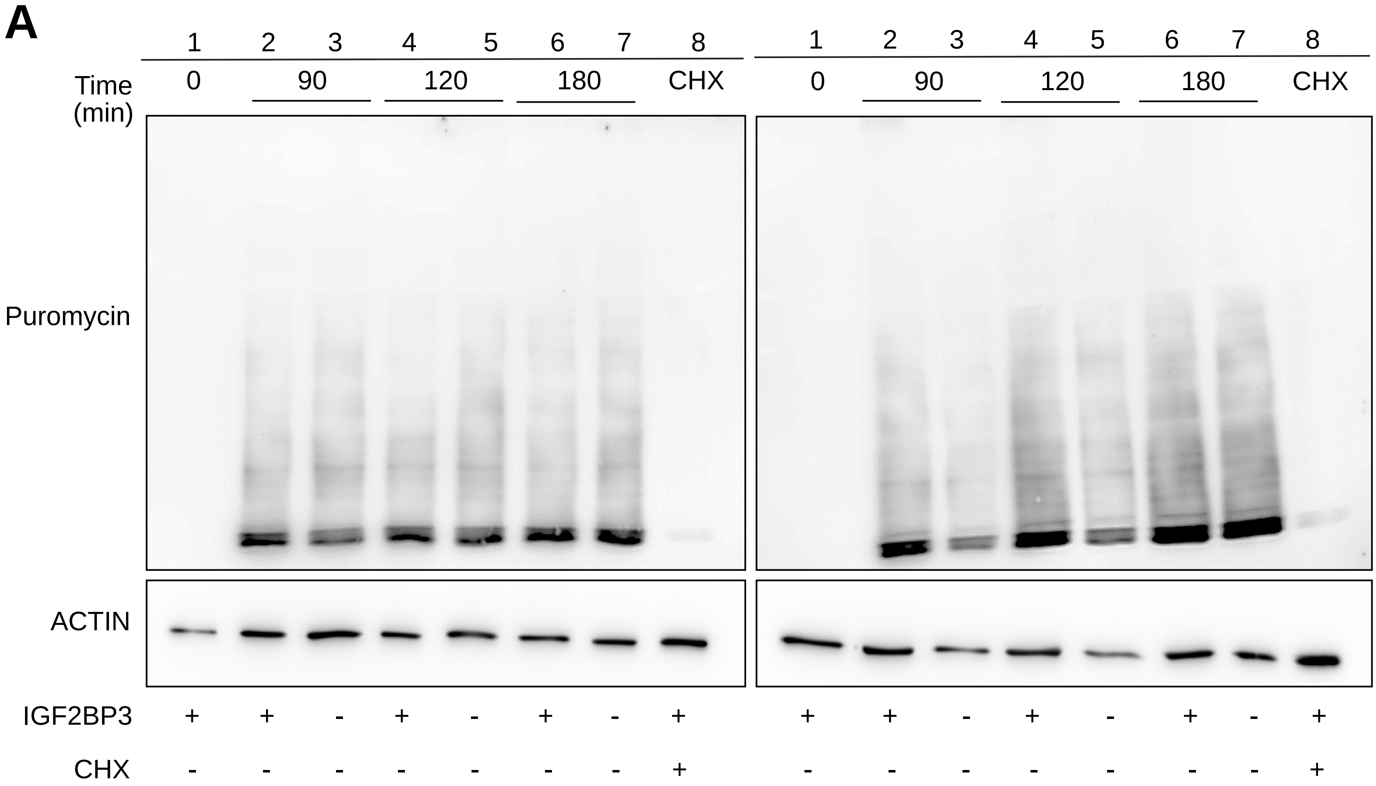


B

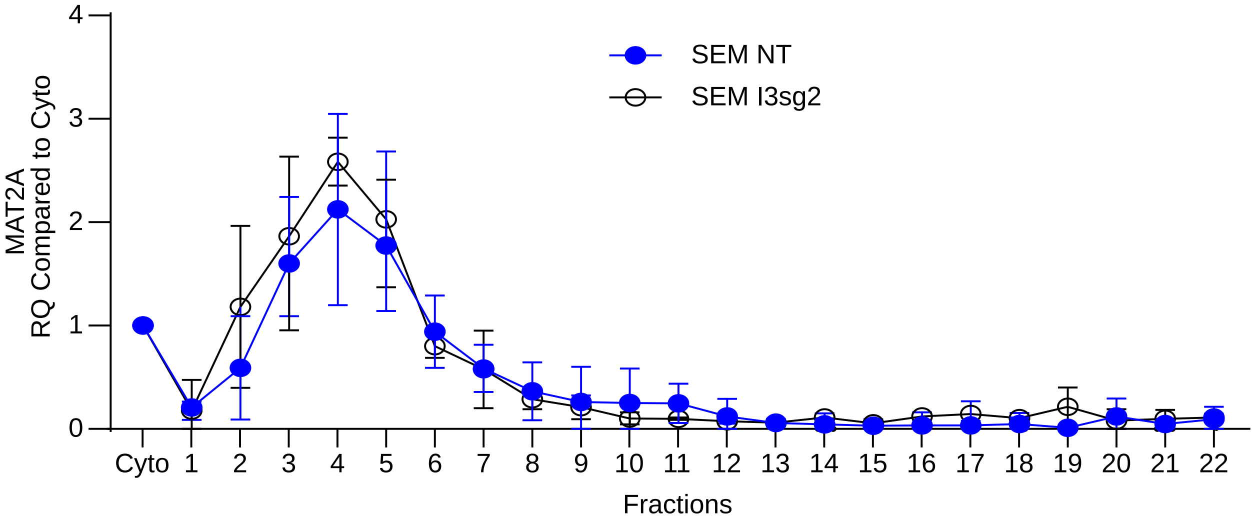


C

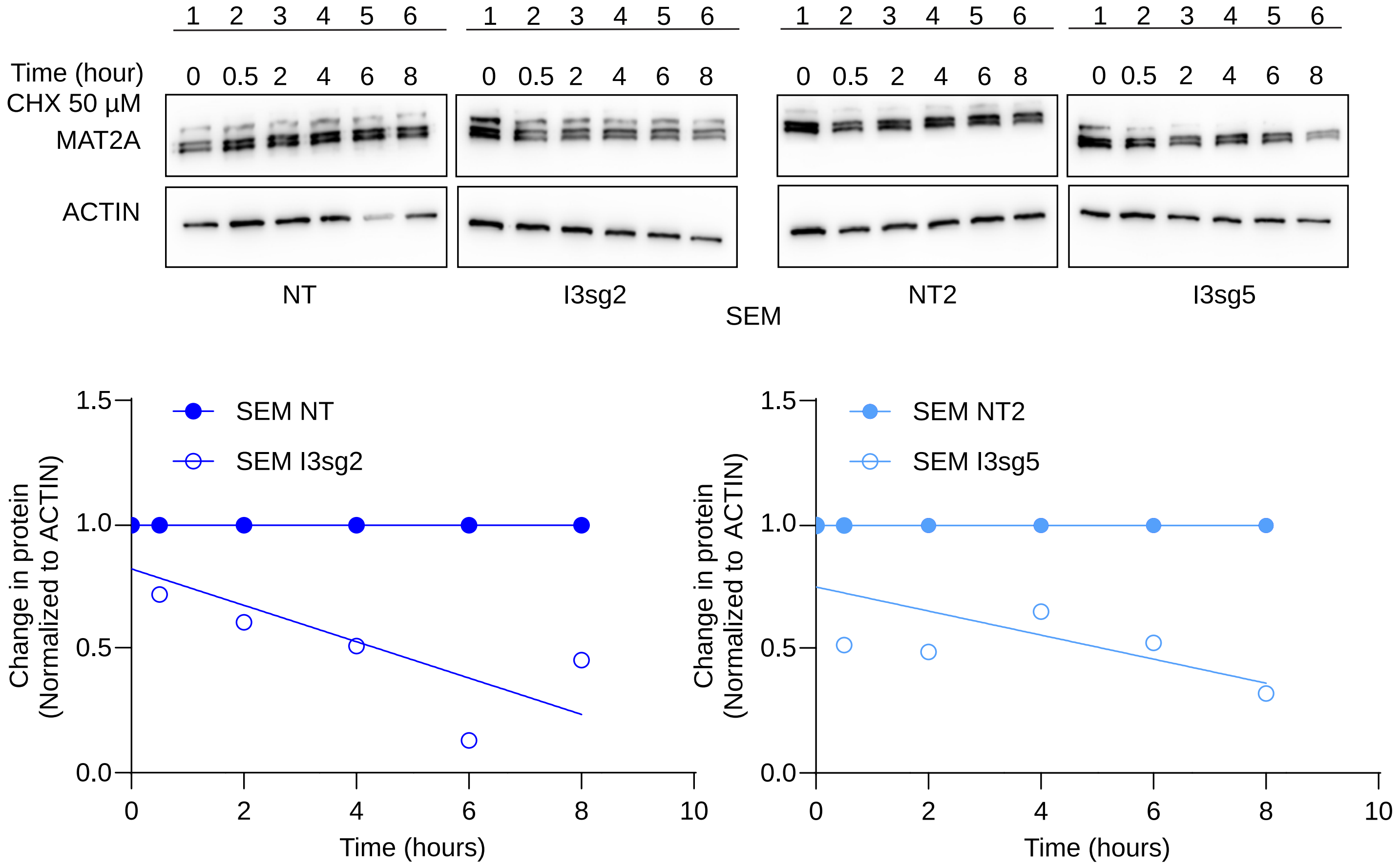

### Supplementary Figure 5

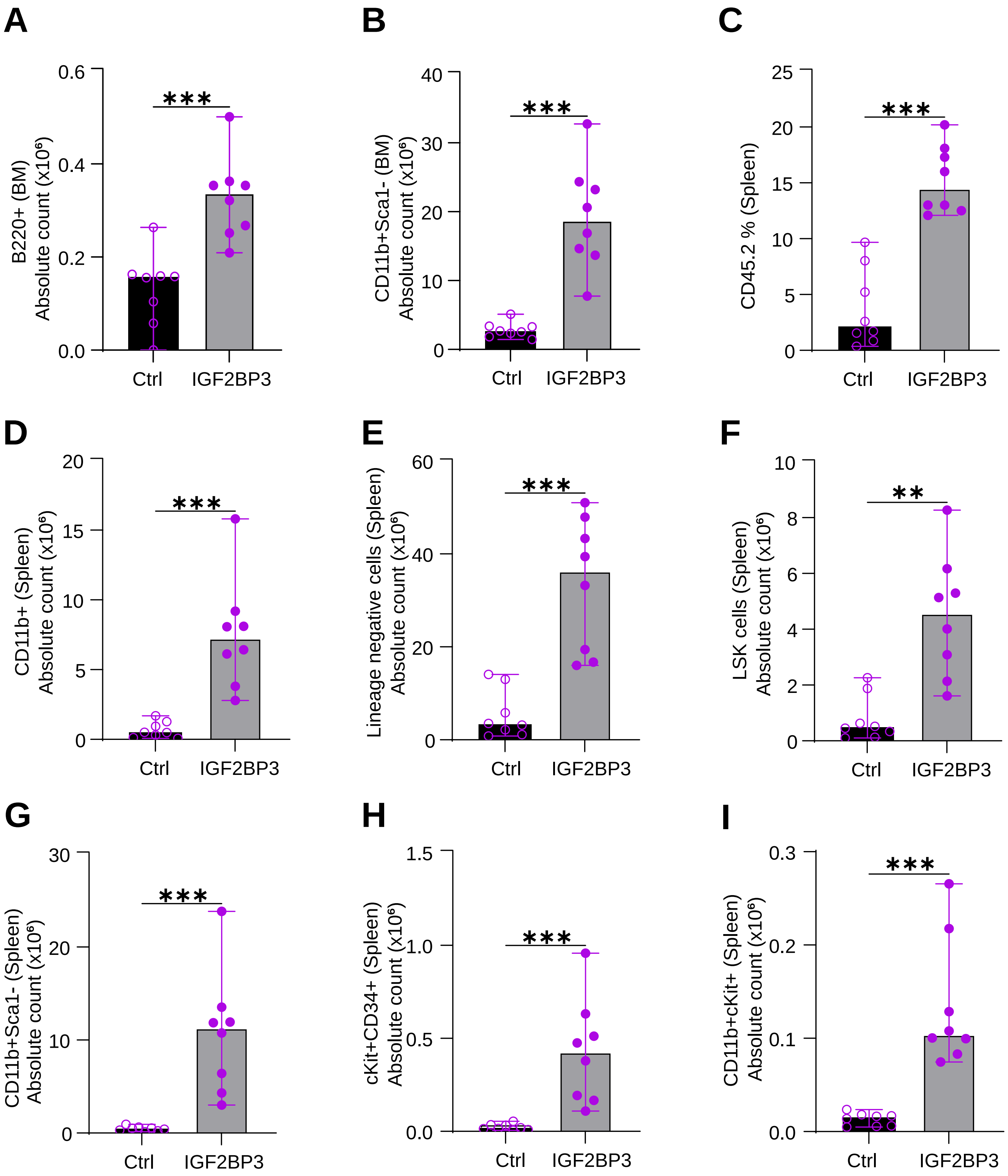
